# Detection of Horizontal Gene Transfer in the Genome of the Choanoflagellate *Salpingoeca rosetta*

**DOI:** 10.1101/2020.06.28.176636

**Authors:** Danielle M. Matriano, Rosanna A. Alegado, Cecilia Conaco

**Affiliations:** Marine Science Institute, University of the Philippines, Diliman; Department of Oceanography, Hawai’i Sea Grant, Daniel K. Inouye Center for Microbial Oceanography: Research and Education, University of Hawai’i at Manoa

**Keywords:** Lateral Gene Transfer, Evolution, Multicellularity

## Abstract

Horizontal gene transfer (HGT), the movement of heritable materials between distantly related organisms, is crucial in eukaryotic evolution. However, the scale of HGT in choanoflagellates, the closest unicellular relatives of metazoans, and its possible roles in the evolution of animal multicellularity remains unexplored. We identified 703 potential HGTs in the *S. rosetta* genome using sequence-based tests. The majority of which were orthologous to bacterial lineages, yet displayed genomic features consistent with the rest of the *S. rosetta* genome – evidence of ancient acquisition events. Putative functions include enzymes involved in amino acid and carbohydrate metabolism, cell signaling, the synthesis of extracellular matrix components, and the detection of bacterial compounds. Functions of candidate HGTs may have contributed to the ability of choanoflagellates to assimilate novel metabolites, thereby supporting adaptation, survival in diverse ecological niches, and response to external cues that are possibly critical in the evolution of multicellularity in choanoflagellates.

## Background

In Bacteria and Archaea, several mechanisms, such as transformation, conjugation, and transduction (1–8), facilitate the introduction of novel genes between neighboring strains and species, known as horizontal (or lateral) gene transfer (HGT) (9,10). Horizontal transfers into eukaryote genomes, however, are thought to be low frequency events, as genetic material must enter the recipient cell’s nucleus to be incorporated into the genome and vertically transmitted (11). While HGT has been difficult to demonstrate in eukaryotic lineages (12), conflicting branching patterns between individual gene histories and species phylogenies (4) has garnered a number of explanations: a gene transfer ratchet to fix prey-derived genes from prey species (“you are what you eat”) (13), movement of DNA from organelles to the nucleus through endosymbiosis (14,15), the presence of “weak-links” or unprotected windows during unicellular or early developmental stages that may enable integration of foreign DNA into eukaryotic genomes (11), and horizontal transposon transfers (16–18). Indeed, HGT may have accelerated genome evolution and innovation in microbial eukaryotes by contributing to species divergence (19–21), metabolic diversity and versatility (22,23), and the establishment of interkingdom genetic exchange (10,19,24,25).

Recently, Choanoflagellatea, a clade of aquatic, free-living, heterotrophic nanoflagellates (26,27), has emerged as a model for understanding the unicellular origins of animals. Choanoflagellates phagocytose detritus and a diverse array of microorganisms and are globally distributed in marine, brackish, and freshwater environments (28). All choanoflagellates have a life history stage characterized by an ovoid unicell capped by a single posterior flagellum that functions to propel the cell in the water column and to generate currents that sweep food items toward an actin microvilli collar (26,27,29). This cellular architecture bears striking similarity to sponge choanocytes (or “collar cells”), leading to speculation on the evolutionary relationship between choanoflagellates and animal (27). Abundant molecular phylogenetic evidence support choanoflagellates as the closest extant unicellular relatives of animals (30–35).

While comparative genomics have revealed a host of animal genes found in common with the Urchoanozoan (28), few studies have investigated the evolutionary history of genes involved in the origins of choanoflagellates and animals, including the possible acquisition of genes from other lineages. Choanoflagellates were historically grouped into two distinct orders based on taxonomy (28,32,33,36): Acanthoecida (families Stephanoecidae and Acanthoecidae), which produce a siliceous cage-like basket exoskeleton or lorica; and non-loricate Craspedida (families Codonosigidae and Salpingoecidae), which produce an organic extracellular sheath or theca (32,33). Previous studies suggest that silicate transport genes and transposable elements in loricate choanoflagellates may have been acquired from microalgal prey (16,37). At least 1,000 genes, including entire enzymatic pathways present in extant choanoflagellates, may have resulted from HGT events (38–46).

The colonial craspedid *Salpingoeca rosetta* has been established as a model organism for origins of animal multicellularity (27,47). In this study, we used sequence-based methods to identify putative HGT events in the *S. rosetta* genome. *S. rosetta* has a rich life history entwined with affiliated environmental bacteria (26,27,48–50). Transient differentiation from the solitary morphotype into chain or rosette colonies as well as sexual mating in choanoflagellates are triggered by distinct prey bacteria (48). We hypothesized that sources of novel horizontal transfers in *S. rosetta* are primarily from food sources - bacterial or microalgal donors consistent with the “you are what you eat” gene ratchet theory (13). We compared the unique gene signatures (i.e. GC content, codon usage bias, intron number) of potential HGTs against the rest of the *S. rosetta* genome to determine whether the genes were ancient or recent transfers. We also assessed the extent of taxonomic representation of the candidate HGTs in other choanoflagellates. Characterizing the magnitude and functions of HGTs in choanoflagellate genomes may provide insight into their role in the evolution and adaptation of the lineage to new ecological niches and lifestyles. Potential HGT events in choanoflagellates can provide unique insights into the evolutionary history of genes involved in the origins of choanoflagellates and animals.

## Results

### Identification and genic architecture of candidate HGTs in S. rosetta

Of the 11,731 reference *S. rosetta* protein coding genes, 4,527 (38.59%) and 4,391 (37.43%) were related to metazoan and *S. rosetta* sequences, respectively (Fig. 1A). Our taxon filter flagged 2,804 (23.90%) genes with highest orthology to sequences from prokaryotes and unicellular non-metazoan taxa. Of these, 1,837 (15.66%) genes had best hits to sequences from prokaryotes, fungi, and unicellular eukaryotic taxa with marine representatives, such as chlorophytes, rhodophytes, haptophytes, and the stramenopile, alveolate, and Rhizaria supergroup (SAR). Best hits from prokaryotes (i.e. bacteria and archaea) and unicellular eukaryotic taxa (i.e. unicellular algae and fungi) were scored as potential HGT candidates and subjected to further analysis.

**Fig. 1.**
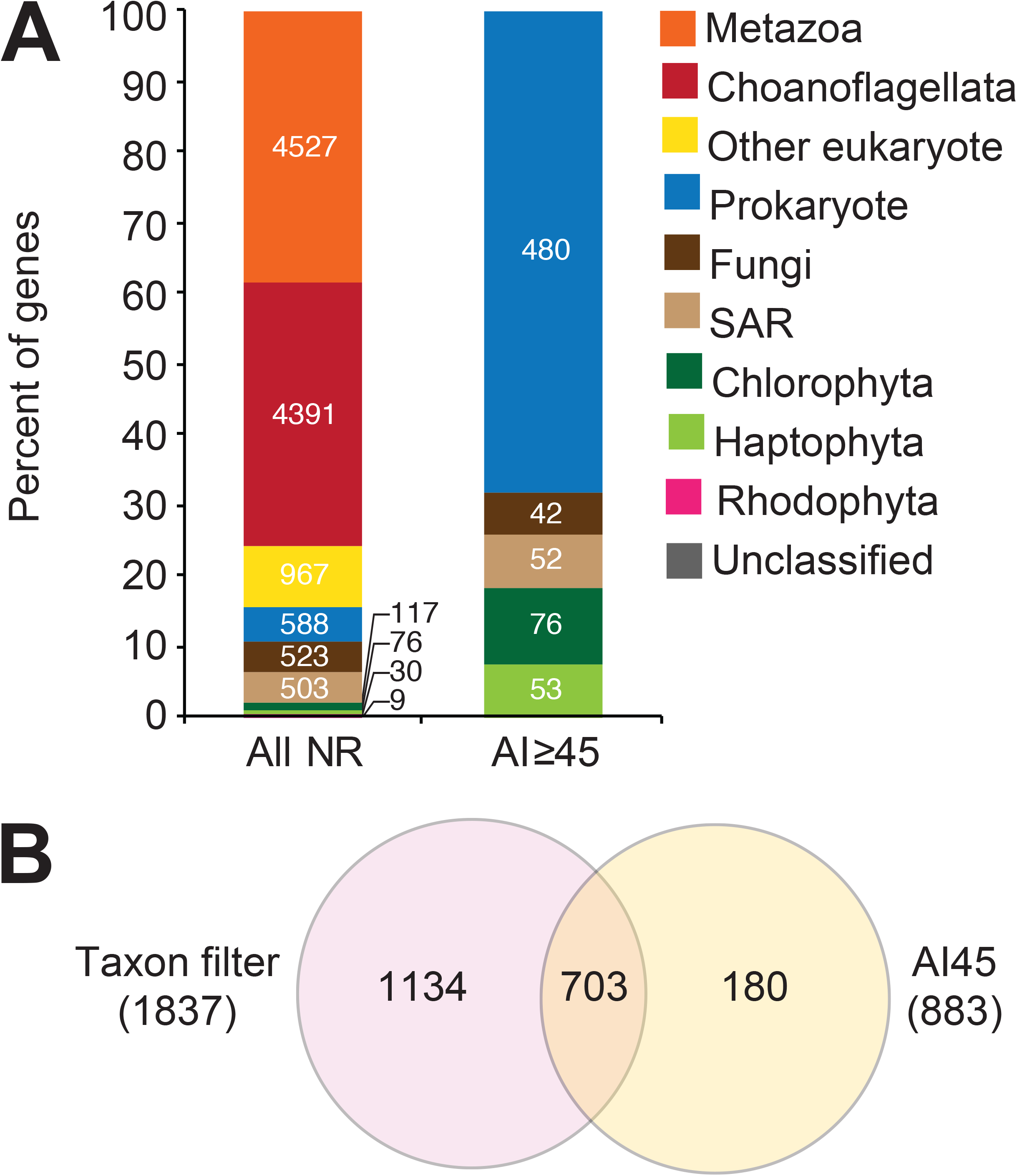
HGT candidates in the *S. rosetta* genome. (A) Number of putative HGTs detected using sequence alignment and Alien Index (AI45) analysis. (B) Taxonomic affiliation of the best sequence match for all *S. rosetta* genes and genes that passed the AI>45 filter.

### 1.2. Identification of candidate HGTs by Alien Index analysis

Using metrics described by Gladyshev et al. (2008), Alien Index (AI) analysis of the 1,837 potential HGT candidate genes identified 703 candidate HGTs from potential bacteria, archaea, and common unicellular marine fungi (i.e. Basidiomycota and Ascomycota) and eukaryotic (i.e. Chlorophyta, Haptophyta, Pelagophyta, and Bacillariophyta) donors (Fig. 1B, Table S1), which included only 38% of the genes flagged by sequence alignment to the NCBI nr database. A total of 111 (16%) of the potential HGTs in *S. rosetta* have orthologs to HGT candidates that were previously identified in the choanoflagellate, *Monosiga brevicollis* (40).

### 1.3. Genic architecture and gene expression of candidate HGTs

In *S. rosetta*, the difference in median protein coding sequence (CDS) length of candidate HGTs (1,509 bp) from the overall median CDS length (1,401 bp) was small but significant (Kruskal-Wallis test, *p* ≤ 0.001; Fig. 2A). Candidate HGTs also had significantly higher GC content at the third codon position (GC3) relative to other *S. rosetta* genes (Fig. 2B; Kruskal-Wallis test, *p* ≤ 0.001), indicating high degeneracy of these frequently recombining genes. However, candidate HGTs were observed to have a similar median number of introns, specifically five, as compared to other *S. rosetta* genes (Fig. 2C). Only 1,103 (8.79 %) of *S. rosetta* genes lacked introns, of which 73 (0.62%) passed the AI ≥ 45 threshold. Codon bias index (CBI), which is a measure of codon usage frequency, was also similar between the putative HGTs and other genes (Fig. 2D). In addition, 662 of the candidate HGTs were expressed in the four life stages of *S. rosetta* (i.e. thecate cells, swimming cells, chain colonies, and rosette colonies), 38 genes were expressed in at least one stage, and only 3 were not detected in any of the stages (Fig. 2E, Table S2), based on data from the study of Fairclough et al. (2013).

**Fig. 2.**
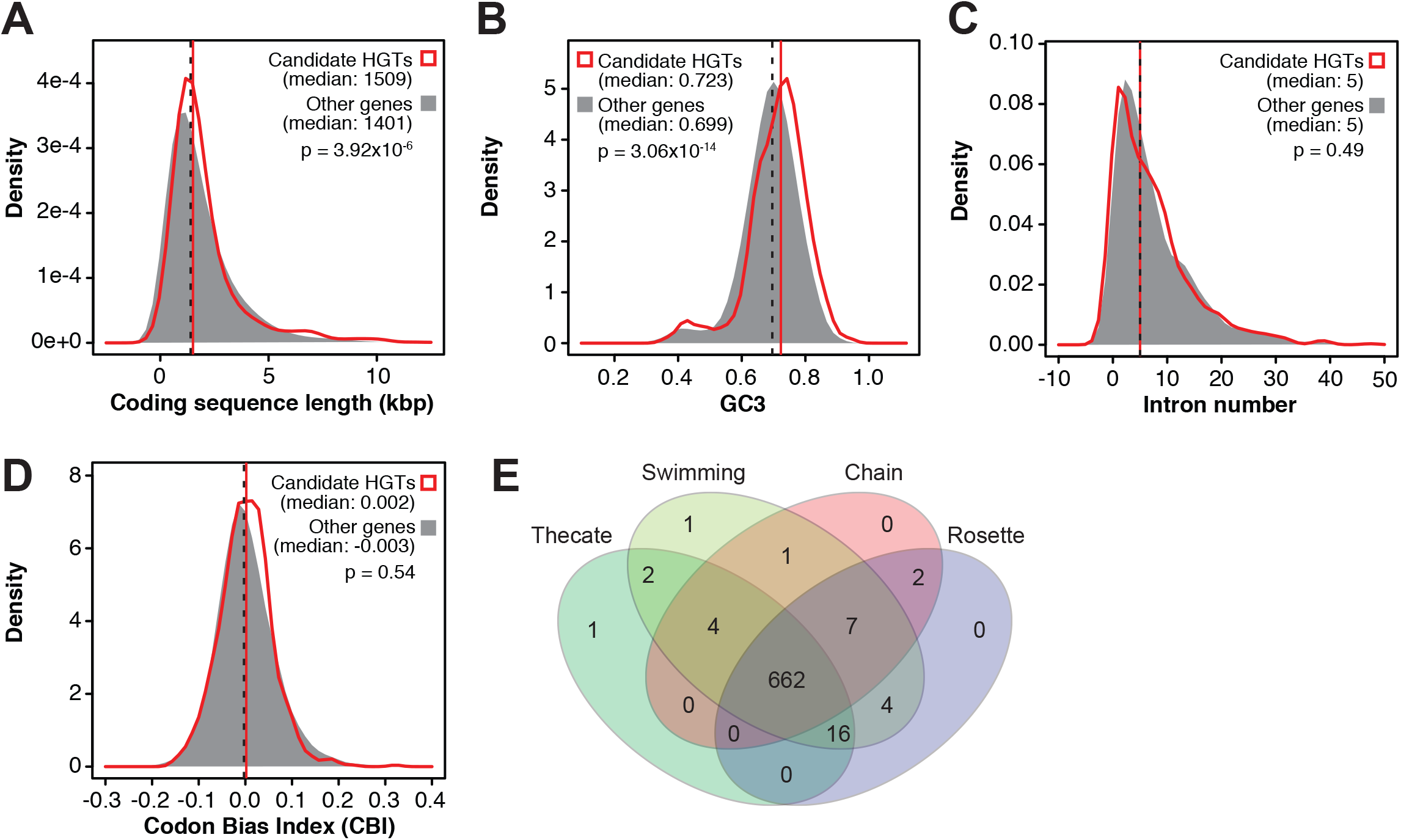
Gene architecture of candidate HGTs. Histograms showing the distribution of (A) coding sequence length (CDS), (B) GC content at the third codon position (GC3), (C) intron number, and (D) codon bias index (CBI) in candidate horizontally transferred genes (red) in comparison to the bulk of *S. rosetta* genes (gray). (E) Number of HGT candidates that are expressed in the indicated life stages of *S. rosetta*.

### Orthologs of candidate HGTs in other taxa

To estimate when horizontally transferred genes in the *S. rosetta* genome were acquired, we assessed the number of orthologs of candidate HGTs in 20 other choanoflagellates and representative eukaryotes, opisthokonts, filasterean, and metazoans based on OrthoMCL groups or ortholog families identified by the study of Richter et al. (2018) (Table S3). Majority of candidate HGTs (517 genes) had orthologs in other eukaryotes (Excavata, Diaphoretickes, and Amoebozoa), 9 in fungi, and 2 in filasterea, likely reflecting genes in donor taxa or gene transfers into older lineages. Thirty six genes had orthologs in animals while 139 were only found in choanoflagellates (Fig. 3A). Most candidate HGTs (443 genes) had orthologs in all choanoflagellate families and 87 were found only in Craspedida (clade 1), which includes *S. rosetta* (Fig. 3B).

**Fig. 3.**
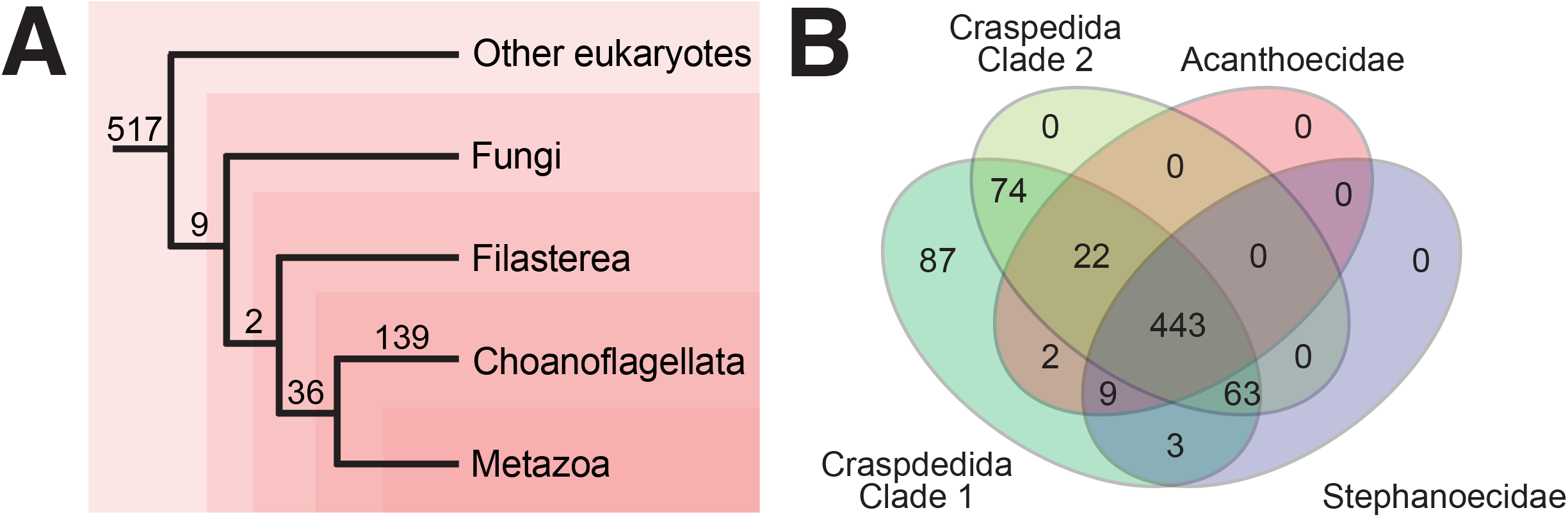
Conservation of candidate HGTs. (A) The number of HGT candidates with orthologs in other lineages. (B) The number of HGT candidates with orthologs in choanoflagellate taxa within orders Craspedida (clade 1 and 2) and Acanthoecida (family Acanthoecidae and Stephanoecidae).

### Potential sources of S. rosetta HGTs

The majority of the candidate HGTs exhibited highest similarity to bacteria (57%), Chlorophyta (15%), Haptophyta (10%), Stramenopiles (10%), and fungal (Ascomycota and Basidiomycota) (8%) sequences (Fig. 4A). Phylogenetic analysis of selected putative HGTs revealed conflicting phylogenetic signals relative to the consensus eukaryote reference phylogeny of Richter et al. (2018) (Fig. 4B-D, Additional File 1), confirming their possible origin from donor taxa. The most common potential bacterial donors were placed in one of the following phyla: Proteobacteria, Terrabacteria, Fibrobacteres, Chlorobi, and Bacteroidetes (FCB) group, and the Planctomycetes, Verrucomicrobia, and Chlamydiae (PVC) group (Fig. 4A). At least 28 sequences may have been derived from marine microorganisms that *S. rosetta* potentially interacts with, including known food sources, such as *Algoriphagus machipongonensis* (Bacteroidetes) (9 genes) and *Vibrio* sp. (Proteobacteria) (13 genes) (Table S4). Other notable potential donors included the Bacteroidetes *Cytophaga* sp. and *Flectobacillus* sp, both of which have been shown to induce colony formation in *S. rosetta* (48).

**Fig. 4.**
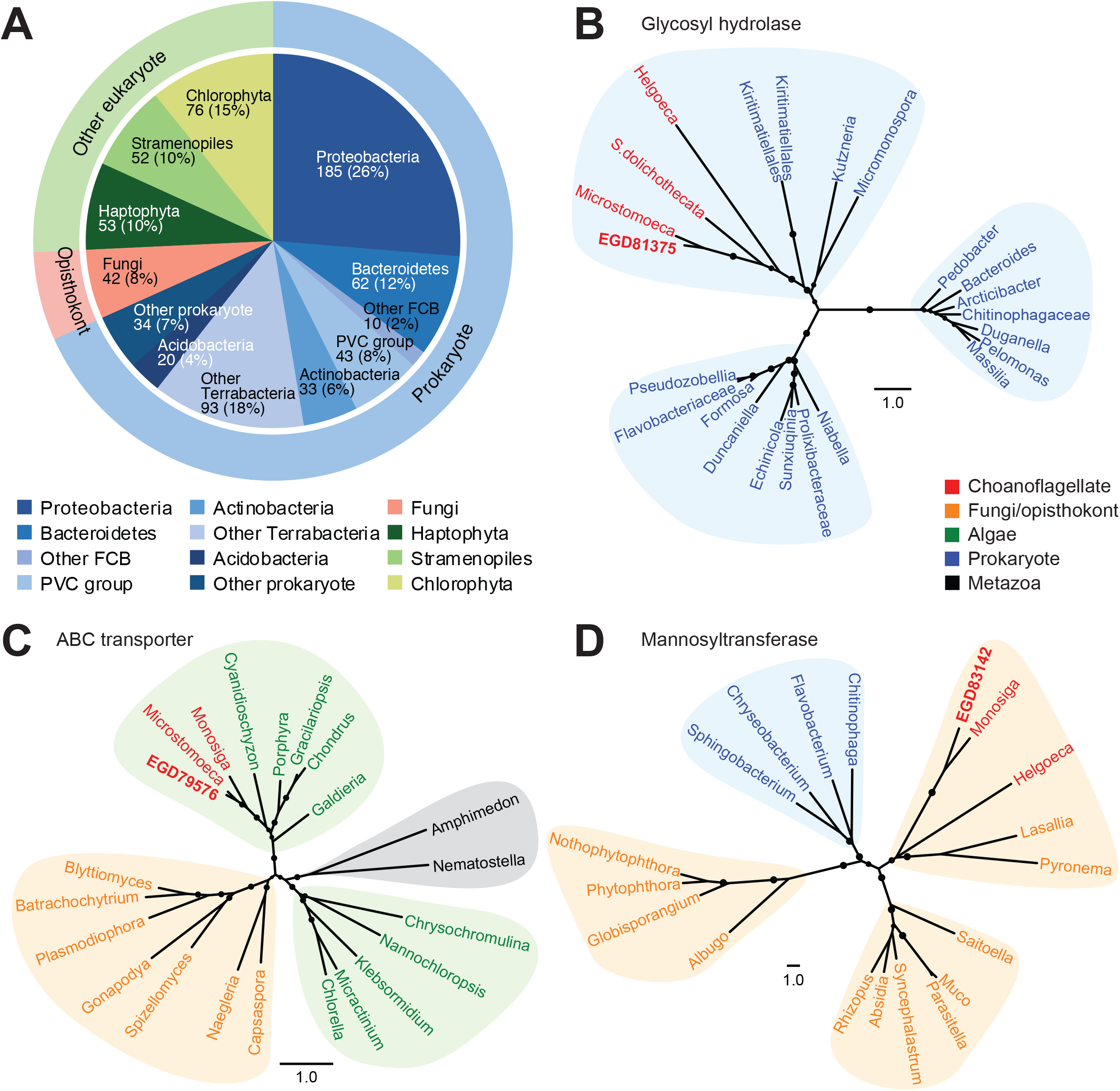
Potential donor phyla of candidate HGTs in *S. rosetta*. (A) Taxon affiliation of candidate HGTs based on the best sequence match for each gene. Bayesian analysis of selected candidate HGTs, including a (B) glycosyl hydrolase of prokaryotic origin, (C) ABC transporter of algal origin, and (D) mannosyltransferase of fungal origin. *S. rosetta* genes are indicated in bold red letters. Genes from other choanoflagellates are shown in red, algae in green, fungi or opisthokonts in orange, prokaryotes in blue, and metazoans in black. Circles at the branches indicate posterior probabilities of 0.70-1.00.

### Functions of candidate HGTs in S. rosetta

Of the 703 candidate HGTs in *S. rosetta*, 595 (85%) had identifiable PFAM domains and 494 (70%) were assigned gene ontology (GO) annotations. At least 539 (76.7%) candidate HGTs contained more than two annotated PFAM domains. The most notable protein domains represented in the set of putative HGTs included enzymes (i.e. protein kinase, short chain dehydrogenase, aminotransferase, ubiquitin-activating enzyme active site, bacterial pre-peptidase C-terminal, sulfatases, hydrolases, oxidoreductases, guanyl cyclase, serine carboxypeptidase, and methyltransferase), transmembrane transporters (i.e. mitochondrial carrier protein, ATP-binding cassette (ABC) transporter, major facilitator superfamily, sugar transporter), ECM-associated domains (i.e. dermatopontin/calcium-binding EGF domain), and signaling domains (i.e. protein kinase, immunoglobin-like, plexins, transcription factor (IPT/TIG)) (Fig. 5A, Table S5).

**Fig. 5.**
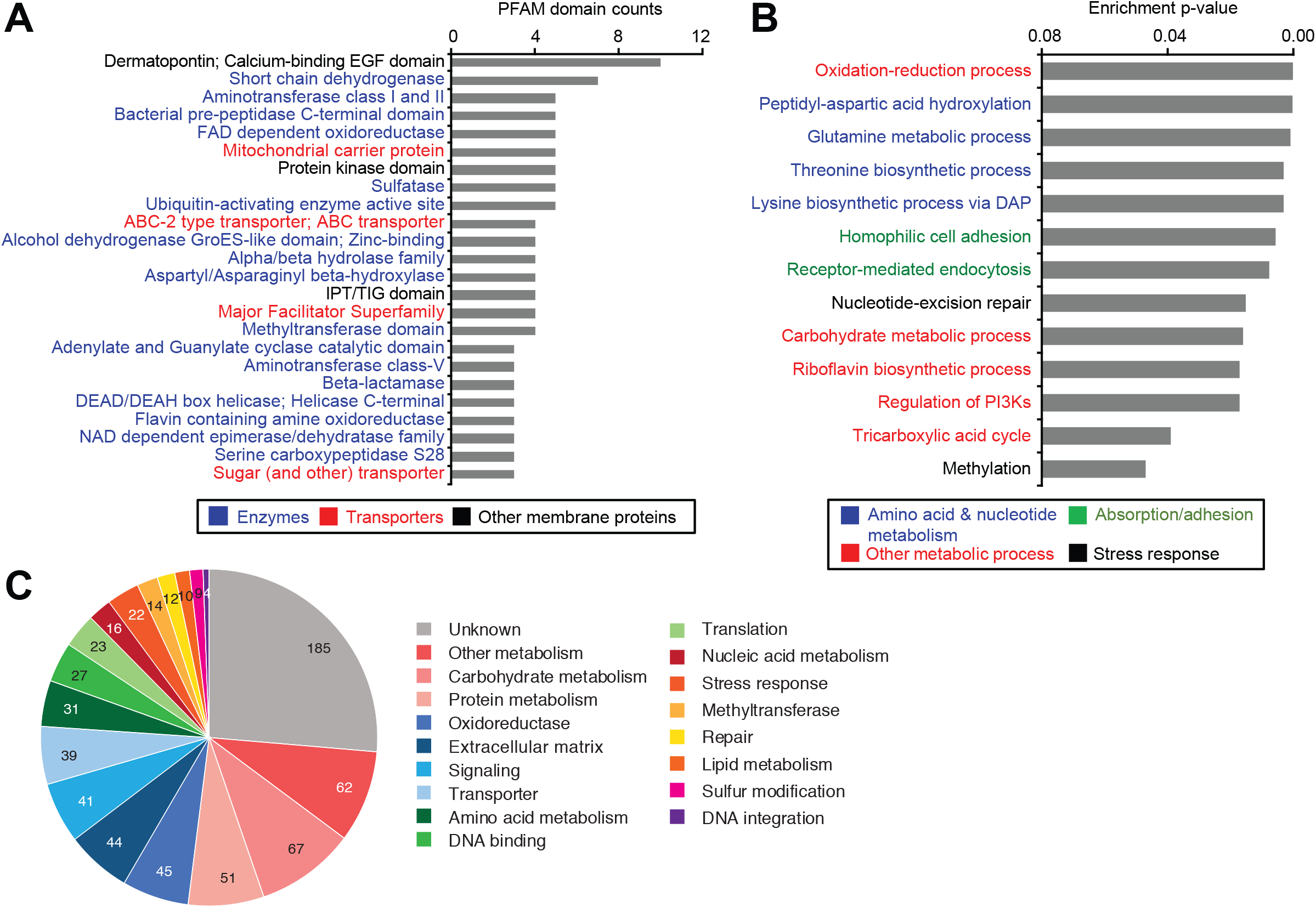
Functional analysis of candidate HGTs. (A) Most common PFAM protein domains in the set of candidate HGTs. (B) Gene ontology functions enriched in the set of putative HGTs (cellular component, CC; molecular function, MF; biological process, BP). Enrichment p-values (*p* ≤ 0.05) for selected functions are shown. Text colors in A and B indicate general functional groupings. (C) Number of candidate HGTs with associated functions based on manual curation.

Gene ontology analysis further supported the finding that majority of HGT candidates were enzymes with a variety of catalytic activities and were associated with cellular membranes where most biosynthetic and energy transduction processes of the cell occur (51) (Fig. 5B, Table S6). More specifically, potential HGTs in *S. rosetta* were associated with functions related to carbohydrate, protein, and lipid metabolic processes. The most common enriched GO terms were related to amino acid metabolism, including peptidyl-aspartic acid hydroxylation, glutamine metabolic process, threonine biosynthetic process, and lysine biosynthetic process via diaminopimelate (DAP). Other enriched functions included oxidation-reduction processes, carbohydrate metabolism, tricarboxylic acid cycle, and methylation. Homophilic cell adhesion, receptor-mediated endocytosis, nucleotide-excision repair, and regulation of phosphatidylinositol 3-kinase (PI3K) signal transduction pathway were also enriched.

### Selected candidate HGTs

To further examine the potential functions of candidate HGTs, the genes were manually curated and assigned to a general cellular function based on GO affiliations and PFAM domains. The most common functions within the set of putative HGTs were general metabolism, metabolism of carbohydrates, proteins, and lipids, oxidoreduction, extracellular matrix components, signaling, and transport (Fig. 5C, Table S7).

Candidate HGTs included a diverse array of transporter families, including ABC transporters, sugar transporters, major facilitator superfamily (MFS), and mitochondrial carrier proteins, among others. Putative HGTs that function in the catabolism or modification of proteins included various families of peptidases. Carbohydrate metabolism related HGTs included 22 glycosyl hydrolase domains, 6 glycosyltransferases, and 3 mannosyltransferases.

Candidate HGTs involved in lysine biosynthesis through the diaminopimelic acid (DAP) pathway included aspartate-semialdehyde dehydrogenase (*asd*), diaminopimelate decarboxylase and aspartate kinase (*lysAC*), and diaminopimelate epimerase (*dapF*). Other enzymes involved in the biosynthesis of histidine, threonine, methionine, cysteine, tryptophan, and arginine were also detected as potential HGTs (Additional file 2). Candidate HGTs related to the metabolism of amino acid is common in other choanoflagellate representatives, sponges, and cnidarians.

Other interesting candidate HGTs encoded stress response-related enzymes, including glutathione S-transferases, peroxidases, heat shock proteins, and thioredoxins. In addition, we identified HGT candidates associated with DNA repair including helicase, ligases, and ribonucleases. Beta-lactamase domain-containing genes that were potentially acquired from bacteria were also retained in the *S. rosetta* genome.

Several ECM-associated genes were identified as potential HGTs, including dermatopontin/calcium-binding EGF (DPT/Ca^2+^ EGF) domain-containing genes. These genes have orthologs in craspedids but all have no orthologs in loricates. Genes containing DPT/Ca^2+^ EGF domains had no orthologs in representative filastereans but were regained in higher animals, including cnidarians and bilaterians.

Two chondroitin sulfate AC lyase genes were detected as potential HGTs. One gene (EGD79853) has orthologs in Craspedida, Acanthoecidae, and Stephanoecidae, yet is absent in *Hartaetosiga balthica* and *Hartaetosiga gracilis.* This gene is conserved in filastereans and sponges. The other chondroitin sulfate AC lyase (EGD79387) gene, however, has no orthologs in other representative filastereans and metazoans.

## Discussion

### Horizontal gene transfer in S. rosetta

Our analysis provides evidence of a rich repertoire of *S. rosetta* genes that may have been acquired through horizontal gene transfer. Recent gene acquisitions are usually distinguished by divergent genetic characteristics, such as GC content, codon usage bias, and genetic architecture (e.g. intron content, coding sequence length) (52,53). Over time, transferred genes undergo sequence changes to adapt to host genome characteristics, enabling improved transcription and translation (54). A majority of *S. rosetta* HGT candidates had gene features similar to the rest of the genome and were expressed, based on the study by Fairclough et al. (2013), indicating that these acquisitions were not recent (34). We noted a higher GC content at the third codon position of the HGT candidate genes, which is a typical marker for genes derived from bacterial sources (55,56), yet the genes did not exhibit a divergent intron count. It is possible that foreign genes with translationally optimal codons and high GC3 content that resembles the GC-rich genome of *S. rosetta* are more likely to be positively selected, as was observed for some horizontally acquired transposable elements (TEs) in this species (16). The prevalence of introns in *S. rosetta* candidate HGTs further indicate adaptation of acquired genes to the intron-rich genome of *S. rosetta*. Moreover, most HGT candidates had orthologs in other choanoflagellate taxa, suggesting that the gene transfer events occurred before divergence of the various choanoflagellate groups. Nevertheless, phylogenetic analysis of the candidate HGTs revealed incongruent gene trees, with candidate HGTs clustering with genes from potential donors, including bacteria, microalgae, and fungi. This suggests that candidate horizontally transferred genes in *S. rosetta* were acquired from multiple prokaryotic and unicellular eukaryotic donors.

Most of the potential donors of HGTs in *S. rosetta* clustered with potential bacterial donors, in contrast to *M. brevicollis* where HGTs were identified as coming mostly from algal donors (40). *S. rosetta* is an active phagotroph of bacteria, such as *A. machipongonensis* and *Vibrio* spp., which have also been shown to influence its development and metabolic processes (26,27,48,50). Thus, the major mechanism of HGT in *S. rosetta* may be through the engulfing of food or associated microbes. The acquisition of genes through ingestion would have also allowed for the transfer of more diverse genetic functions as opposed to a more selective one via endosymbiosis. The predation of bacteria corroborate with the “you are what you eat” gene transfer ratchet theory, which suggests that the evolution of the nuclear genome of most protists was driven by acquisition of exogenous genes by phagotrophy or engulfment and acquisition of gene fragments from their food sources (13). HGT events may also be facilitated by the activity of TEs, which are abundant in the genome of *S. rosetta* (16). However, it should be noted that because some taxonomic lineages are underrepresented in sequence databases and since some HGTs may have been integrated into the host genome for a long time, the identification of the donor species for HGTs is not straightforward.

### Functions of HGTs in S. rosetta

Most candidate HGTs in *S. rosetta* were orthologous to operational genes, such as enzymes that function in amino acid, carbohydrate, and lipid biosynthesis, as well as genes that function in intercellular signaling and in the establishment and modification of ECM components. Based on the complexity hypothesis proposed by Jain et al. (1999), gene transferability is dependent on two factors: gene function and protein-protein interaction/network interaction (22,23,57–63). Operational genes are more likely to be passed horizontally because they can function independently of other genes (40,64–69). On the other hand, informational genes or genes involved in transcription and translational processes, physically interact with more gene products, limiting their functionality when transferred individually and reducing the possibility that they will be successfully retained as HGTs (58).

The acquisition of genes from a vast gene pool enhances genetic variability in free-living organisms. This variability may have conferred upon *S. rosetta* the ability to explore and establish new niches and adapt to various environmental conditions by mediating interactions with other organisms in the environment and facilitating life stage transitions. The contribution of novel functions acquired via HGT may be higher in choanoflagellates like *M. brevicollis* and *S. rosetta* that have retained fewer ancestral gene families compared to other choanoflagellates (70).

Novel combinations of protein domains could potentially enhance catalytic efficiency and functional novelty of enzymes. The genome of *S. rosetta* contains a rich complement of multidomain genes that may have contributed to its unique biology and morphology (31,34). It is possible that some of these molecular innovations emerged through gene fusion or domain shuffling events that incorporated pre-existing domains with domains acquired from horizontally transferred genes (71).

### Candidate HGTs potentially contribute to metabolism and defense

Many of the putative HGTs in *S. rosetta* contribute to nutrient acquisition and metabolic processes by enabling efficient use of available organic substrates. Diverse transporters facilitate uptake and transport of the building blocks for metabolic pathways (i.e. sugars, phosphates, and amino acids) or the excretion of biomolecules and toxins (40,72,73). Proteases and glycosyl hydrolases break down proteins and complex sugars (74–76). Glycosyl hydrolases, in particular, break down complex sugars and proteoglycans (i.e. heparan sulfate proteoglycans, chondroitin sulfate proteoglycans, and hyaluronans) (77), as well as plant materials and matrix polysaccharides of biofilms (74,75). These enzymes are important for nutrient acquisition in carbohydrate-rich environments, including mud core samples, possibly rich in plant materials, where *S. rosetta* was isolated (27,78). Conservation of horizontally acquired glycosyl hydrolases in phagotrophic choanoflagellates suggest their importance in digesting various food sources and adapting to an environment high in plant biomass (40). Transporters, proteases, and glycosyl hydrolases have also been flagged as candidate HGTs in *M. brevicollis,* bdelloid rotifers, sponge, rumen ciliates, and fungi (40,65,66,78,79).

HGTs with functions related to amino acid biosynthesis contribute to the metabolic flexibility of *S. rosetta*. Some of these enzymes, particularly those involved in the DAP pathway of lysine biosynthesis, as well as those involved in the biosynthesis of arginine, threonine and methionine, have previously been identified as potential HGTs in *M. brevicollis* (40,41). Conservation of these potential HGTs in the choanoflagellate lineage and their absence in most metazoans, filastereans, and fungi suggest that they were likely transferred into older lineages and retained in the choanoflagellate lineage. Retention of genes involved in amino acid metabolism in choanoflagellates may contribute to unique metabolic competencies. On the other hand, these genes have been lost in the animal stem lineage (80,81) as multicellular animals rely on direct acquisition of essential amino acids from their diet (82). Cnidarians, however, regained the ability to synthesize aromatic amino acids tryptophan, phenylalanine, and other aromatic compounds through HGT events (80).

*S. rosetta,* like other marine organisms, are exposed to multiple biotic and abiotic stressors, including temperature, salinity, and pH fluctuations, osmotic and oxidative stress, and xenobiotics. The retention of horizontally acquired genes such as oxidoreductases, protein-folding chaperones, repair enzymes, and antibiotic degrading enzymes, may protect against cellular damage and strengthen organismal defenses against environmental stressors and potential pathogens (43).

### Candidate HGTs potentially regulate intercellular interactions

*S. rosetta* has several life stages regulated by extrinsic factors. Its genome harbors multiple adhesion receptors and cell membrane enzymes for substrate attachment, cell-cell communication, and colony formation that may regulate these life stage transitions (26,34,48–50,70). Several of these genes were identified as potential HGTs in *S. rosetta*. HGTs with dermatopontin (DPT/Ca^2+^ EGF) domains may function similar to dermatopontin, which is an acidic multifunctional matrix protein that promotes cell adhesion, ECM collagen fibrillogenesis, and cell assembly mediated by cell surface integrin binding (83–87). It is also a major component of the organic matrix of biomineralized tissues (e.g. mussels) (88). It is possible that expression of dermatopontin domain-containing genes in thecate cells is involved in controlling specific cell behaviors, such as phagocytosis, mating, or colony formation in *S. rosetta.* Absence of genes with DPT/Ca^2+^ EGF domains in some loricate choanoflagellates further highlights the possible importance of these genes in the production of the organic theca in Craspedida.

*S. rosetta* also has a rich repertoire of glycosyl hydrolases, glycosyltransferases, and mannosyltransferases that were potentially acquired through horizontal transfer. These enzymes are known to sculpt the ECM of animals by changing the structure of the extracellular matrix components and by modifying functional groups on molecules to regulate their interactions (89,90). Glycosyl hydrolases may degrade proteoglycans to control the mating process of *S. rosetta* (50,91). Glycosyltransferases regulate key signaling and adhesion proteins like cadherins and integrins (92–94). In particular, mannosyltransferases modify mannose groups on glycoproteins, which mediate the binding of C-type lectin domain proteins, such as the *rosetteless* gene that is crucial in rosette formation (95).

The *S. rosetta* genome contains two chondroitin sulfate lyase genes, both of which are candidate HGTs from potential bacterial donors. It was suggested that these proteins may be involved in endogenous processes for regulating mating in *S. rosetta* (50). It is also possible that GAG lyases contribute to the ability of choanoflagellates to modify cell walls and extracellular matrix proteins, which may facilitate transitions from one life stage to the next. Chondroitin sulfate lyase cleaves chondroitin sulfate glycosaminoglycans (GAGs) via an elimination mechanism resulting in disaccharides or oligosaccharides (96). GAGs are typically found as side chains on proteoglycans on cell membranes and the ECM of animal tissues where they regulate processes such as adhesion, differentiation, migration, proliferation, and cell-cell communication (96). The proteoglycan, chondroitin sulfate, was found to partially suppress the adhesive properties of dermatopontin (83–87), suggesting that breakdown of chondroitin sulfate through lyase activity may promote stronger cell adhesion. The bacterium, *Vibrio fischerii*, produces a chondroitin lyase called *EroS* (“extracellular regulator of sex”) that induces mating in the choanoflagellate, *S. rosetta* (50). The conservation of chondroitin lyase genes in *S. rosetta* and most choanoflagellate representatives suggest that these genes may be uncovered unique adaptive mechanisms in choanoflagellates. Further studies on the expression of these genes in different environmental conditions are needed in order to better understand their potential functions in the host.

As with other studies on the detection of HGTs in eukaryotes, it is important to note that the current work is limited by the lack of broader taxon representation in publicly available databases. In addition, parametric tests may not accurately estimate the number of HGT events, particularly for ancient transfer events. Moreover, it can be difficult to distinguish HGT events from gene gain or loss events in the last universal common ancestor as both result in patchy distribution of genes in the species tree. These limitations may have resulted in overestimation of the number of horizontally acquired genes in *S. rosetta*. Further development of methods to effectively filter out false positives and fine tune the results of HGT analysis is needed to reveal the true extent of horizontally acquired genes in the last common ancestor of choanoflagellates and animals.

## Conclusions

We detected a rich repertoire of genes potentially acquired through horizontal transfer in the genome of *S. rosetta*. Most of these genes were possibly ancient horizontal transfers gained prior to the divergence of choanoflagellates, as evidenced by similar genomic signatures and expression motifs to the host, as well as conservation in other choanoflagellate taxa. Our results show that the gain of genes from the microorganisms that *S. rosetta* interacts with may have played a key role in the diversification of cellular metabolic processes and contributed novel functions that enhanced the catalytic ability of enzymes thereby allowing *S. rosetta* to colonize diverse ecological niches. In addition, the acquisition of genes in *S. rosetta* involved in extra-cellular senses and sensory response to its environment, through the induction of reproduction and multicellular development, may have helped shape choanoflagellate evolution and multicellular life stages. We anticipate that future expression analysis of these promising candidate HGTs will provide a more in-depth understanding on their potential roles in choanoflagellates and animal multicellularity.

## Materials and Methods

### Identification of HGTs by Sequence Alignment

The 11,731 predicted *S. rosetta* protein sequences downloaded from the Ensembl protist database (97) (http://www.ensembl.org/; last accessed October, 2018) were aligned to the complete NCBI non-redundant (nr) protein database (http://www.ncbi.nlm.nih.gov/; last accessed October, 2018) using Diamond(98) local sequence alignment with a threshold E-value of 1×10^−5^. The taxonomic affiliation of hits were retrieved from NCBI taxonomy (http://www.ncbi.nlm.nih.gov/taxonomy), and a Python (99) script was used to collect hits with specific taxon IDs. Only sequences with the lowest E-value corresponding to bacteria, archaea, (i.e. Chlorophyta, Stramenopiles, Bacillariophyta, Pelagophyta, and Haptophta), unicellular fungi (i.e. Ascomycota and Basidiomycota), or protozoa were considered as potential HGTs; peptides with no hits to archaea, bacteria, unicellular algae, unicellular fungi and other protozoans were disregarded from further analysis.

### Determining the Alien Index Scores of Candidate HGTs

Alien Index (AI) analysis quantitatively measures how well the *S. rosetta* protein sequences align to non-metazoan versus metazoan protein sequences (65). The E-value of the best sequence alignment match of *S. rosetta* peptides against all metazoan or non-metazoan gene sequences from the NCBI nr database were used to compute the AI score of each gene using the formula (65):

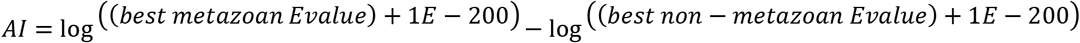

In cases where *S. rosetta* sequences had no hits to non-metazoans (excluding choanoflagellates) or metazoans, the E-value was set to 1. Genes that scored ≥45 were classified as foreign, 0 ≤ AI ≤ 45 as indeterminate, and less than 0 as metazoan genes (65).

### Genomic feature analysis of candidate HGTs and identification of gene orthologs

Genomic features of putative HGTs, specifically GC content at the third codon position (GC3) and codon usage, as represented by the codon bias index (CBI), were determined using correspondence analysis on CodonW (100). Coding sequence (CDS) lengths and intron numbers were determined from published information on the *S. rosetta* genome on Ensembl (97). To determine if there were mean differences between the genomic features of candidate HGTs compared to all *S. rosetta* genes, we performed Kruskal-Wallis test in RStudio version 1.2.1335 (101) with R version 3.6.1 (101). To identify orthologs of candidate HGTs, we obtained OrthoMCL groups data from the study of Richter et al. (2018).

### Gene expression analysis

To examine expression of candidate HGTs, we obtained transcriptome data from the study of Fairclough et al. (2013). Sequence libraries representing various life stages of *S. rosetta* were downloaded from the NCBI Short Read Archive (rosette cells, SRX042054; chain cells, SRX042047; thecate cells, SRX042052; swimming cells, SRX042053). Reads were mapped against the predicted cDNA sequences of *S. rosetta* using kallisto (102) with default settings to obtain gene expression values in transcripts per million reads (TPM).

### Protein domain and gene ontology analysis

Identification of protein domains and gene ontology associations was conducted on Blast2GO (103) to determine possible functions of candidate HGTs in *S. rosetta*. Gene ontology terms associated with each predicted peptide were determined from its best Diamond hit at an E-value ≤ 1×10^−5^ to the UniProt database (104). Enriched functions in the set of putative HGTs versus non-HGTs were identified using the Fisher’s exact test in topGO (105) in R package version 3.6.1 (101). Only functions with an FDR-corrected p-value ≤ 0.05 were considered statistically significant.

### Phylogenetic analysis of candidate HGTs

Peptide sequences of candidate HGTs were aligned with representative sequences from selected metazoa, eukaryotic, and prokaryotic taxa using MAFFT version 7 (106) then trimmed using Gblocks (107) with default settings to remove ambiguous and divergent protein alignments. For each protein sequence, Akaike information criterion (AIC) and Bayesian information criterion (BIC) scores were calculated in MEGAX (108). The best evolutionary model for phylogeny between different models tested for the protein sequences was determined by using the calculated AIC and BIC scores that had the smallest score difference, in this case mtrev substitution model was consistently used. Markov Chain Monte Carlo (MCMC) parameters of each analysis were set to 100,000 generations sampled every 100 trees. By default, the first 25% of the trees were discarded as burn-in. Bayesian phylogenetic trees were constructed using MrBayes 3.2.6 with posterior probabilities of 0.70-1.00 indicated on selected branches (109). Trees were edited online using Interactive Tree Of Life (iTOL) (110).

## Additional Data

**Table S1.** Summary of HGT analyses for all *S. rosetta* genes.

**Table S2.** Expression values of candidate HGTs

**Table S3.** Orthologs of candidate HGTs

**Table S4.** Candidate HGTs from selected potential bacterial donors

**Table S5.** PFAM domain architectures in candidate HGTs

**Table S6.** Gene ontology enrichment in candidate HGTs

**Table S7.** Associated functions of candidate HGTs

**Additional file 1. Phylogenetic analysis of selected candidate HGTs in *S. rosetta***. Peptide sequences of candidate HGTs were aligned to representative sequences from selected metazoa, eukaryotic, and prokaryotic taxa were using MAFFT version 7. Phylogenetic trees were generated using MrBayes 3.2.6 with posterior probabilities of 0.70-1.00 indicated on selected branches.

**Additional file 2. Candidate HGTs in amino acid biosynthetic pathways.** Genes flagged as HGTs in *S. rosetta* are shown in red lines, while those detected as HGTs in both *S. rosetta* and *M. brevicollis* are shown in blue lines. Text colors indicate potential donors of candidate HGTs.

## Abbreviations

ABC transporter: Adenosine triphosphate-binding cassette transporter
AI: Alien Index
Asd: aspartate-semialdehyde dehydrogenase
CBI: Codon bias index
CDS: coding sequence
DapF: diaminopimelate epimerase
DPT/Ca^2+^ EGF: Dermatopontin/calcium binding epidermal growth factor domain
ECM: Extracellular matrix
EroS: extracellular regulator of sex
FAD/NAD: Flavin adenine dinucleotide/ nucleotide adenine dinucleotide binding domain
FPKM: Fragments Per Kilobase of transcript per Million mapped reads
GAG lyase: glycosaminoglycan lyase
GO: Gene Ontology
HGT: Horizontal gene transfer
HSP: heat shock protein
HTT: Horizontal transposon transfers
LGT: Lateral gene transfer
LysC-lysA fusion: diaminopimelate decarboxylase and aspartate kinase
MCMC: Markov Chain Monte Carlo
ML: Maximum likelihood
NHL repeats: Ncl-1, HT2A and lin-41 (NHL) repeats
PFAM: Protein family
TE: transposable element

## Declarations

### Consent for publication

Not applicable.

### Availability of data and materials

The datasets generated during and/or analysed during the current study are available in: Mateo-Matriano D, Alegado RA, Conaco C. Detection of Horizontal Gene Transfer in the Choanoflagellate *Salpingoeca rosetta* data sets. figshare. 2020. [DOI: 10.6084/m9.figshare.12286658.v1]

### Authors’ contributions

DM and CC together conceived and designed the study. DM and CC conducted the analyses and drafted the manuscript. CC and RA advised on data analyses and helped to draft and edit the manuscript. All authors read and approved the final manuscript.

## Acknowledgements

We acknowledge the Core Facility for Bioinformatics, Philippine Genome Center (CFB-PGC) especially Joshua Dizon and Francis Tablizo for helping with the construction of the databases and the scripts used in this project. We greatly appreciate the insightful comments and suggestions of Dr. Cheryl Andam (College of Life Sciences and Agriculture UNH), Dr. Deo Onda (UP, MSI), Dr. Ron Leonard Dy (UP, NIMBB) and Becca Lensing (UH, C-MORE) on the analysis and interpretation of the data.

## References

1. Lorenz MG, Wackernagel W. Bacterial gene transfer by natural genetic transformation in the environment. Microbiol Rev. 1994;58(3):563–602.

2. Dubnau D. DNA Uptake in Bacteria. Annu Rev Microbiol. 1999;53(1):217–44.

3. Chen I, Dubnau D. DNA uptake during bacterial transformation. Vol. 2, Nature Reviews Microbiology. 2004. p. 241–9.

4. Heinemann JA, Sprague GF. Bacterial conjugative plasmids mobilize DNA transfer between bacteria and yeast. Nature. 1989;340(6230):205–9.

5. Llosa M, Gomis-Rüth FX, Coll M, De la Cruz F. Bacterial conjugation: A two-step mechanism for DNA transport. Mol Microbiol. 2002;45(1):1–8.

6. Haas AL, Baboshina O, Williams B, Schwartz LM. Coordinated induction of the ubiquitin conjugation pathway accompanies the developmentally programmed death of insect skeletal muscle. J Biol Chem. 1995;270(16):9407–12.

7. Norman A, Hansen LH, Sørensen SJ. Conjugative plasmids: Vessels of the communal gene pool. Vol. 364, Philosophical Transactions of the Royal Society B: Biological Sciences. 2009. p. 2275–89.

8. Kyndt T, Quispe D, Zhai H, Jarret R, Ghislain M, Liu Q, et al. The genome of cultivated sweet potato contains Agrobacterium T-DNAs with expressed genes: An example of a naturally transgenic food crop. Proc Natl Acad Sci U S A. 2015;112(18):5844–9.

9. Keeling PJ, Palmer JD. Horizontal gene transfer in eukaryotic evolution. Vol. 9, Nature Reviews Genetics. 2008. p. 605–18.

10. Soucy SM, Huang J, Gogarten JP. Horizontal gene transfer: Building the web of life. Vol. 16, Nature Reviews Genetics. 2015. p. 472–82.

11. Huang J. Horizontal gene transfer in eukaryotes: The weak-link model. BioEssays. 2013;35(10):868–75.

12. Takeuchi N, Kaneko K, Koonin E V. Horizontal gene transfer can rescue prokaryotes from Muller’s ratchet: Benefit of DNA from dead cells and population subdivision. G3 Genes, Genomes, Genet. 2014;4(2):325–39.

13. Ford Doolittle W. You are what you eat: A gene transfer ratchet could account for bacterial genes in eukaryotic nuclear genomes. Vol. 14, Trends in Genetics. 1998. p. 307–11.

14. O’Malley MA. Endosymbiosis and its implications for evolutionary theory. Proc Natl Acad Sci U S A. 2015;112(33):10270–7.

15. Margulis L, Chapman M, Guerrero R, Hall J. The last eukaryotic common ancestor (LECA): Acquisition of cytoskeletal motility from aerotolerant spirochetes in the Proterozoic Eon. Proc Natl Acad Sci U S A. 2006;103(35):13080–5.

16. Southworth J, Grace CA, Marron AO, Fatima N, Carr M. A genomic survey of transposable elements in the choanoflagellate Salpingoeca rosetta reveals selection on codon usage. Vol. 10, Mobile DNA. 2019.

17. Pace JK, Gilbert C, Clark MS, Feschotte C. Repeated horizontal transfer of a DNA transposon in mammals and other tetrapods. Proc Natl Acad Sci U S A. 2008;105(44):17023–8.

18. Mizrokhi LJ, Mazo AM. Evidence for horizontal transmission of the mobile element jockey between distant Drosophila species. Proc Natl Acad Sci U S A. 1990;87(23):9216–20.

19. Williams D, Fournier GP, Lapierre P, Swithers KS, Green AG, Andam CP, et al. A Rooted Net of Life. Vol. 6, Biology Direct. 2011.

20. Andam CP, Gogarten JP. Biased gene transfer and its implications for the concept of lineage. Biol Direct. 2011;6.

21. Huang J, Gogarten JP. Ancient horizontal gene transfer can benefit phylogenetic reconstruction. Trends Genet. 2006;22(7):361–6.

22. Jain R, Rivera MC, Lake JA. Horizontal gene transfer among genomes: The complexity hypothesis. Proc Natl Acad Sci U S A. 1999;96(7):3801–6.

23. Jain R, Rivera MC, Moore JE, Lake JA. Horizontal gene transfer accelerates genome innovation and evolution. Mol Biol Evol. 2003;20(10):1598–602.

24. Swithers KS, Gogarten JP, Fournier GP. Trees in the web of life. Vol. 8, Journal of Biology. 2009.

25. Redrejo-Rodriǵuez M, Munõz-Espín D, Holguera I, Menciá M, Salas M. Functional eukaryotic nuclear localization signals are widespread in terminal proteins of bacteriophages. Proc Natl Acad Sci U S A. 2012;109(45):18482–7.

26. Dayel MJ, King N. Prey capture and phagocytosis in the choanoflagellate Salpingoeca rosetta. PLoS One. 2014;9(5).

27. Dayel MJ, Alegado RA, Fairclough SR, Levin TC, Nichols SA, McDonald K, et al. Cell differentiation and morphogenesis in the colony-forming choanoflagellate Salpingoeca rosetta. Dev Biol. 2011;357(1):73–82.

28. Richter DJ, Nitsche F. Choanoflagellatea. In: Handbook of the Protists: Second Edition. 2017. p. 1479–96.

29. Hoffmeyer TT, Burkhardt P. Choanoflagellate models — Monosiga brevicollis and Salpingoeca rosetta. Vol. 39, Current Opinion in Genetics and Development. 2016. p. 42–7.

30. Lang BF, O’Kelly C, Nerad T, Gray MW, Burger G. The closest unicellular relatives of animals. Curr Biol. 2002;12(20):1773–8.

31. Ruiz-Trillo I, Roger AJ, Burger G, Gray MW, Lang BF. A phylogenomic investigation into the origin of Metazoa. Mol Biol Evol. 2008;25(4):664–72.

32. Carr M, Richter DJ, Fozouni P, Smith TJ, Jeuck A, Leadbeater BSC, et al. A six-gene phylogeny provides new insights into choanoflagellate evolution. Mol Phylogenet Evol. 2017;107:166–78.

33. Nitsche F, Carr M, Arndt H, Leadbeater BSC. Higher level taxonomy and molecular phylogenetics of the Choanoflagellatea. J Eukaryot Microbiol. 2011;58(5):452–62.

34. Fairclough SR, Chen Z, Kramer E, Zeng Q, Young S, Robertson HM, et al. Premetazoan genome evolution and the regulation of cell differentiation in the choanoflagellate Salpingoeca rosetta. Genome Biol. 2013;14(2):1–15.

35. King N, Westbrook MJ, Young SL, Kuo A, Abedin M, Chapman J, et al. The genome of the choanoflagellate Monosiga brevicollis and the origin of metazoans. Nature. 2008;451(7180):783–8.

36. Jeuck A, Arndt H, Nitsche F. Extended phylogeny of the Craspedida (Choanomonada). Eur J Protistol. 2014;50(4):430–43.

37. Marron AO, Alston MJ, Heavens D, Akam M, Caccamo M, Holland PWH, et al. A family of diatom-like silicon transporters in the siliceous loricate choanoflagellates. Proc R Soc B Biol Sci. 2013;280(1756).

38. Bapteste E, Moreira D, Philippe H. Rampant horizontal gene transfer and phospho-donor change in the evolution of the phosphofructokinase. Gene. 2003;318(1-2):185–91.

39. Malik SB, Ramesh MA, Hulstrand AM, Logsdon JM. Protist homologs of the meiotic Spo11 gene and topoisomerase VI reveal an evolutionary history of gene duplication and lineage-specific loss. Mol Biol Evol. 2007;24(12):2827–41.

40. Yue J, Sun G, Hu X, Huang J. The scale and evolutionary significance of horizontal gene transfer in the choanoflagellate Monosiga brevicollis. BMC Genomics. 2013;14(1).

41. Sun G, Huang J. Horizontally acquired DAP pathway as a unit of self-regulation. J Evol Biol. 2011;24(3):587–95.

42. Maruyama S, Matsuzaki M, Misawa K, Nozaki H. Cyanobacterial contribution to the genomes of the plastid-lacking protists. BMC Evol Biol. 2009;9(1).

43. Nedelcu AM, Miles IH, Fagir AM, Karol K. Adaptive eukaryote-to-eukaryote lateral gene transfer: Stress-related genes of algal origin in the closest unicellular relatives of animals. J Evol Biol. 2008;21(6):1852–60.

44. Nedelcu AM, Blakney AJC, Logue KD. Functional replacement of a primary metabolic pathway via multiple independent eukaryote-to-eukaryote gene transfers and selective retention. J Evol Biol. 2009;22(9):1882–94.

45. Tucker RP, Beckmann J, Leachman NT, Schöler J, Chiquet-Ehrismann R. Phylogenetic analysis of the teneurins: Conserved features and premetazoan ancestry. Mol Biol Evol. 2012;29(3):1019–29.

46. Torruella G, Suga H, Riutort M, Peretó J, Ruiz-Trillo I. The evolutionary history of lysine biosynthesis pathways within eukaryotes. J Mol Evol. 2009;69(3):240–8.

47. Fairclough SR, Dayel MJ, King N. Multicellular development in a choanoflagellate. Vol. 20, Current Biology. 2010.

48. Alegado RA, Brown LW, Cao S, Dermenjian RK, Zuzow R, Fairclough SR, et al. A bacterial sulfonolipid triggers multicellular development in the closest living relatives of animals. Elife. 2012;2012(1).

49. Woznica A, Cantley AM, Beemelmanns C, Freinkman E, Clardy J, King N. Bacterial lipids activate, synergize, and inhibit a developmental switch in choanoflagellates. Proc Natl Acad Sci U S A. 2016;113(28):7894–9.

50. Woznica A, Gerdt JP, Hulett RE, Clardy J, King N. Mating in the Closest Living Relatives of Animals Is Induced by a Bacterial Chondroitinase. Cell. 2017;170(6):1175–1183.e11.

51. Ray S, Kassan A, Busija AR, Rangamani P, Patel HH. The plasma membrane as a capacitor for energy and metabolism. Vol. 310, American Journal of Physiology - Cell Physiology. 2016. p. C181–92.

52. Daubin V, Lerat E, Perrière G. The source of laterally transferred genes in bacterial genomes. Genome Biol. 2003;4(9).

53. Lawrence JG, Ochman H. Molecular archaeology of the Escherichia coli genome. Proc Natl Acad Sci U S A. 1998;95(16):9413–7.

54. Langille MGI, Hsiao WWL, Brinkman FSL. Detecting genomic islands using bioinformatics approaches. Vol. 8, Nature Reviews Microbiology. 2010. p. 373–82.

55. Hildebrand F, Meyer A, Eyre-Walker A. Evidence of selection upon genomic GC-content in bacteria. PLoS Genet. 2010;6(9).

56. Hershberg R, Petrov DA. Evidence that mutation is universally biased towards AT in bacteria. PLoS Genet. 2010;6(9).

57. Ford Doolittle W. Lateral genomics. Vol. 24, Trends in Biochemical Sciences. 1999.

58. Rivera MC, Jain R, Moore JE, Lake JA. Genomic evidence for two functionally distinct gene classes. Proc Natl Acad Sci U S A. 1998;95(11):6239–44.

59. Sicheritz-Ponten T. A phylogenomic approach to microbial evolution. Nucleic Acids Res. 2001;29(2):545–52.

60. Gogarten JP, Senejani AG, Zhaxybayeva O, Olendzenski L, Hilario E. Inteins: Structure, Function, and Evolution. Annu Rev Microbiol. 2002;56(1):263–87.

61. Brown JR. Ancient horizontal gene transfer. Vol. 4, Nature Reviews Genetics. 2003. p. 121–32.

62. Wellner A, Lurie MN, Gophna U. Complexity, connectivity, and duplicability as barriers to lateral gene transfer. Genome Biol. 2007;8(8).

63. Lercher MJ, Pál C. Integration of horizontally transferred genes into regulatory interaction networks takes many million years. Mol Biol Evol. 2008;25(3):559–67.

64. Yue J, Hu X, Huang J. Origin of plant auxin biosynthesis. Vol. 19, Trends in Plant Science. 2014. p. 764–70.

65. Gladyshev EA, Meselson M, Arkhipova IR. Massive horizontal gene transfer in bdelloid rotifers. Science (80-). 2008;320(5880):1210–3.

66. Conaco C, Tsoulfas P, Sakarya O, Dolan A, Werren J, Kosik KS. Detection of prokaryotic genes in the amphimedon queenslandica genome. PLoS One. 2016;11(3).

67. Boschetti C, Carr A, Crisp A, Eyres I, Wang-Koh Y, Lubzens E, et al. Biochemical Diversification through Foreign Gene Expression in Bdelloid Rotifers. PLoS Genet. 2012;8(11).

68. Lal D, Lal R. Evolution of mercuric reductase (merA) gene: A case of horizontal gene transfer. Microbiology. 2010;79(4):500–8.

69. Eyres I, Boschetti C, Crisp A, Smith TP, Fontaneto D, Tunnacliffe A, et al. Horizontal gene transfer in bdelloid rotifers is ancient, ongoing and more frequent in species from desiccating habitats. BMC Biol. 2015;13(1).

70. Richter DJ, Fozouni P, Eisen MB, King N. Gene family innovation, conservation and loss on the animal stem lineage. Elife. 2018;7.

71. Leadbeater BSC. The choanoflagellates: Evolution, biology and ecology. The Choanoflagellates: Evolution, Biology and Ecology. 2014. 1–315 p.

72. Walmsley AR, Barrett MP, Bringaud F, Gould GW. Sugar transporters from bacteria, parasites and mammals: Structure-activity relationships. Vol. 23, Trends in Biochemical Sciences. 1998. p. 476–81.

73. Goffeau A, De Hertogh B. ABC Transporters. In: Encyclopedia of Biological Chemistry: Second Edition. 2013. p. 7–11.

74. Lombard V, Golaconda Ramulu H, Drula E, Coutinho PM, Henrissat B. The carbohydrate-active enzymes database (CAZy) in 2013. Nucleic Acids Res. 2014;42(D1).

75. Michikawa M, Ichinose H, Momma M, Biely P, Jongkees S, Yoshida M, et al. Structural and biochemical characterization of glycoside hydrolase family 79 β-glucuronidase from Acidobacterium capsulatum. J Biol Chem. 2012;287(17):14069–77.

76. van der Hoorn RAL. Plant Proteases: From Phenotypes to Molecular Mechanisms. Annu Rev Plant Biol. 2008;59(1):191–223.

77. Tan L, Eberhard S, Pattathil S, Warder C, Glushka J, Yuan C, et al. An Arabidopsis cell wall proteoglycan consists of pectin and arabinoxylan covalently linked to an arabinogalactan protein. Plant Cell. 2013;25(1):270–87.

78. Ricard G, McEwan NR, Dutilh BE, Jouany JP, Macheboeuf D, Mitsumori M, et al. Horizontal gene transfer from bacteria to rumen ciliates indicates adaptation to their anaerobic, carbohydrates-rich environment. BMC Genomics. 2006;7.

79. Garcia-Vallvé S, Romeu A, Palau J. Horizontal gene transfer of glycosyl hydrolases of the rumen fungi. Mol Biol Evol. 2000;17(3):352–61.

80. Starcevic A, Akthar S, Dunlap WC, Shick JM, Hranueli D, Cullum J, et al. Enzymes of the shikimic acid pathway encoded in the genome of a basal metazoan, Nematostella vectensis, have microbial origins. Proc Natl Acad Sci U S A. 2008;105(7):2533–7.

81. López-Escardó D, Grau-Bové X, Guillaumet-Adkins A, Gut M, Sieracki ME, Ruiz-Trillo Reconstruction of protein domain evolution using single-cell amplified genomes of uncultured choanoflagellates sheds light on the origin of animals. Philos Trans R Soc B Biol Sci. 2019;374(1786).

82. Payne SH, Loomis WF. Retention and loss of amino acid biosynthetic pathways based on analysis of whole-genome sequences. Eukaryot Cell. 2006;5(2):272–6.

83. Kuroda K, Okamoto O, Shinkai H. Dermatopontin expression is decreased in hypertrophic scar and systemic sclerosis skin fibroblasts and is regulated by transforming growth factor- β1, interleukin-4, and matrix collagen. J Invest Dermatol. 1999;112(5):706–10.

84. Okamoto O, Fujiwara S. Dermatopontin, a novel player in the biology of the extracellular matrix. Vol. 47, Connective Tissue Research. 2006. p. 177–89.

85. Lewandowska K, Choi HU, Rosenberg LC, Sasse J, Neame PJ, Culp LA. Extracellular matrix adhesion-promoting activities of a dermatan sulfate proteoglycan-associated protein (22K) from bovine fetal skin. J Cell Sci. 1991;99(3):657–68.

86. Lewandowska K, Choi HU, Rosenberg LC, Zardi L, Culp LA. Fibronectin-mediated adhesion of fibroblasts: Inhibition by dermatan sulfate proteoglycan and evidence for a cryptic glycosaminoglycan-binding domain. J Cell Biol. 1987;105(3):1443–54.

87. Winnemoller M, Schon P, Vischer P, Kresse H. Interactions between thrombospondin and the small proteoglycan decorin: Interference with cell attachment. Eur J Cell Biol. 1992;59(1):47–55.

88. Marxen JC, Nimtz M, Becker W, Mann K. The major soluble 19.6 kDa protein of the organic shell matrix of the freshwater snail Biomphalaria glabrata is an N-glycosylated dermatopontin. Biochim Biophys Acta - Proteins Proteomics. 2003;1650(1-2):92–8.

89. Wetzel LA, Levin TC, Hulett RE, Chan D, King GA, Aldayafleh R, et al. Predicted glycosyltransferases promote development and prevent spurious cell clumping in the choanoflagellate S. Rosetta. Elife. 2018;7.

90. Larson BT, Ruiz-Herrero T, Lee S, Kumar S, Mahadevan L, King N. Biophysical principles of choanoflagellate self-organization. Proc Natl Acad Sci U S A. 2020;117(3):1303–11.

91. Trincone A, Tramice A, Giordano A, Andreotti G. Glycoside hydrolases in aplysia fasciata: Analysis and applications. Biotechnol Genet Eng Rev. 2008;25(1):129–48.

92. Sawaguchi S, Varshney S, Ogawa M, Sakaidani Y, Yagi H, Takeshita K, et al. O-GlcNAc on NOTCH1 EGF repeats regulates ligand-induced Notch signaling and vascular development in mammals. Elife. 2017;6.

93. Stratford M. Yeast flocculation: Receptor definition by mnn mutants and concanavalin A. Yeast. 1992;8(8):635–45.

94. Naderer T, Vince J, McConville M. Surface Determinants of Leishmania Parasites and their Role in Infectivity in the Mammalian Host. Curr Mol Med. 2005;4(6):649–65.

95. Levin TC, Greaney AJ, Wetzel L, King N. The Rosetteless gene controls development in the choanoflagellate S. rosetta. Elife. 2014;3.

96. Zhang F, Zhang Z, Linhardt RJ. Glycosaminoglycans. In: Handbook of Glycomics. 2010. p. 59–80.

97. Kersey PJ, Allen JE, Allot A, Barba M, Boddu S, Bolt BJ, et al. Ensembl Genomes 2018: An integrated omics infrastructure for non-vertebrate species. Nucleic Acids Res. 2018;46(D1):D802–8.

98. Buchfink B, Xie C, Huson DH. Fast and sensitive protein alignment using DIAMOND. Vol. 12, Nature Methods. 2014. p. 59–60.

99. Python Software Foundation. Python Language Reference, version 3.5. Python Software Foundation. 2016.

100. Wang M, Zhang J, Zhou JH, Chen HT, Ma LN, Ding YZ, et al. Analysis of codon usage in type 1 and the new genotypes of duck hepatitis virus. BioSystems. 2011;106(1):45–50.

101. 3.5.1. RDCT. A Language and Environment for Statistical Computing. R Found Stat Comput [Internet]. 2018;2:https://www.R-project.org. Available from: http://www.r-project.org

102. Bray NL, Pimentel H, Melsted P, Pachter L. Near-optimal probabilistic RNA-seq quantification. Nat Biotechnol. 2016;34(5):525–7.

103. Conesa A, Götz S, García-Gómez JM, Terol J, Talón M, Robles M. Blast2GO: A universal tool for annotation, visualization and analysis in functional genomics research. Bioinformatics. 2005;21(18):3674–6.

104. Bateman A. UniProt: A worldwide hub of protein knowledge. Nucleic Acids Res. 2019;47(D1):D506–15.

105. Alexa A, Rahnenfuhrer J. topGO: topGO: Enrichment analysis for Gene On-tology. R package version 2.26.0. 2009. p. R package version 2.22.0.

106. Katoh K, Standley DM. MAFFT multiple sequence alignment software version 7: Improvements in performance and usability. Mol Biol Evol. 2013;30(4):772–80.

107. Castresana J. Selection of conserved blocks from multiple alignments for their use in phylogenetic analysis. Mol Biol Evol. 2000;17(4):540–52.

108. Kumar S, Stecher G, Li M, Knyaz C, Tamura K. MEGA X: Molecular evolutionary genetics analysis across computing platforms. Mol Biol Evol. 2018;35(6):1547–9.

109. Ronquist F, Teslenko M, Van Der Mark P, Ayres DL, Darling A, Höhna S, et al. Mrbayes 3.2: Efficient bayesian phylogenetic inference and model choice across a large model space. Syst Biol. 2012;61(3):539–42.

110. Letunic I, Bork P. Interactive Tree of Life (iTOL) v4: Recent updates and new developments. Nucleic Acids Res. 2019;47(W1).

