## Additional File 1 for "Detection of Horizontal Gene Transfer in the Genome of the Choanoflagellate *Salpingoeca rosetta*"

**Additional file 1. Selected unrooted phylogenetic trees of candidate HGTs in *S. rosetta*.** Phylogenetic trees were generated using MrBayes 3.2.6 with posterior probabilities of 0.70-1.00 indicated as dots on selected branches. Sequences were aligned using Blastp search in NCBI.

---

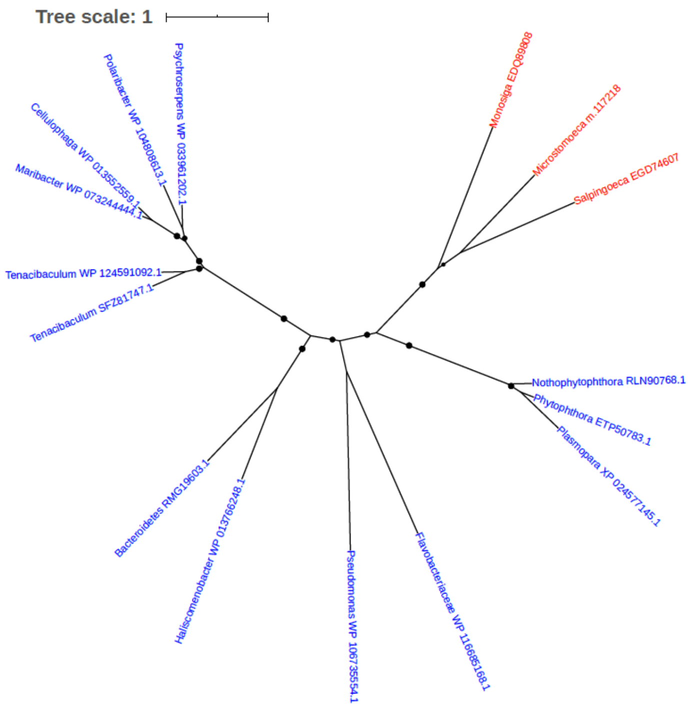


Figure S1. Phylogeny of 2,4-dienoyl-CoA reductase FadH2 (EGD74607)

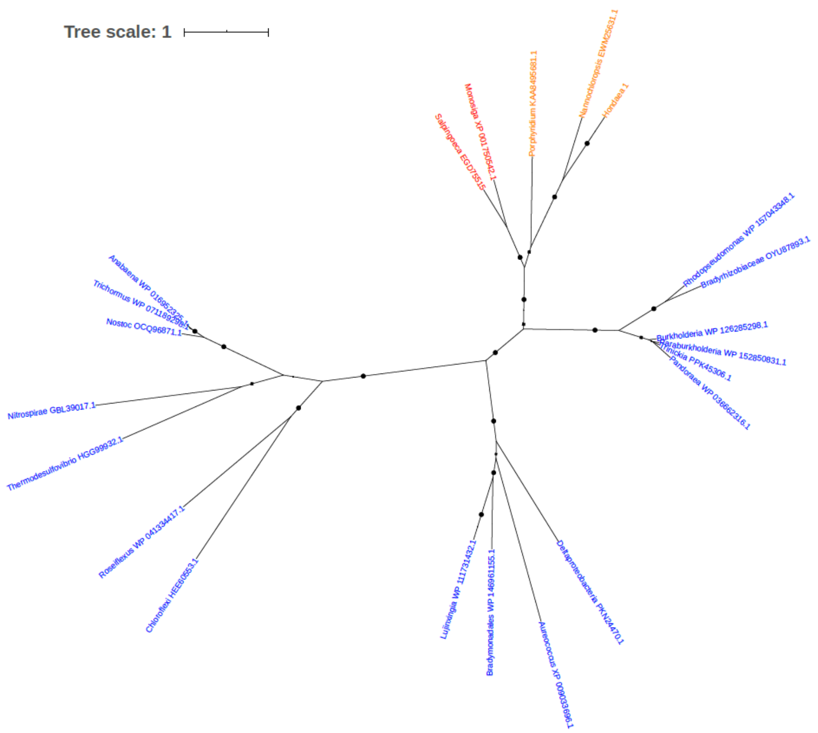


Figure S2. Phylogeny of 3-dehydroquinate synthase (EGD75515)

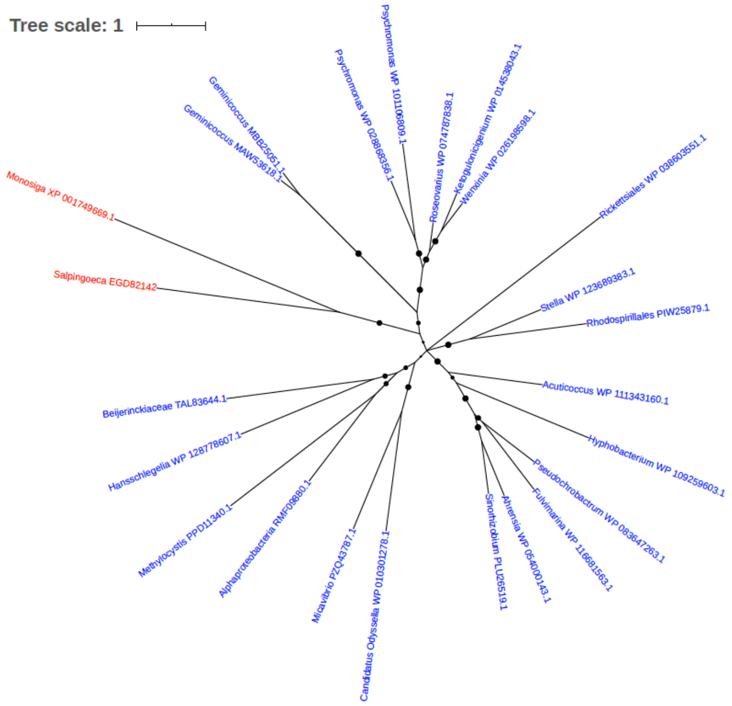


Figure S3. Phylogeny of 3-demethylubiquinone-9 3-methyltransferase (EGD82142)

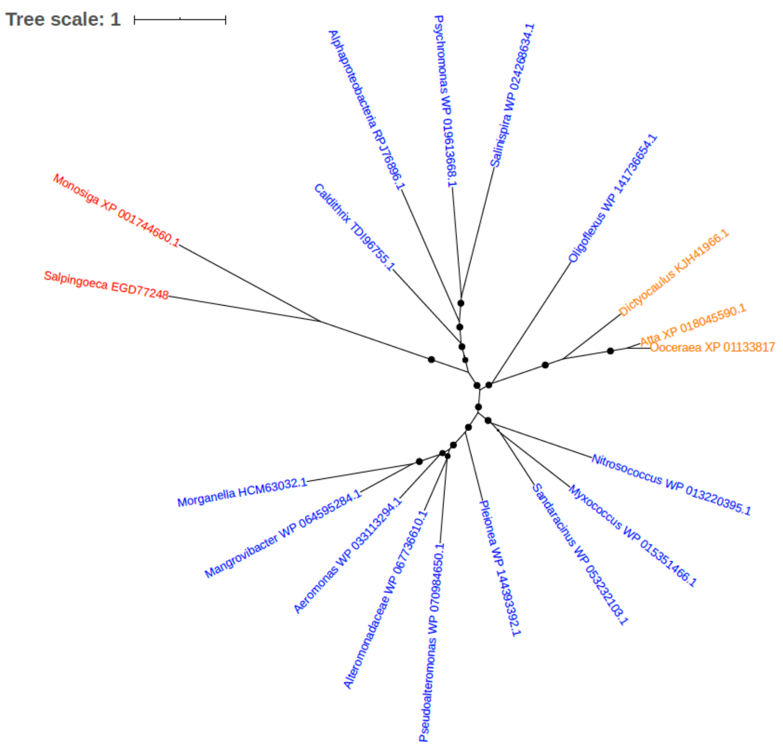


Figure S4. Phylogeny of 3-hydroxybutyryl-CoA dehydratase (EGD77248)

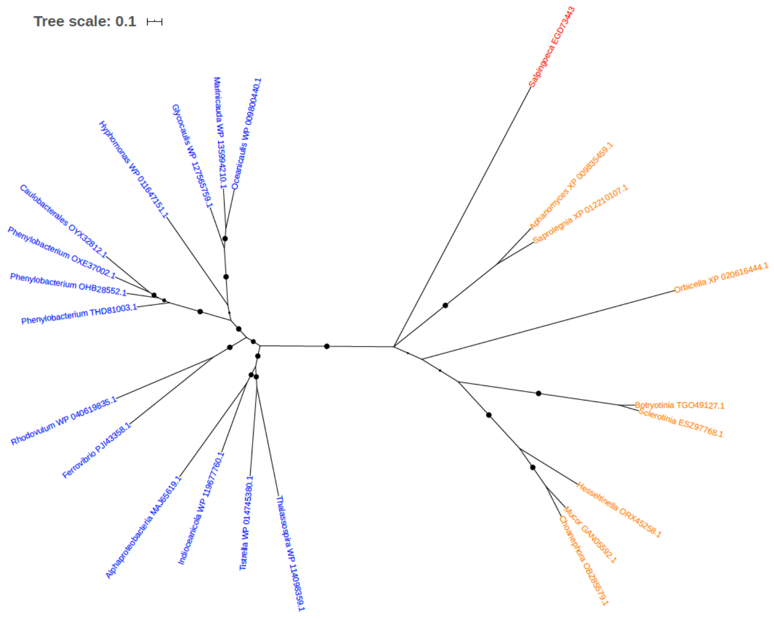


Figure S5. Phylogeny of 3-oxoacyl-[acyl-carrier-protein] synthase (EGD73443)

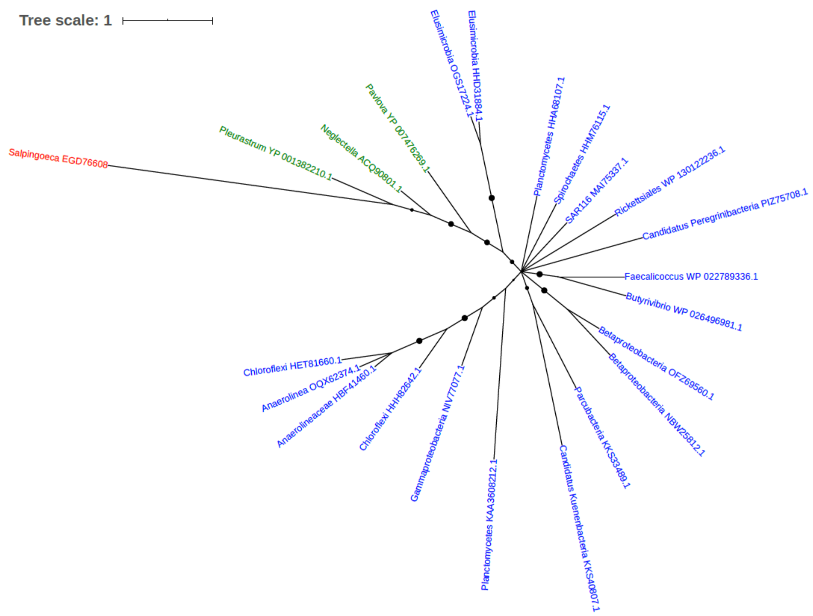


Figure S6. Phylogeny of 30S ribosomal protein S11 (EGD76608)

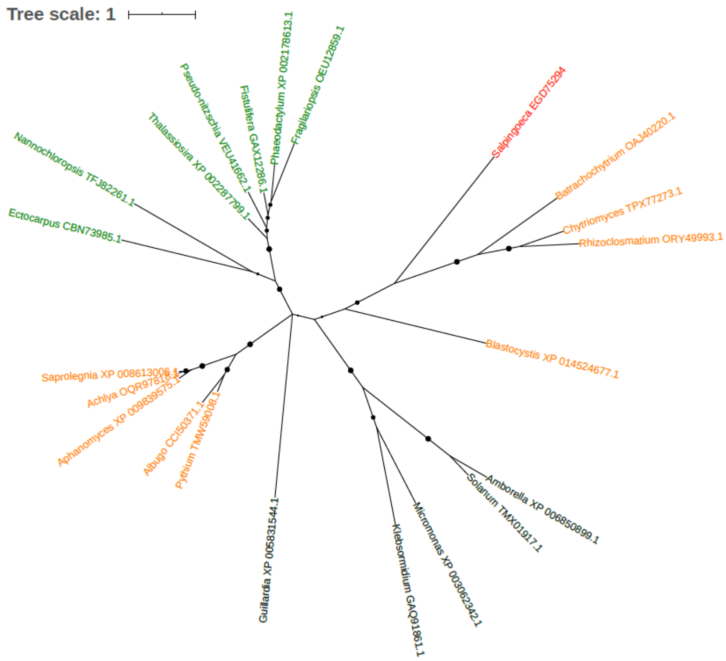


Figure S7. Phylogeny of 30S ribosomal protein S16 (EGD75394)

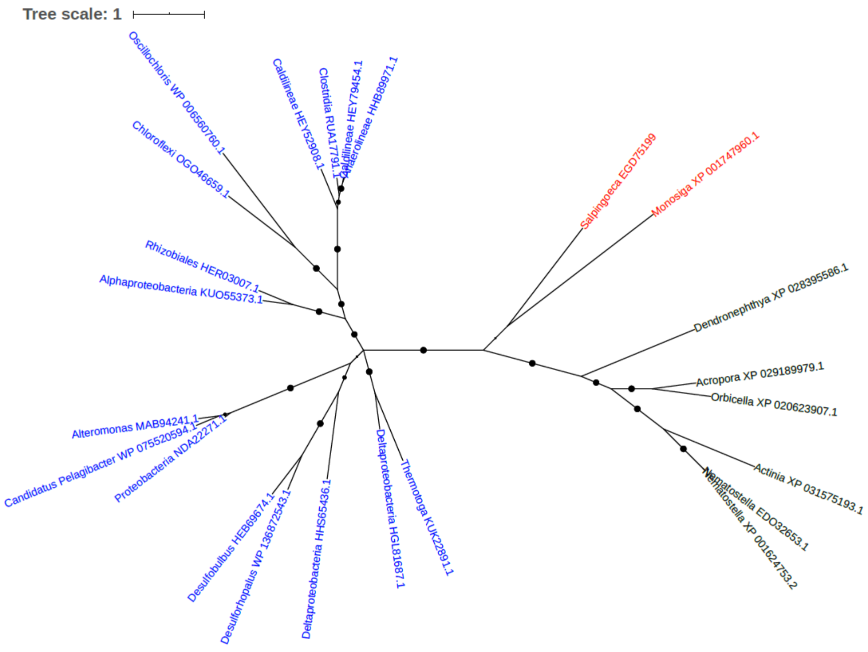


Figure S8. Phylogeny of 30S ribosomal protein S7 (EGD75199)

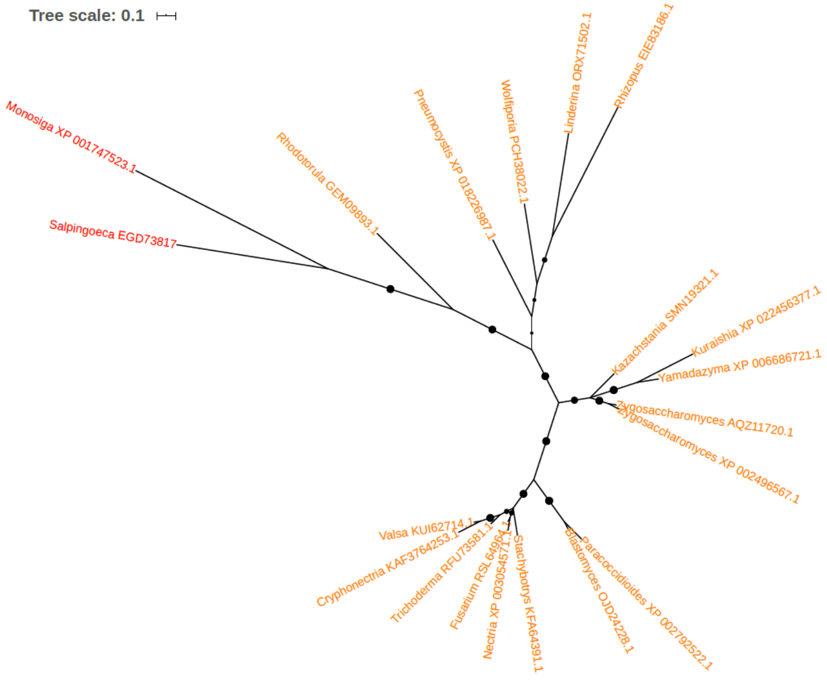


Figure S9. Phylogeny of 40S ribosomal protein S21 (EGD73817)

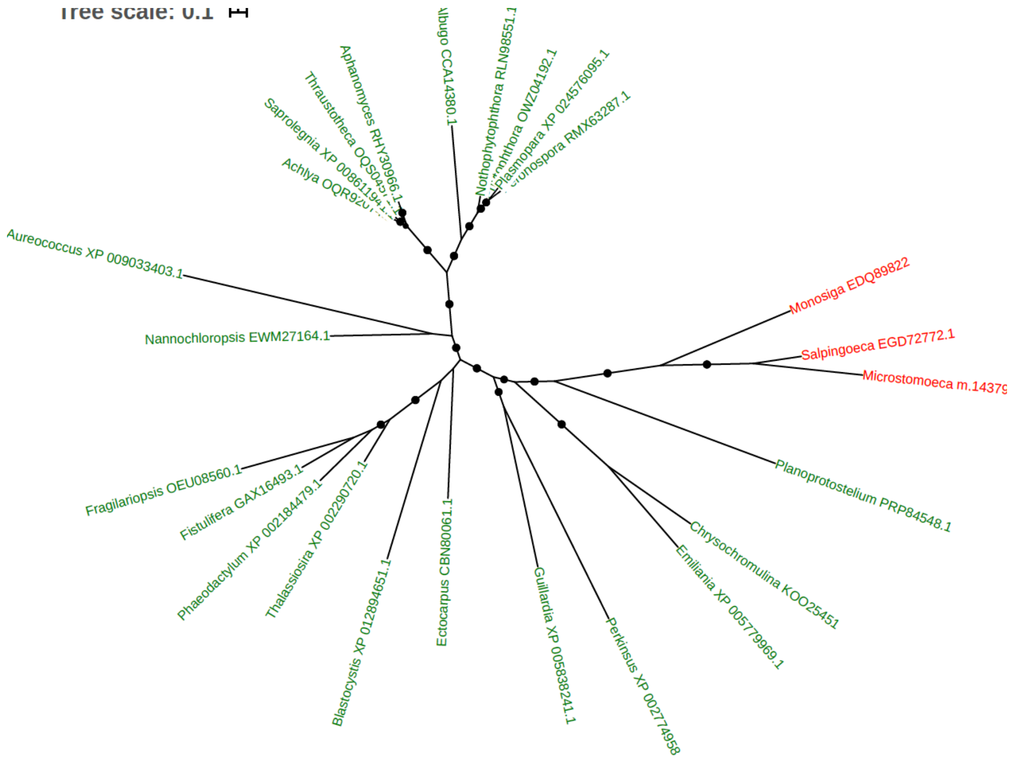


Figure S10. Phylogeny of 5-aminoimidazole-4-carboxamide ribonucleotide transformylase (EGD72772)

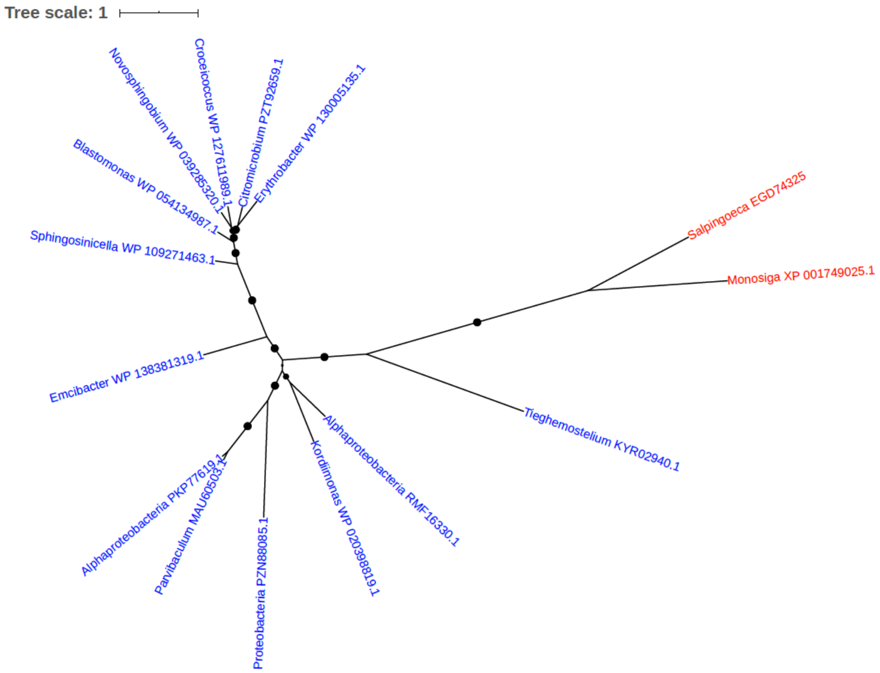


Figure S11. Phylogeny of 5-aminolevulinic acid synthase (EGD74325)

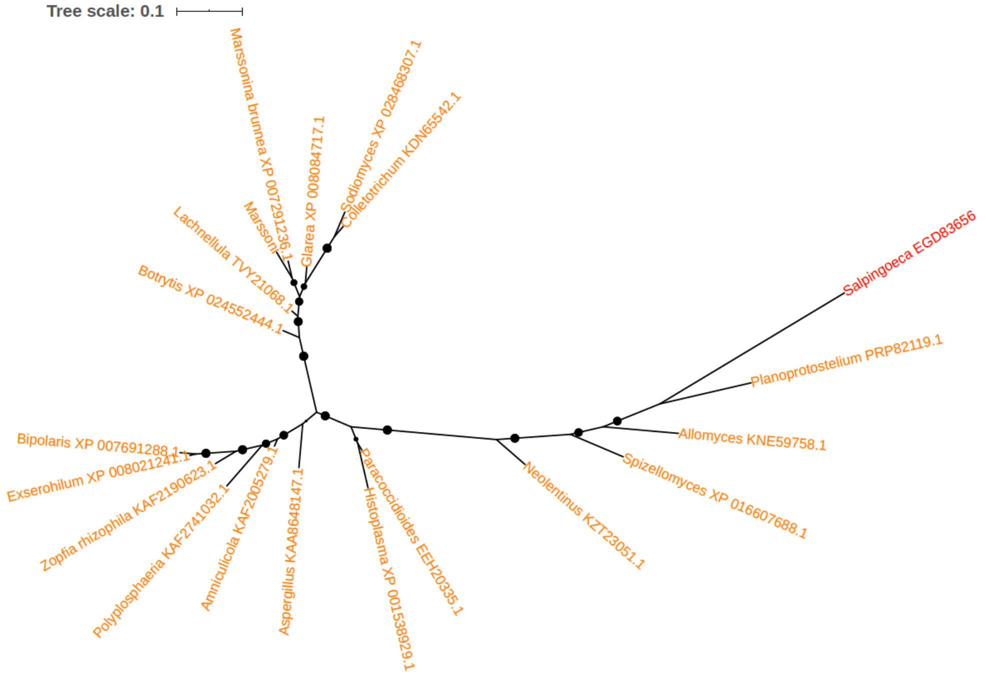


Figure S12. Phylogeny of 60S ribosomal protein L43 (EGD83656)

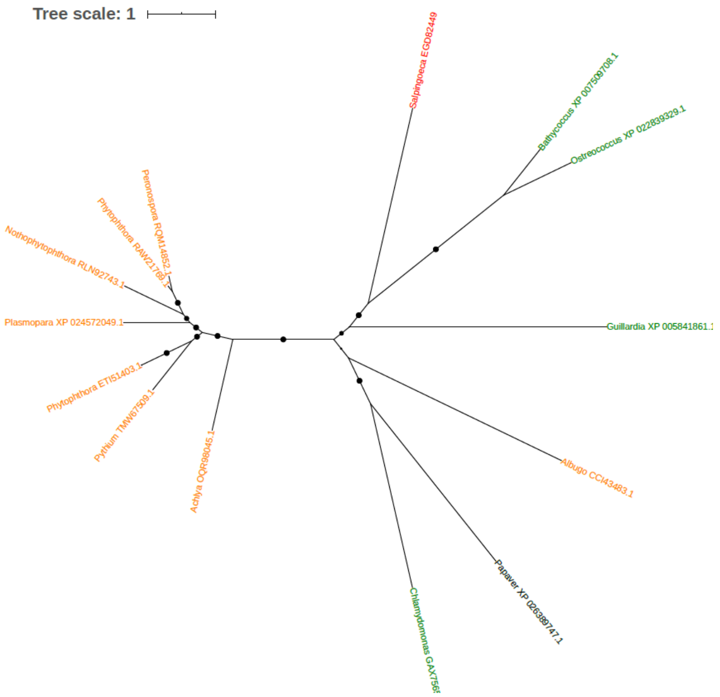


Figure S13. Phylogeny of ABC transporter (EGD82449)

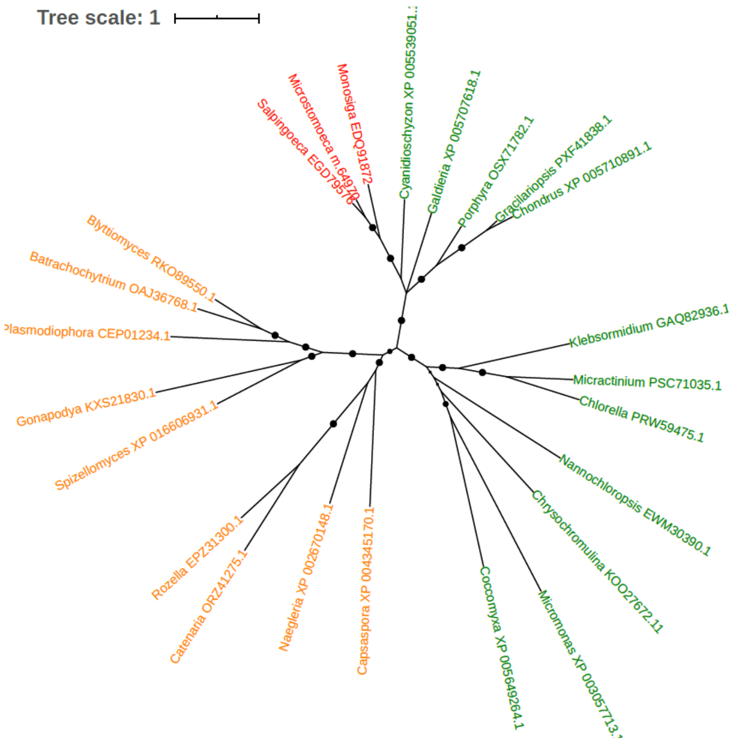


Figure S14. Phylogeny of ABC transporter (EGD79576)

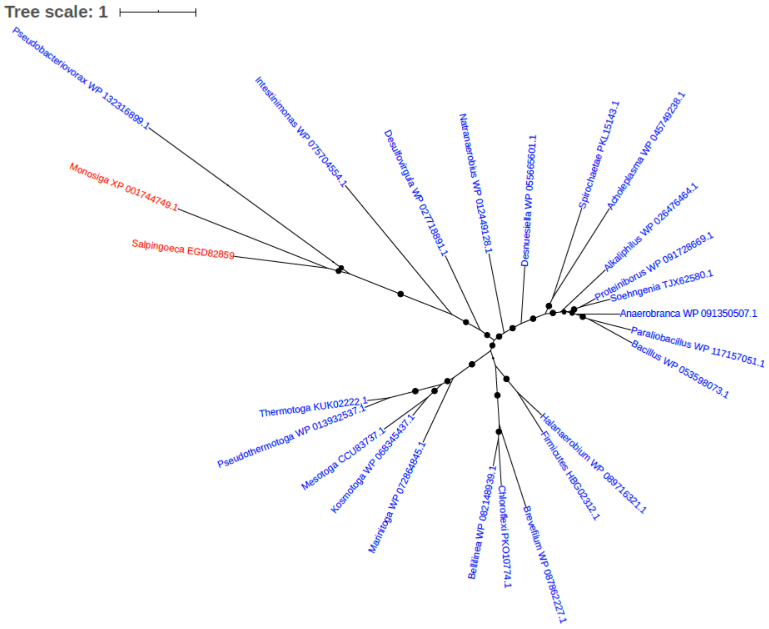


Figure S15. Phylogeny of Acetyl-CoA hydrolase/transferase (EGD82859)

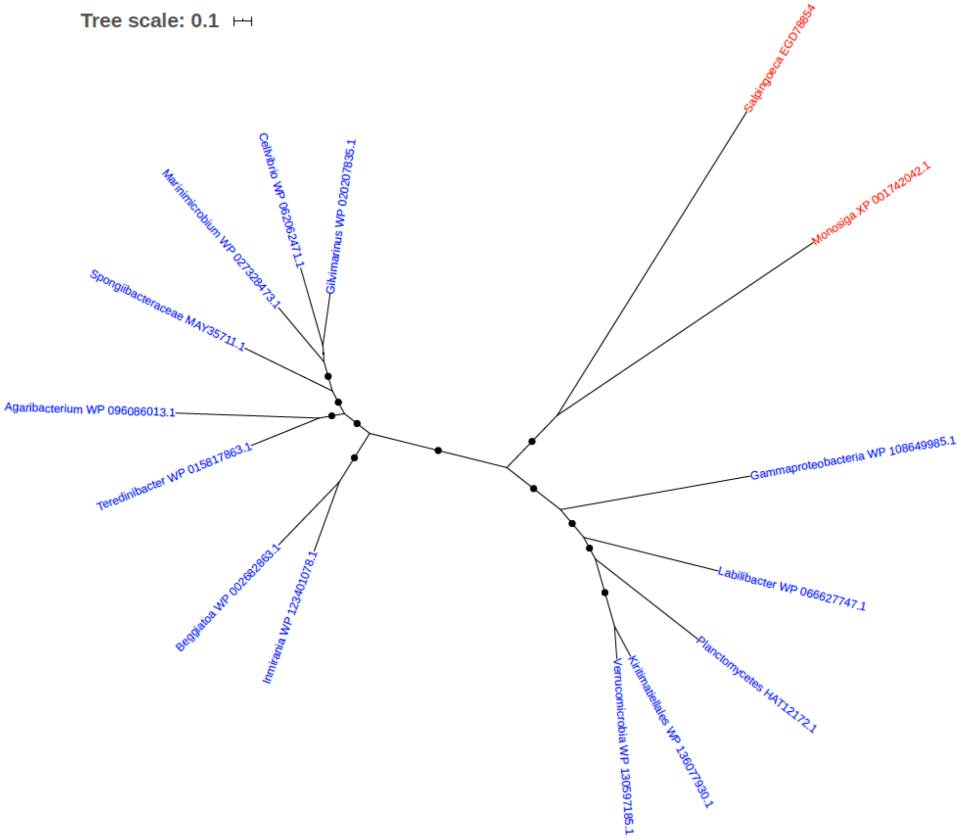


Figure S16. Phylogeny of Acetyl-coenzyme A synthetase (EC 6.2.1.1) (EGD78854)

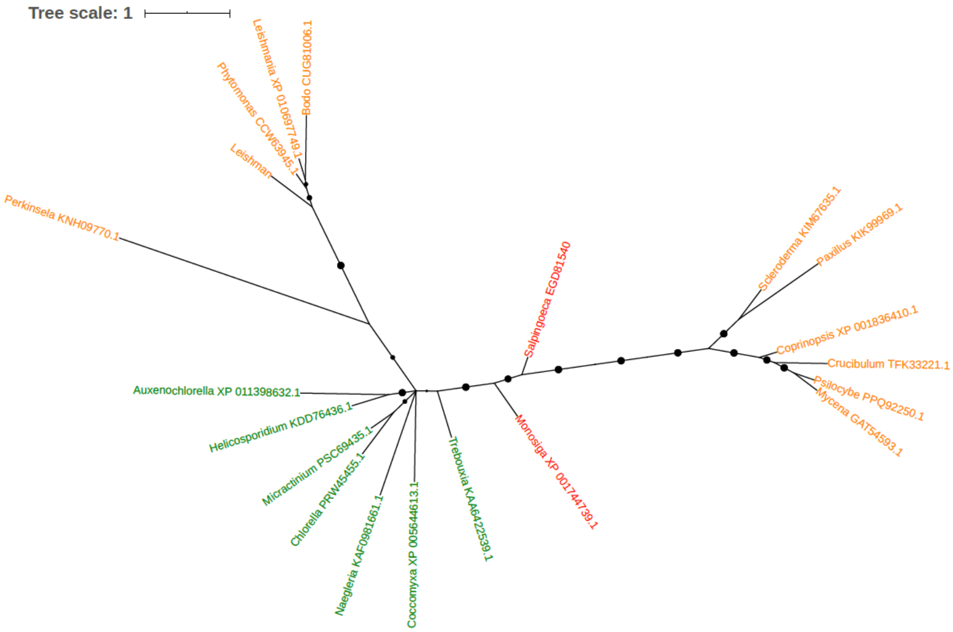


Figure S17. Phylogeny of Acyl-CoA binding protein (EGD81540)

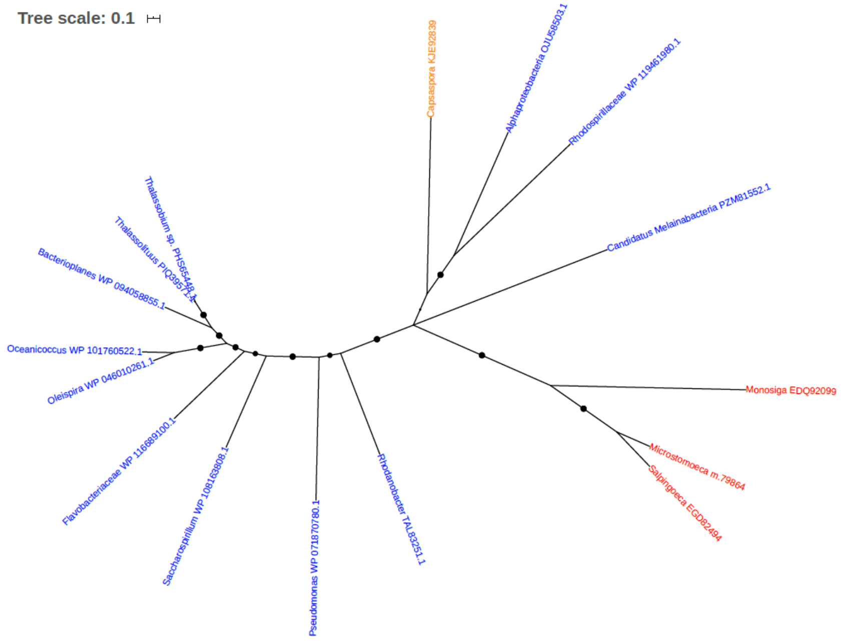


Figure S18 Phylogeny of Acyl-CoA dehydrogenase (EGD82494)

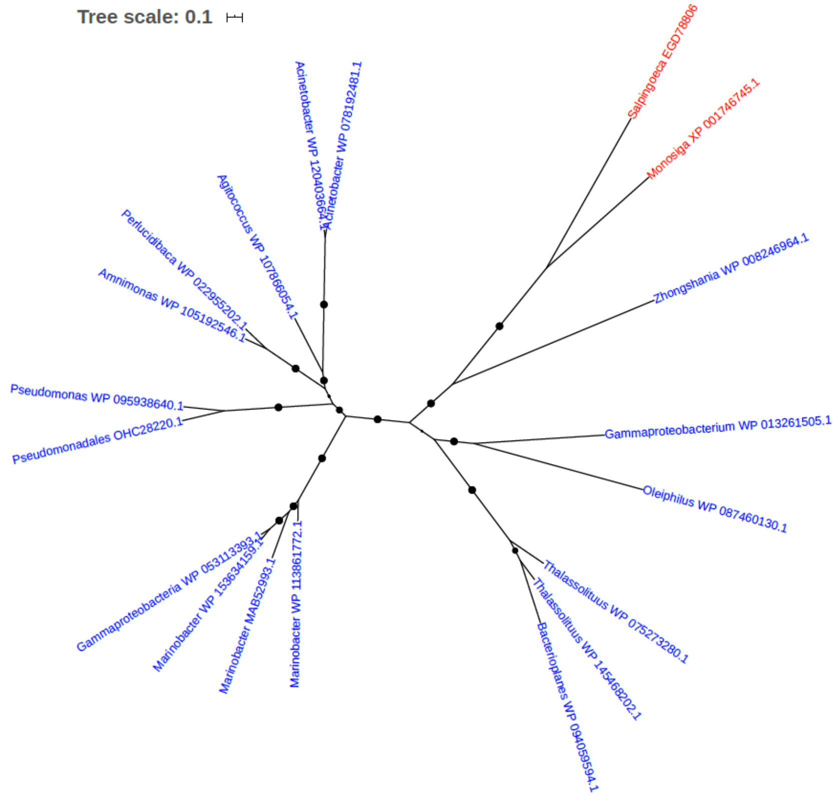


Figure S19. Phylogeny of Acyl-CoA dehydrogenase (EGD78806)

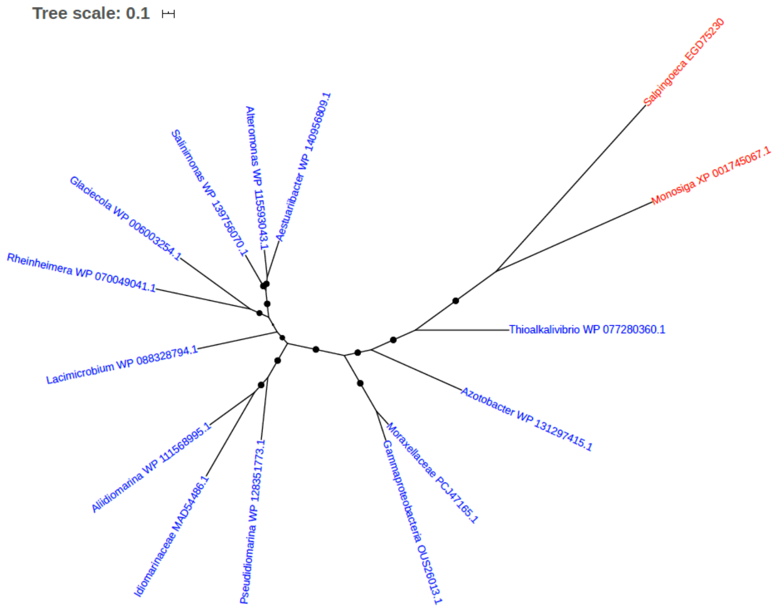


Figure S20. Phylogeny of Adenylosuccinate lyase (ASL) (EC 4.3.2.2) (Adenylosuccinase) (EGD75230)

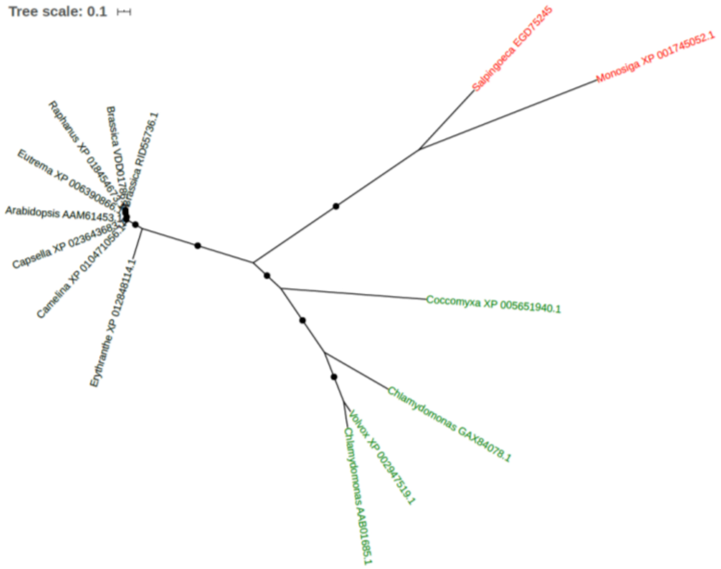


Figure S21. Phylogeny of Alanine aminotransferase (EGD75245)

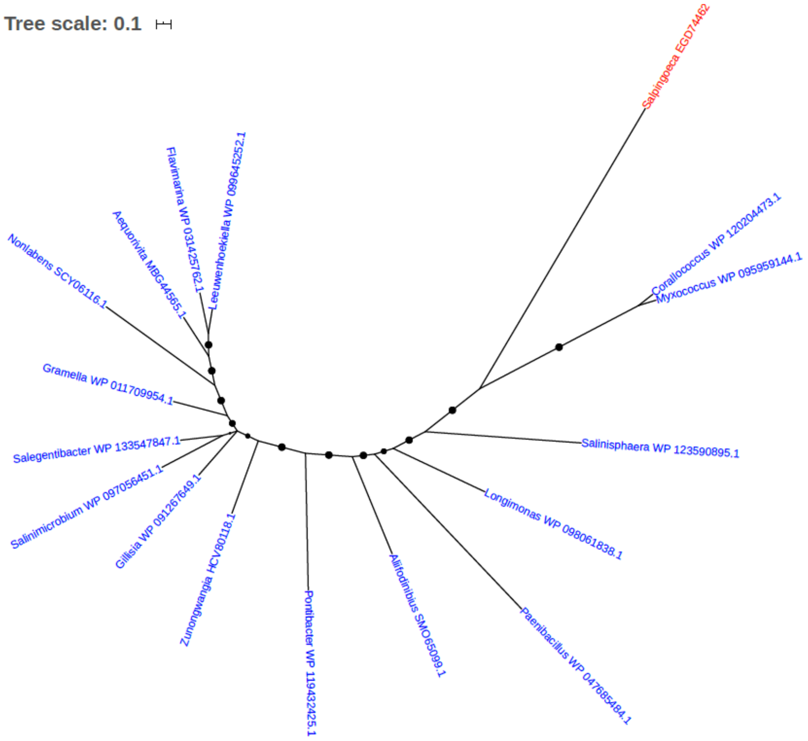


Figure S22. Phylogeny of Alcohol dehydrogenase (EGD74462)

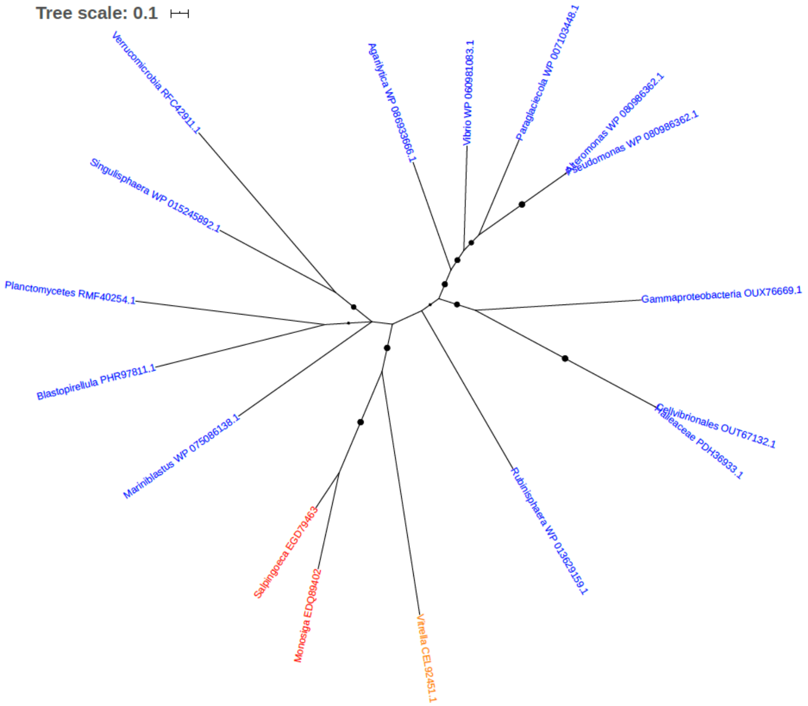


Figure S23. Phylogeny of Aldehyde reductase (EGD79463)

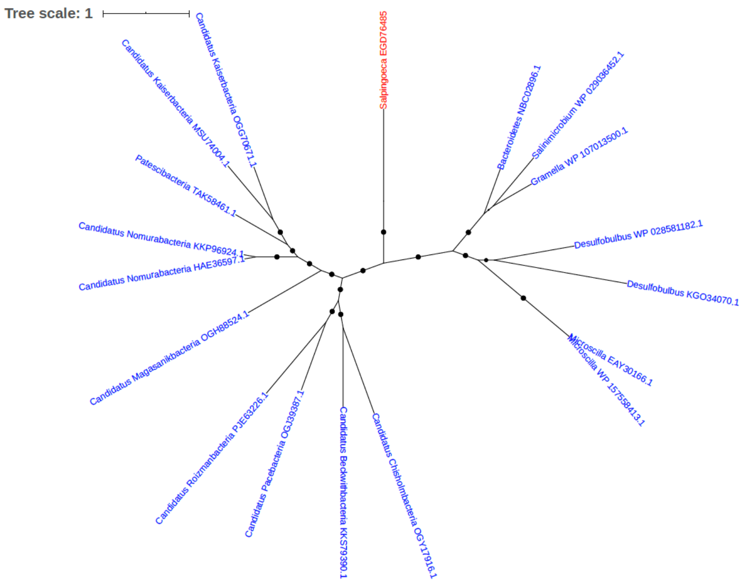


Figure S24. Phylogeny of Aldo/keto reductase (EGD76485)

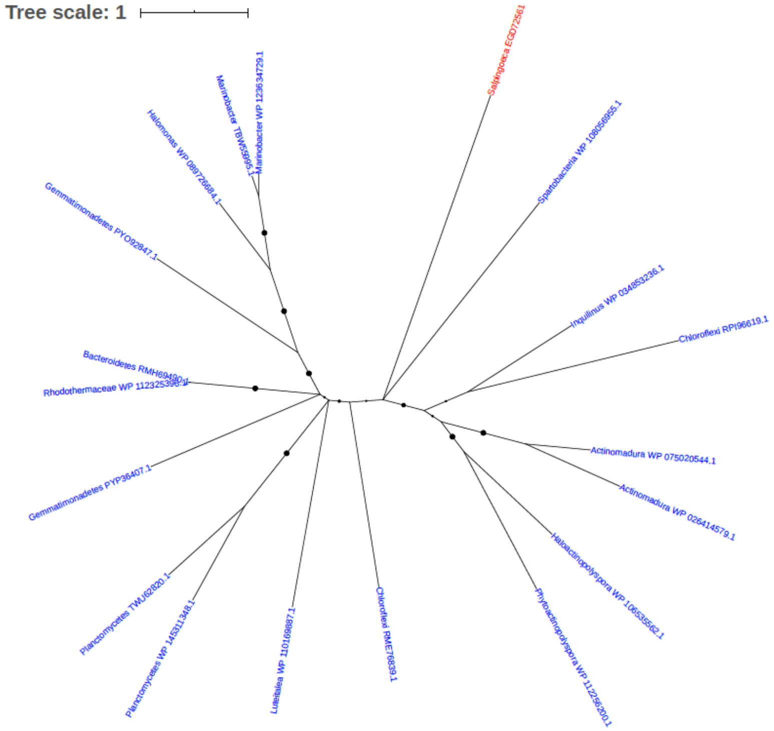


Figure S25. Phylogeny of Aldose 1-epimerase (EC 5.1.3.3) (Galactose mutarotase) (EGD72561)

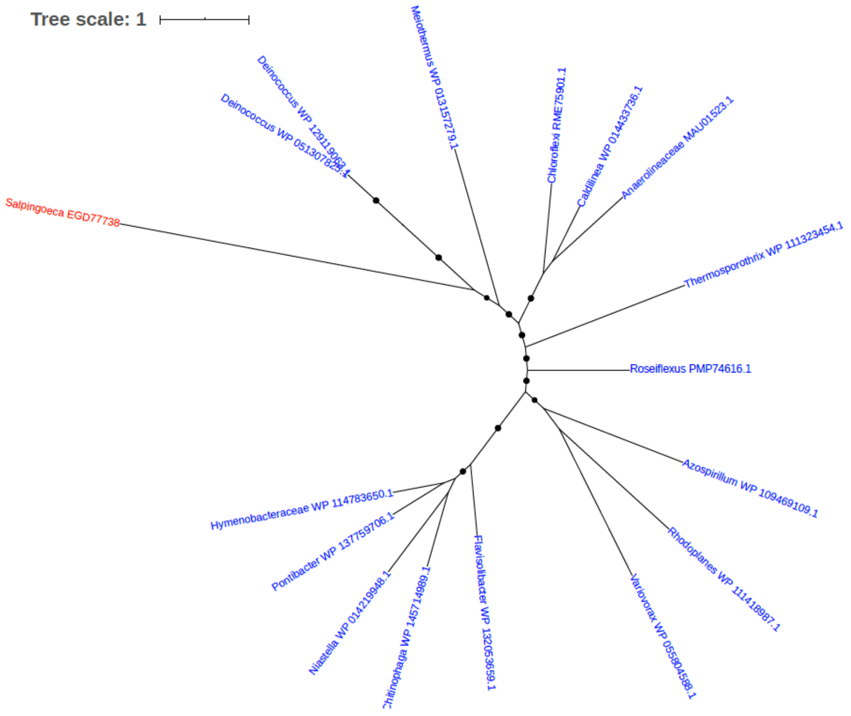


Figure S26. Phylogeny of Alpha amylase (EGD77738)

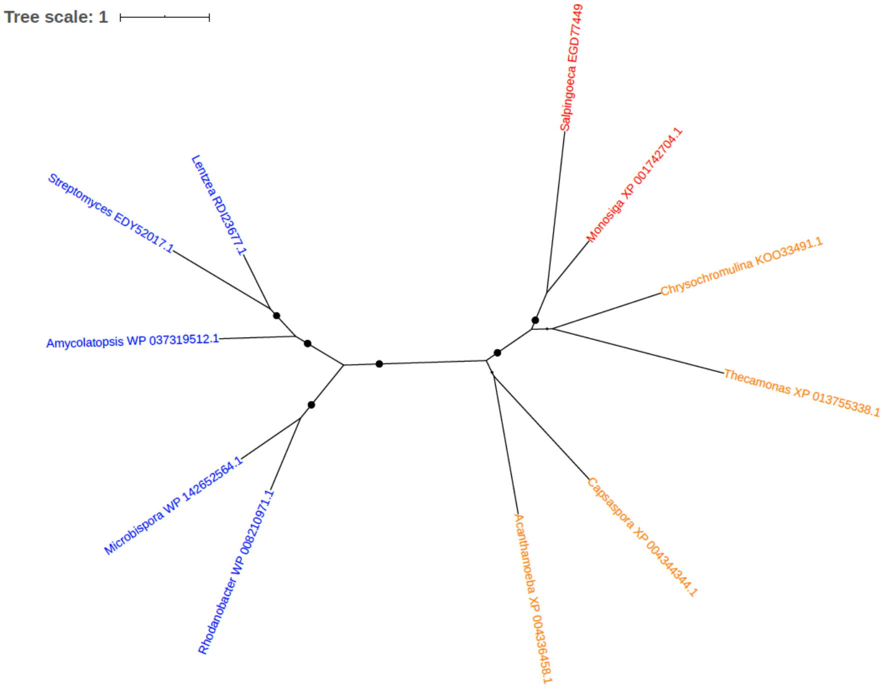


Figure S27. Phylogeny of Alpha-galactosidase (EC 3.2.1.22) (Melibiase) (EGD77449)

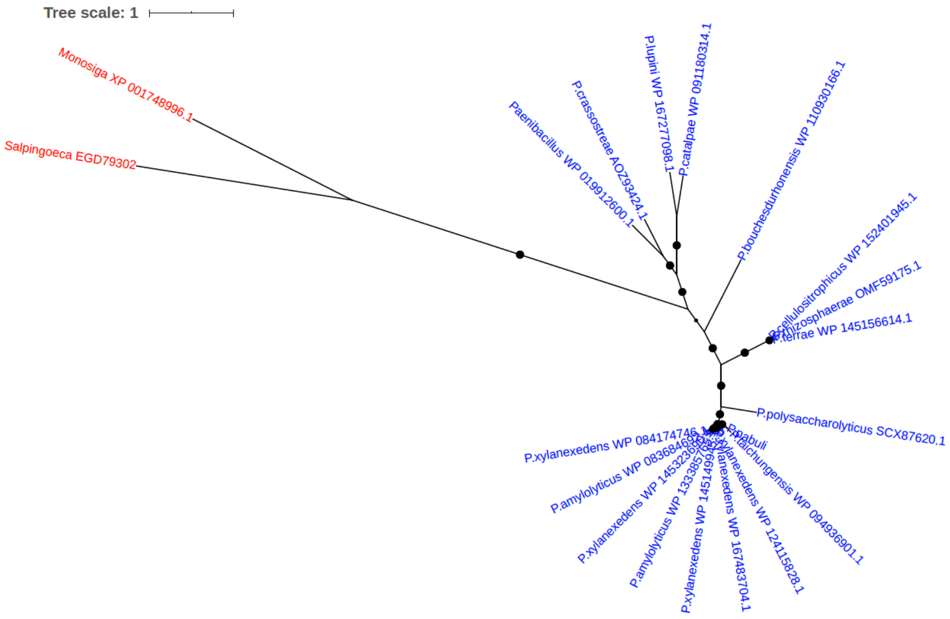


Figure S28. Phylogeny of Alpha-L-arabinofuranosidase B (EGD79302)

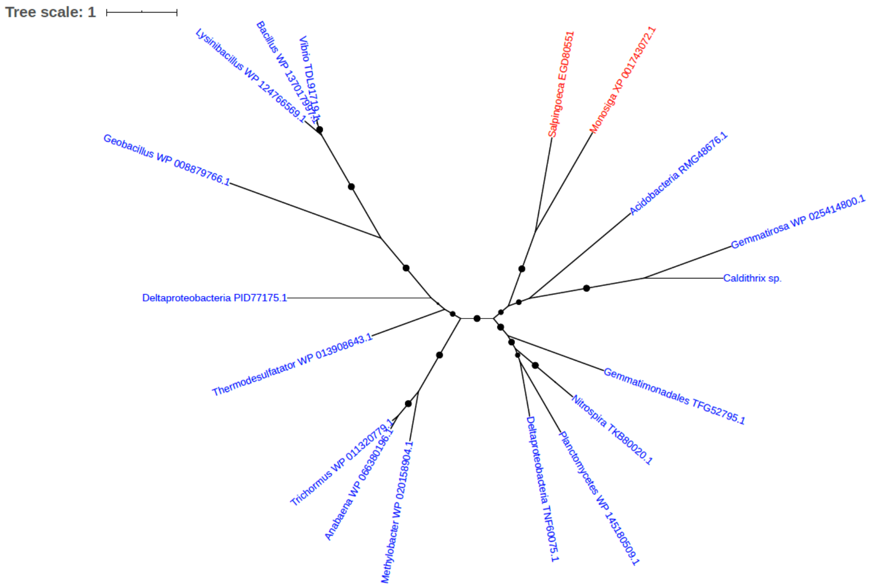


Figure S29. Phylogeny of Amidohydrolase (EGD80551)

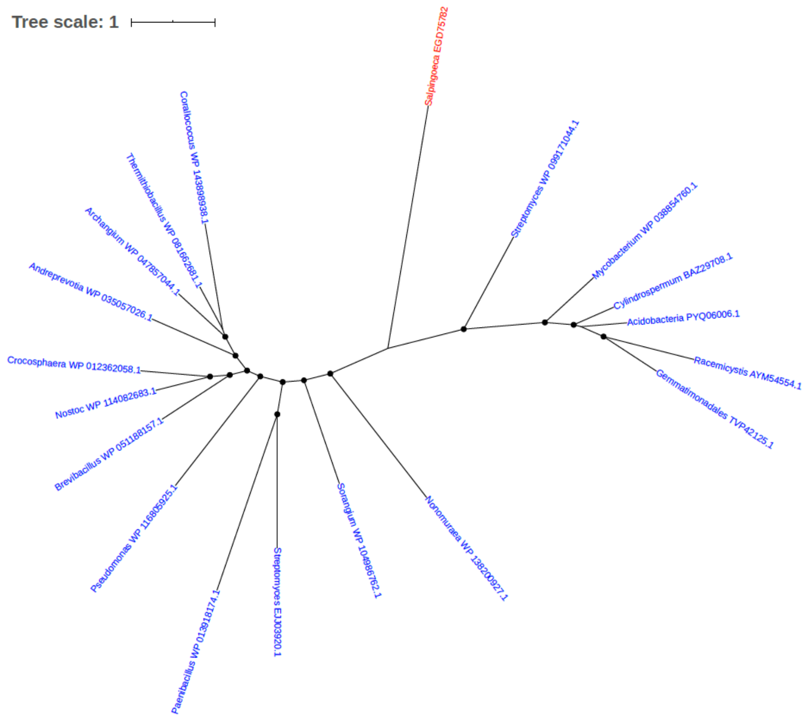


Figure S30. Phylogeny of Amino acid adenylation domain-containing protein (EGD75782)

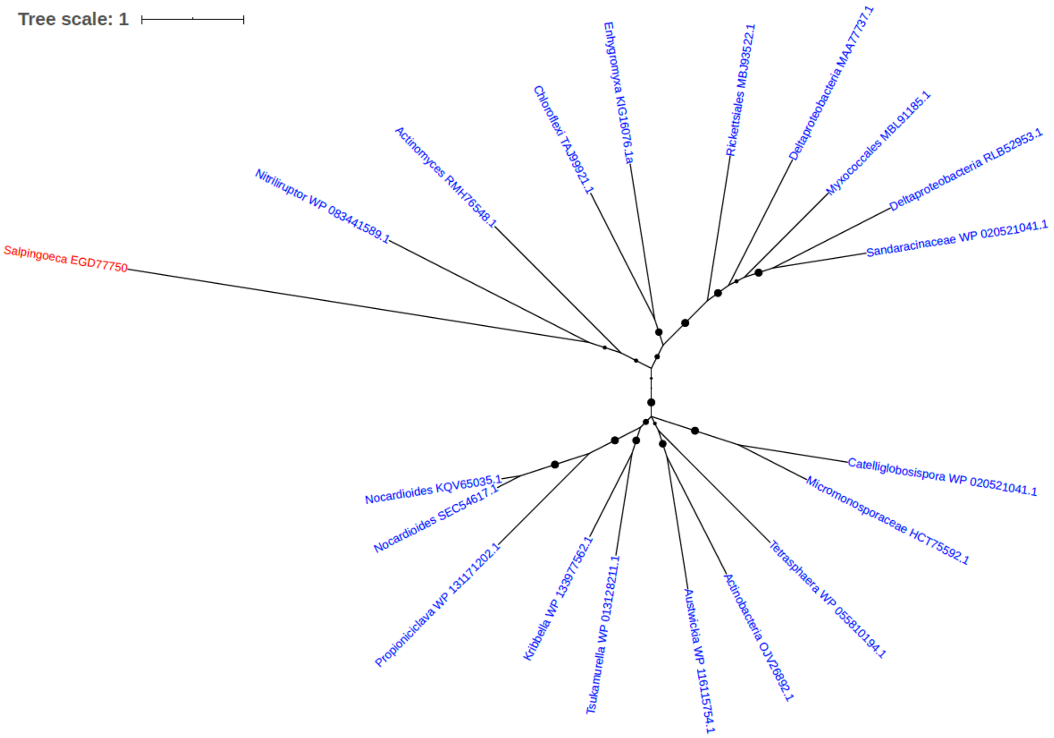


Figure S31. Phylogeny of Aminotransferase (EGD77750)

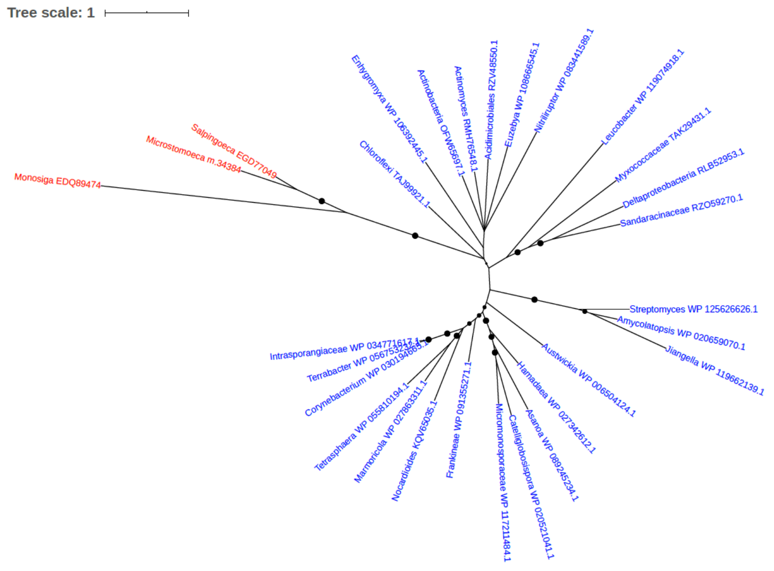

Figure S32. Phylogeny of Aminotransferase (EGD77049)

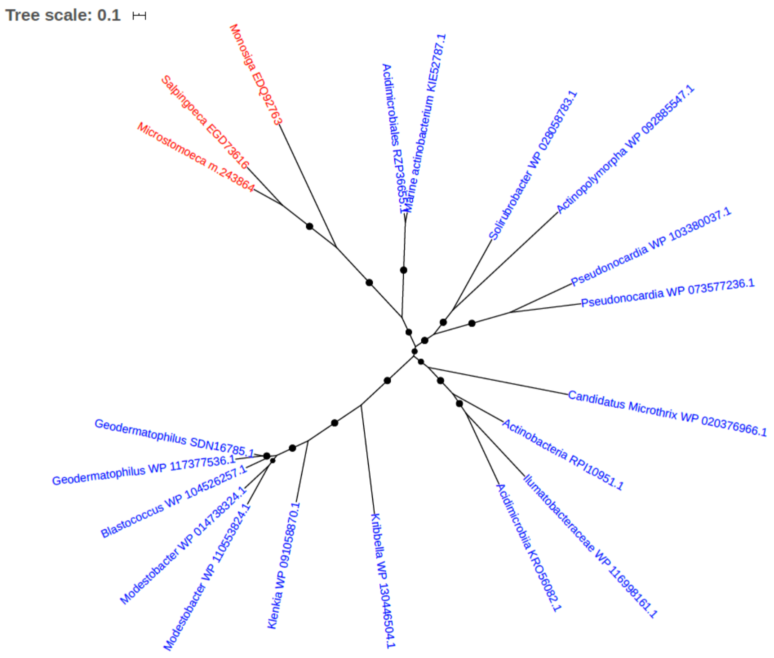


Figure S33. Phylogeny of AMP-dependent synthetase and ligase (EGD73616)

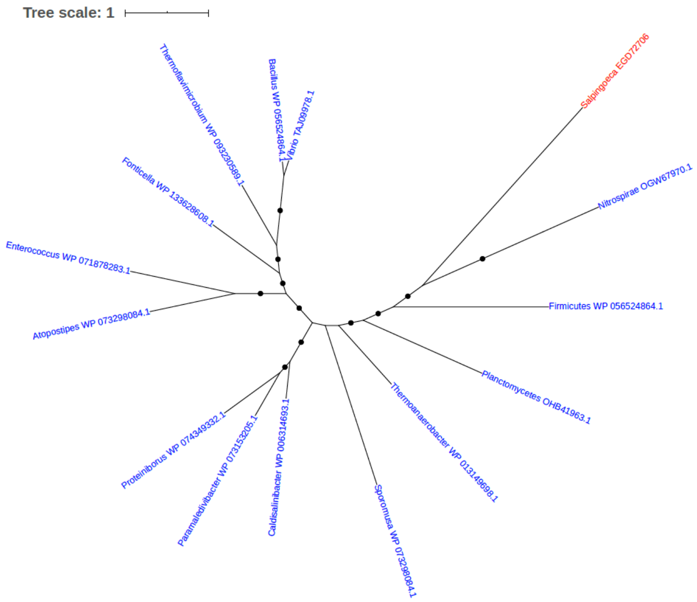


Figure S34. Phylogeny of Anhydro-N-acetylmuramic acid kinase (EGD72706)

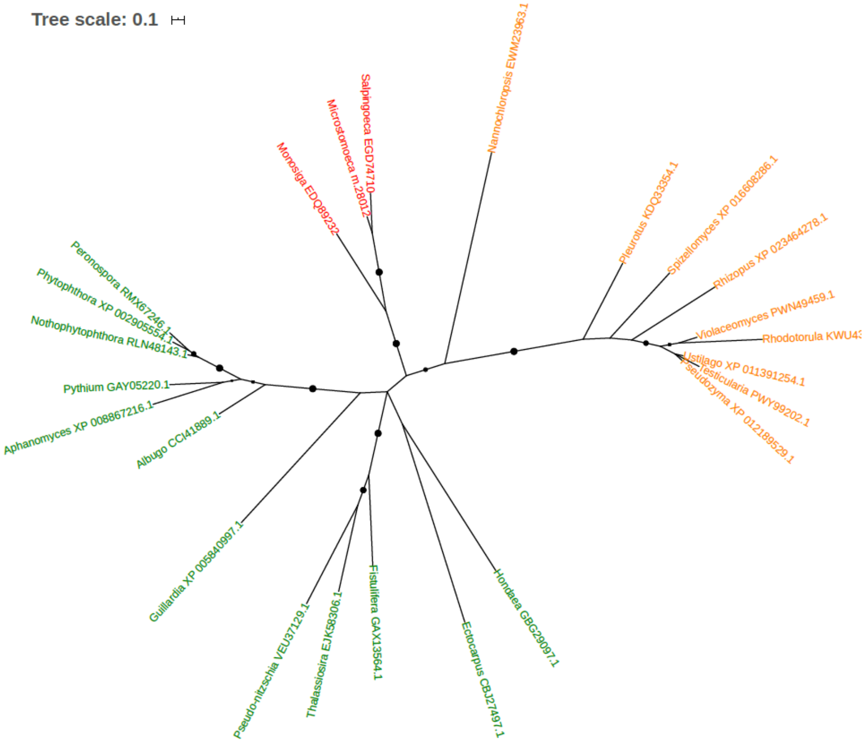


Figure S35. Phylogeny of Arginase (EC 3.5.3.1) (EGD74710)

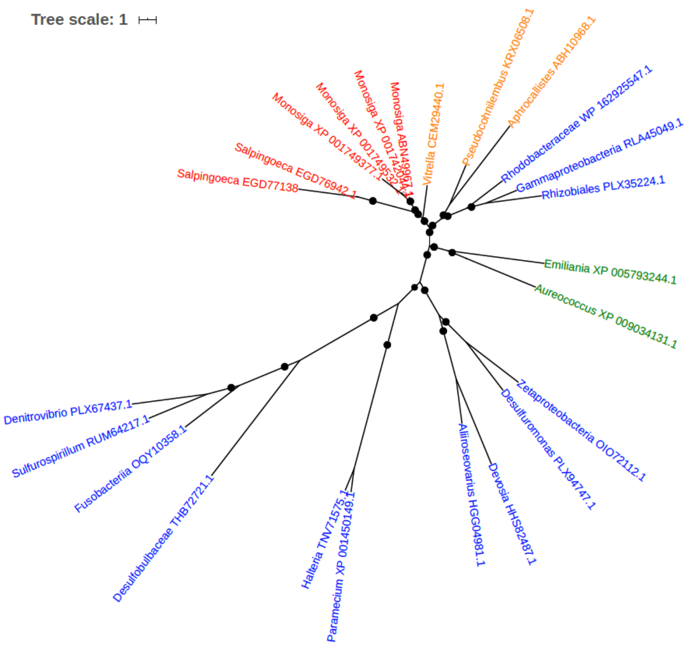


Figure S36. Phylogeny of Arginine kinase (EGD77138, EGD76942)

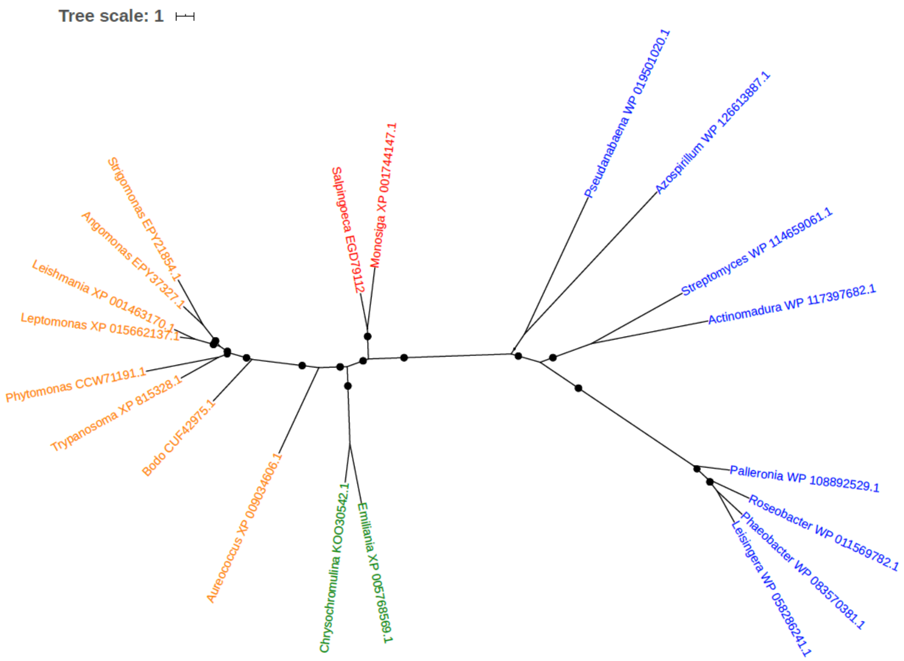


Figure S37. Phylogeny of Arginine N-methyltransferase (EGD79112)

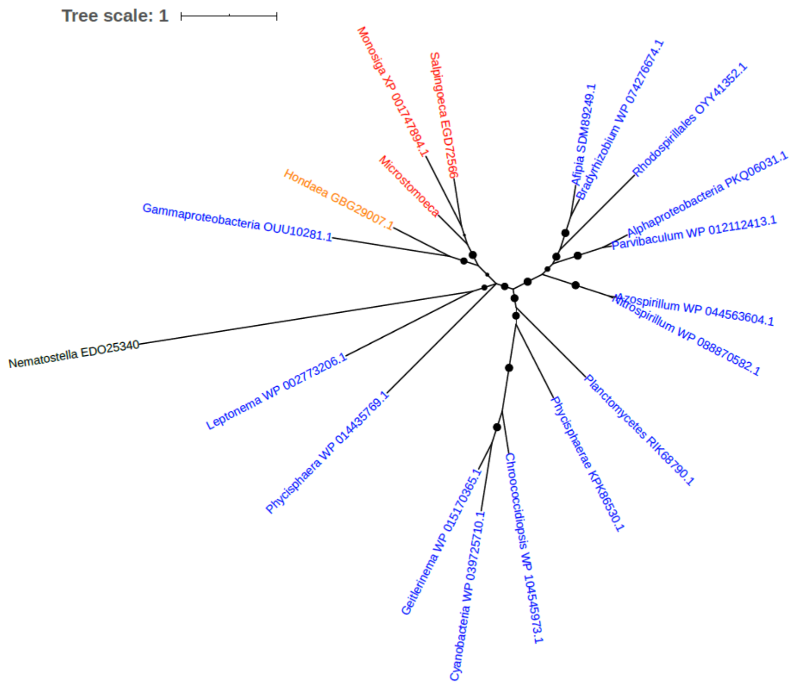


Figure S38. Phylogeny of Aspartate-semialdehyde dehydrogenase (EGD72566)

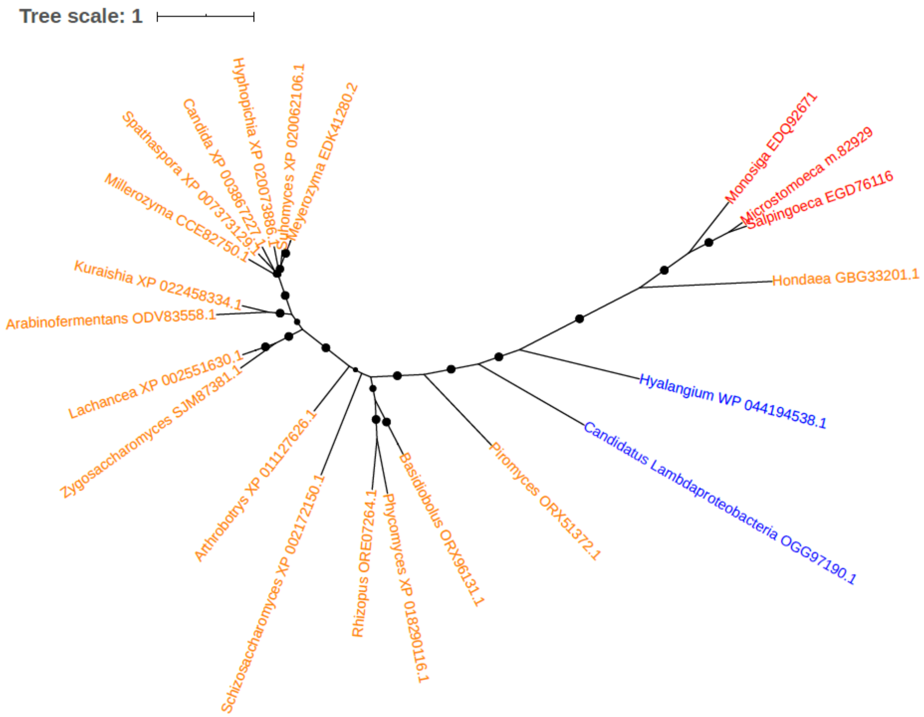


Figure S39. Phylogeny of Aspartokinase (EGD76112)

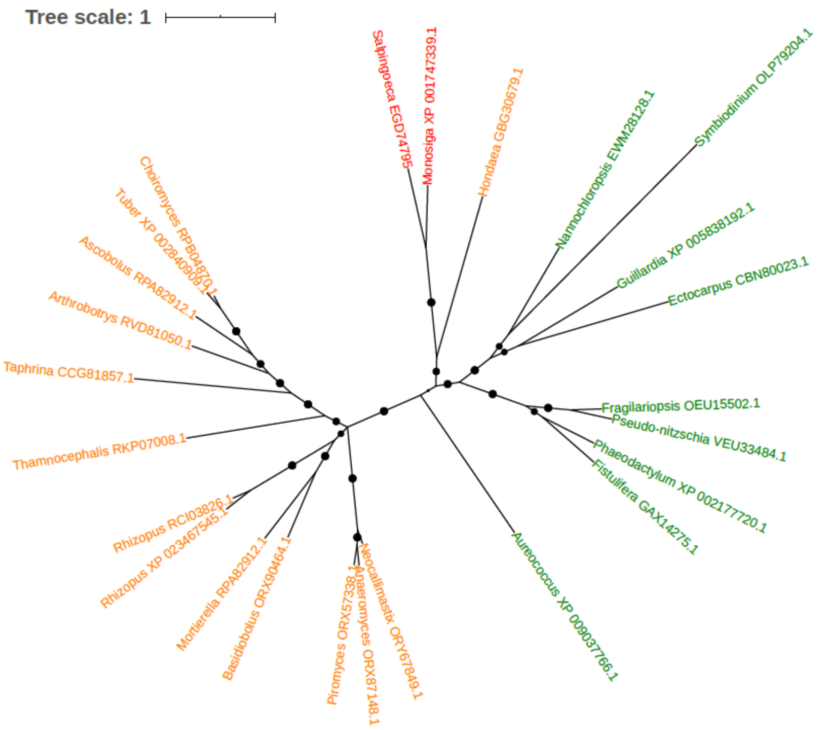


Figure S40. Phylogeny of ATP phosphoribosyltransferase (EGD74795)

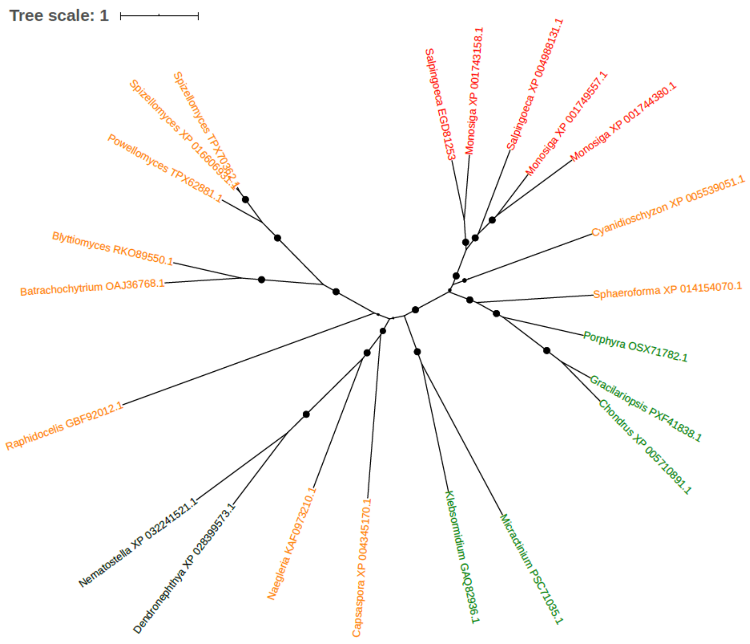


Figure S41. Phylogeny of ATP-binding cassette transporter G2 (EGD81253)

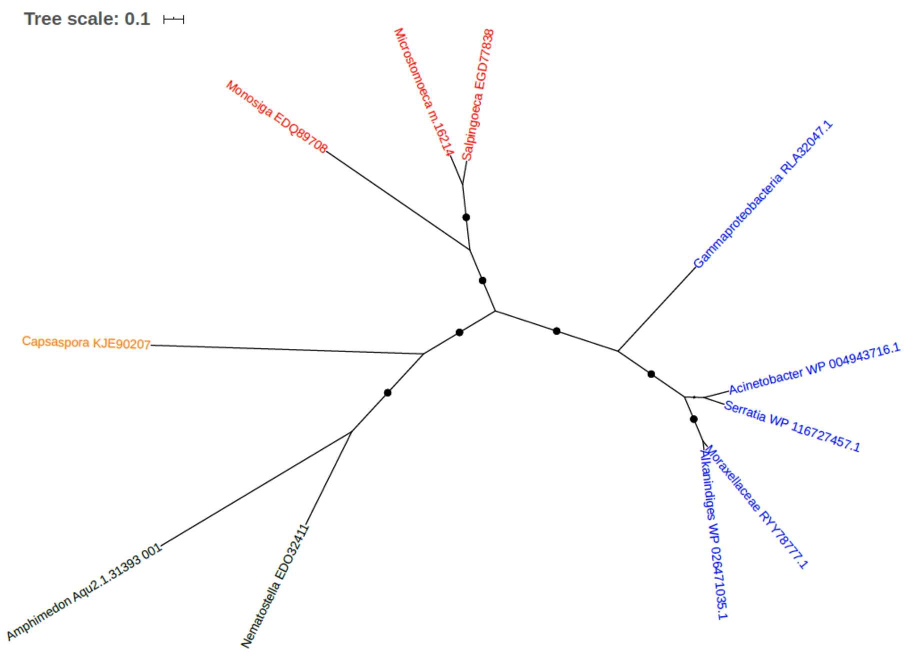


Figure S42. Phylogeny of ATP-dependent protease ATP-binding subunit (EGD77838)

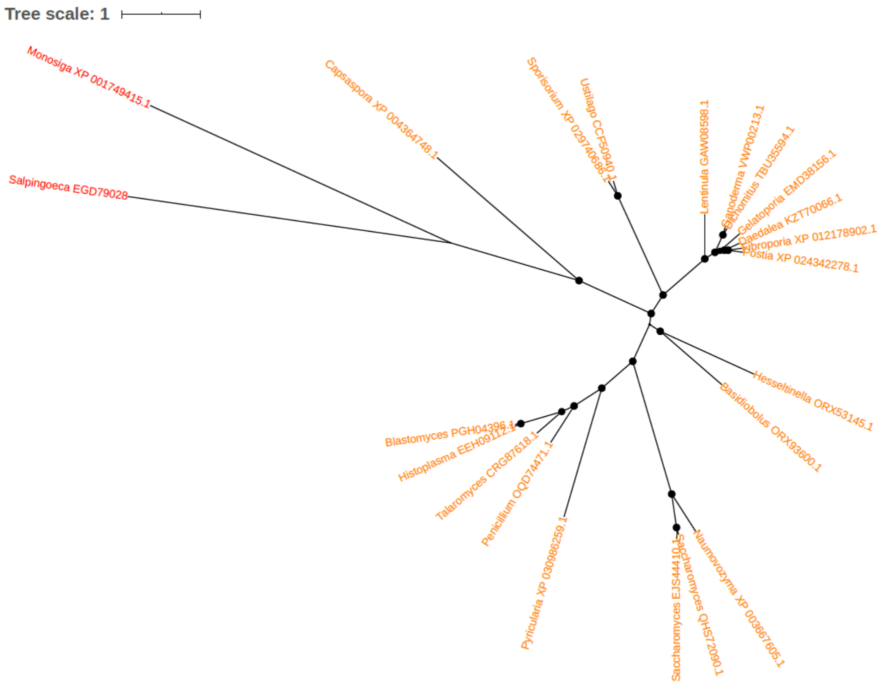


Figure S43. Phylogeny of Autophagy-related protein 9 (EGD79028)

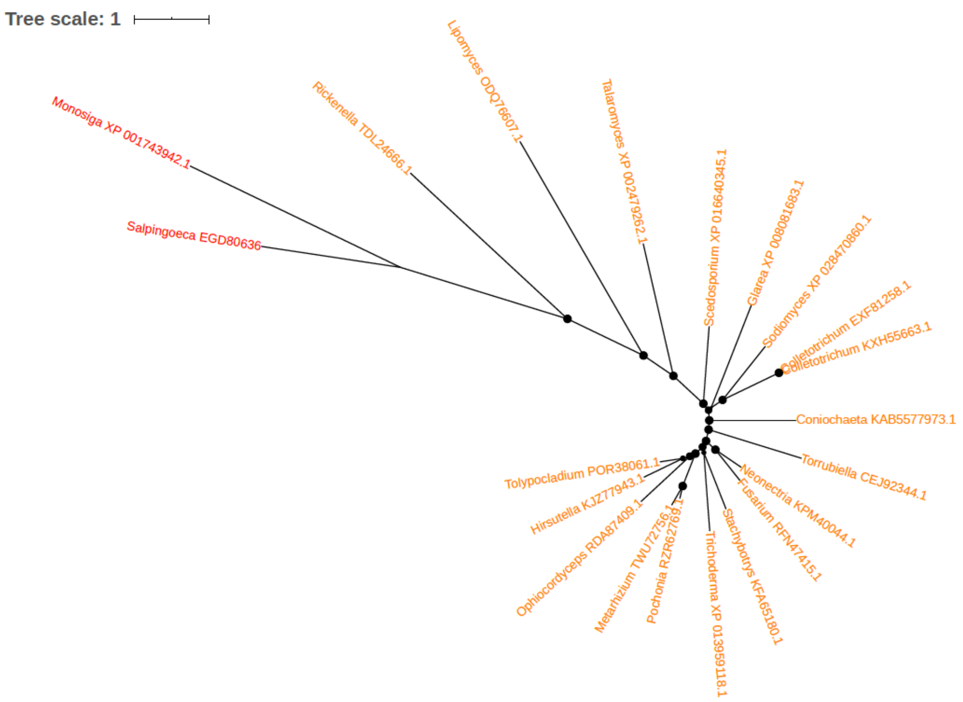


Figure S44. Phylogeny of Baculoviral IAP repeat-containing protein 5.1 (EGD80636)

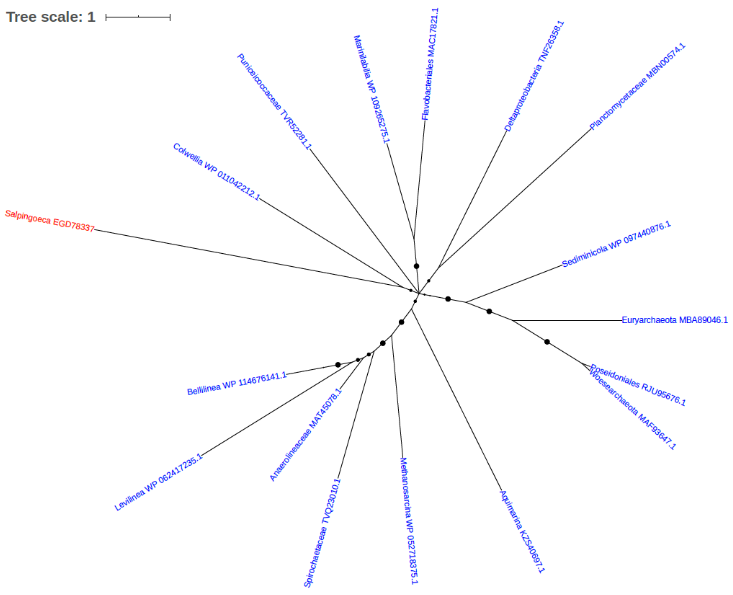


Figure S45. Phylogeny of Beta-Ig-H3/fasciclin (EGD78337)

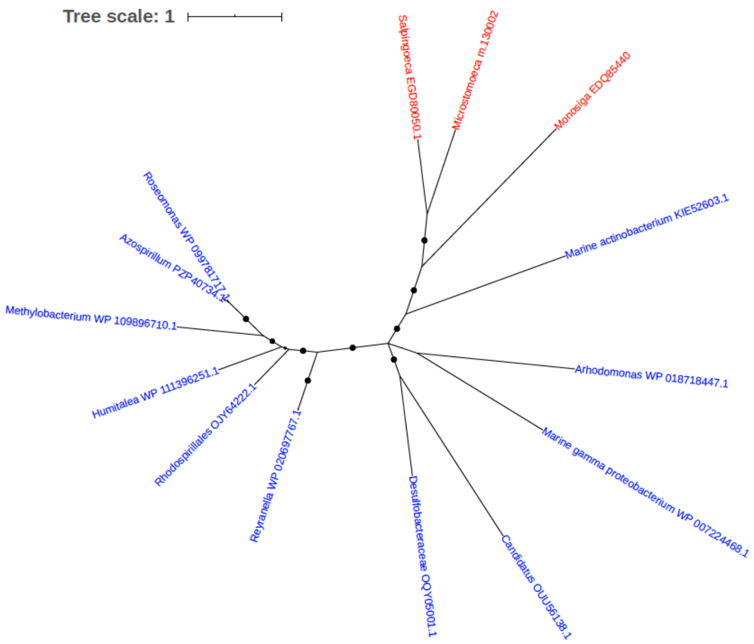


Figure S46. Phylogeny of Beta-lactamase (EGD80050)

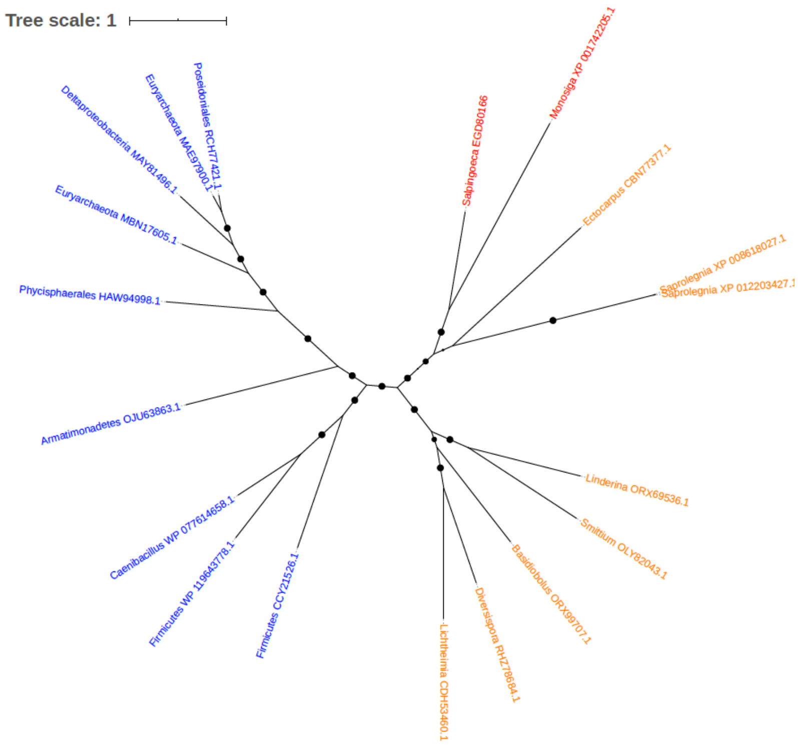


Figure S47. Phylogeny of Bleomycin hydrolase (EGD80166)

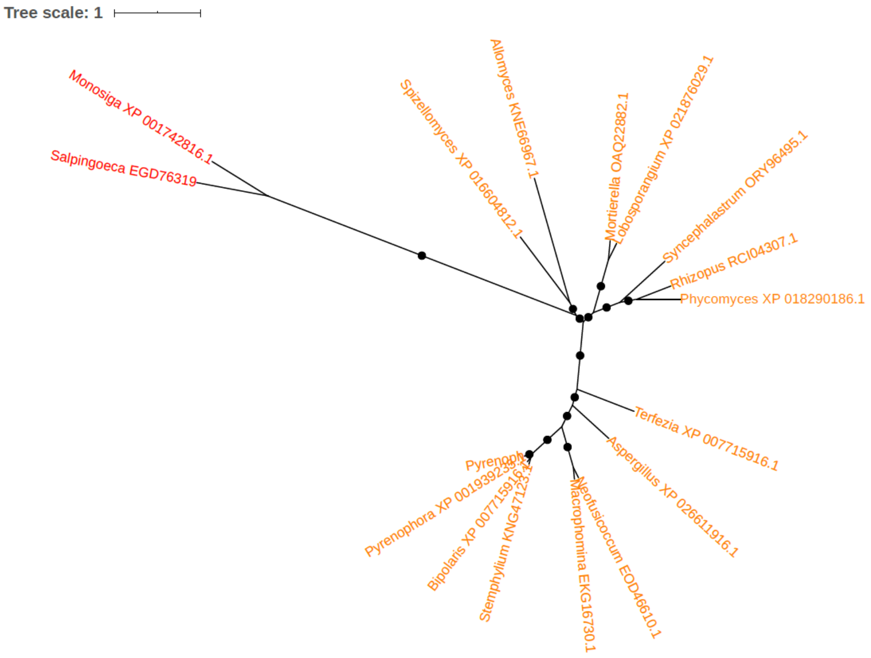


Figure S48. Phylogeny of Carbamylphosphate synthetase 1 (EGD76319)

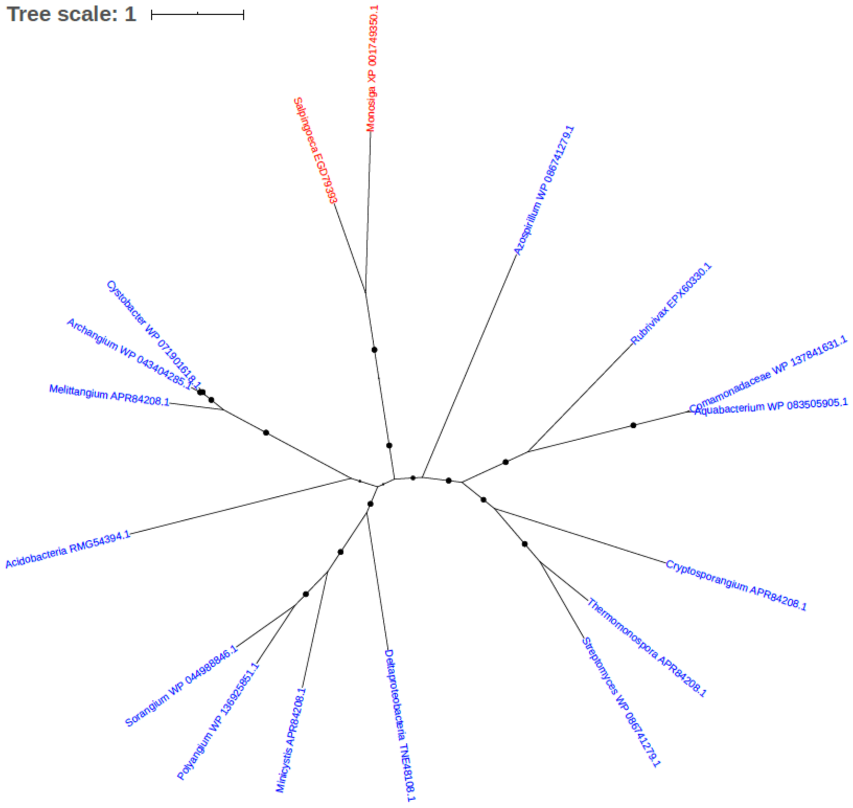


Figure S49. Phylogeny of Carboxylic ester hydrolase (EC 3.1.1.-) (EGD79393)

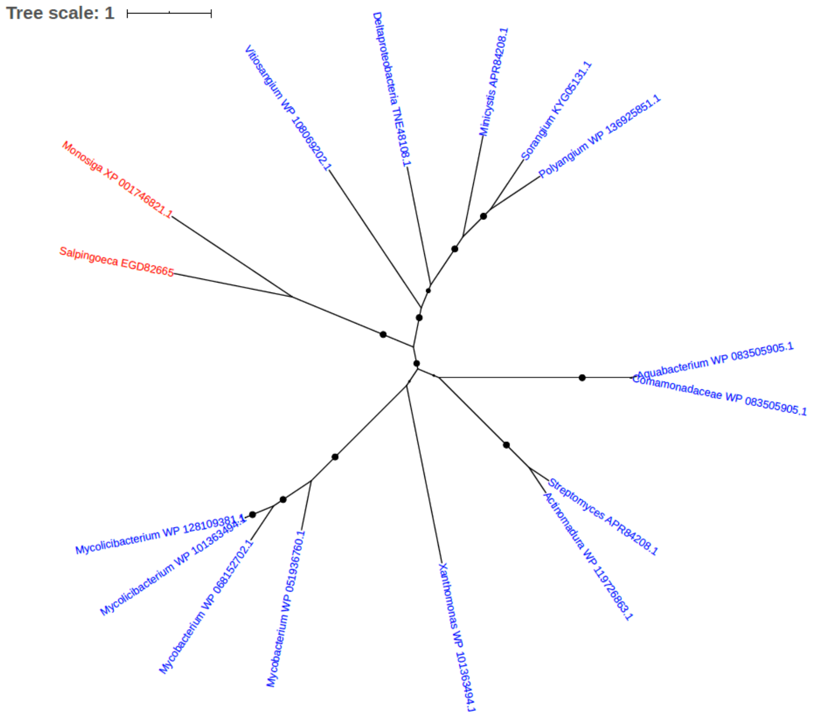


Figure S50. Phylogeny of Carboxylic ester hydrolase (EC 3.1.1.-) (EGD82665)

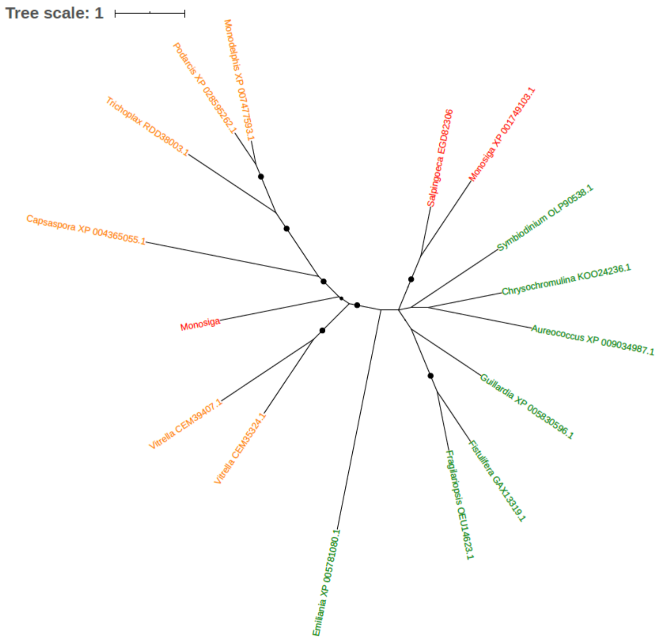


Figure S51. Phylogeny of Carboxypeptidase (EC 3.4.16.-) (EGD82306)

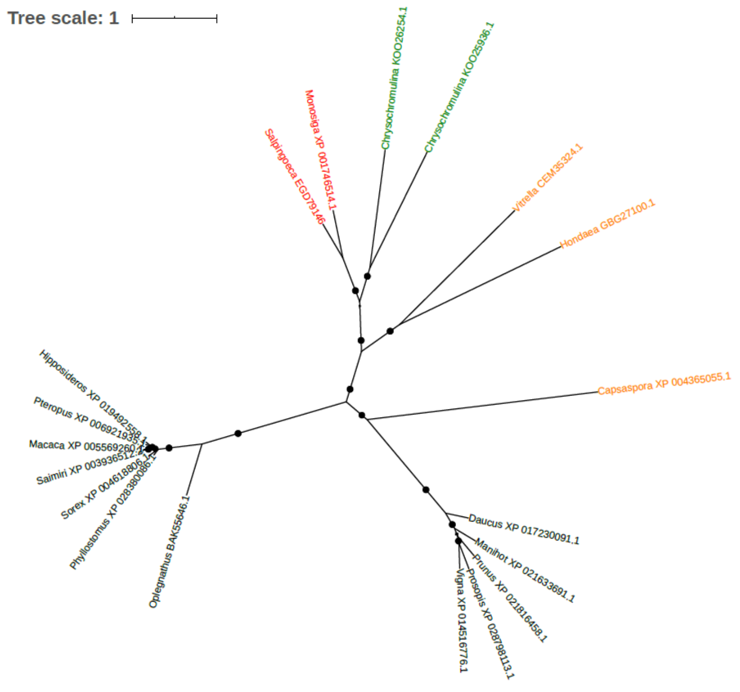


Figure S52. Phylogeny of Carboxypeptidase (EC 3.4.16.-) (EGD79146)

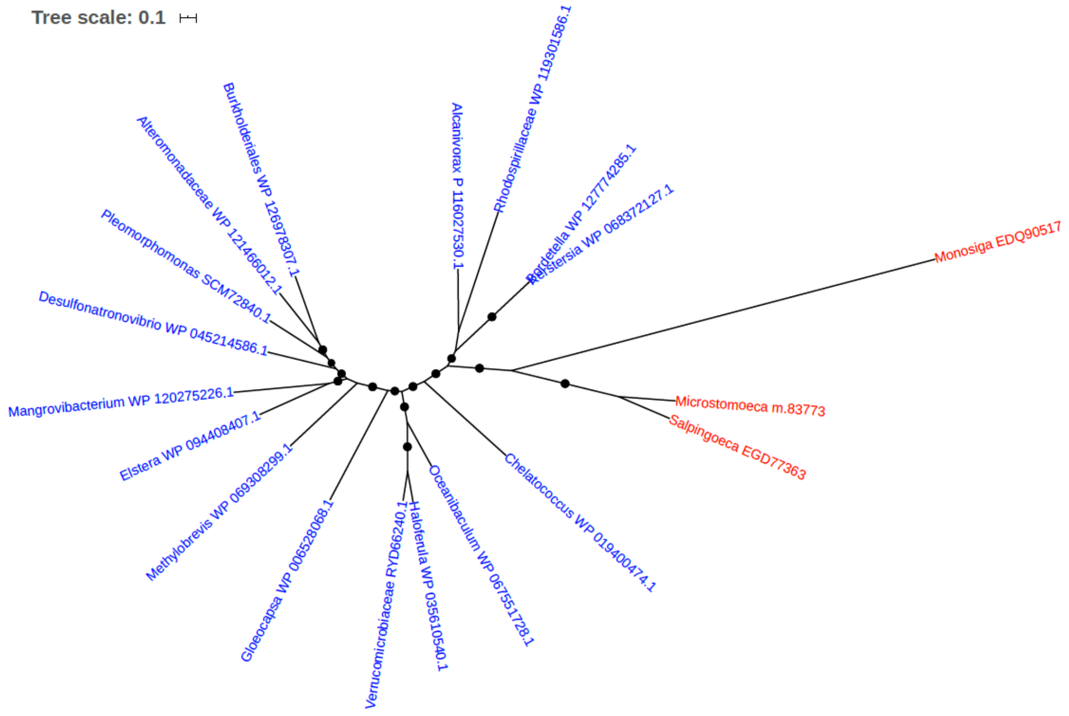


Figure S53. Phylogeny of Catalase (EC 1.11.1.6) (EGD77363)

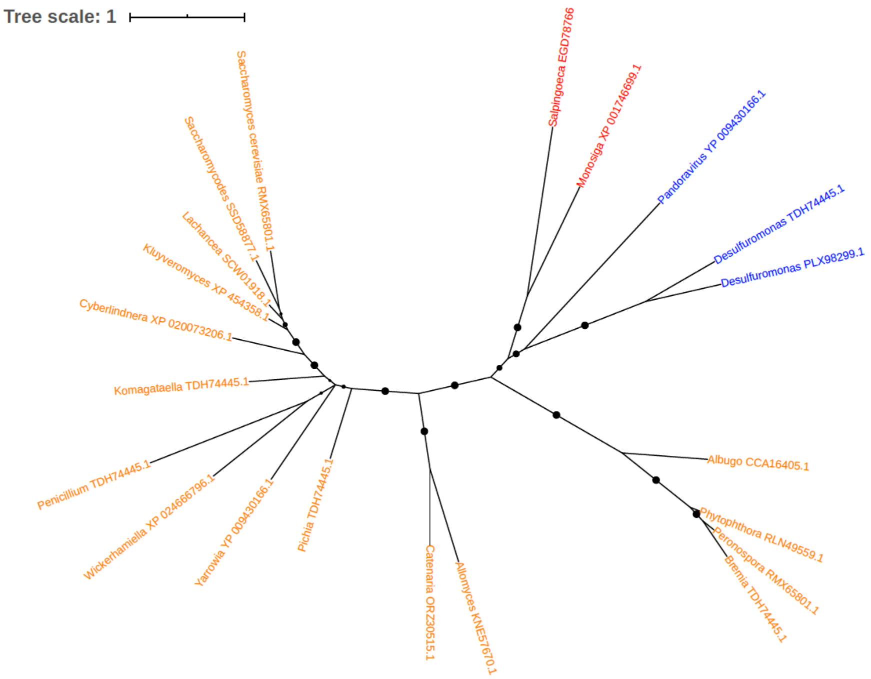


Figure S54. Phylogeny of Chorismate synthase (EGD78766)

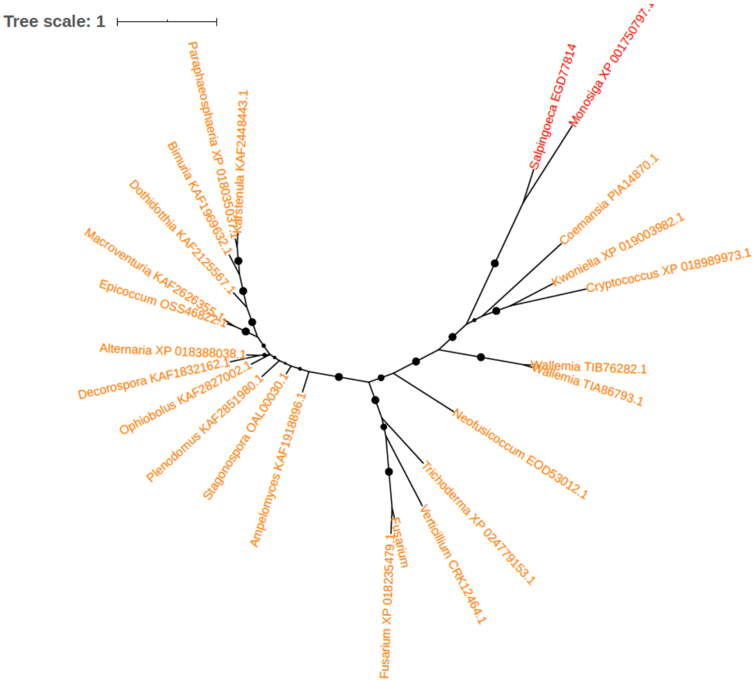


Figure S55. Phylogeny of CHY zinc finger domain-containing protein (EGD77814)

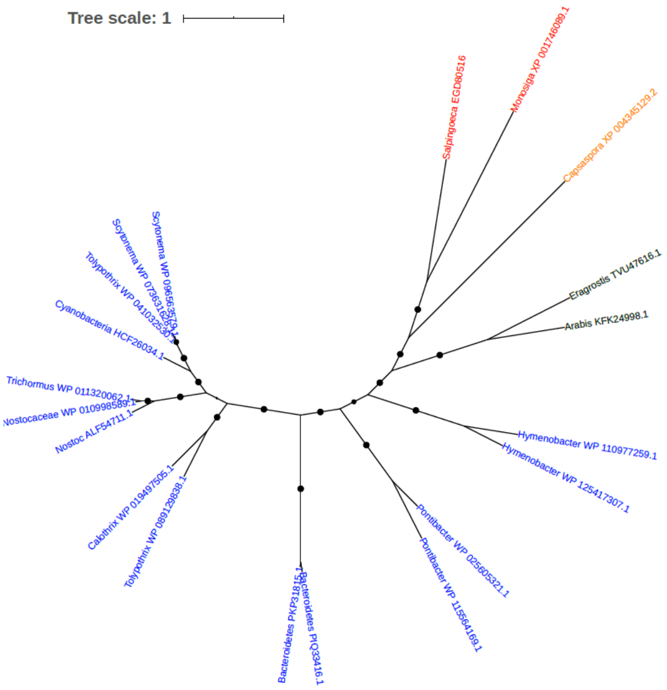


Figure S56. Phylogeny of Cinnamyl-alcohol dehydrogenase family/CAD family (EGD80516)

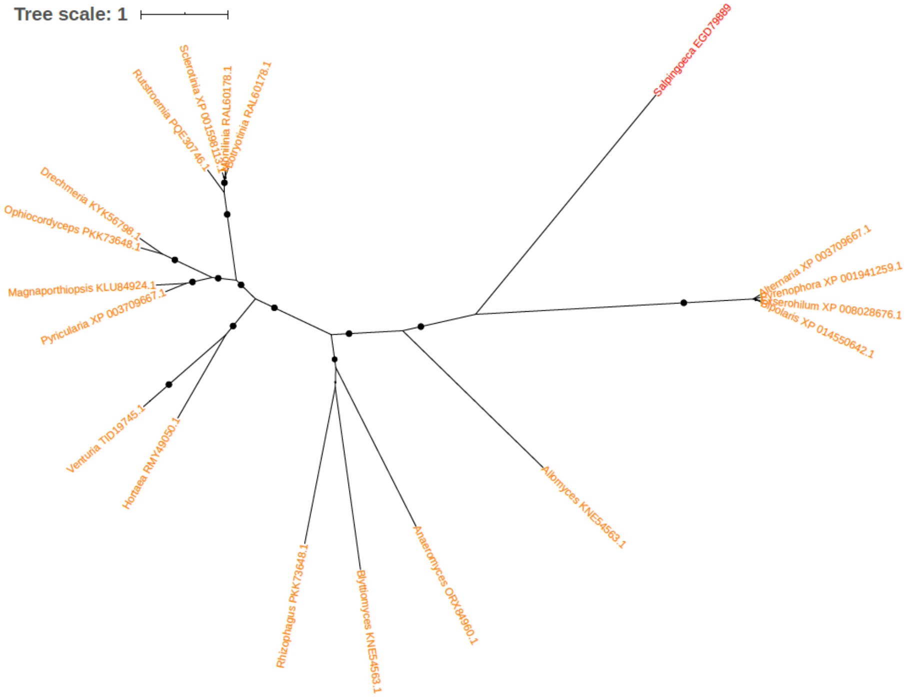


Figure S57. Phylogeny of CMGC/CDK protein kinase (EGD79889)

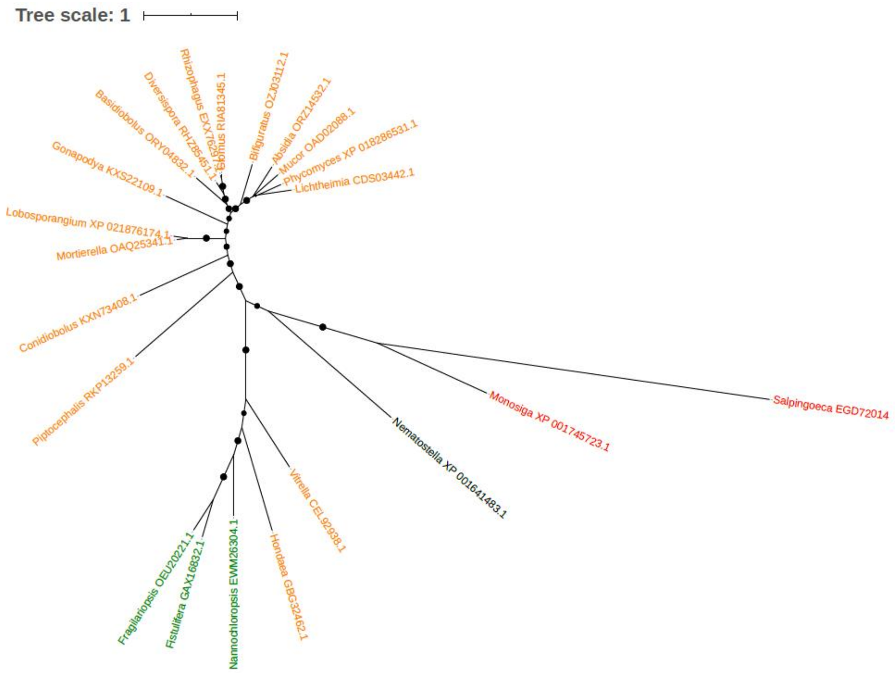


Figure S58. Phylogeny of CMGC/CDK/CDK5 protein kinase (EGD72014)

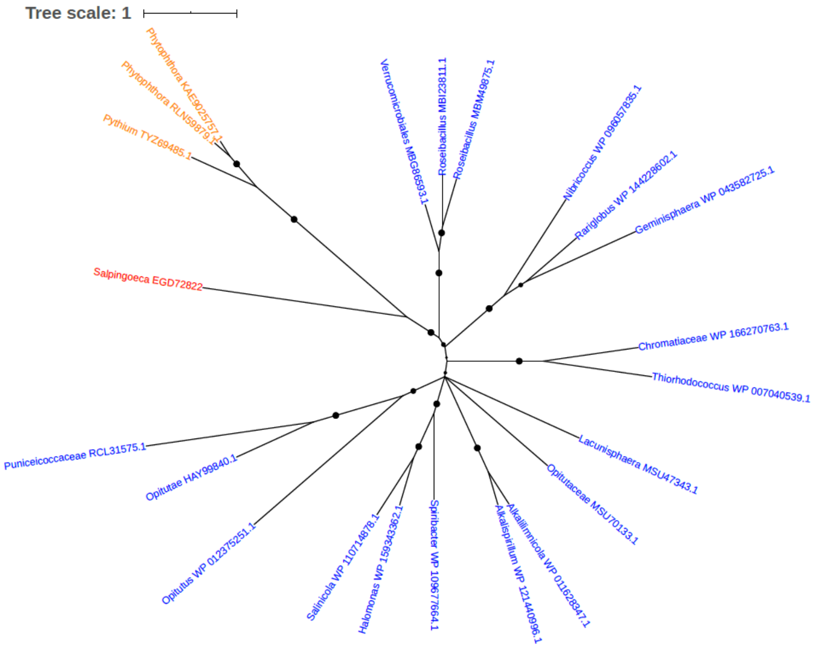


Figure S59. Phylogeny of Cobalamin biosynthesis CbiX protein (EGD72822)

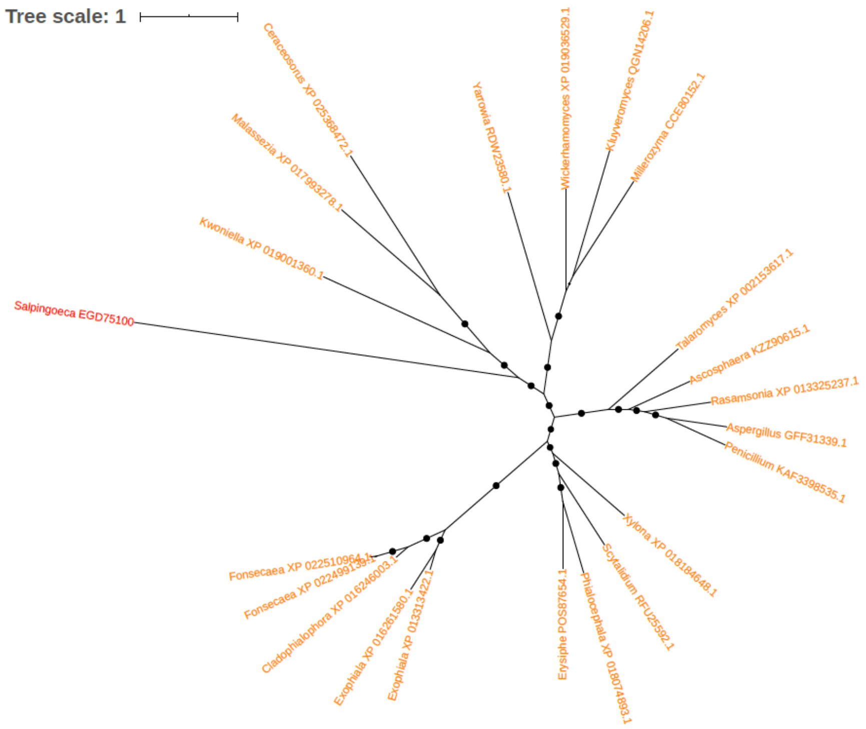


Figure S60. Phylogeny of CobW/P47K family protein (EGD75100)

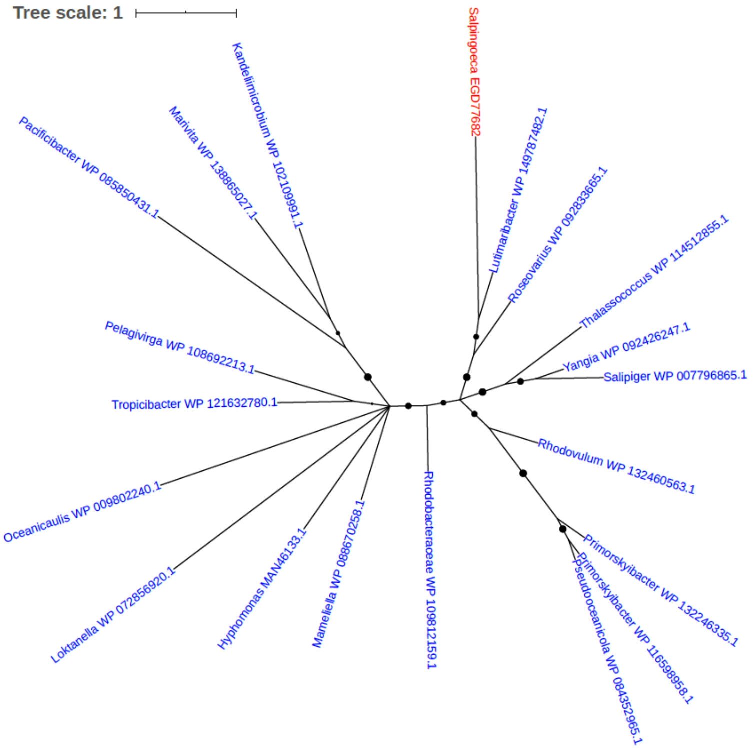


Figure S61. Phylogeny of Cobyric acid synthase (EGD77682)

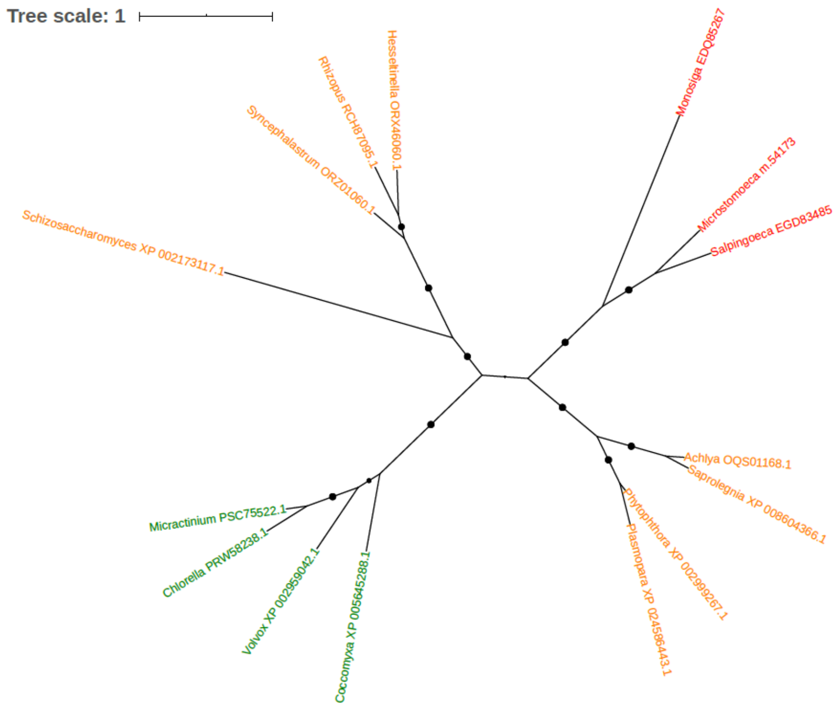


Figure S62. Phylogeny of Cysteine synthase (EGD82485)

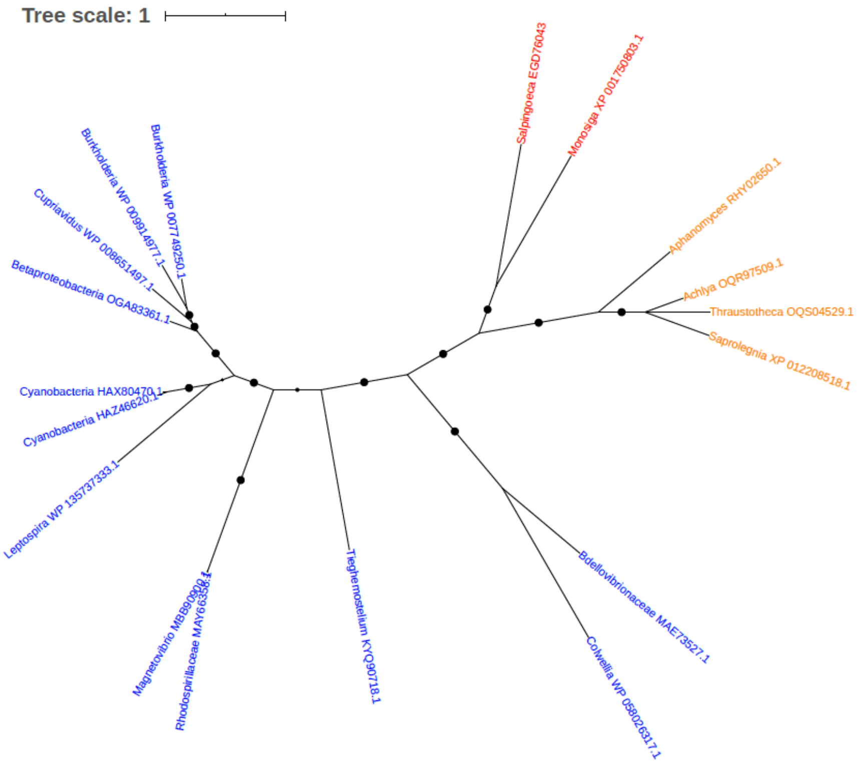


Figure S63. Phylogeny of Cysteine synthase (EGD76043)

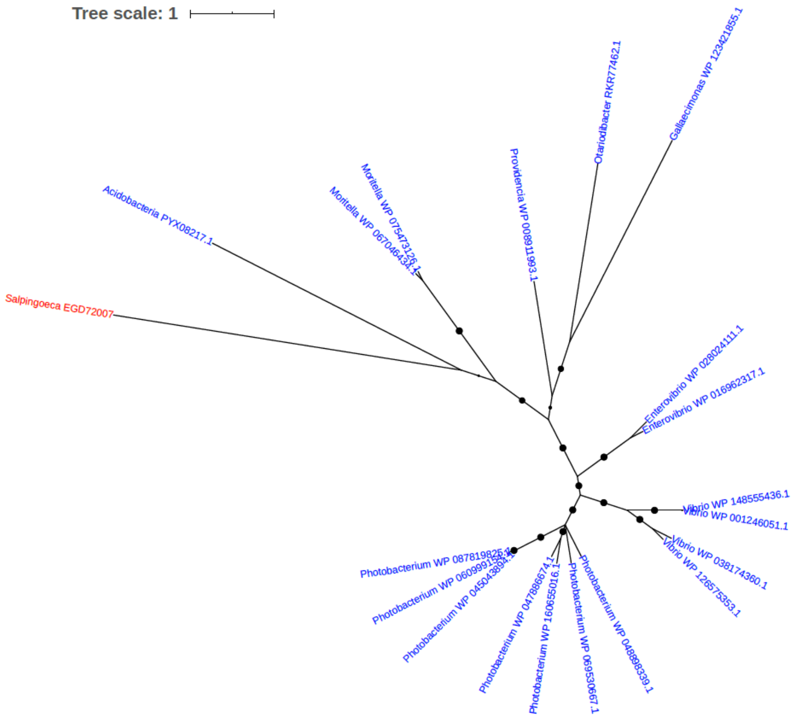


Figure S64. Phylogeny of Cytidine deaminase (EGD72007)

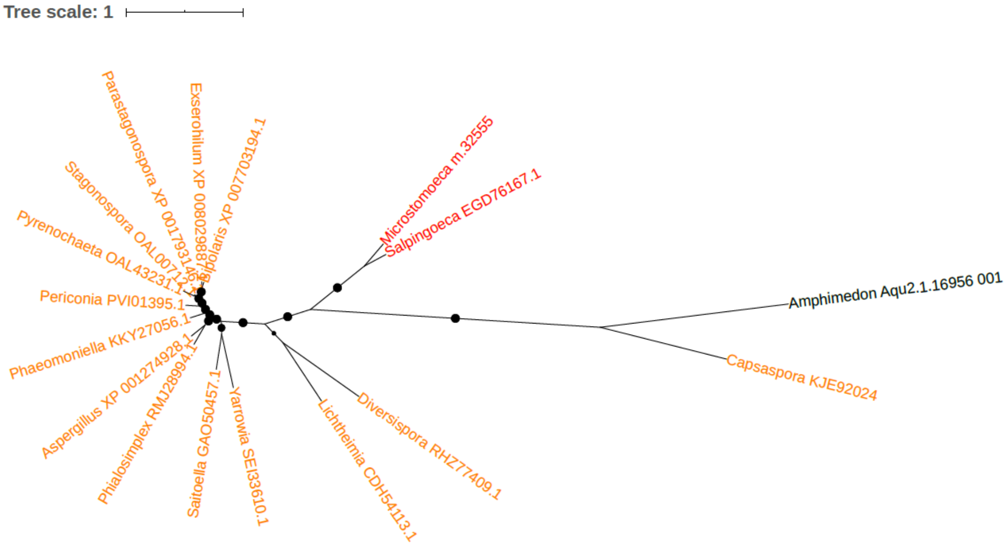


Figure S65. Phylogeny of Cytochrome b2 (EGD76167)

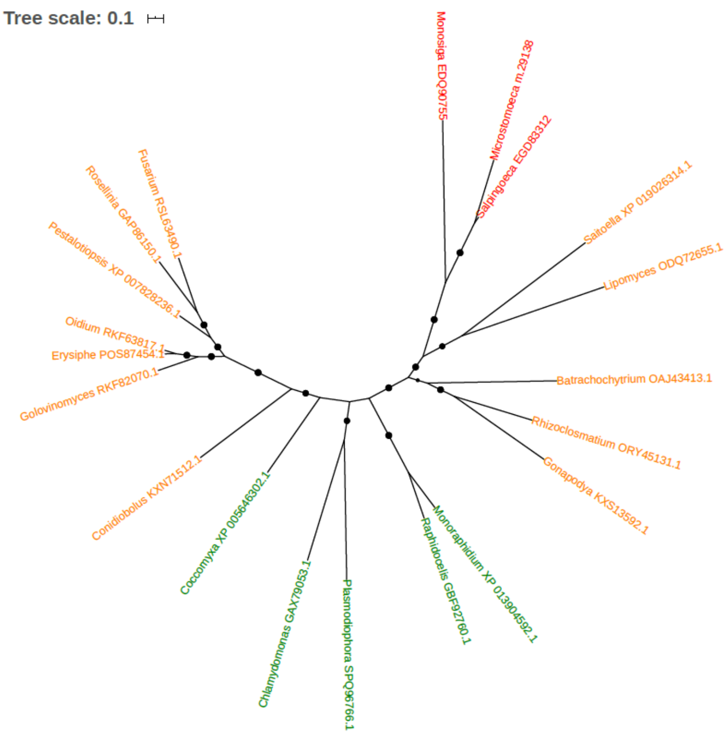


Figure S66. Phylogeny of Cytochrome c peroxidase (EGD83312)

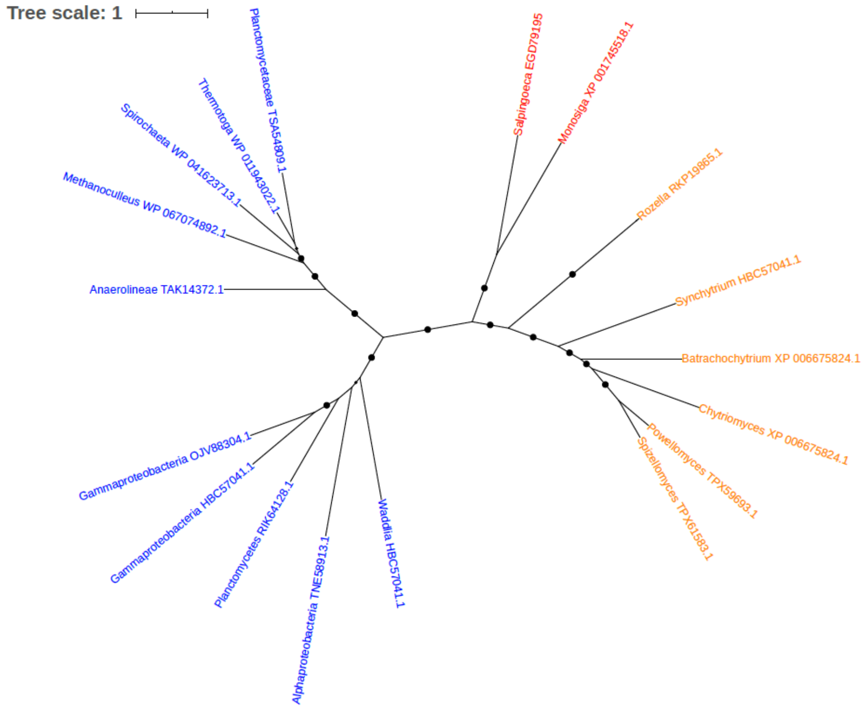


Figure S67. Phylogeny of D-alanine-D-alanine ligase (EGD79195)

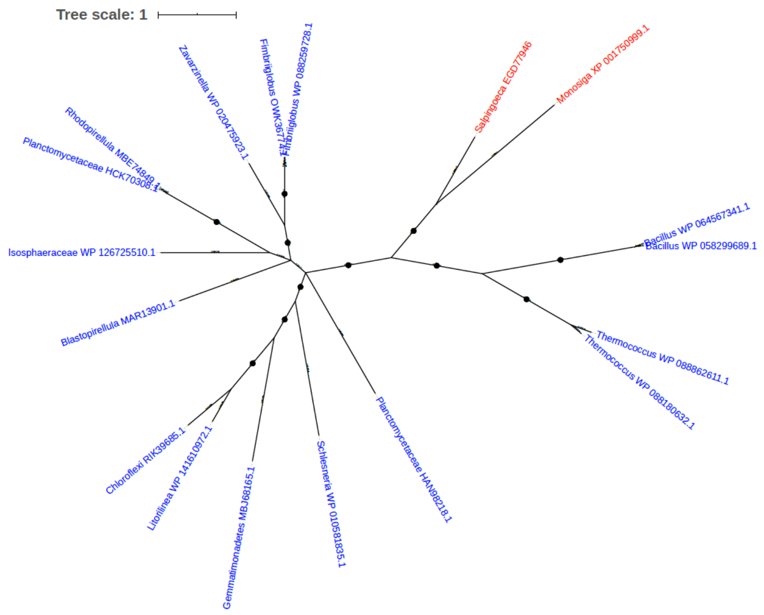


Figure S68. Phylogeny of D-isomer specific 2-hydroxyacid dehydrogenase (EGD77946)

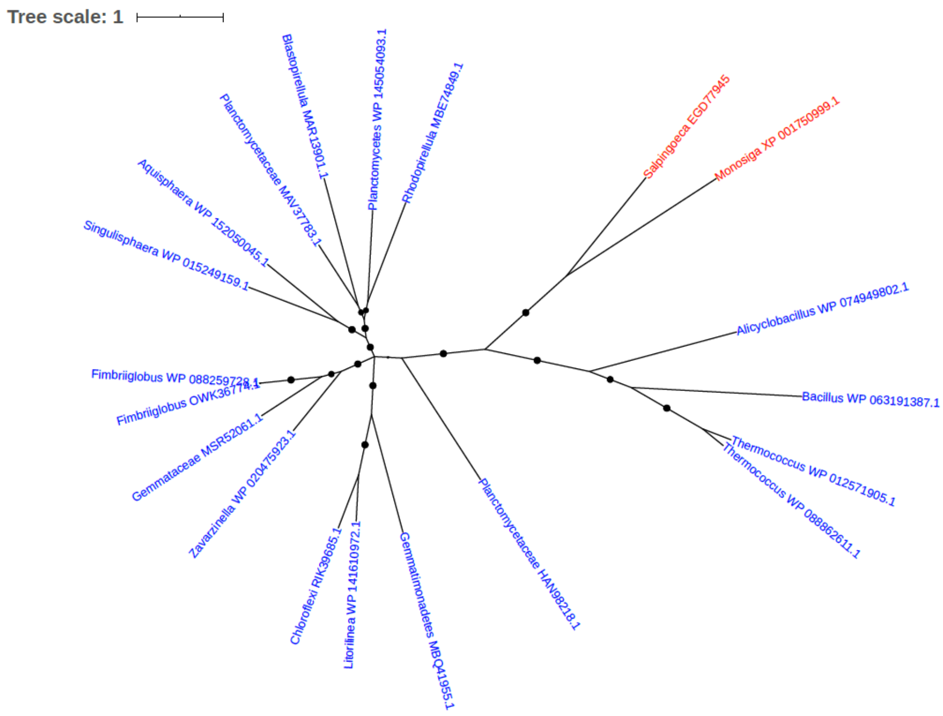


Figure S69. Phylogeny of D-isomer specific 2-hydroxyacid dehydrogenase (EGD77945)

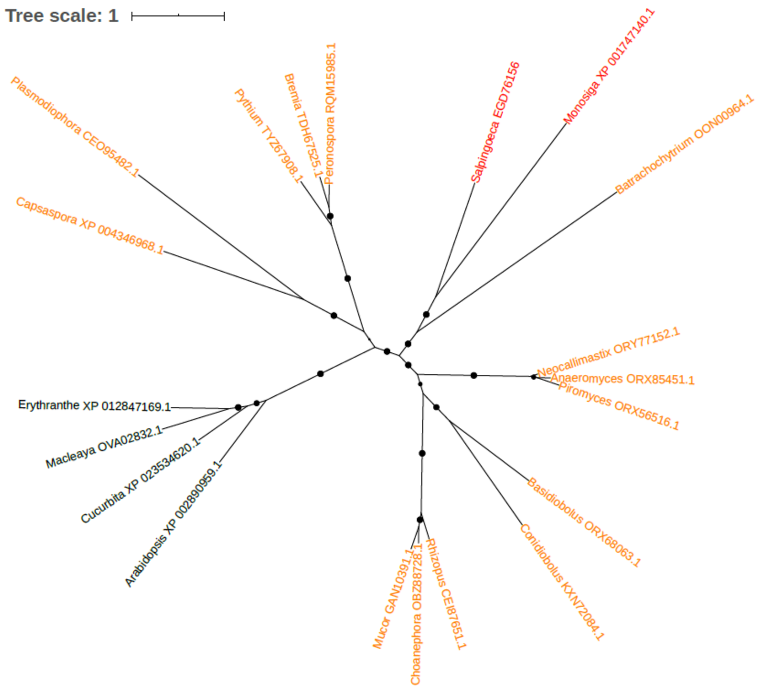


Figure S70. Phylogeny of DEAD box polypeptide 17 isoform 2 (EGD76156)

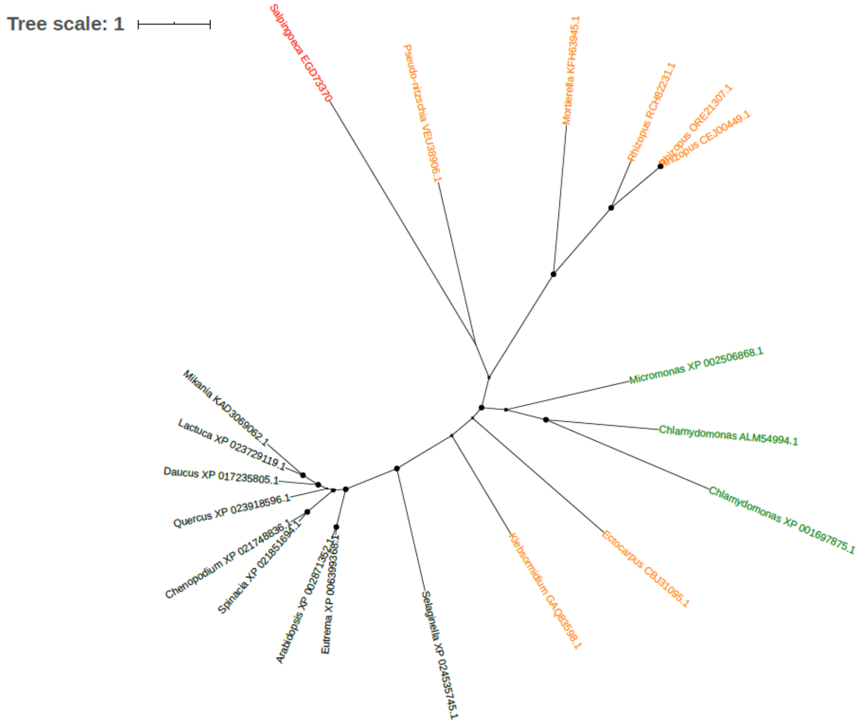


Figure S71. Phylogeny of DEAD-box ATP-dependent RNA helicase 26 (EGD73370)

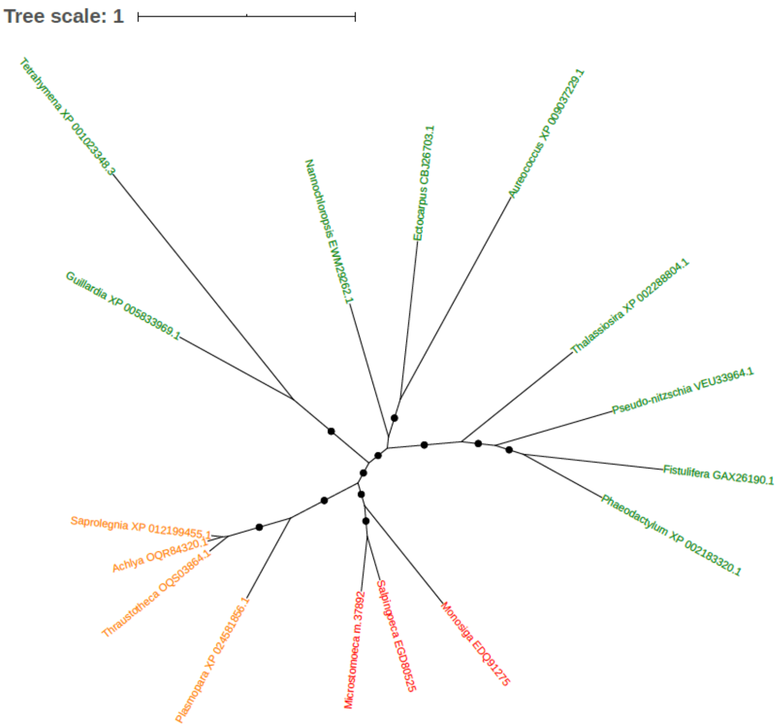


Figure S72. Phylogeny of Dehydrogenase (EGD80525)

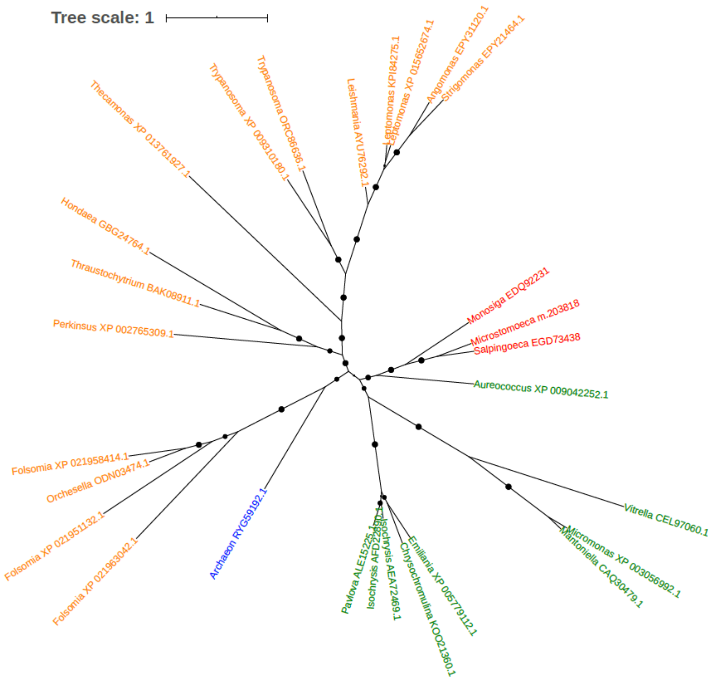


Figure S73. Phylogeny of Delta-5 desaturase (EGD73438)

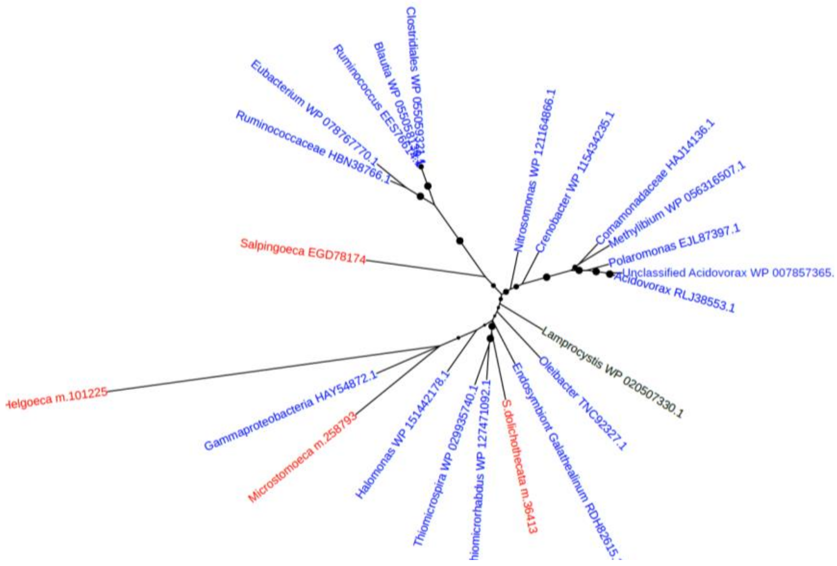


Figure S74. Phylogeny of Diaminopimelate epimerase (EGD78174)

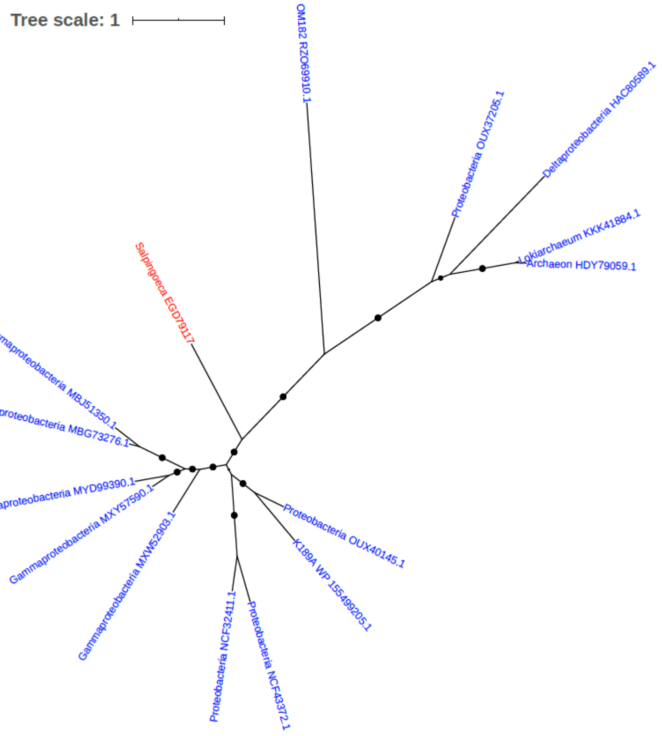


Figure S75. Phylogeny of Diaminopimelate epimerase (EGD79117)

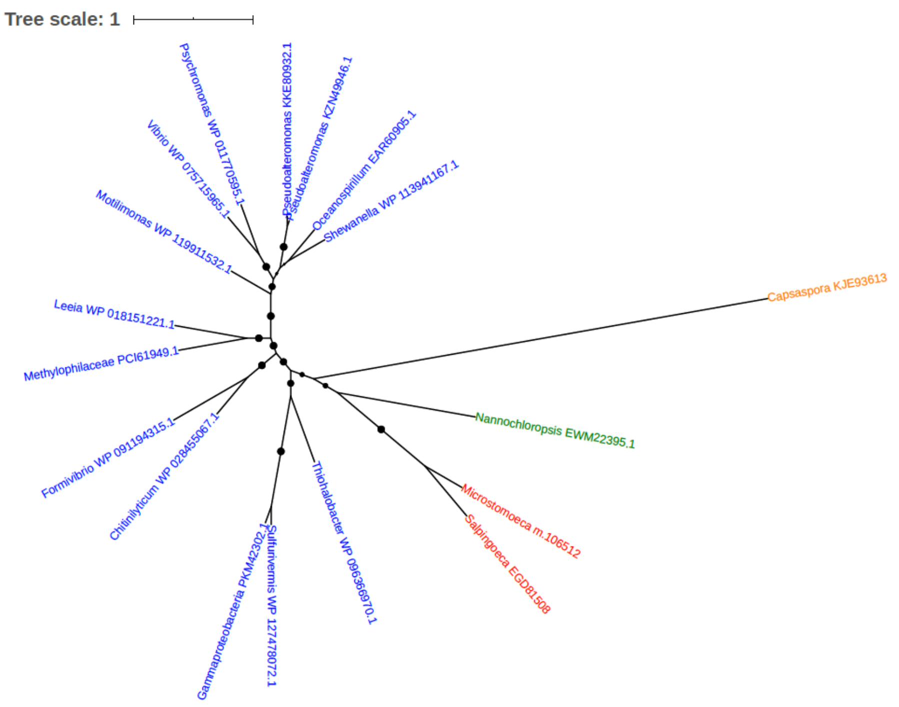


Figure S76. Phylogeny of Dihydroorotase (EGD81508)

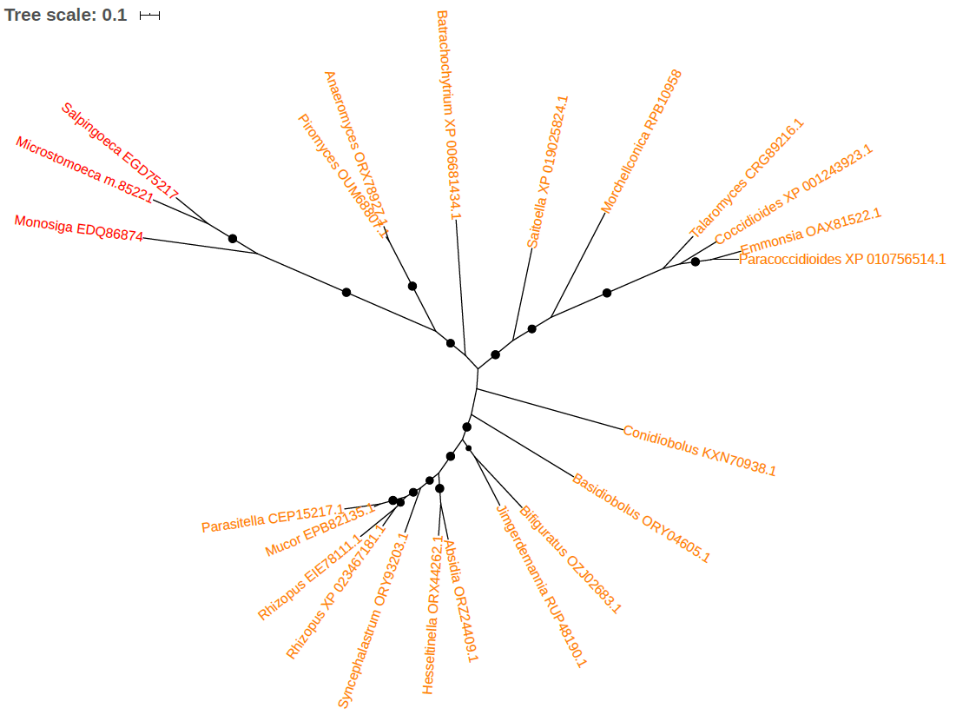


Figure S77. Phylogeny of Diphthine synthase (EGD75217)

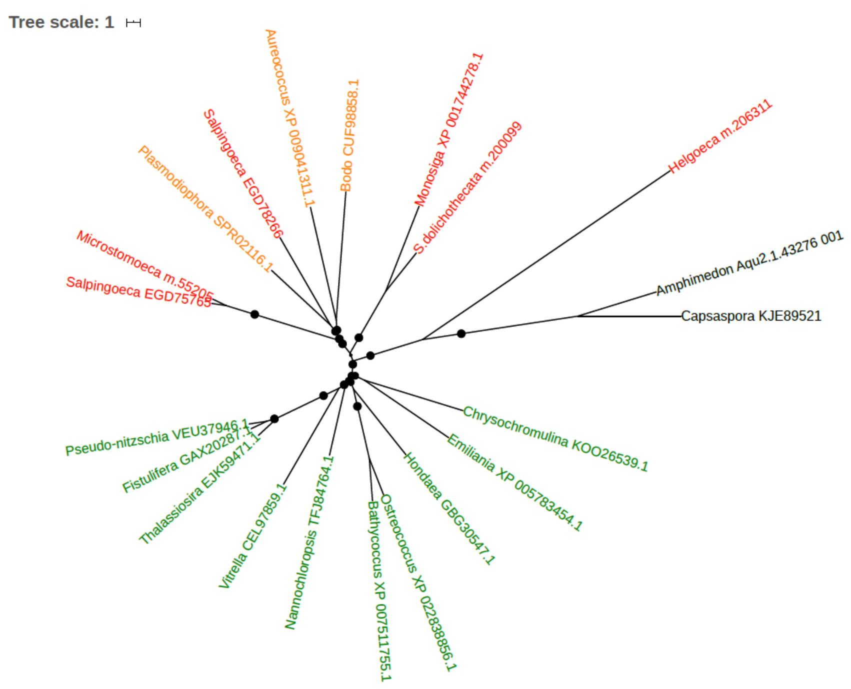


Figure S78. Phylogeny of Disulfide-isomerase (EGD75765)

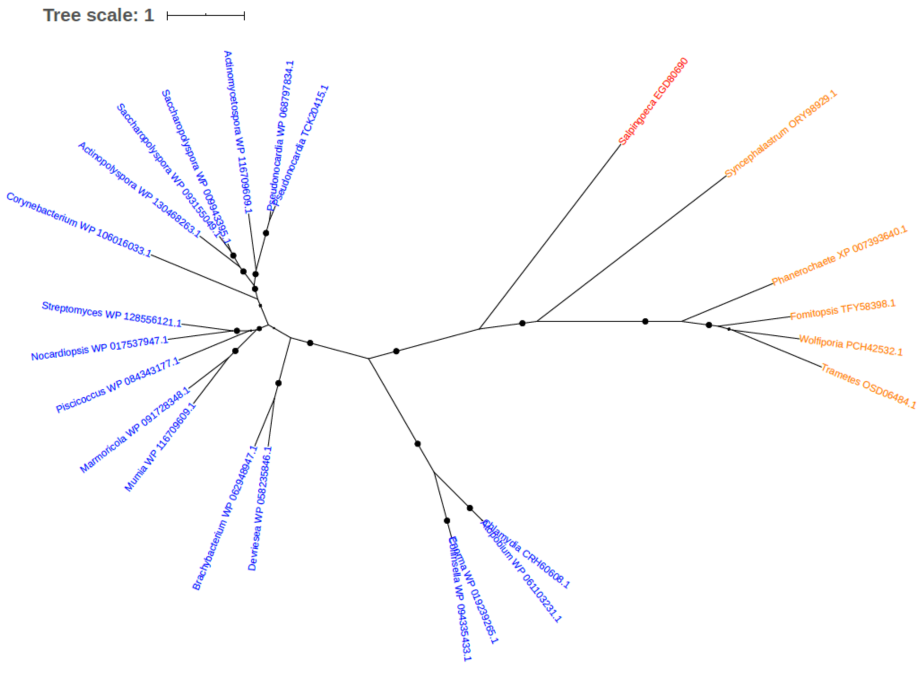


Figure S79. Phylogeny of DNA and RNA helicase (EGD80690)

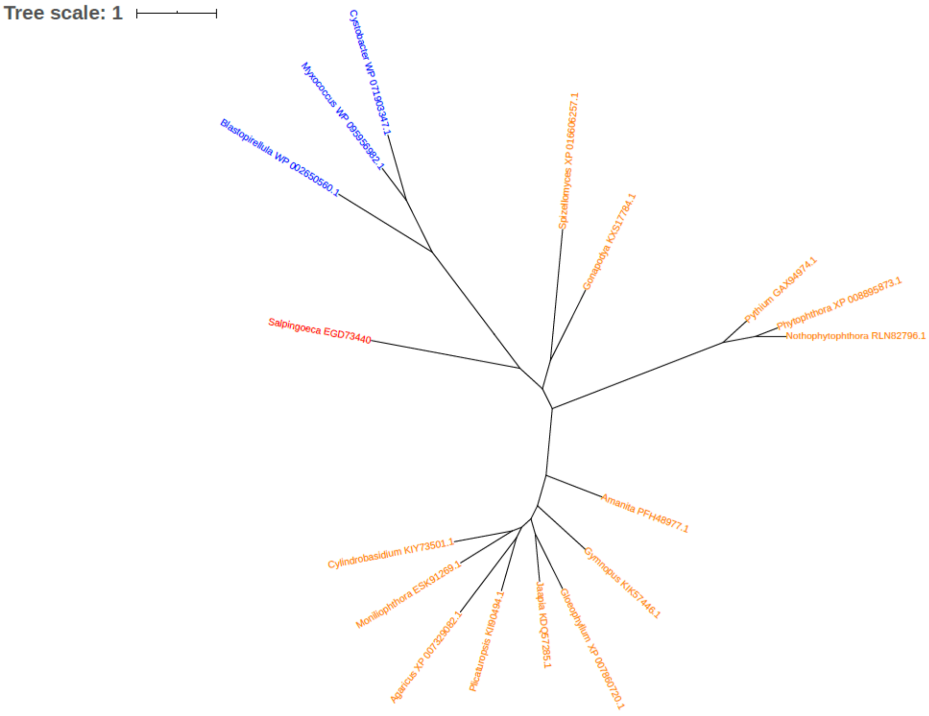


Figure S80. Phylogeny of DNA ligase (EGD73440)

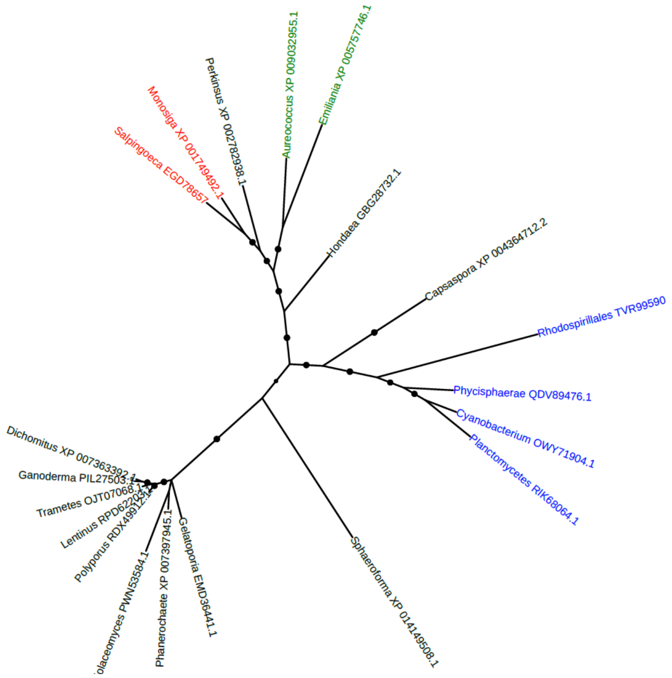


Figure S81. Phylogeny of Dolichol-phosphate mannosyltransferase subunit 1 (EC 2.4.1.83) (EGD78657)

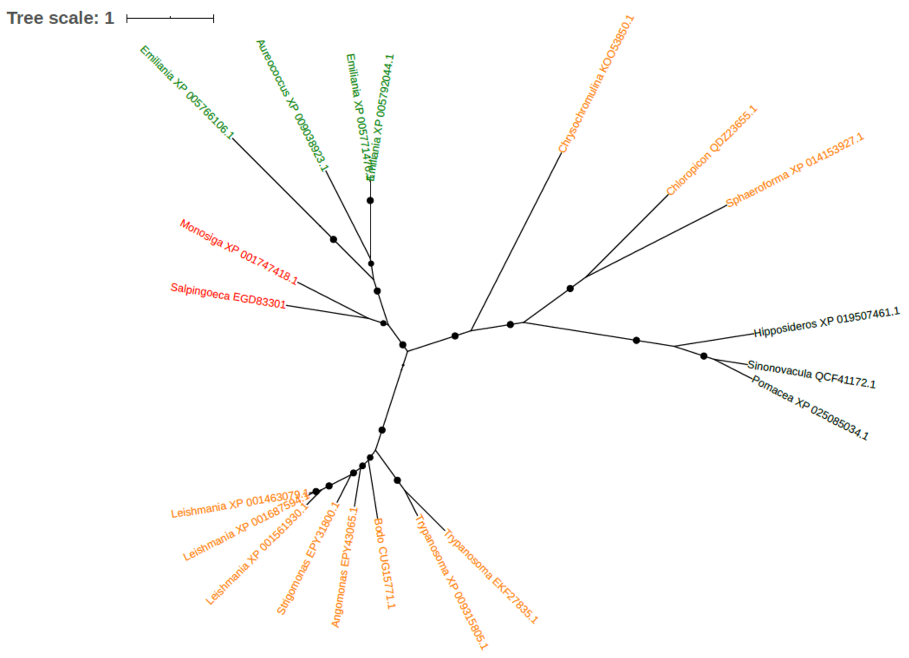


Figure S82. Phylogeny of Elongation of fatty acids protein (EC 2.3.1.199) (Very-long-chain 3-oxoacyl-CoA synthase) (EGD83301)

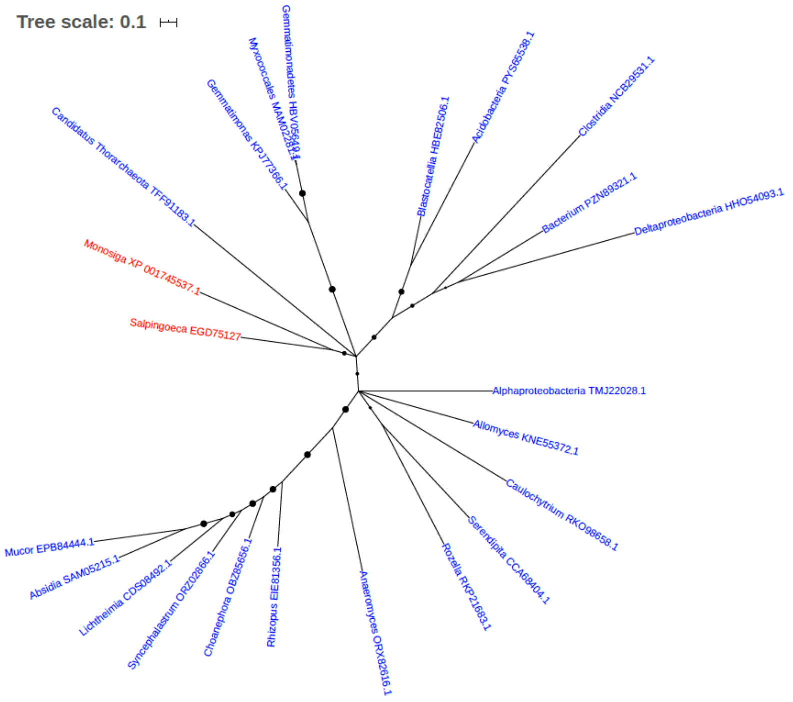


Figure S83. Phylogeny of Endoribonuclease L-PSP (EGD75127)

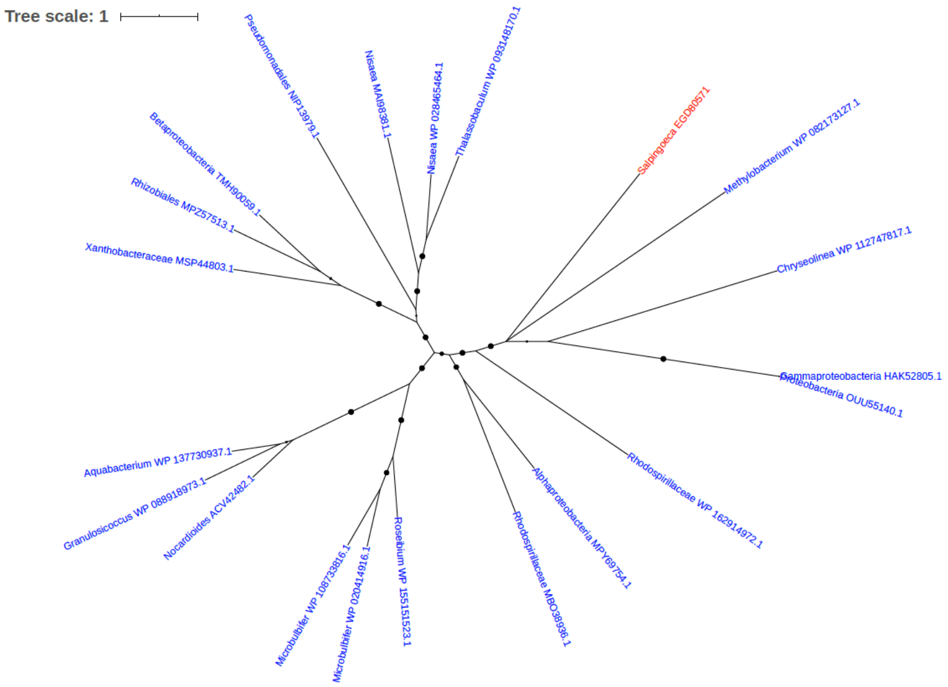


Figure S84. Phylogeny of Esterase (EGD80571)

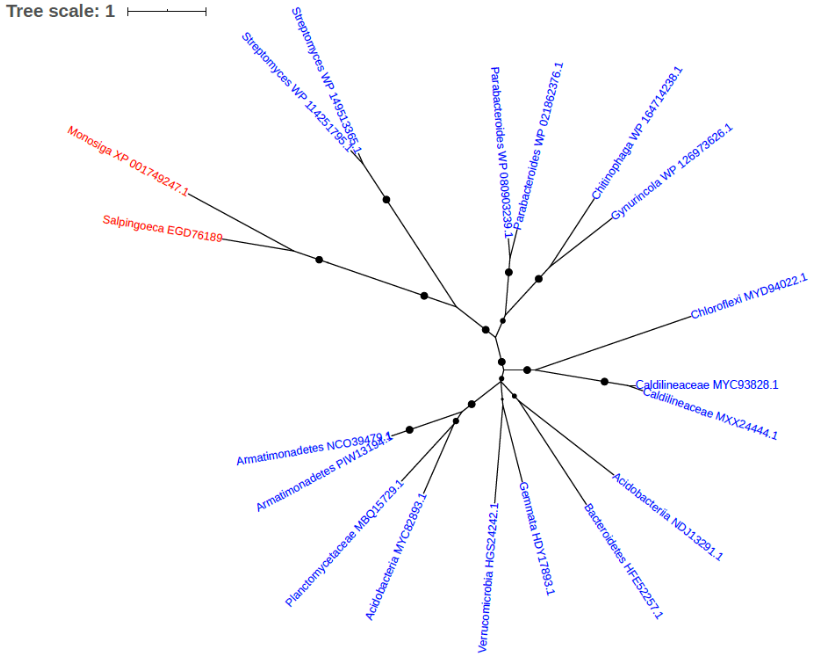


Figure S85. Phylogeny of Exo-alpha-sialidase (EGD76189)

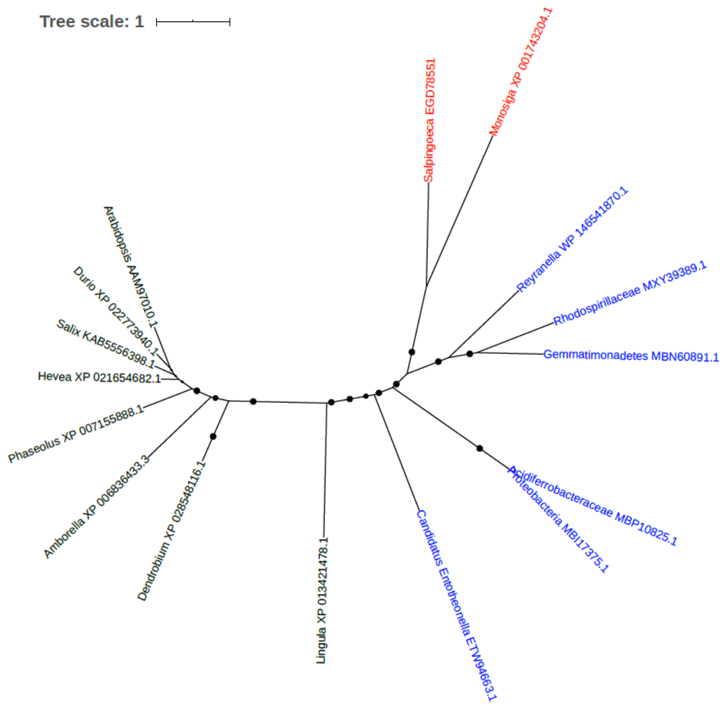


Figure S86. Phylogeny of FAD-dependent oxidoreductase (EGD78551)

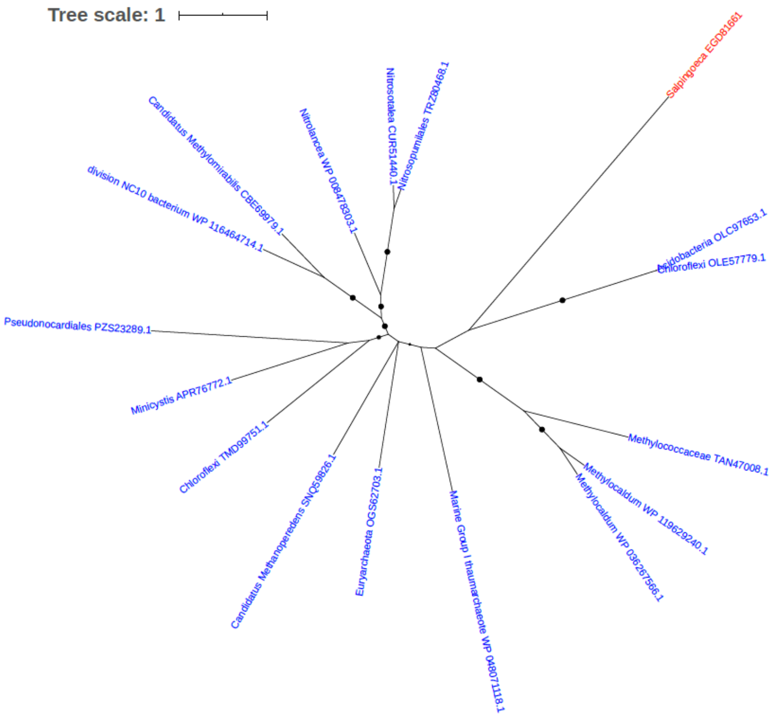


Figure S87. Phylogeny of Ferroxidase (EGD81661)

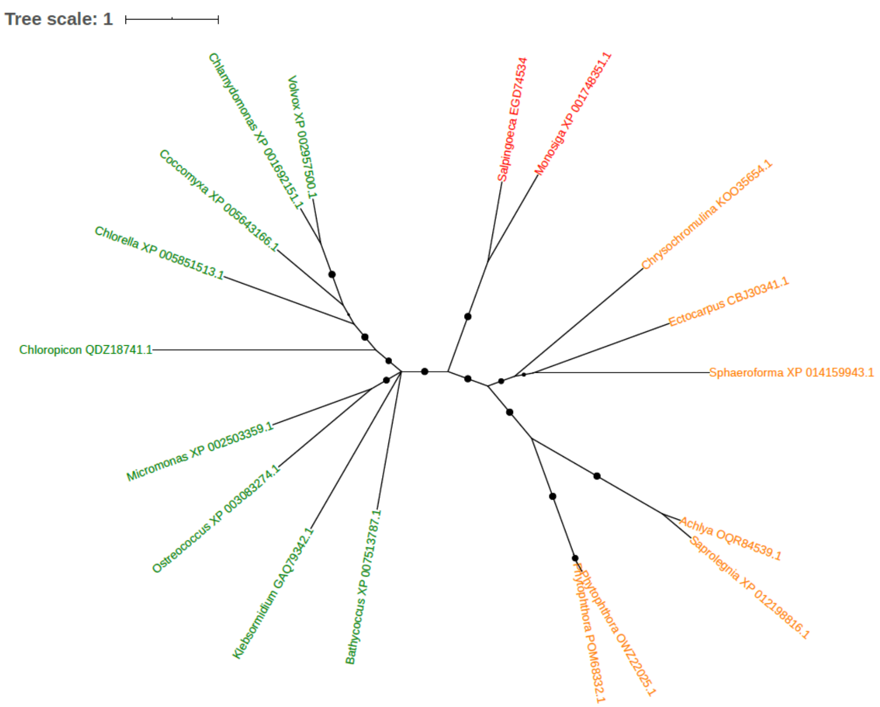


Figure S88. Phylogeny of Flagellar associated protein (EGD74534)

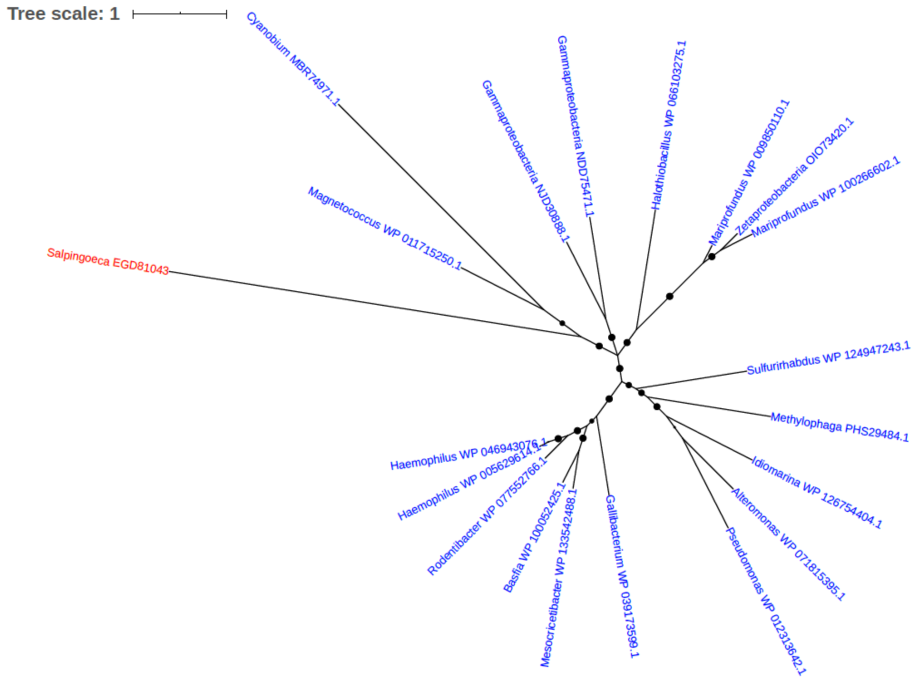


Figure S89. Phylogeny of FolD protein (EGD81043)

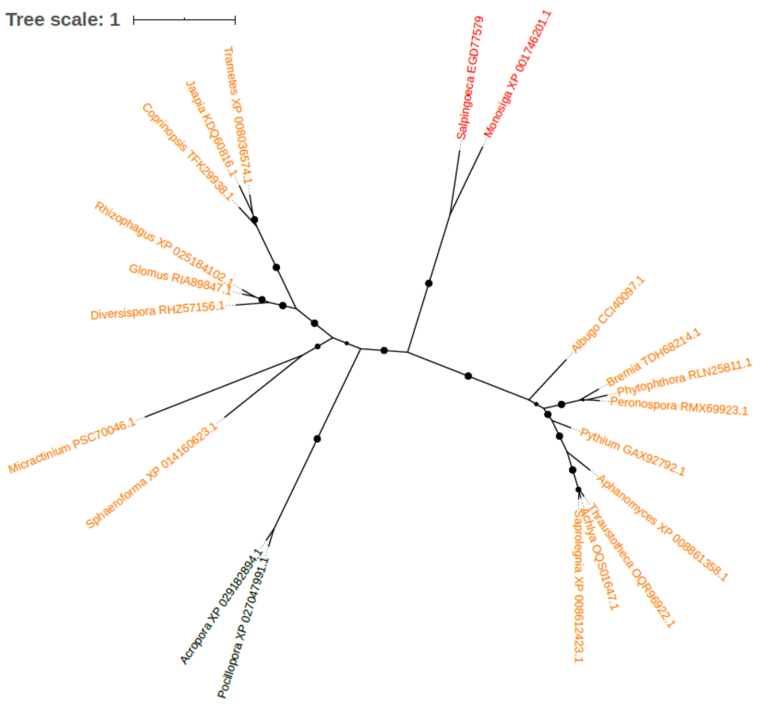


Figure S90. Phylogeny of Fructose-1,6-bisphosphatase (EGD77579)

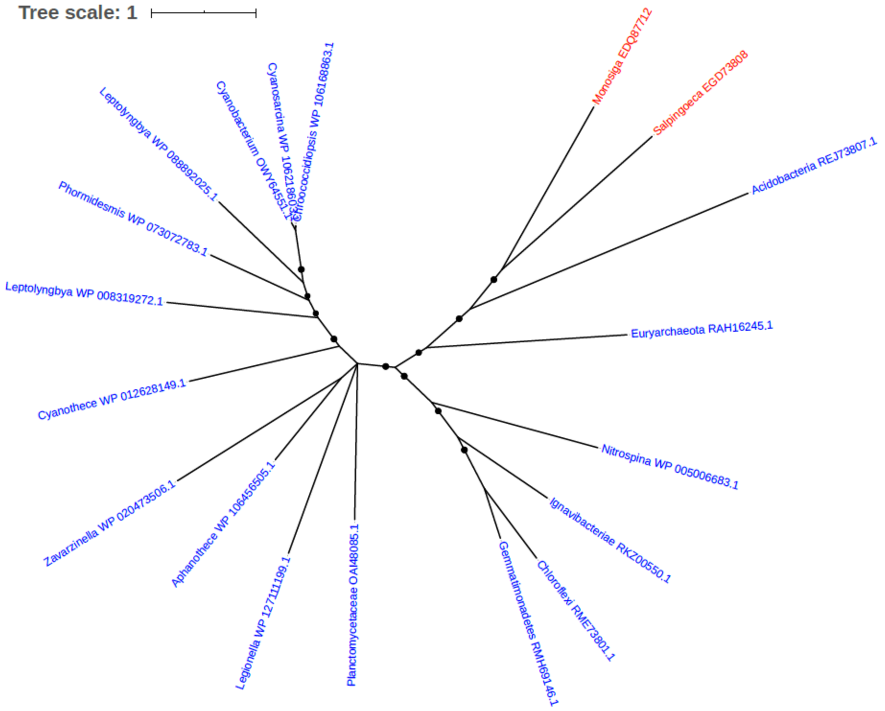


Figure S91. Phylogeny of Fumarate hydratase (EGD73808)

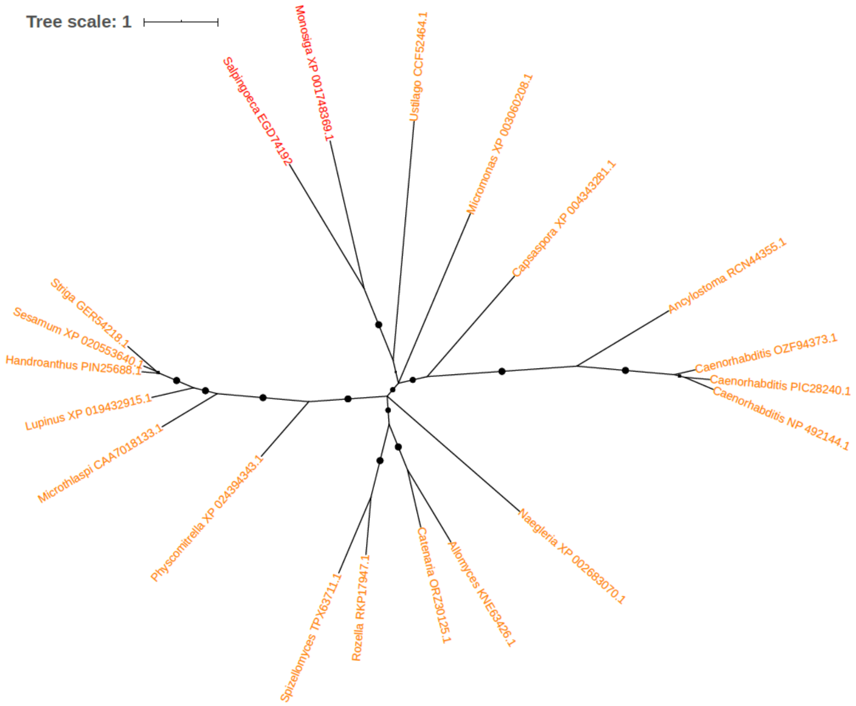


Figure S92. Phylogeny of Glucosamine 6-phosphate N-acetyltransferase (EC 2.3.1.4) (EGD74192)

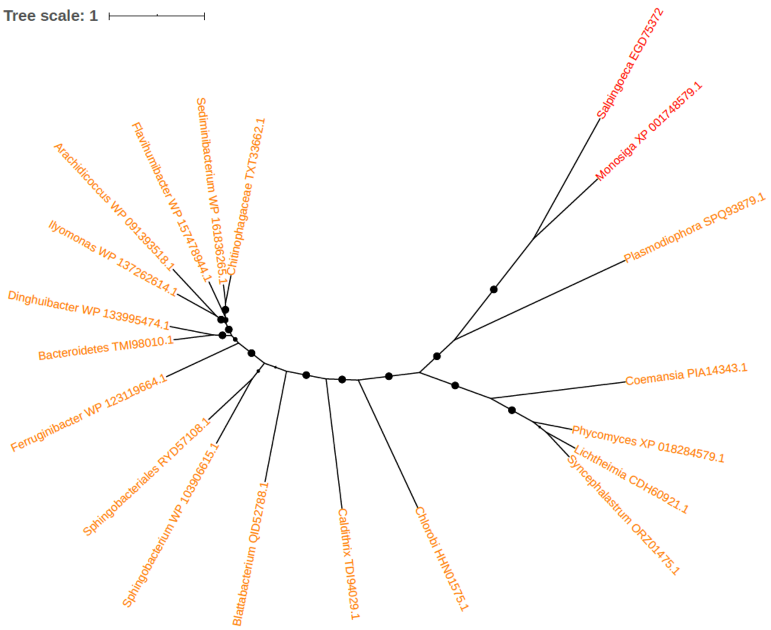


Figure S93. Phylogeny of Glutamate dehydrogenase (EGD75372)

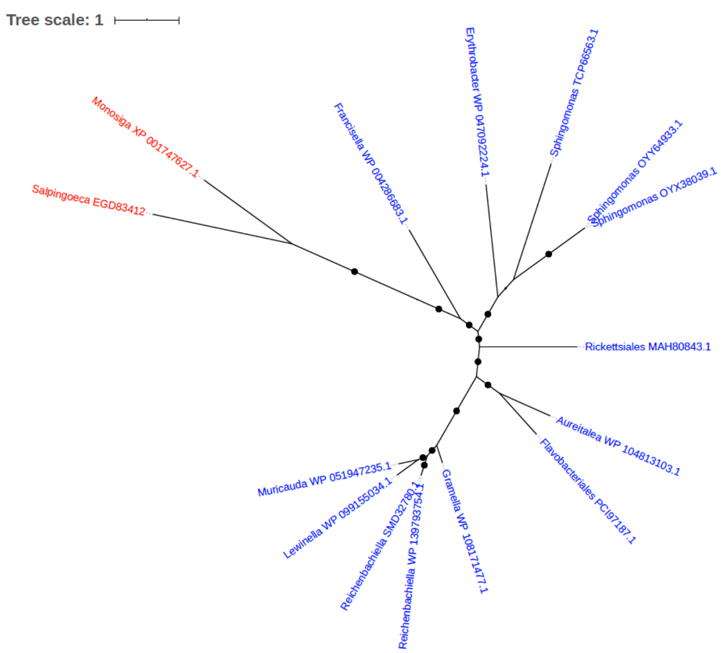


Figure S94. Phylogeny of Glutaminase (EGD83412)

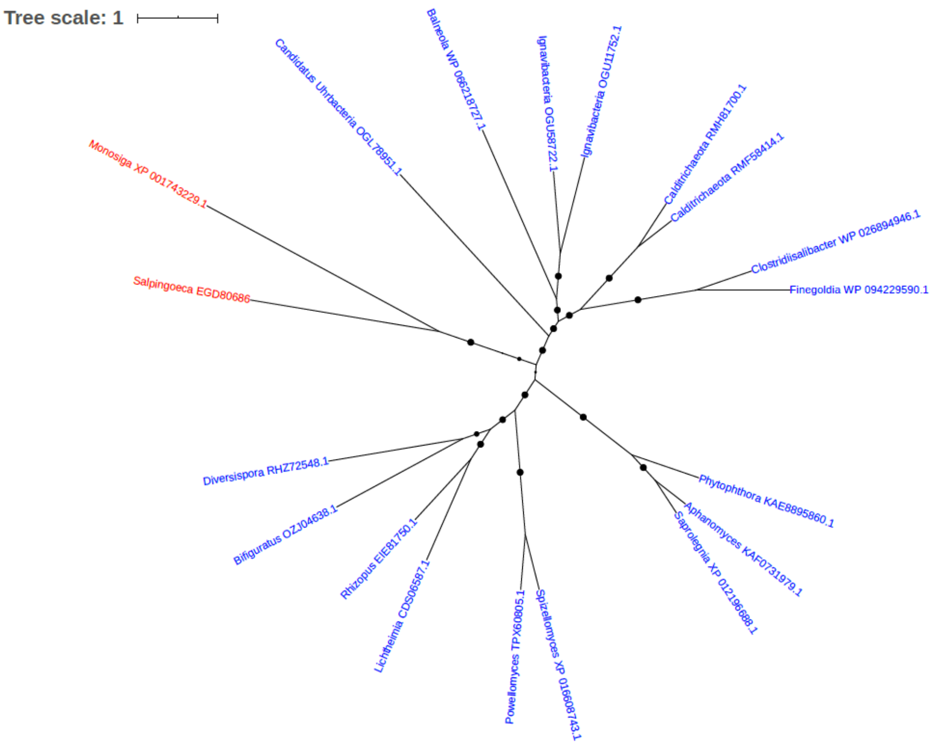


Figure S95. Phylogeny of Glutamyl-tRNA synthetase (EGD80686)

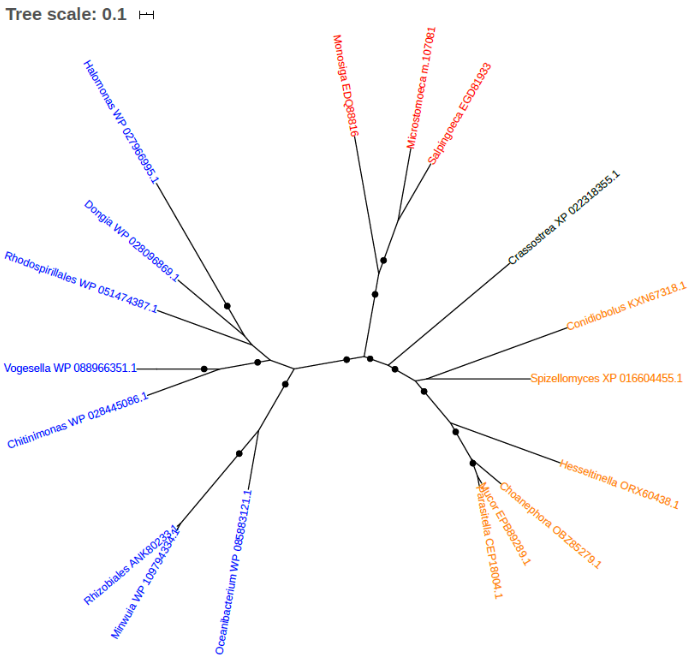


Figure S96. Phylogeny of Glutaryl-CoA dehydrogenase (EGD81933)

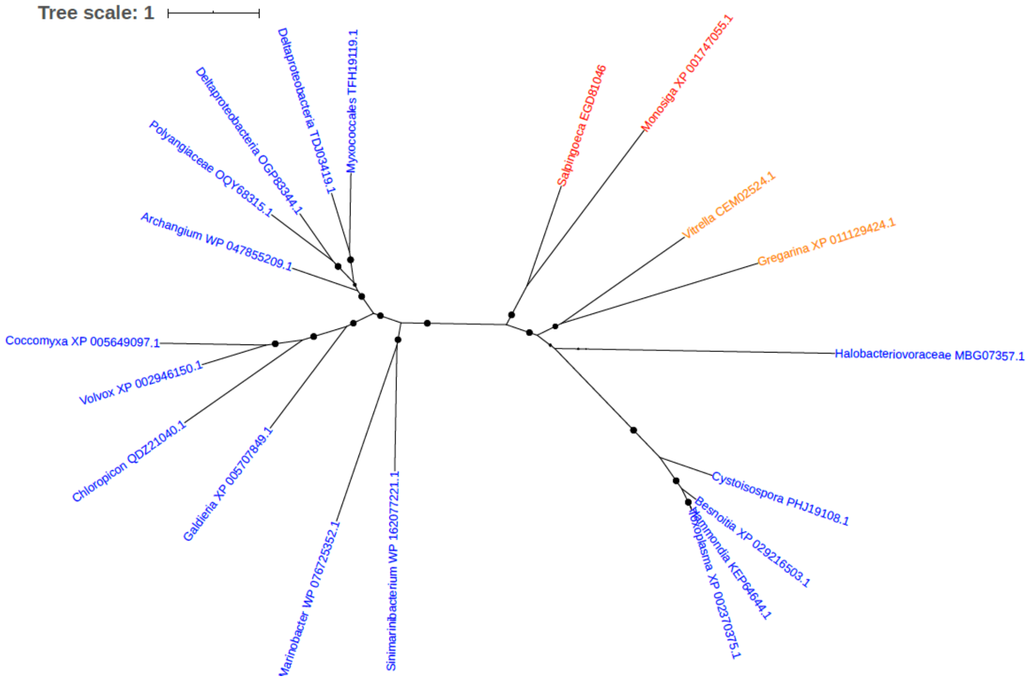


Figure S97. Phylogeny of Glutathione S-transferase domain-containing protein (EGD81046)

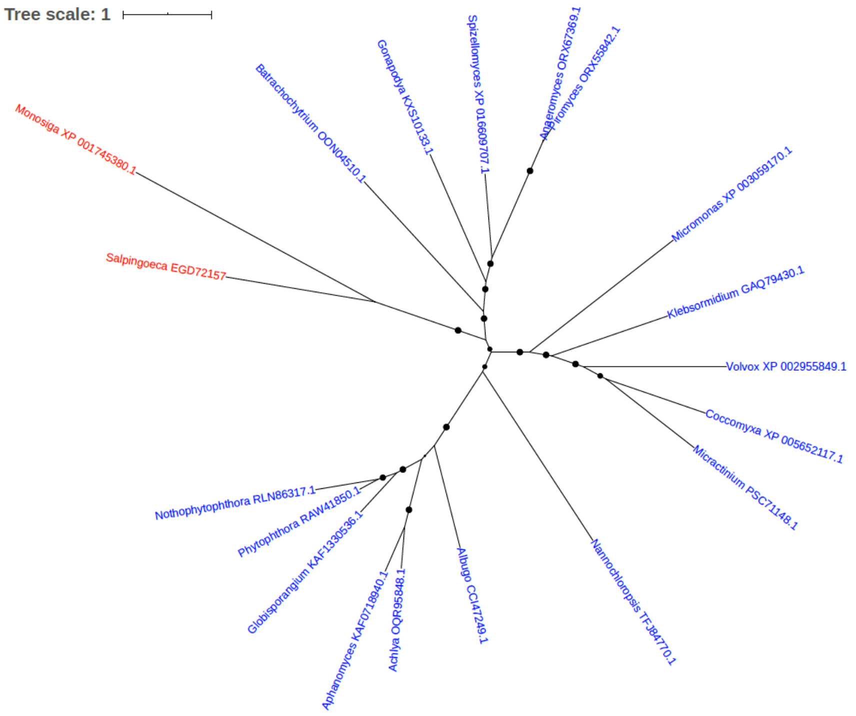


Figure S98. Phylogeny of Glutathione synthetase (GSH-S) (EC 6.3.2.3) (EGD72157)

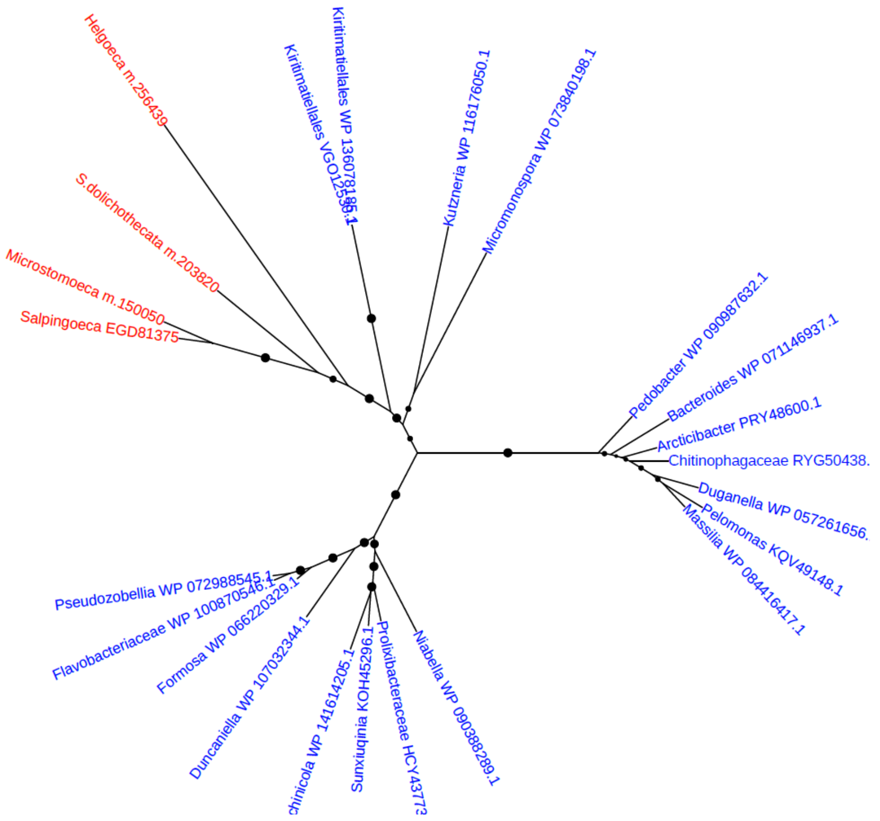


Figure S99. Phylogeny of Glycosyl hydrolase, family 88 (EGD81375)

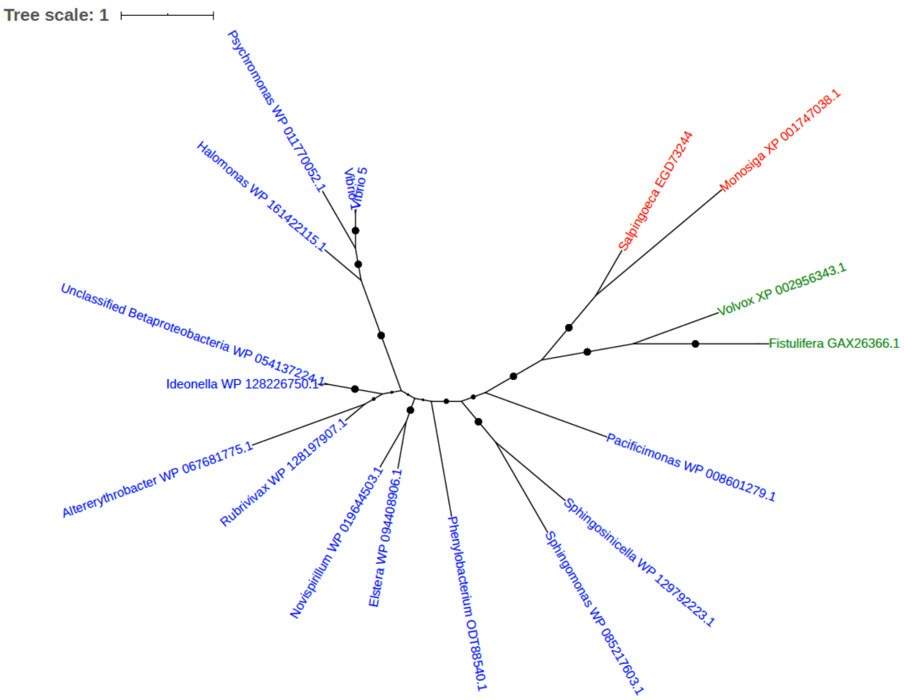


Figure S100. Phylogeny of Glyoxalase/bleomycin resistance protein/dioxygenase (EGD73244)

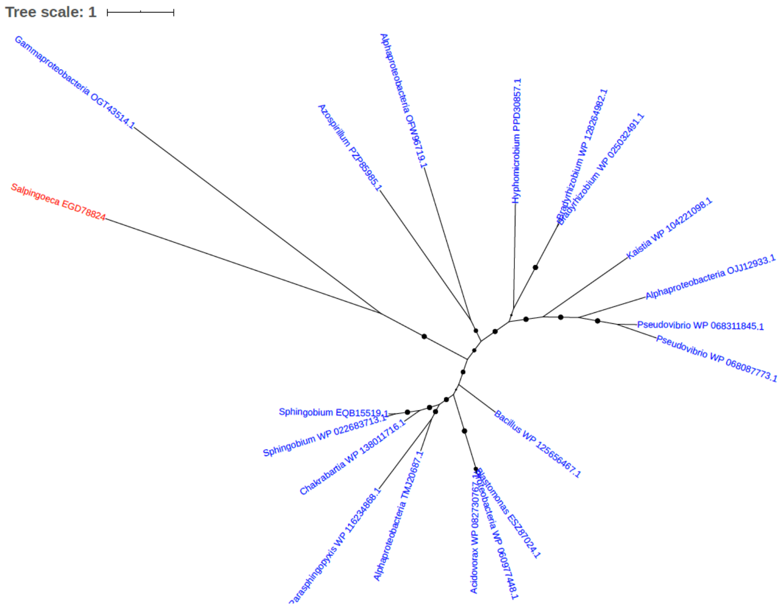


Figure S101. Phylogeny of GTPase EngB (EGD78824)

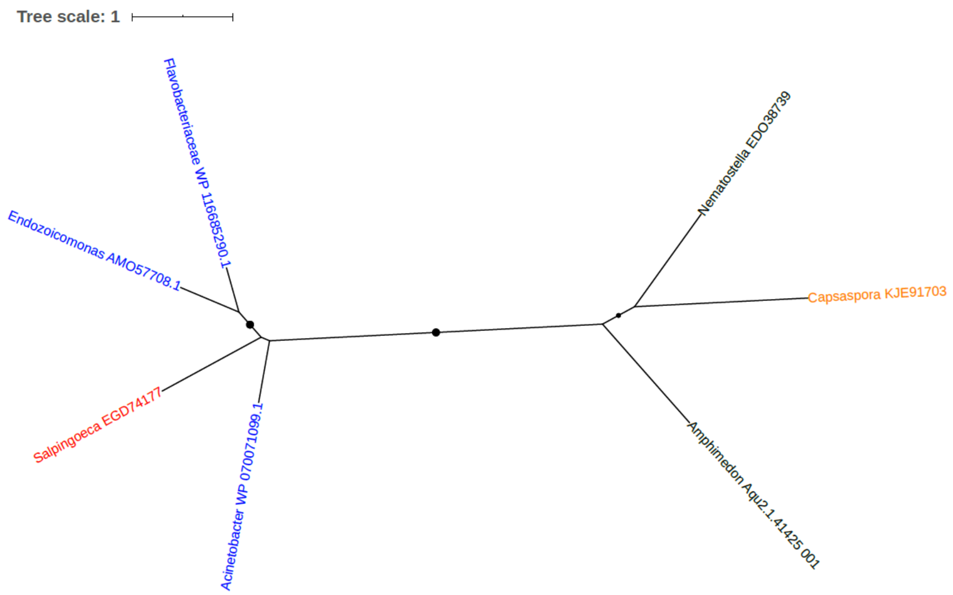


Figure S102. Phylogeny of Guanine deaminase (Guanase) (EC 3.5.4.3) (Guanine aminohydrolase) (EGD4177)

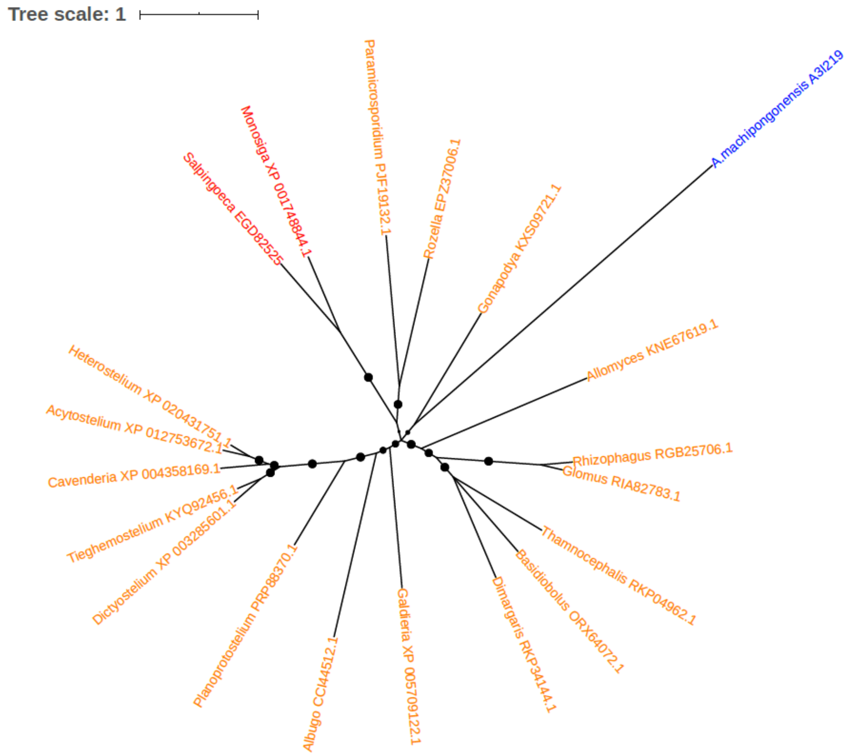


Figure S103. Phylogeny of Heat shock protein (EGD82515)

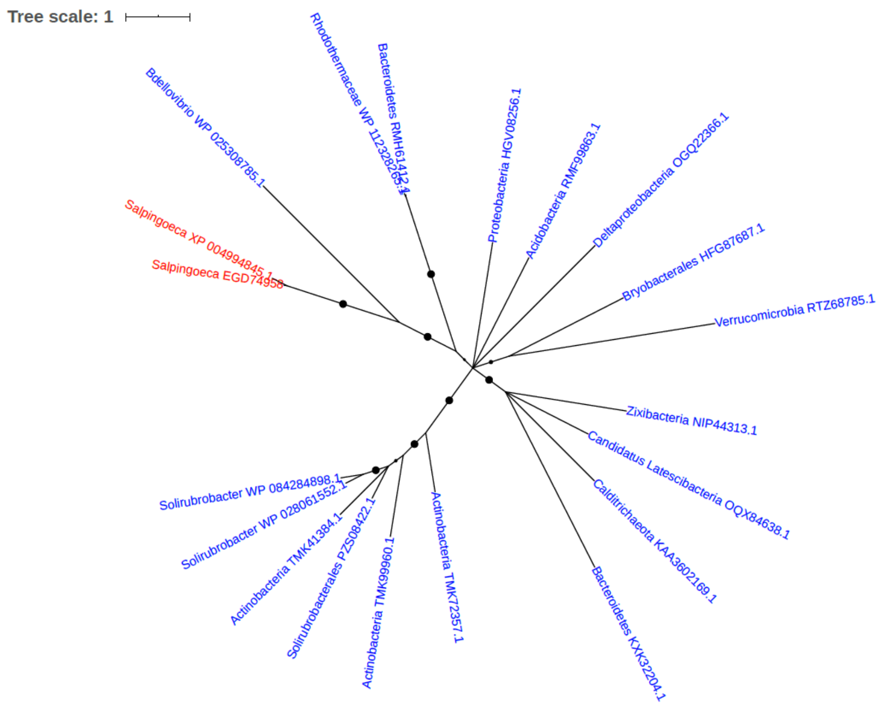


Figure S104. Phylogeny of Heat shock protein (EGD74958)

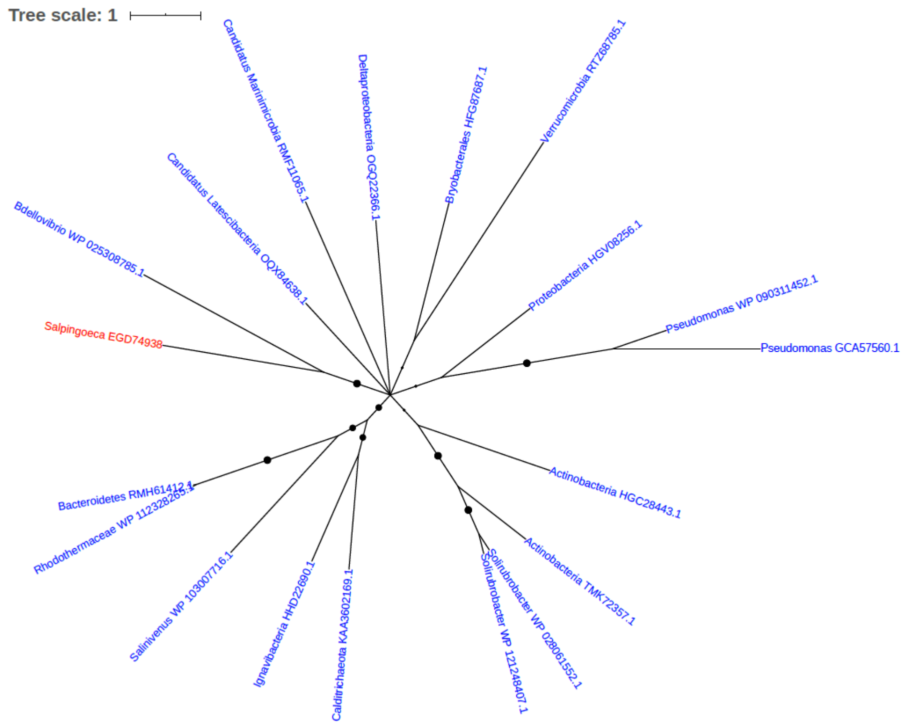


Figure S105. Phylogeny of Heat shock protein (EGD74938)

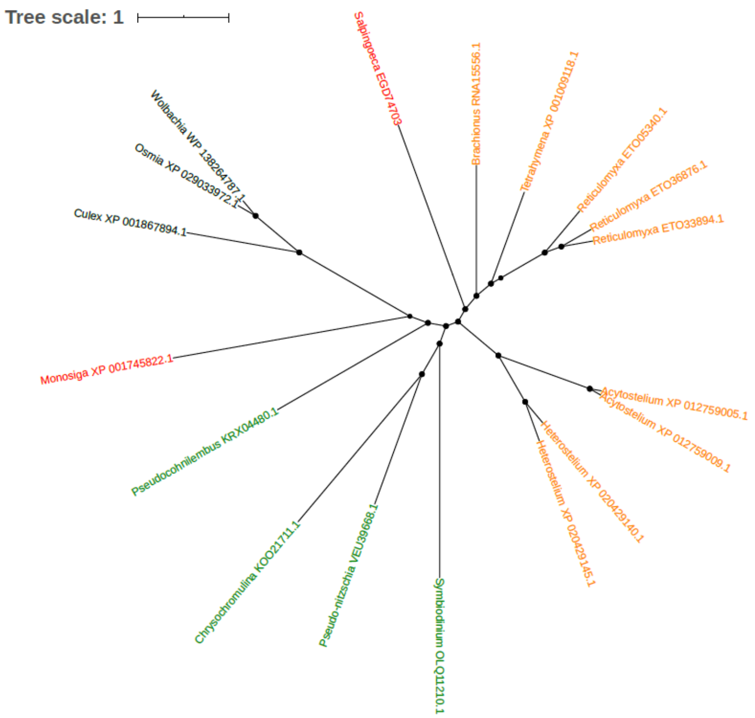


Figure S106. Phylogeny of Helicase (EGD74703)

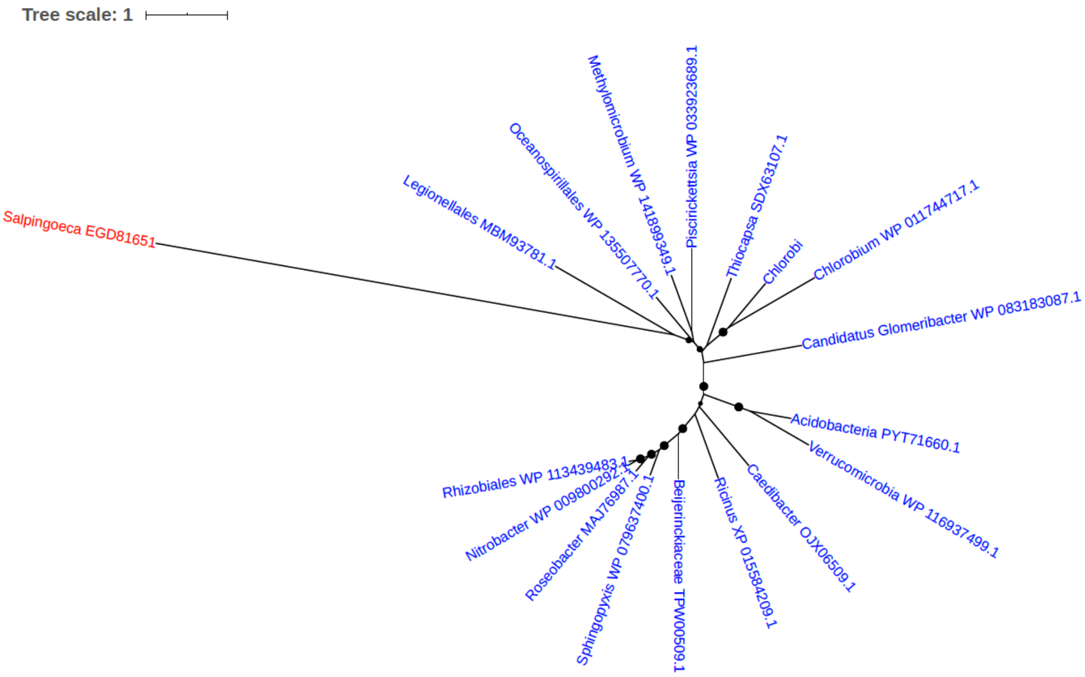


Figure S107. Phylogeny of Helicase RecD/TraA (EGD81651)

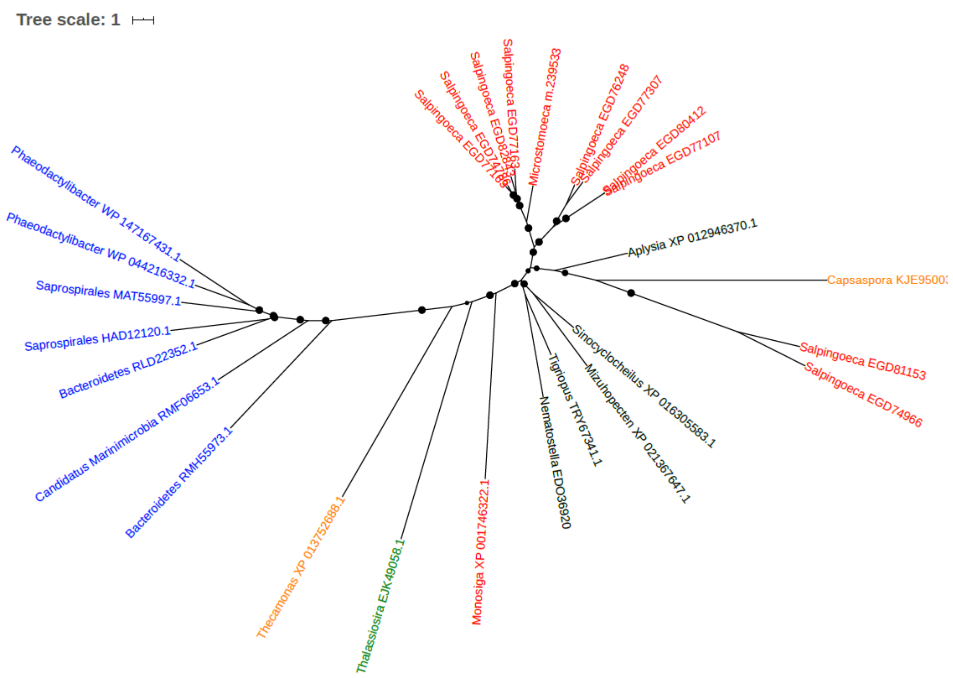


Figure S108. Phylogeny of Uncharacterized protein (EGD77163, EGD74786, EGD82843, EGD76248, EGD76248, EGD77307, EGD80412, EGD77107)

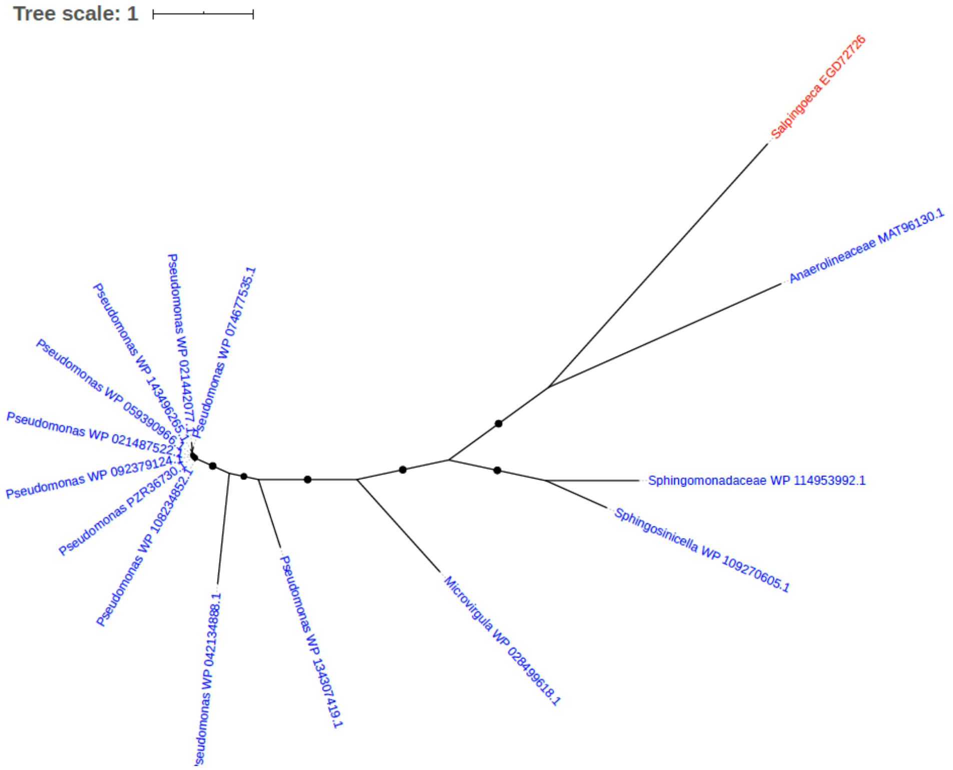


Figure S109. Phylogeny of Histone deacetylase superfamily protein (EGD72726)

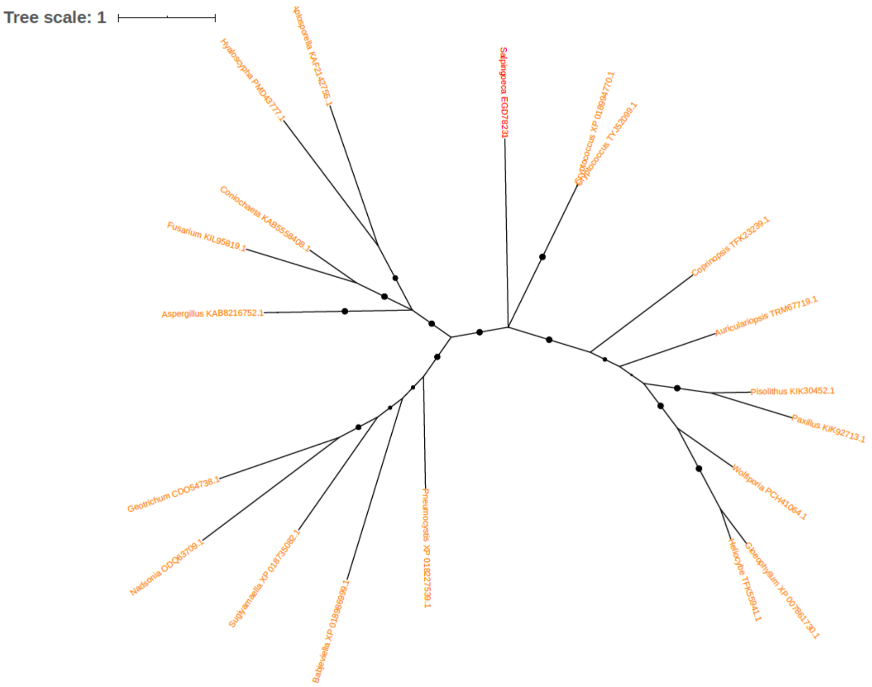


Figure S110. Phylogeny of Histone H3 (EGD78231)

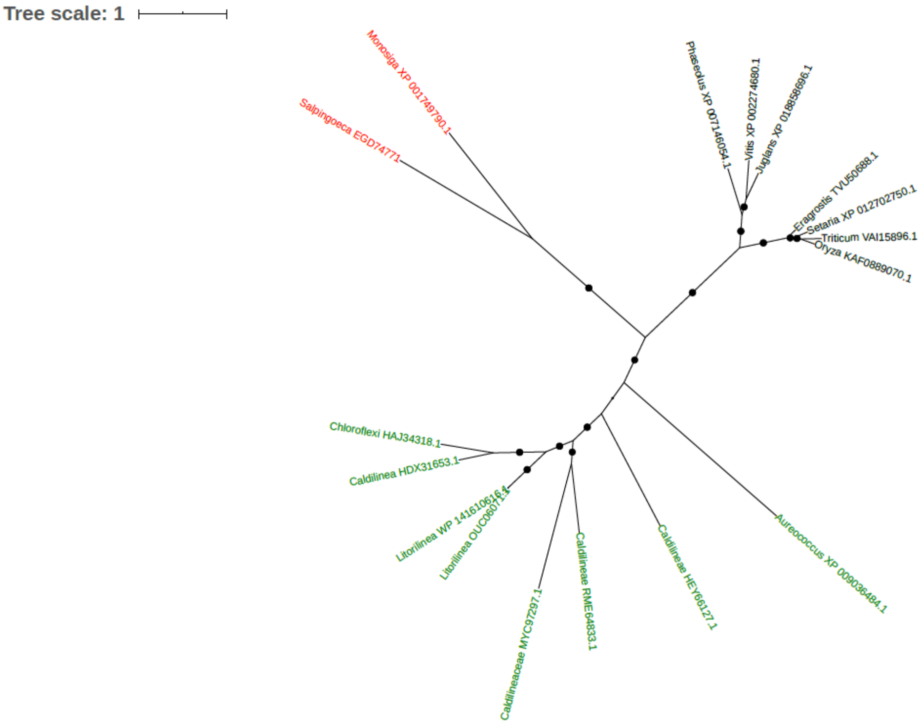


Figure S111. Phylogeny of Homoserine dehydrogenase (EGD74771)

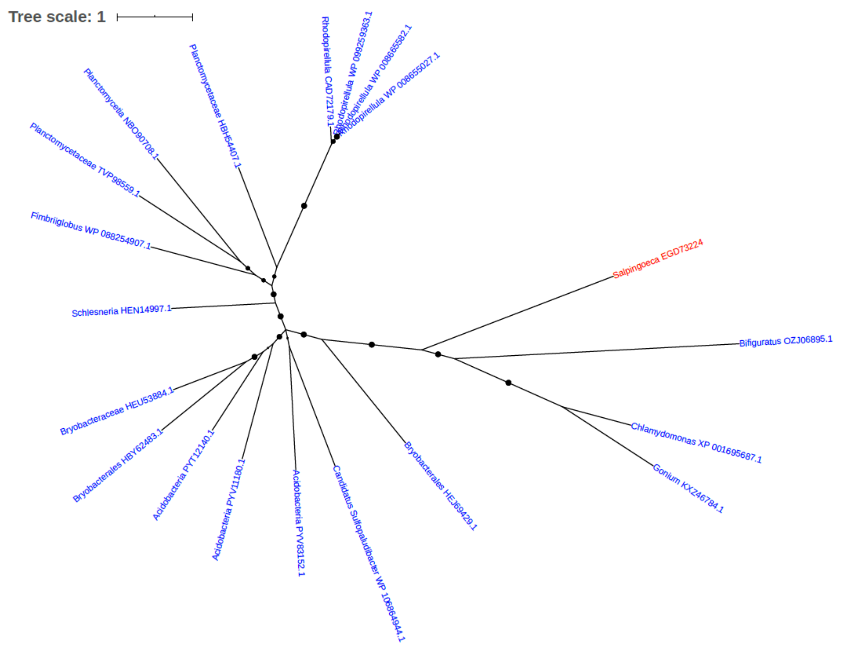


Figure S112. Phylogeny of Hydroxyneurosporene-O-methyltransferase (EGD73224)

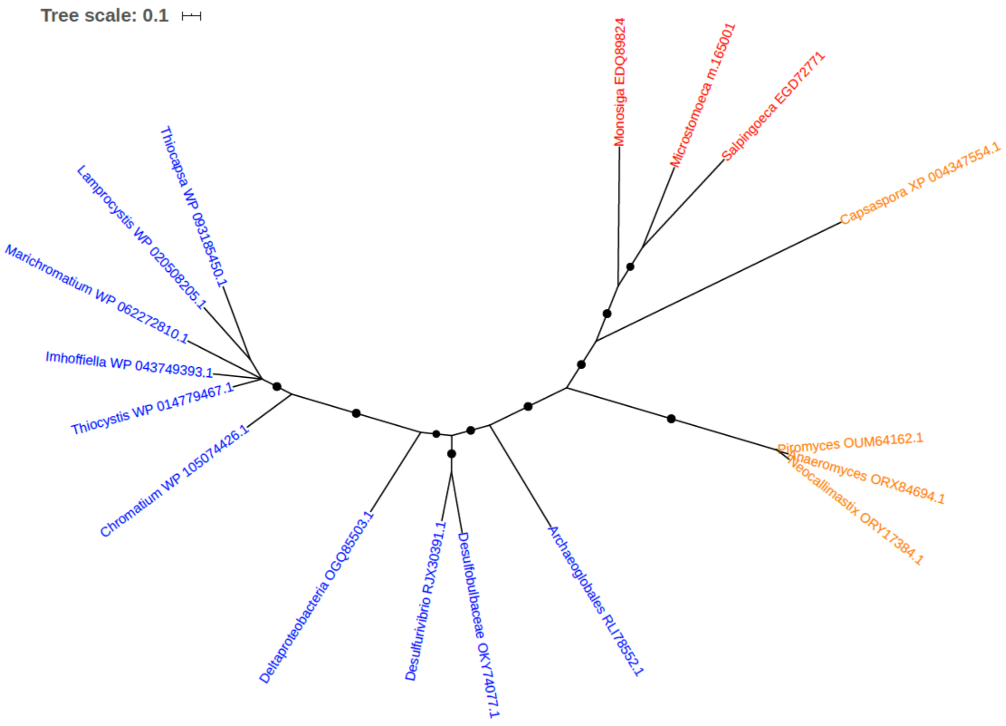


Figure S113. Phylogeny of Imidazole glycerol phosphate synthase hisHF [Includes: Glutaminase (EC 3.5.1.2); Cyclase] (EGD72771)

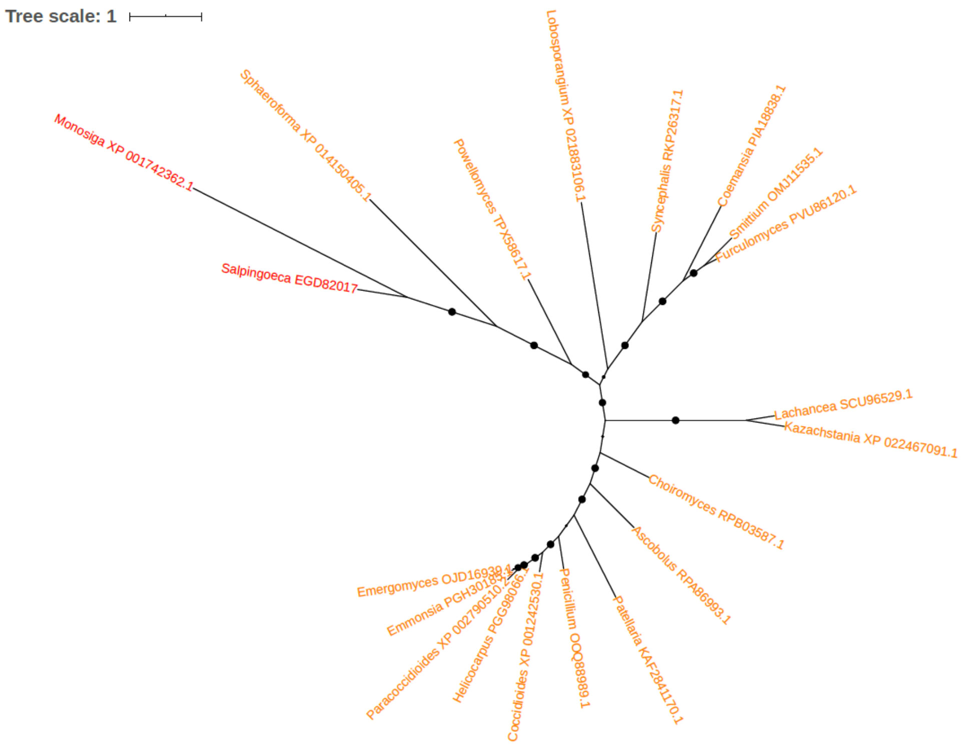


Figure S114. Phylogeny of Import inner membrane translocase subunit TIM22 (EGD82017)

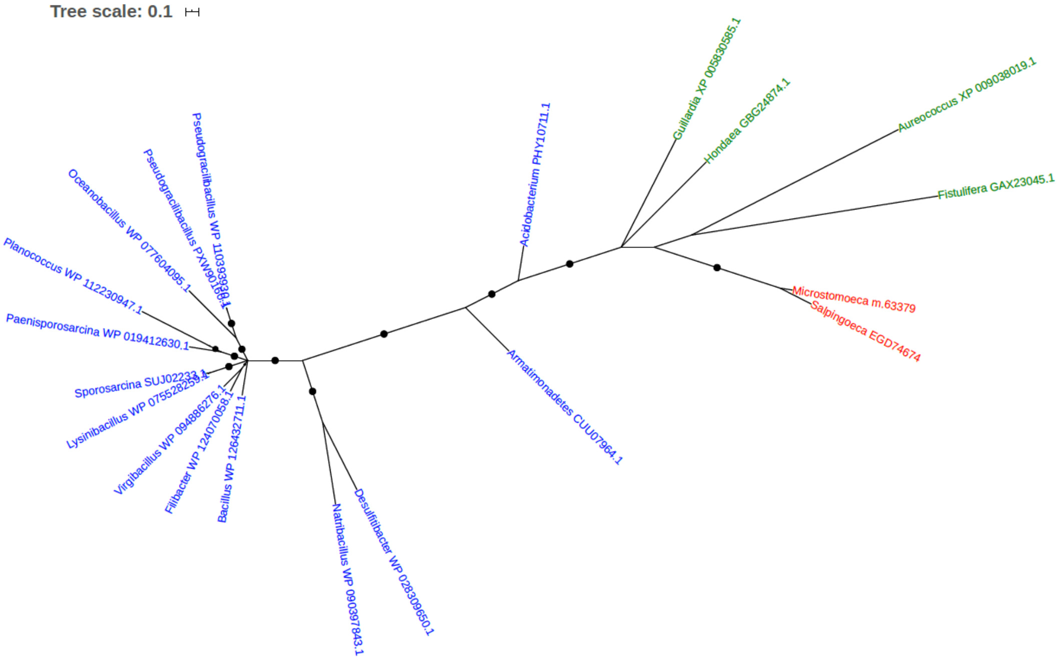


Figure S115. Phylogeny of Inner membrane protein (EGD74674)

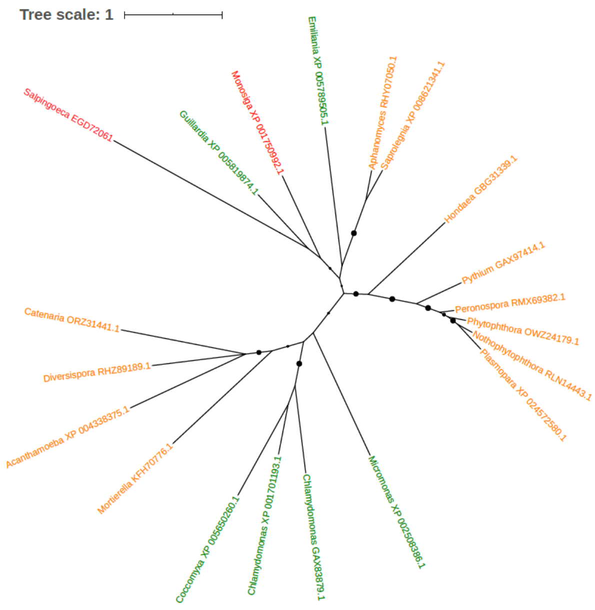


Figure S116. Phylogeny of Inosine triphosphate pyrophosphatase (ITPase) (Inosine triphosphatase) (EC 3.6.1.9) (EGD72061)

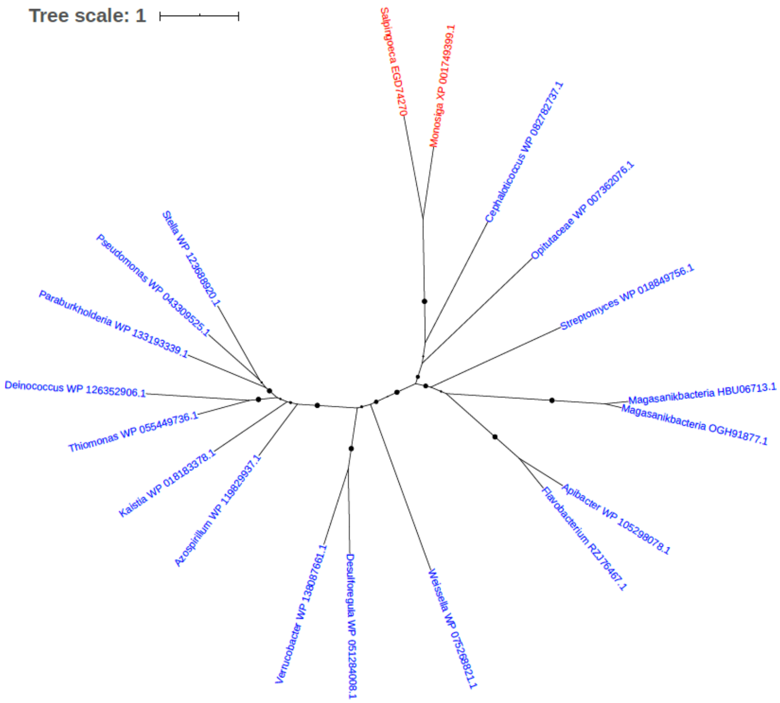


Figure S117. Phylogeny of Integral membrane protein TerC (EGD74270)

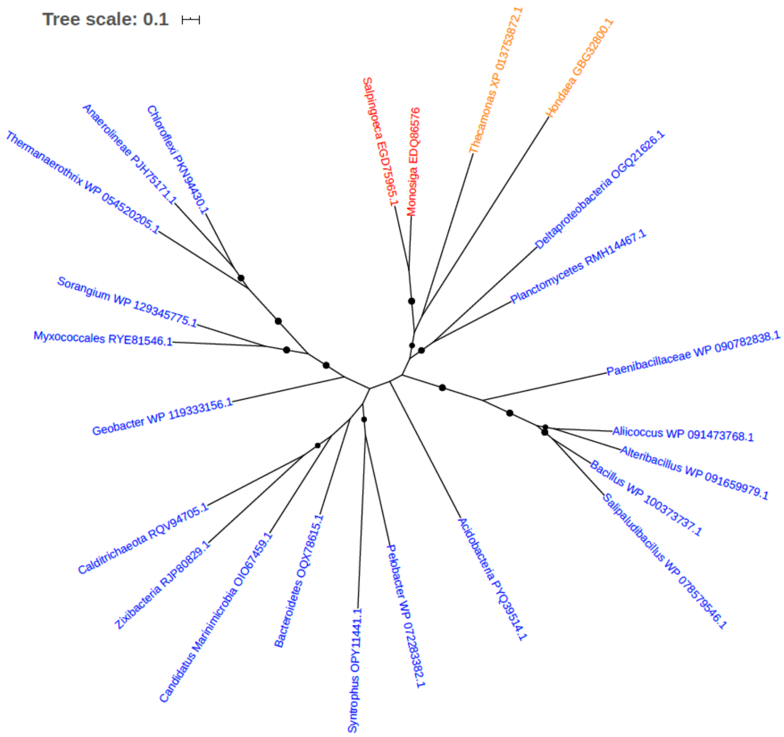


Figure S118. Phylogeny of Isocitrate dehydrogenase [NADP] (EC 1.1.1.42) (EGD75965)

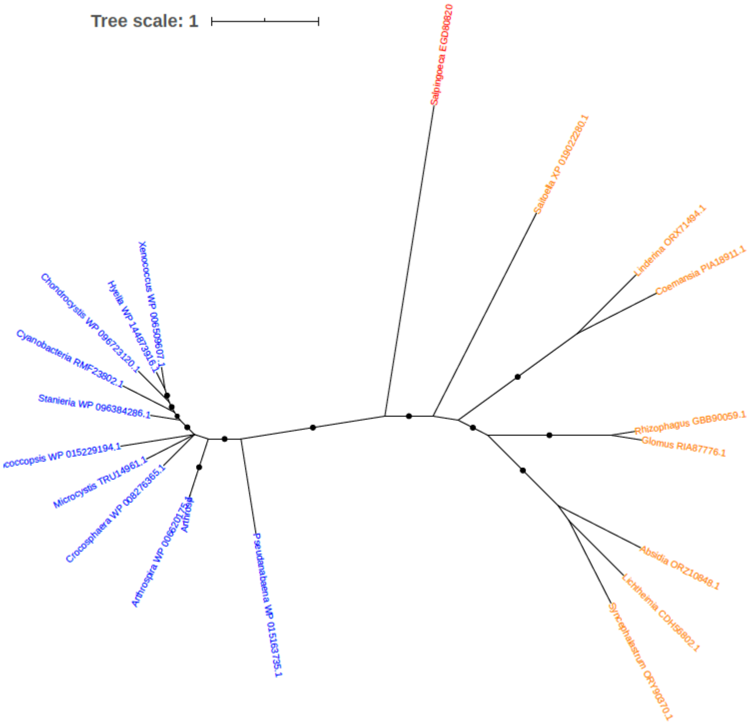


Figure S119. Phylogeny of Isoleucyl-tRNA synthetase (EGD80820)

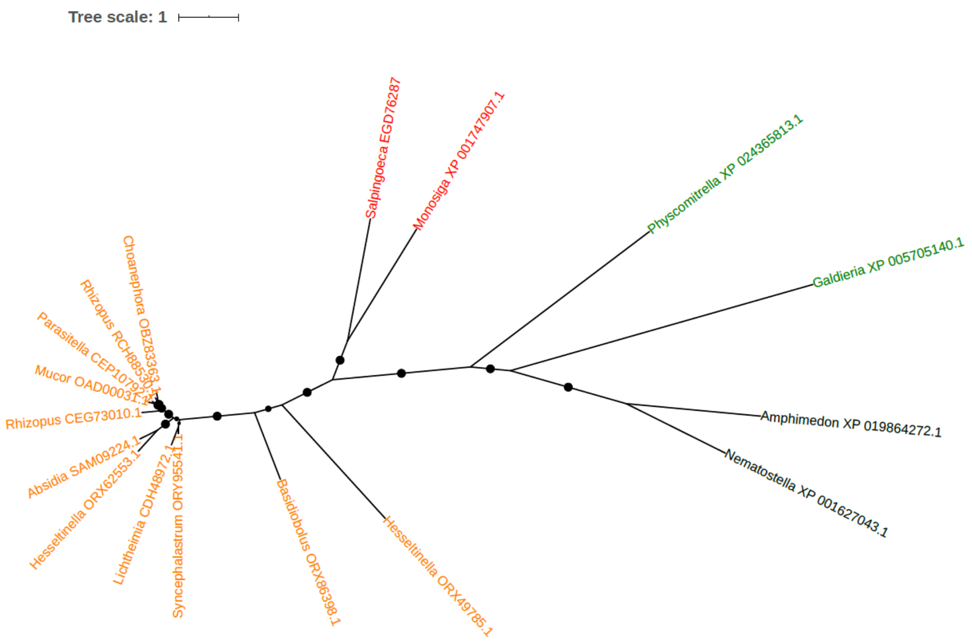


Figure S120. Phylogeny of L-2-hydroxyglutarate dehydrogenase (EGD76287)

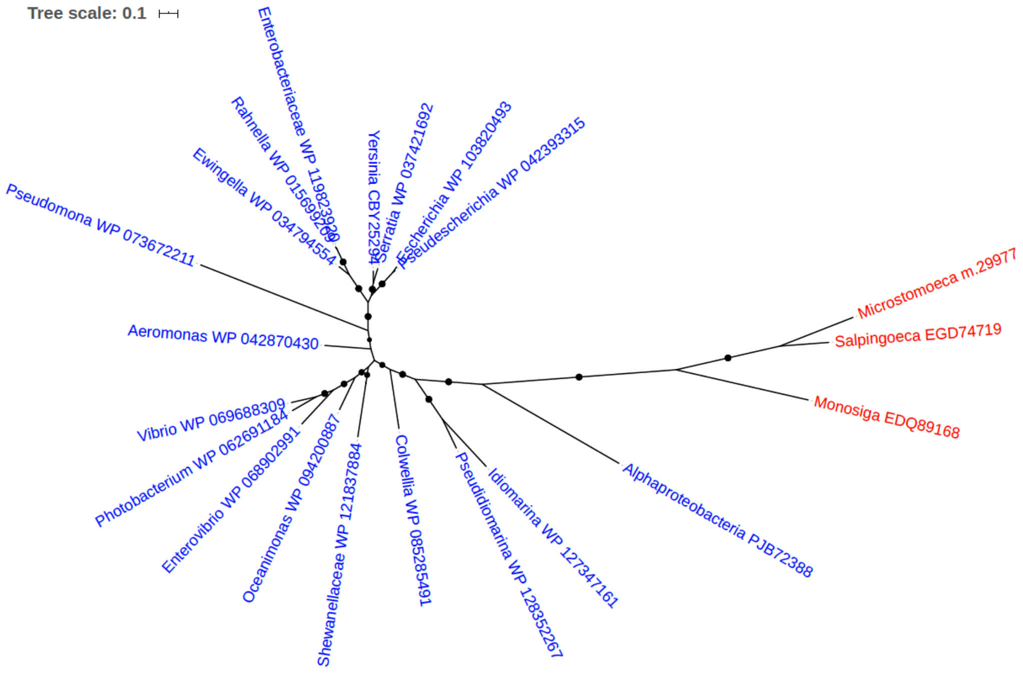


Figure S121. Phylogeny of L-threonine 3-dehydrogenase (EGD74719)

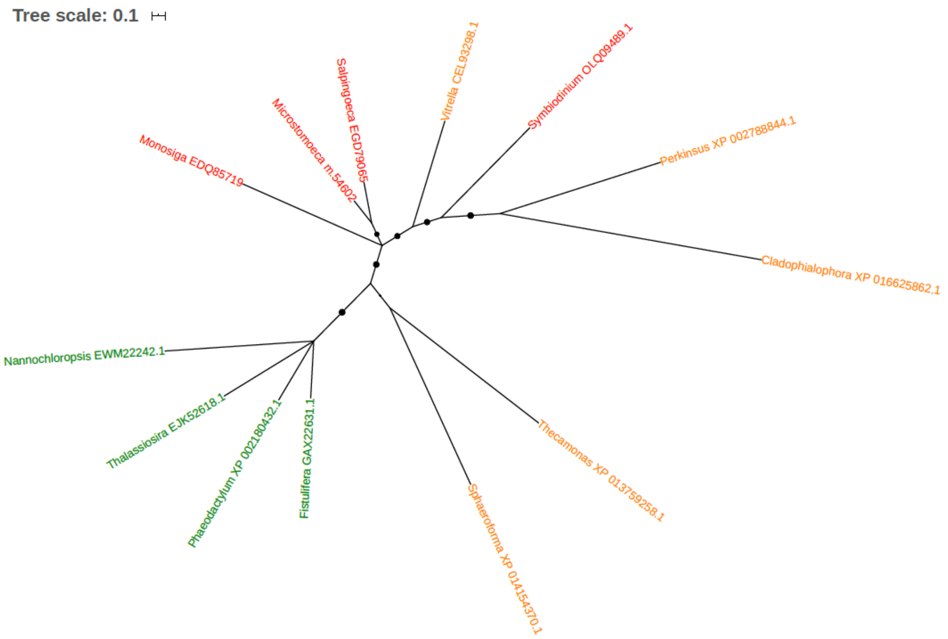


Figure S122. Phylogeny of Lactoylglutathione lyase (EC 4.4.1.5) (Glyoxalase I) (EGD79065)

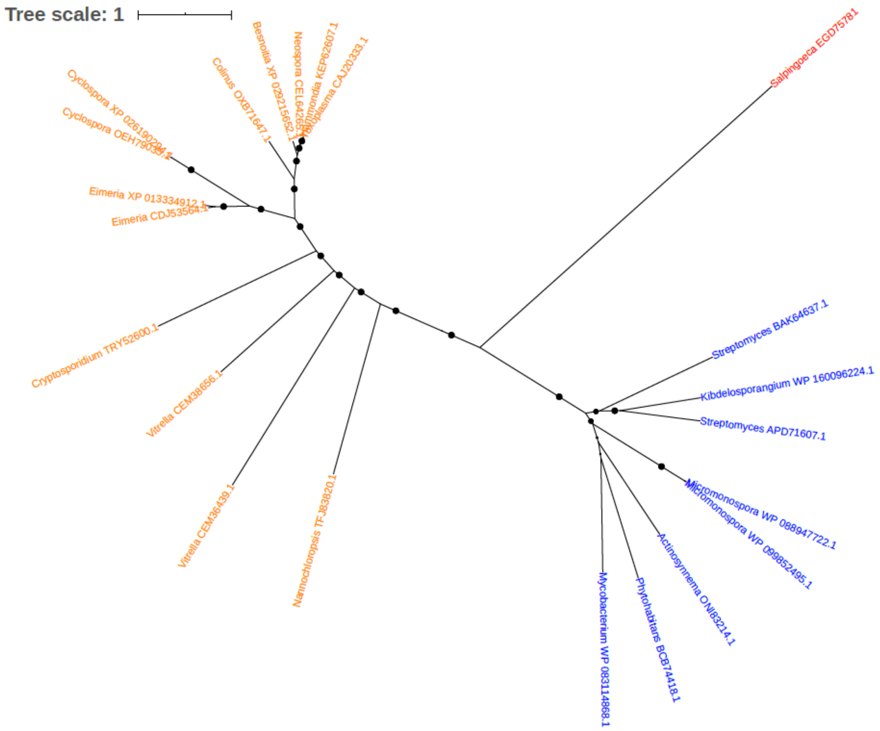


Figure S123. Phylogeny of Lankamycin synthase (EGD75781)

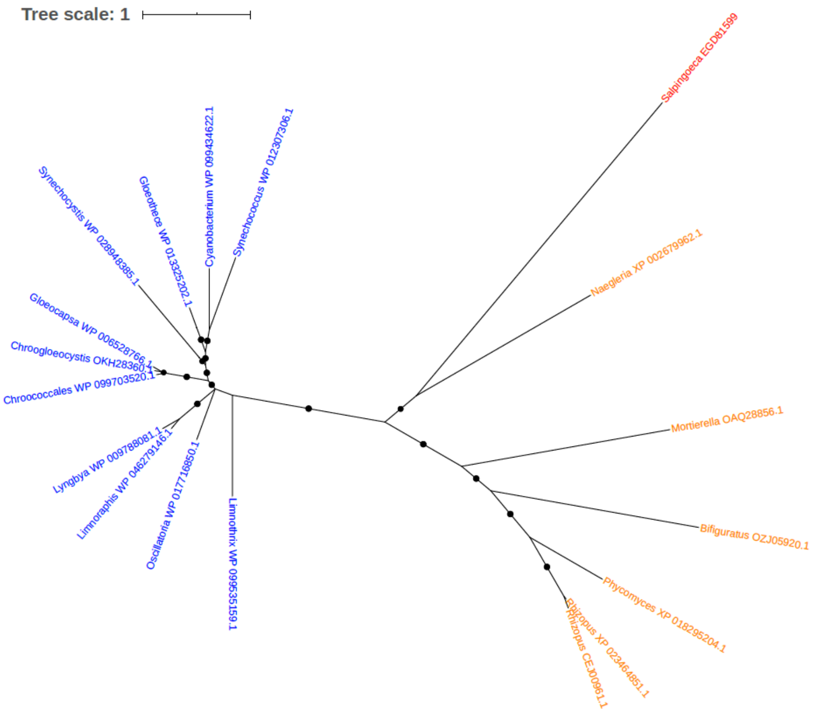


Figure S124. Phylogeny of Leucyl-tRNA synthetase (EGD81599)

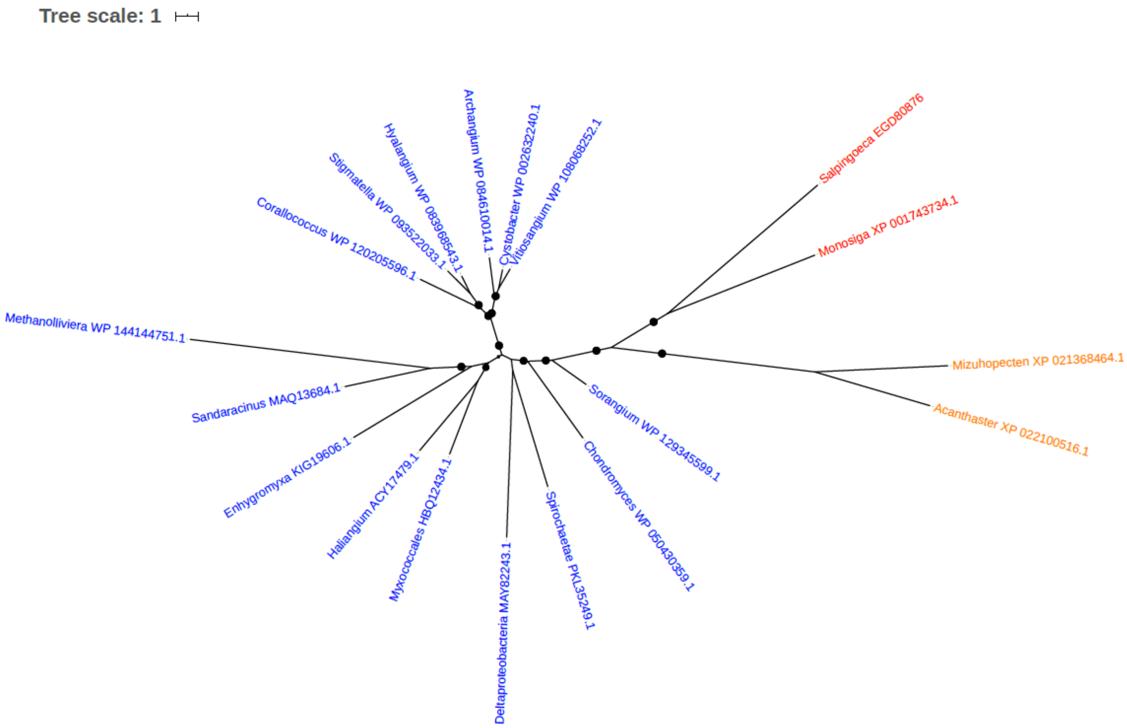


Figure S125. Phylogeny of Major facilitator superfamily transporter permease (EGD80876)

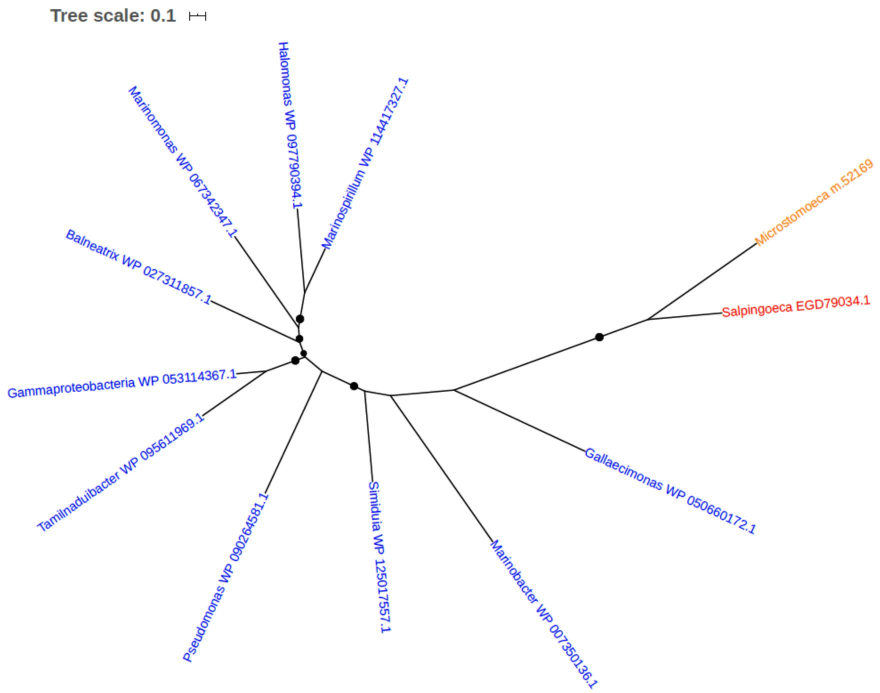


Figure S126. Phylogeny of Malic enzyme (EGD79034)

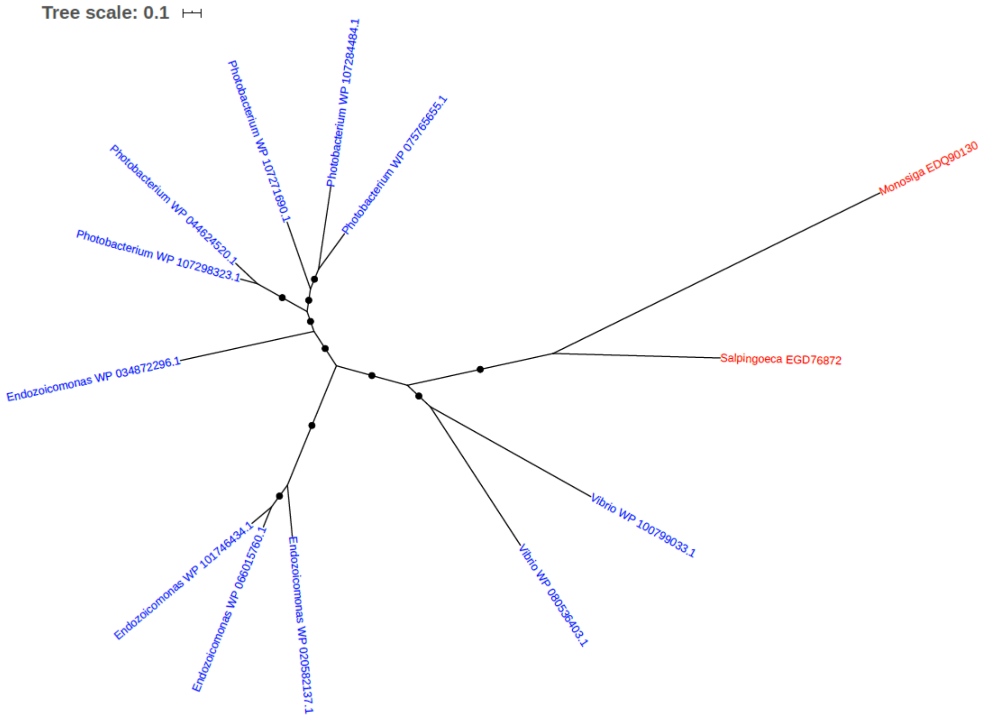


Figure S127. Phylogeny of Maltodextrin glucosidase (EGD76872)

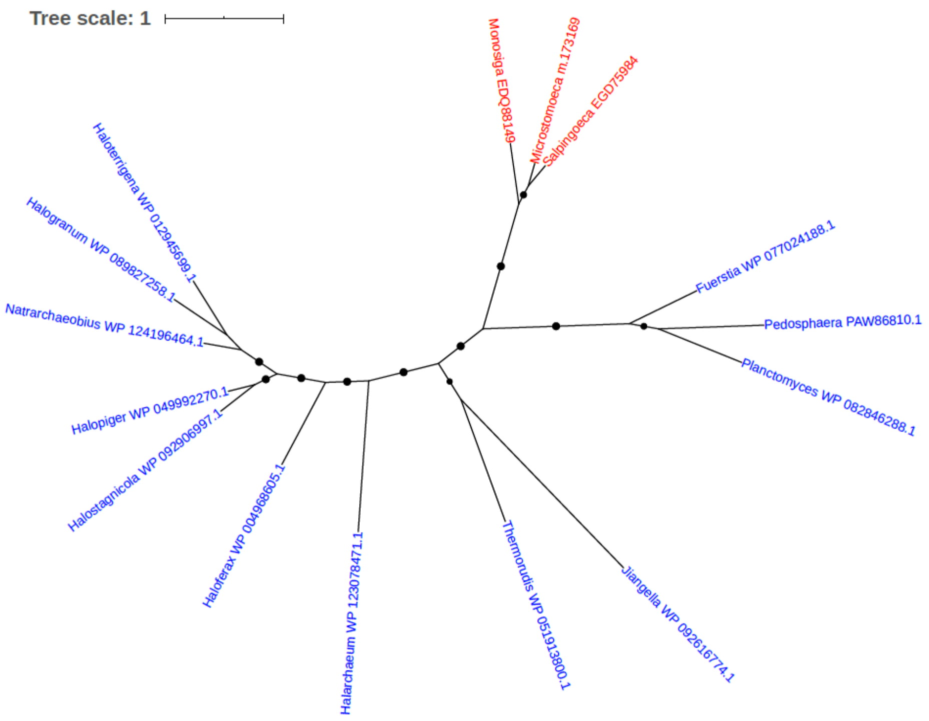


Figure S128. Phylogeny of Mannonate dehydratase (EGD75984)

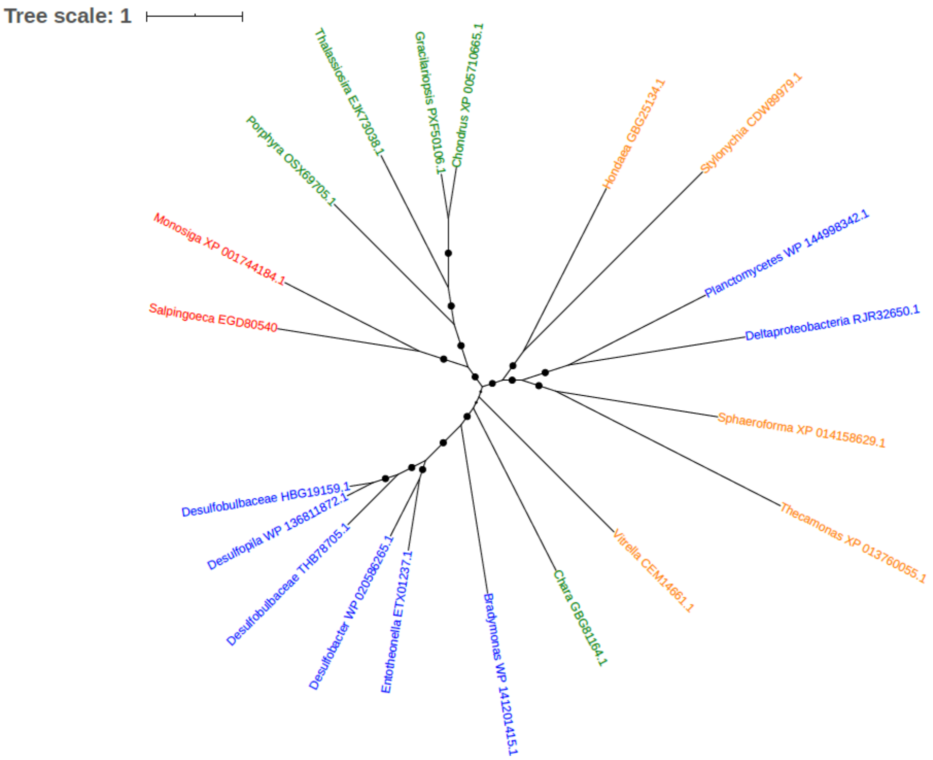


Figure S129. Phylogeny of Mercuric reductase (EGD80540)

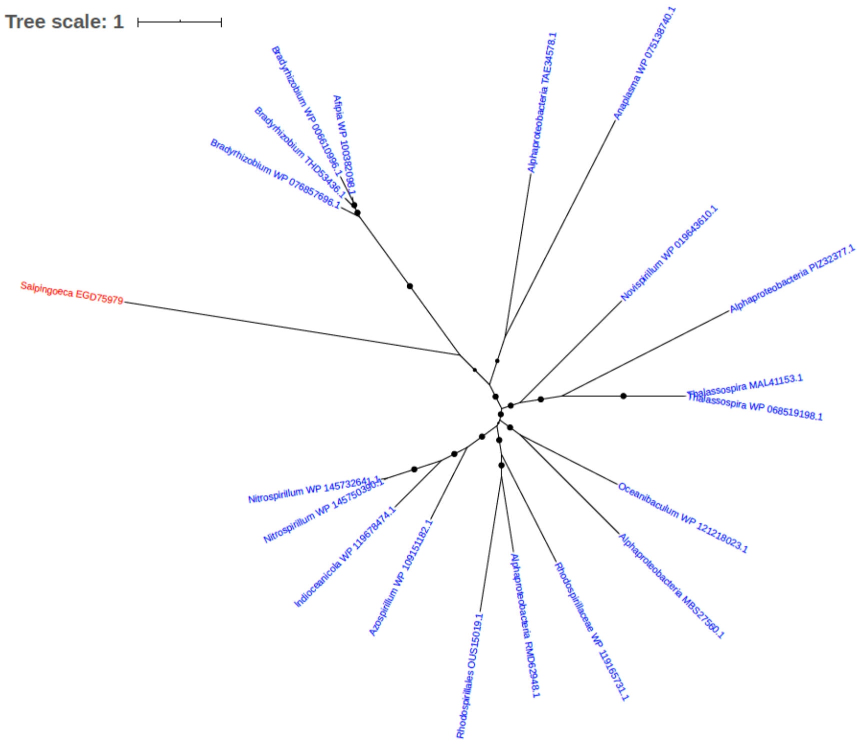


Figure S130. Phylogeny of Metallo-beta-lactamase superfamily (EGD75979)

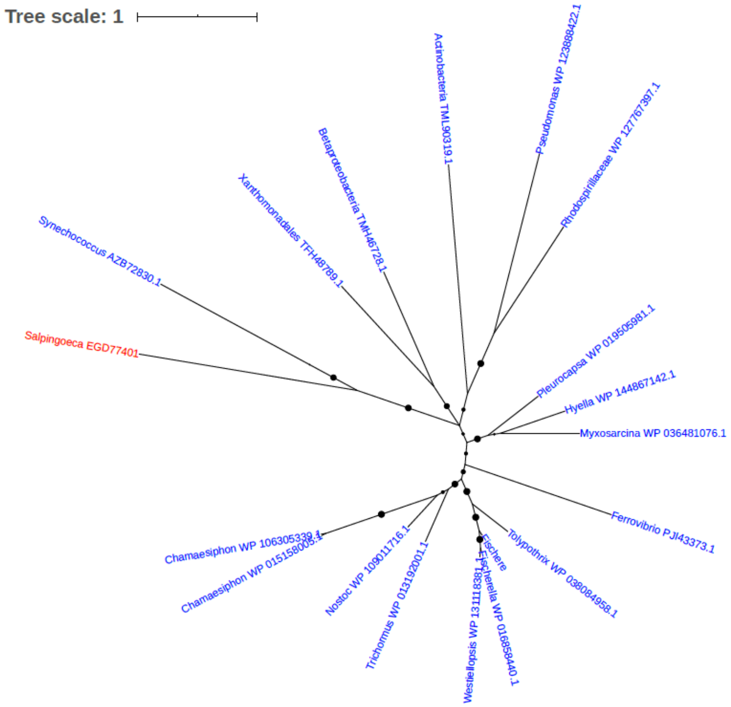


Figure S131. Phylogeny of Methionine-S-sulfoxide reductase (EGD77401)

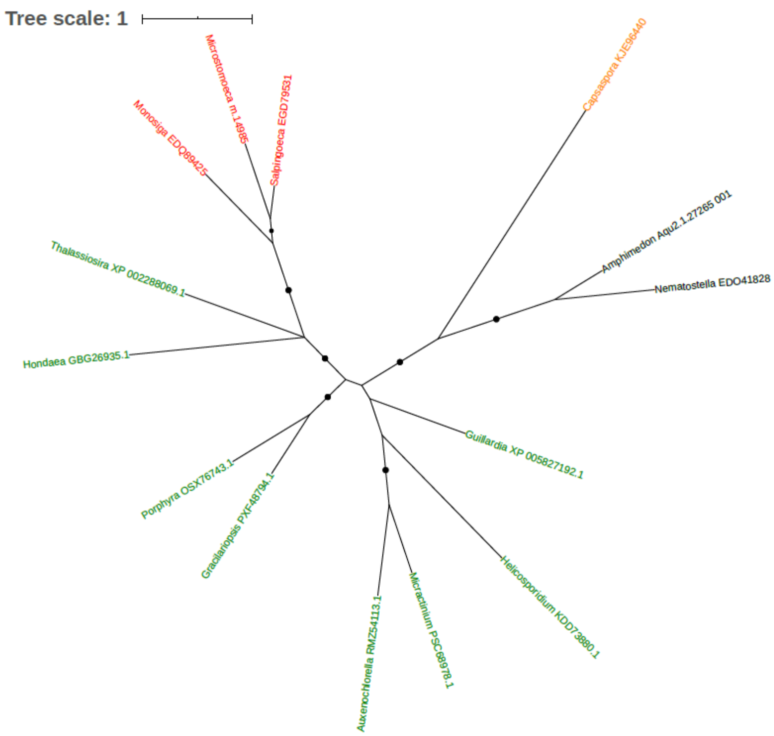


Figure S132. Phylogeny of Methylenetetrahydrofolate reductase (EGD79531)

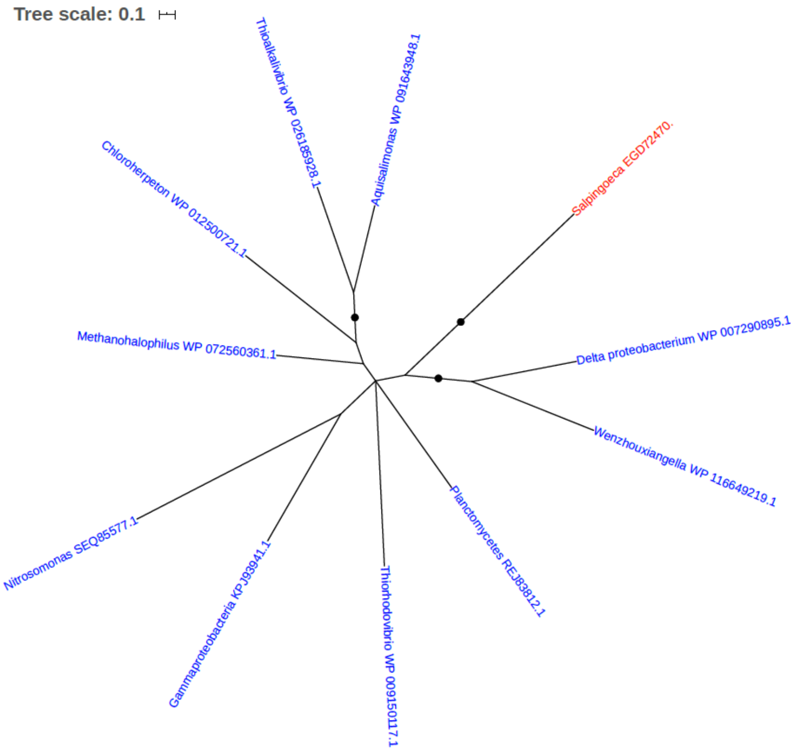


Figure S133. Phylogeny of Methyltransferase type 11 (EGD72470)

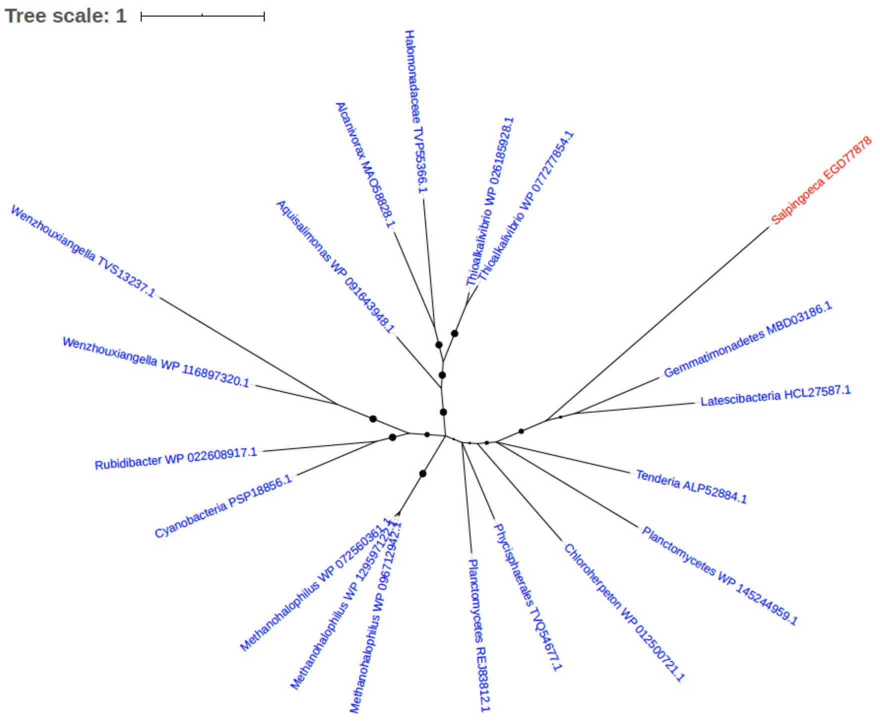


Figure S134. Phylogeny of Methyltransferase type 11 (EGD77878)

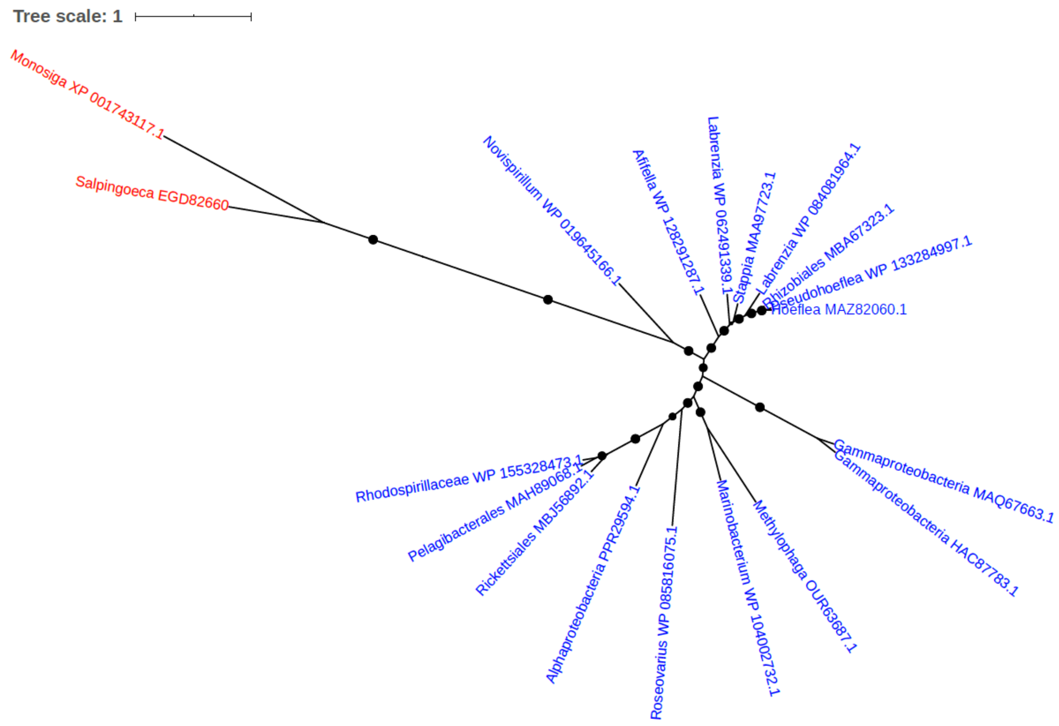


Figure S135. Phylogeny of MFS permease (EGD82660)

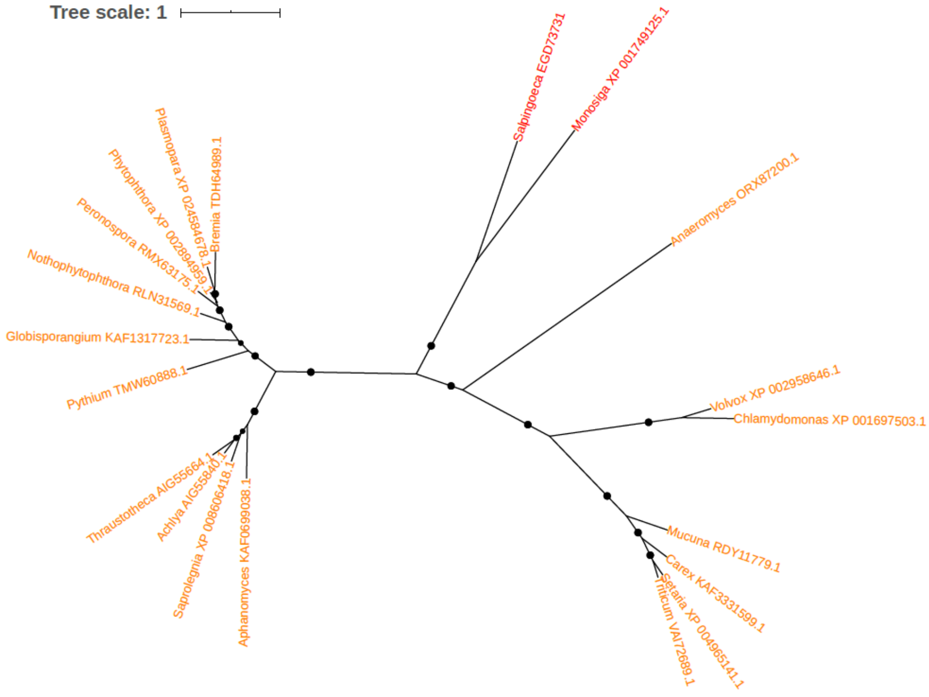


Figure S136. Phylogeny of Mitochondrion protein (EGD73731)

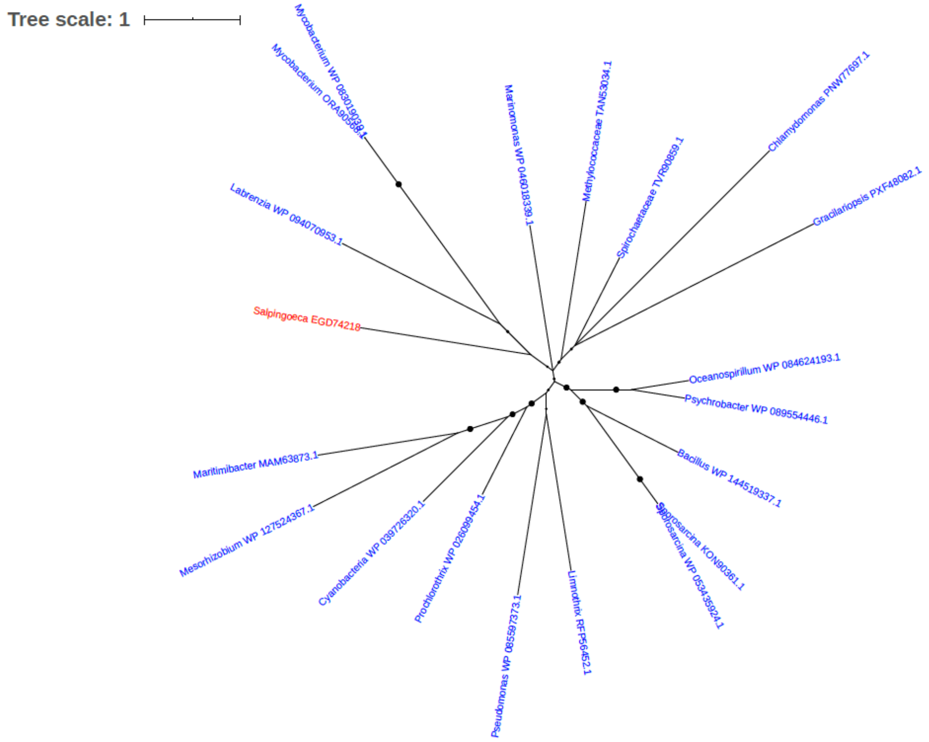


Figure S137. Phylogeny of Monoamine oxidase (EGD74218)

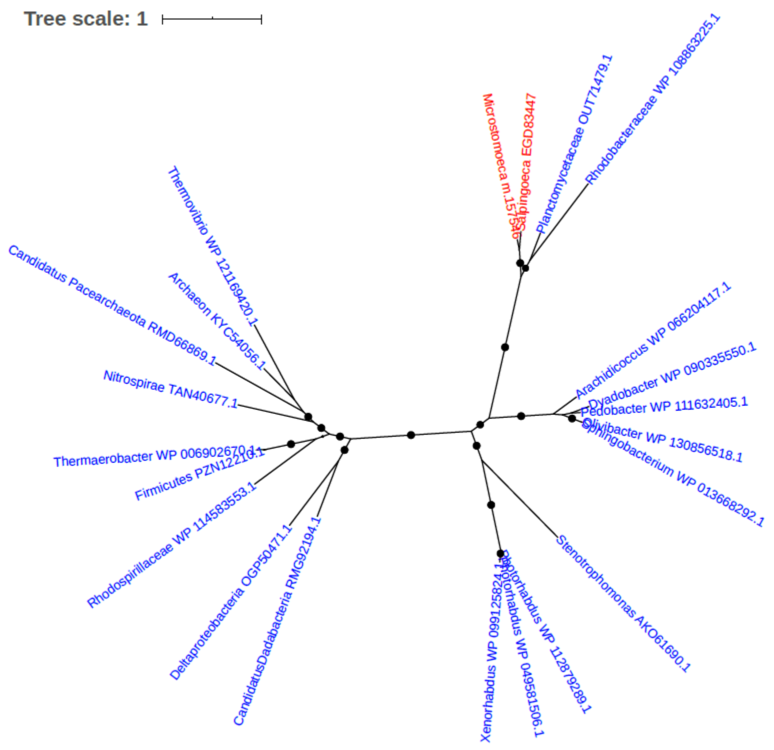


Figure S138. Phylogeny of Myo-inositol-1-phosphate synthase (EGD83447)

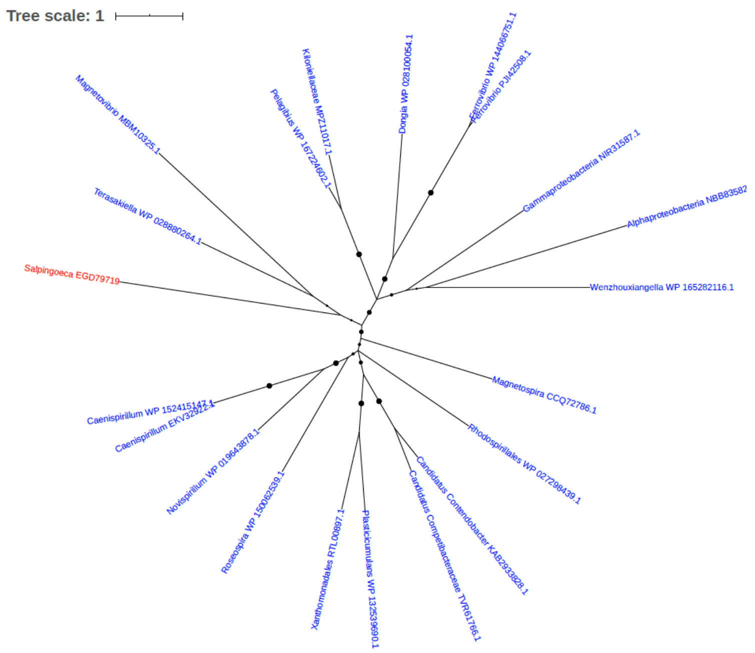


Figure S139. Phylogeny of N-formylglutamate amidohydrolase (EGD79719)

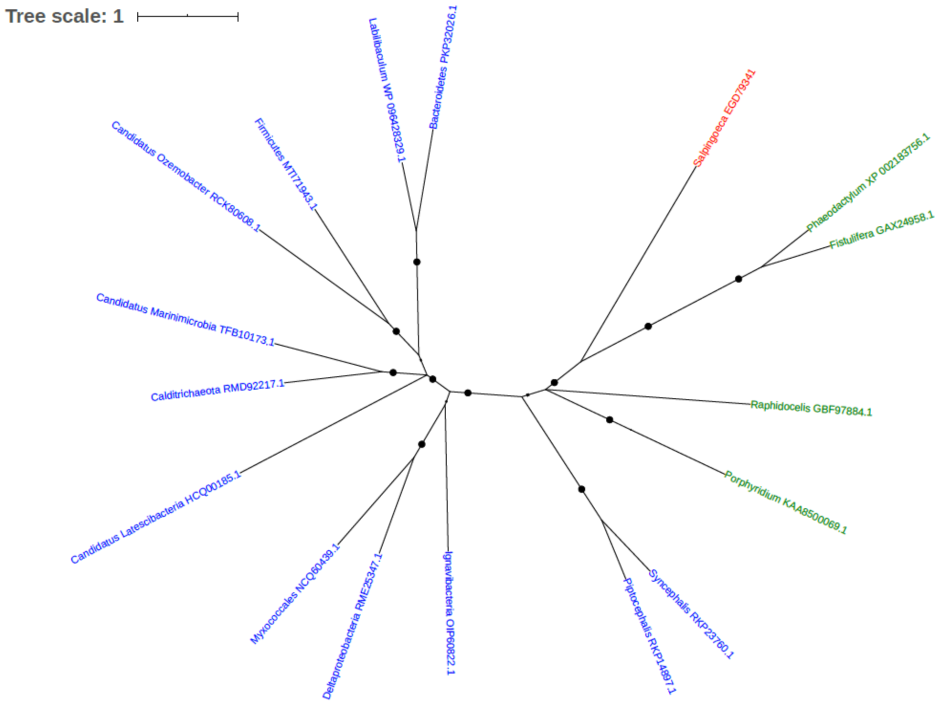


Figure S140. Phylogeny of Na+/H+ antiporter family protein (EGD79341)

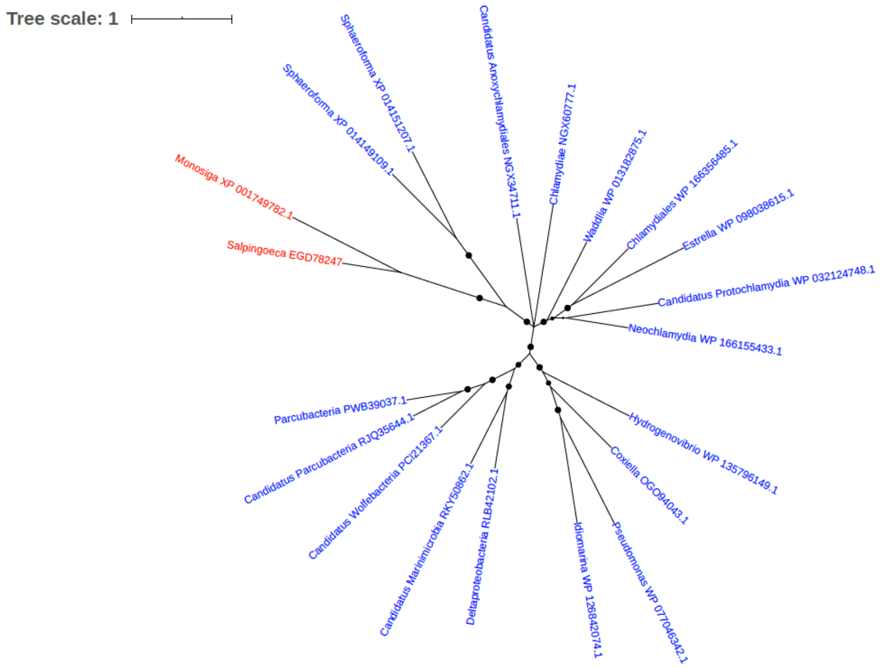


Figure S141. Phylogeny of NAD-dependent epimerase/dehydratase (EGD78247)

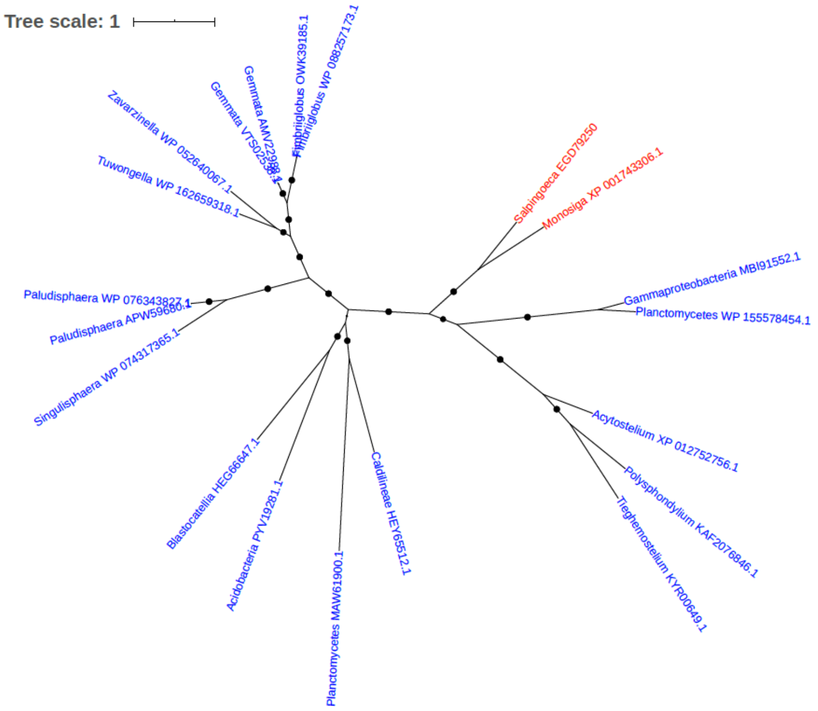


Figure S142. Phylogeny of NAD-dependent epimerase/dehydratase (EGD79250)

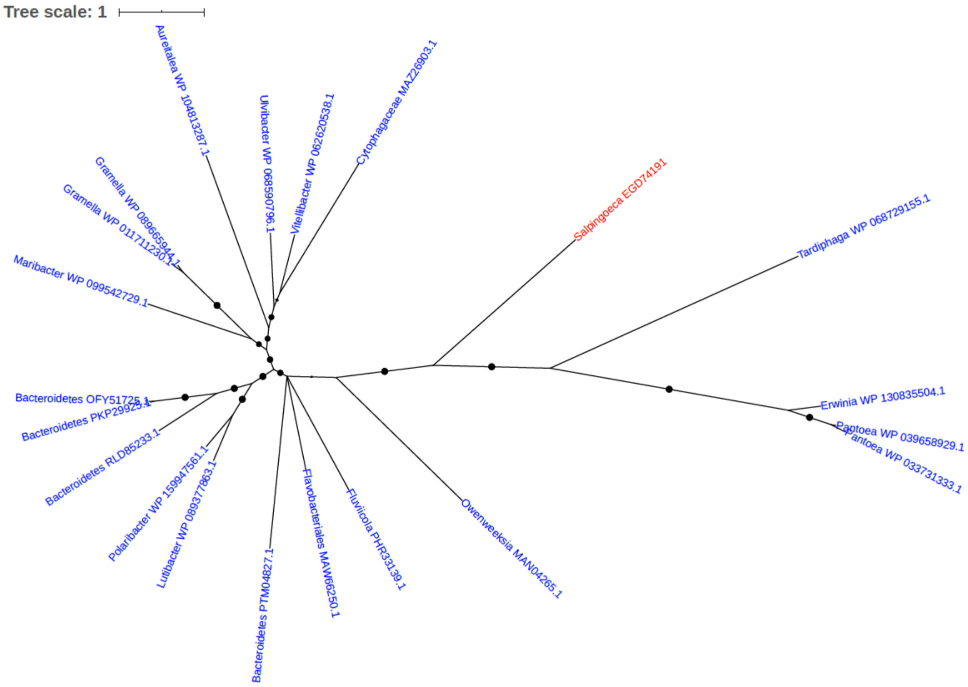


Figure S143. Phylogeny of NAD-dependent epimerase/dehydratase (EGD74191)

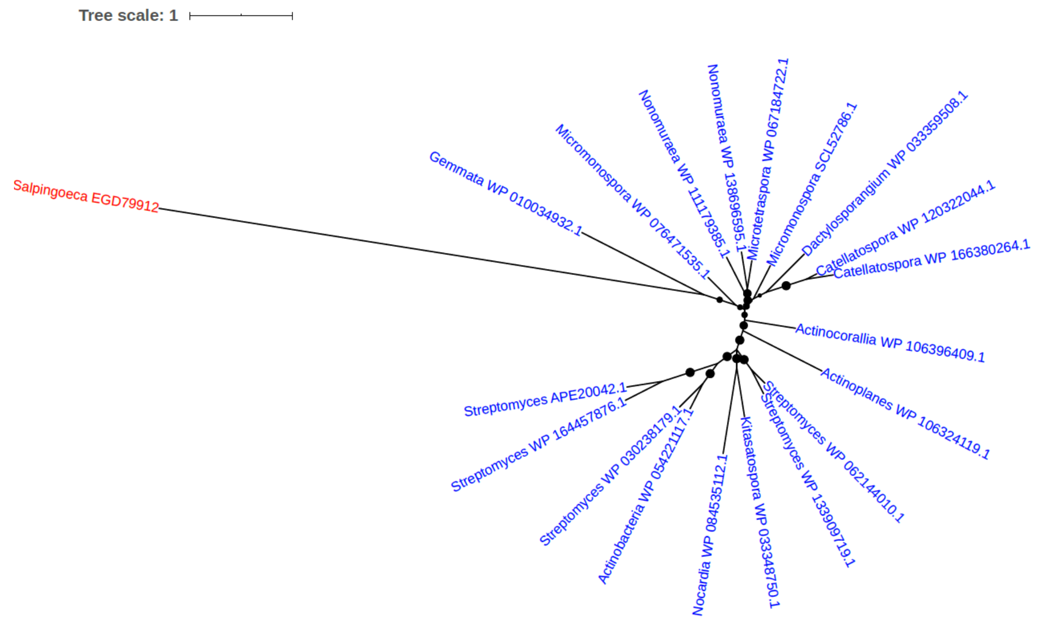


Figure S144. Phylogeny of NAD(P)(+)--arginine ADP-ribosyltransferase (EC 2.4.2.31) (Mono(ADP-ribosyl)transferase) (EGD79912)

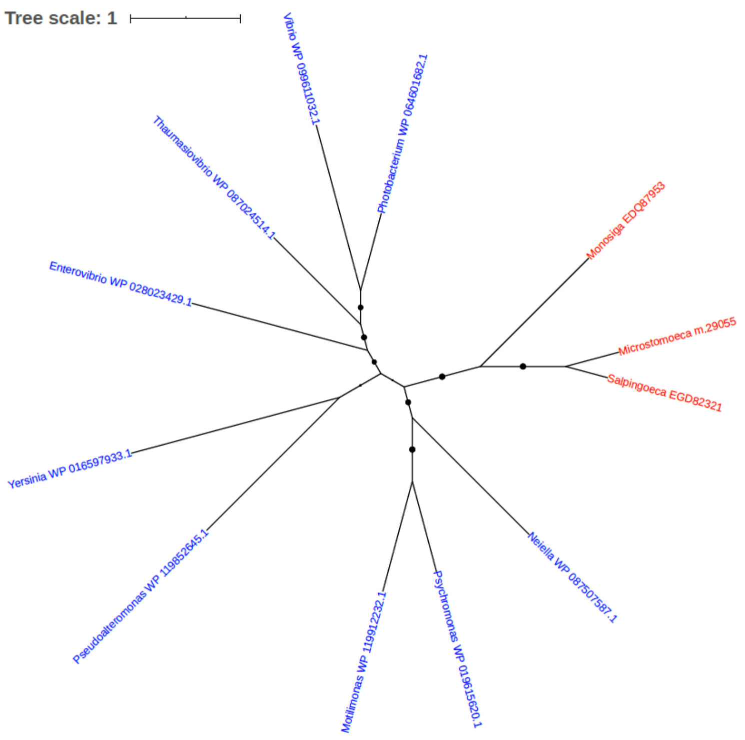


Figure S145. Phylogeny of NAD(P)H-hydrate epimerase (EC 5.1.99.6) (NAD(P)HX epimerase) (EGD82321)

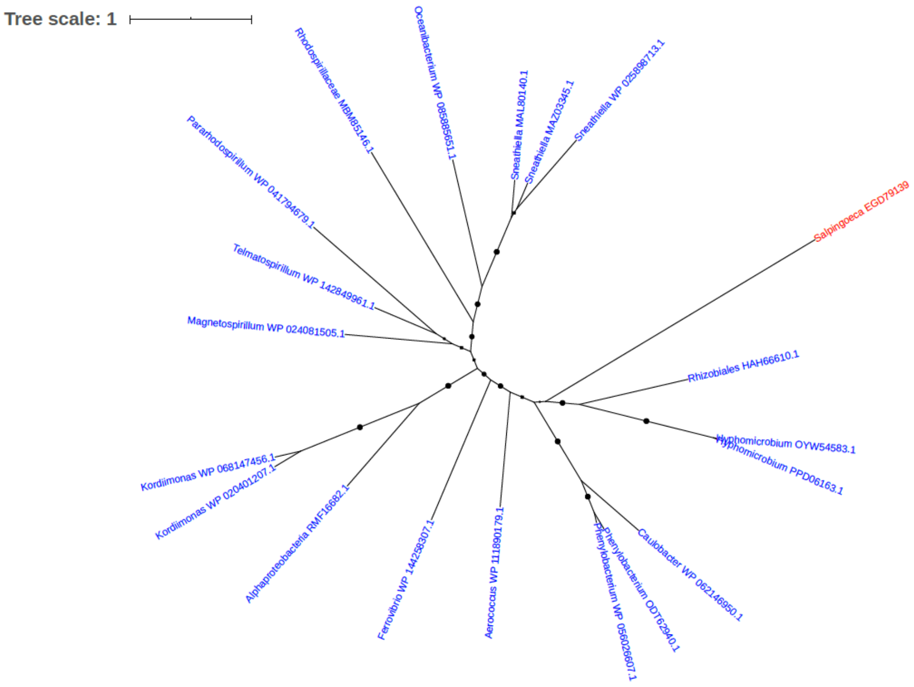


Figure S146. Phylogeny of NAD+ kinase (EGD79139)

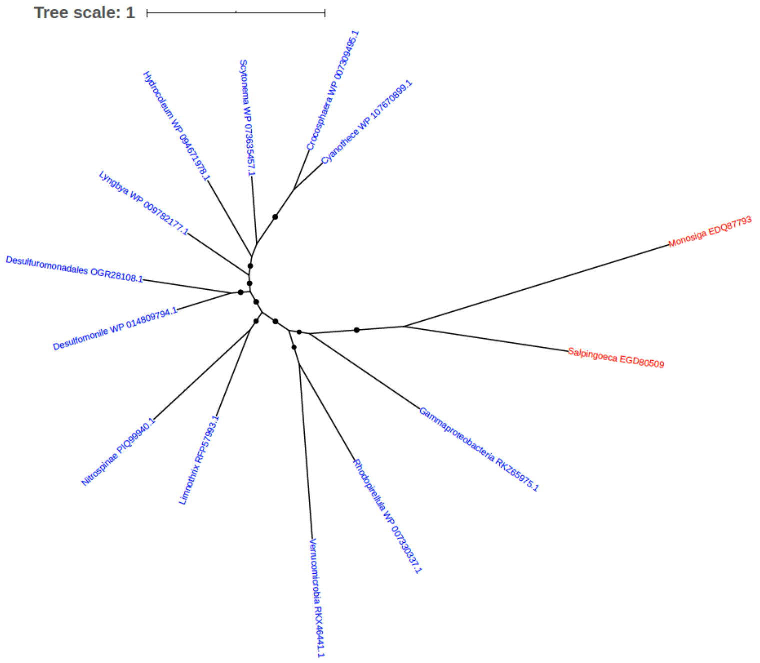


Figure S147. Phylogeny of NADH:flavin oxidoreductase (EGD80509)

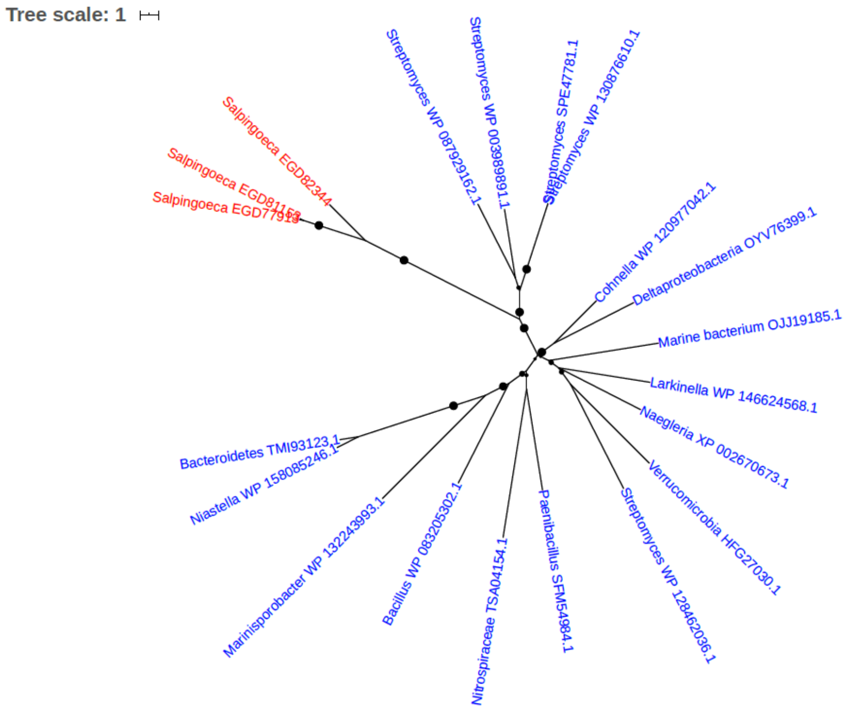


Figure S148. Phylogeny of NHL repeat protein (EGD77913, EGD81153, EGD82344)

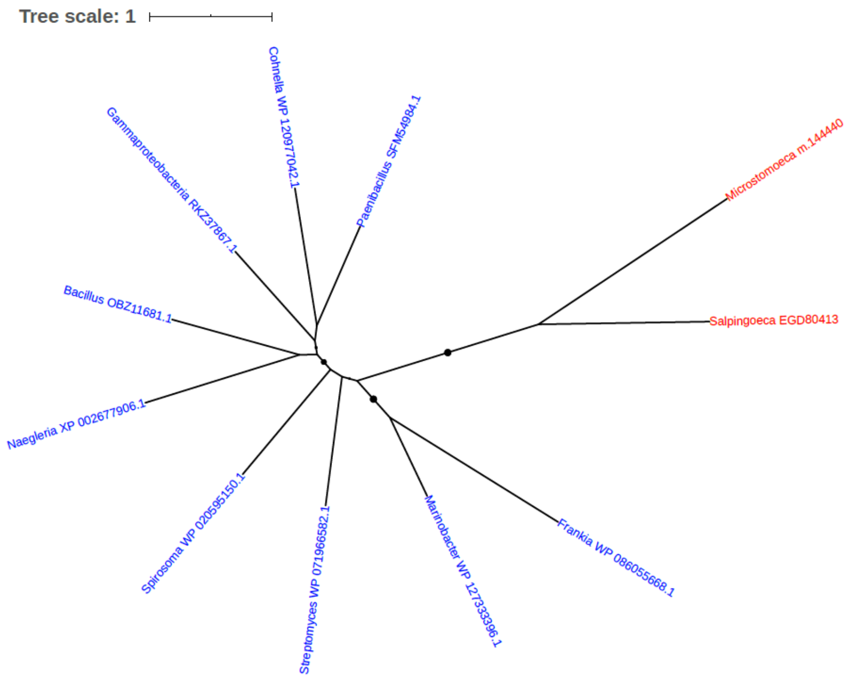


Figure S149. Phylogeny of NHL repeat-containing protein (EGD80413)

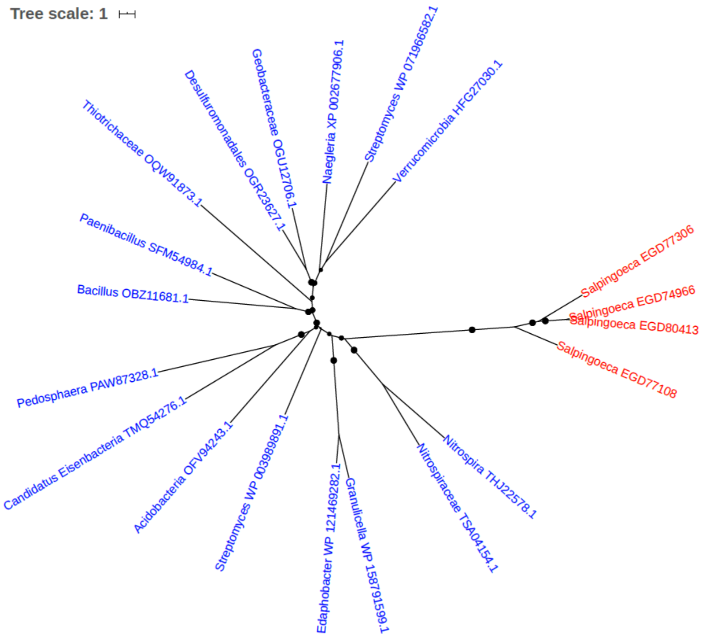


Figure S150. Phylogeny of NHL repeat-containing protein (EGD77108, EGD80413, EGD74966, EGD77306)

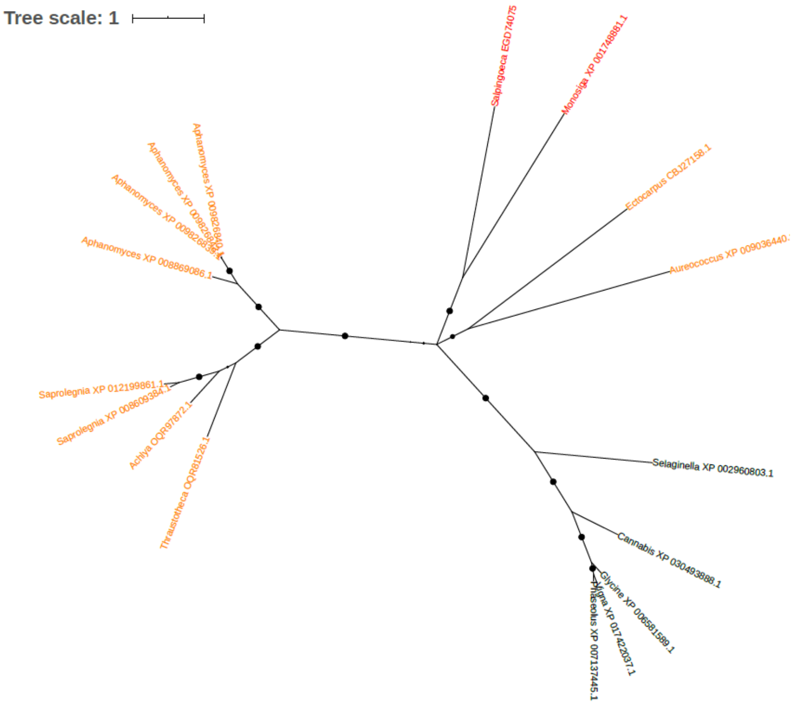


Figure S151. Phylogeny of OGG1 protein type 2c (EGD74075)

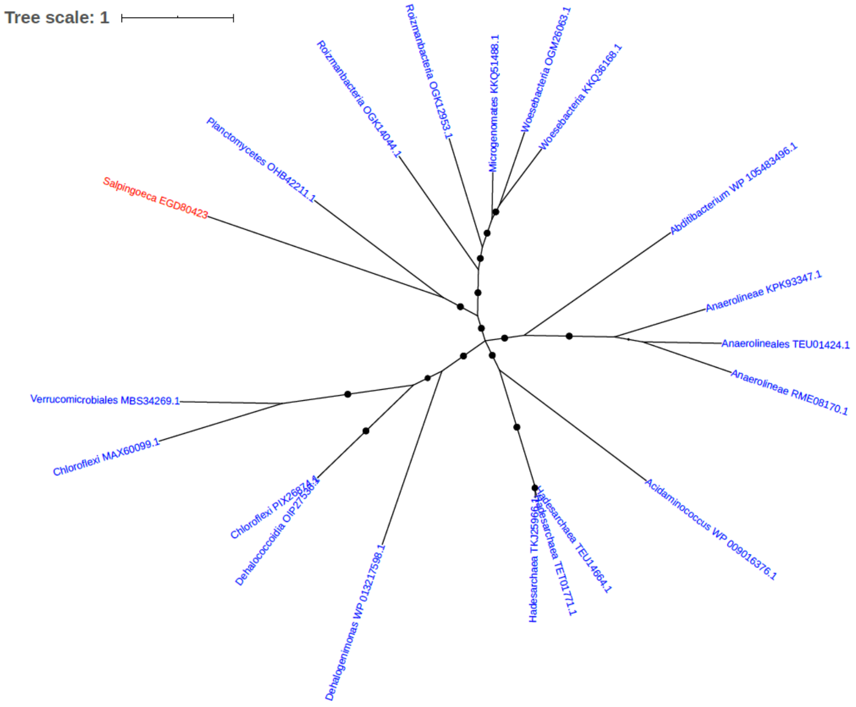


Figure S152. Phylogeny of Ornithine carbamoyltransferase (EGD80423)

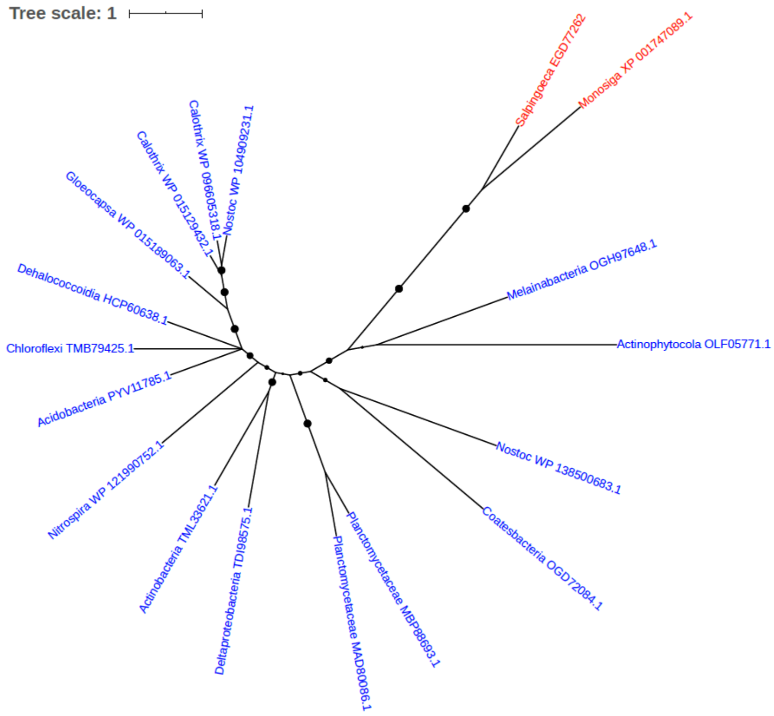


Figure S153. Phylogeny of Oxidoreductase (EGD77262)

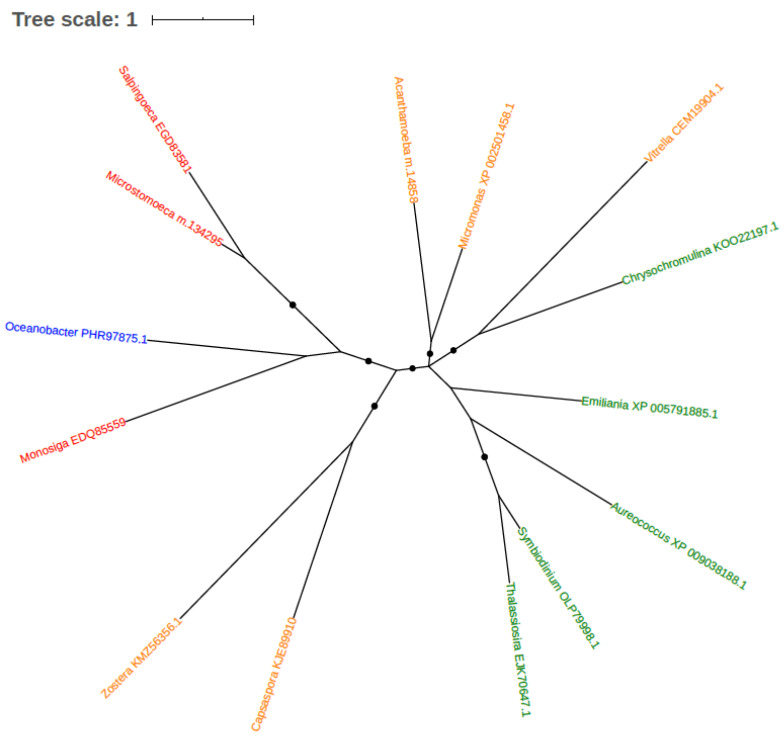


Figure S154. Phylogeny of Oxidoreductase (EGD83581)

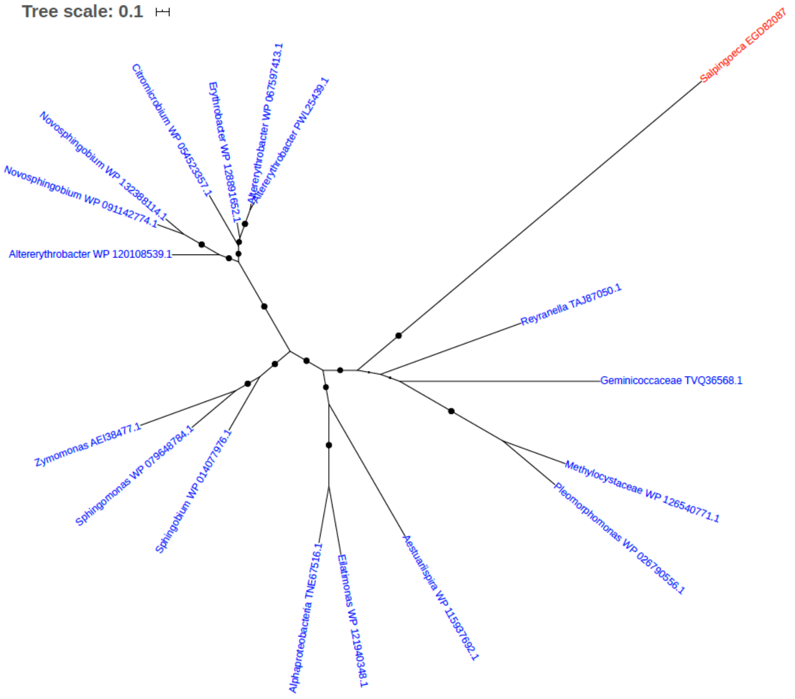


Figure S155. Phylogeny of Peptide chain release factor 2 (EGD82087)

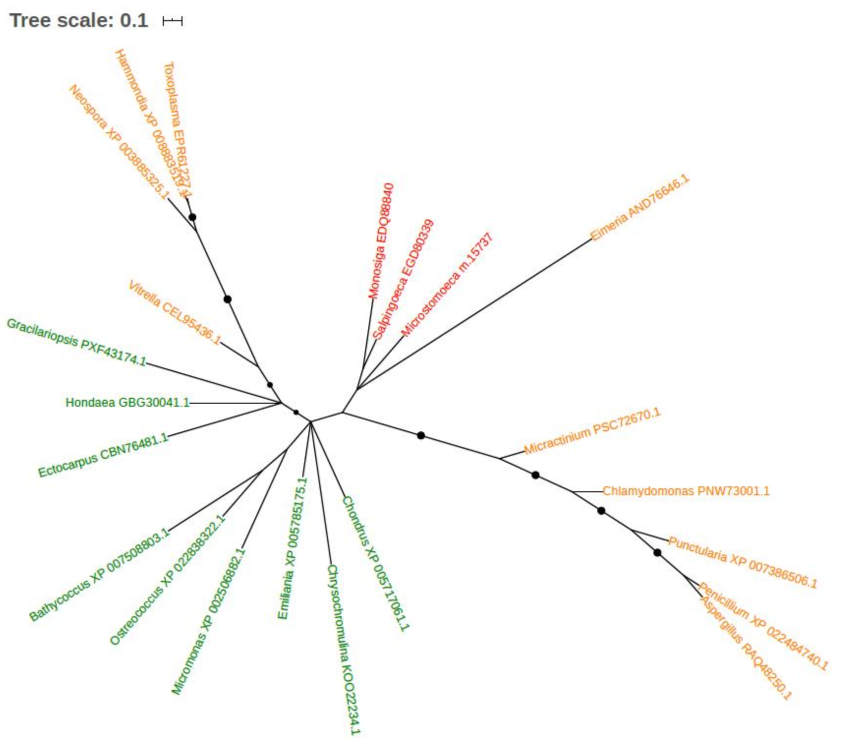


Figure S156. Phylogeny of Peptide-methionine (R)-S-oxide reductase (EC 1.8.4.12) (EGD80339)

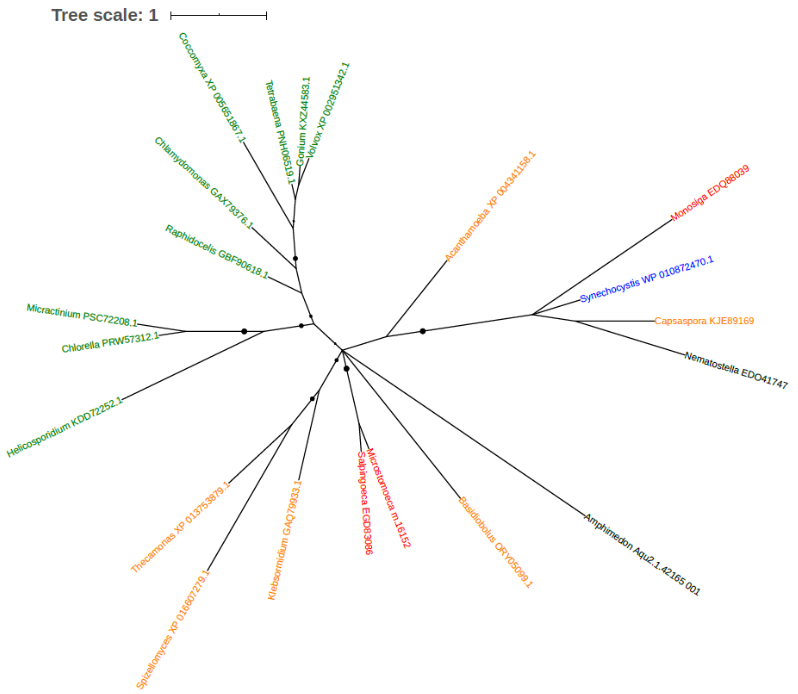


Figure S157. Phylogeny of Peptidyl-prolyl cis-trans isomerase (PPIase) (EC 5.2.1.8) (EGD83086)

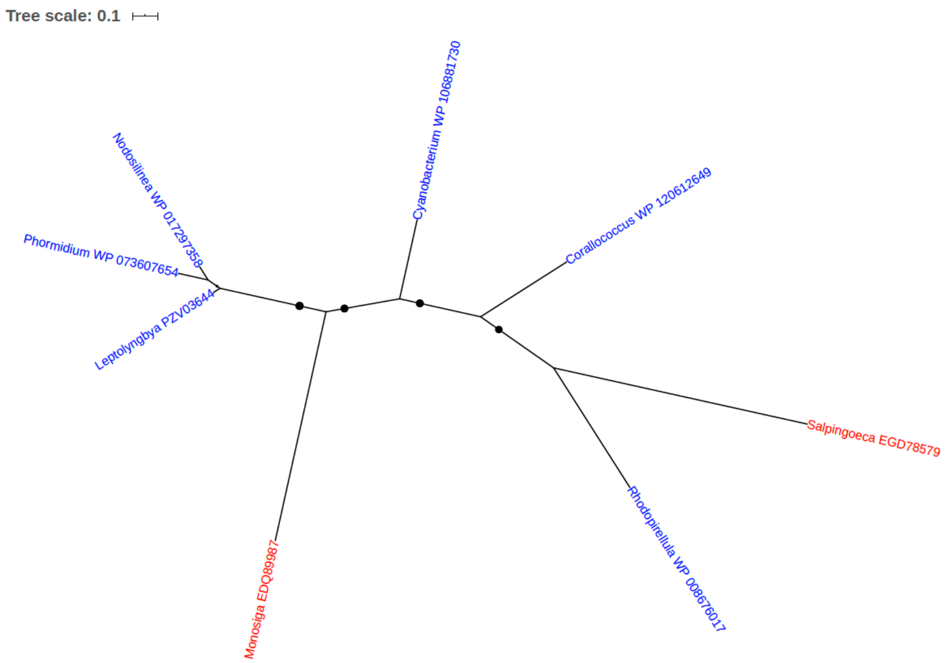


Figure S158. Phylogeny of Peptidyl-prolyl cis-trans isomerase (PPIase) (EGD78579)

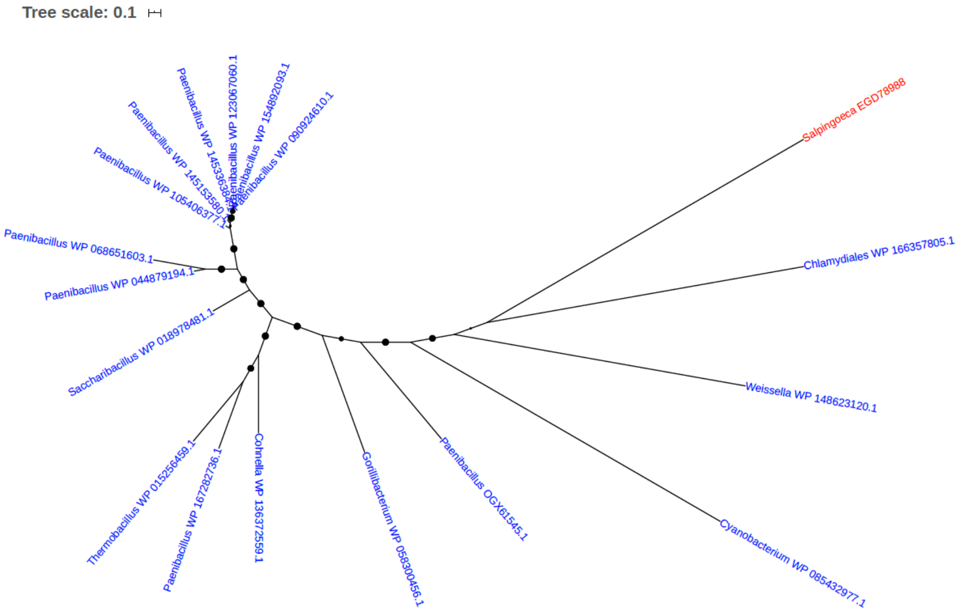


Figure S159. Phylogeny of Peptidyl-tRNA hydrolase (EGD78988)

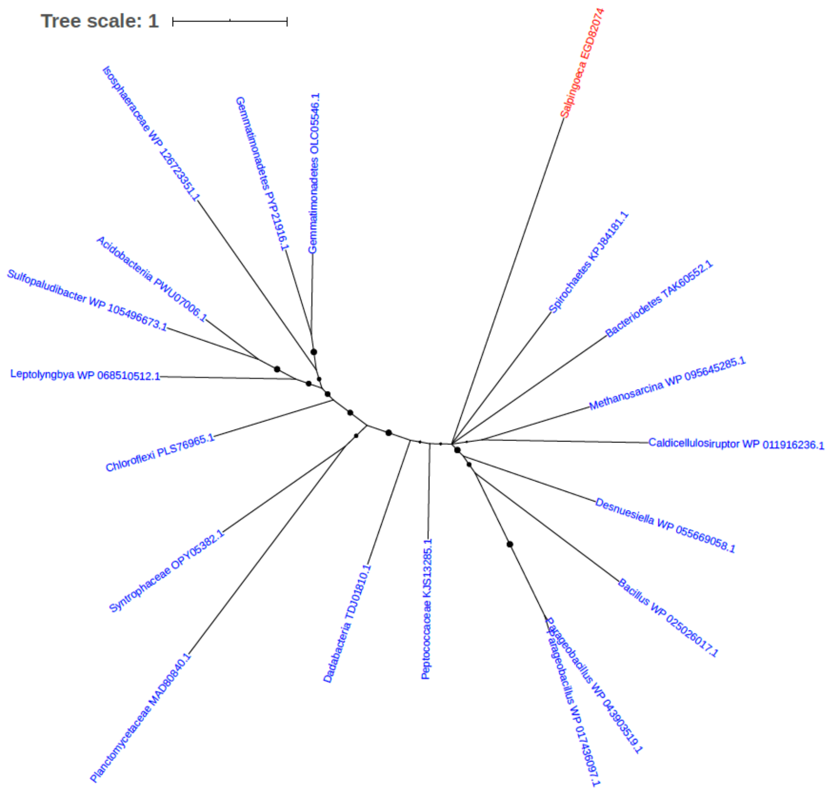


Figure S160. Phylogeny of Phenazine biosynthesis protein PhzF family protein (EGD82074)

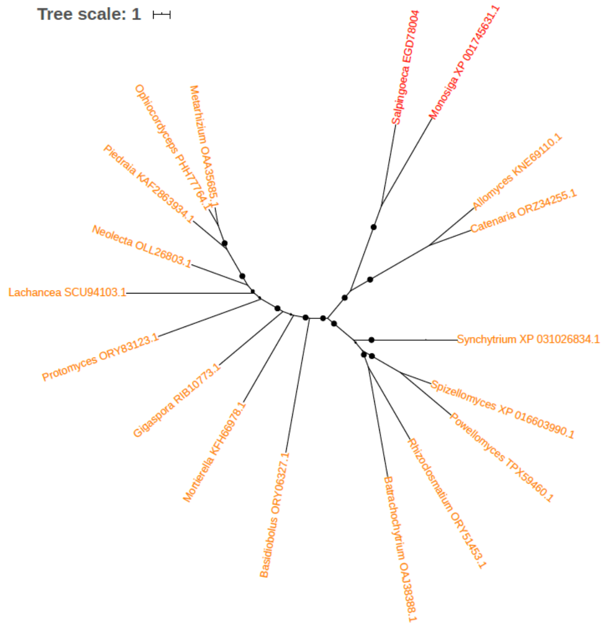


Figure S161. Phylogeny of Phosphatase type 2C (EGD78004)

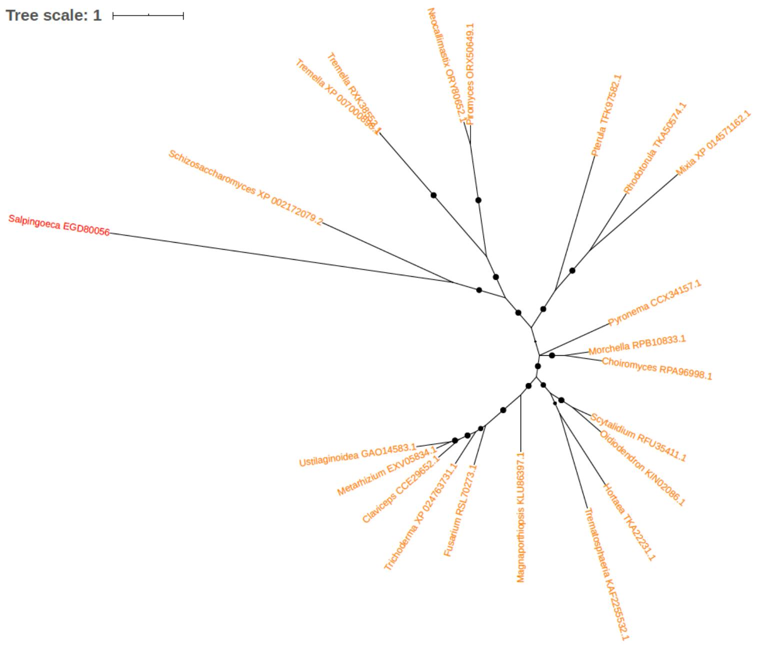


Figure S162. Phylogeny of Phosphatidylethanolamine N-methyltransferase (PEAMT) (PEMT) (EC 2.1.1.17) (EC 2.1.1.71) (Phospholipid methyltransferase) (PLMT) (EGD80056)

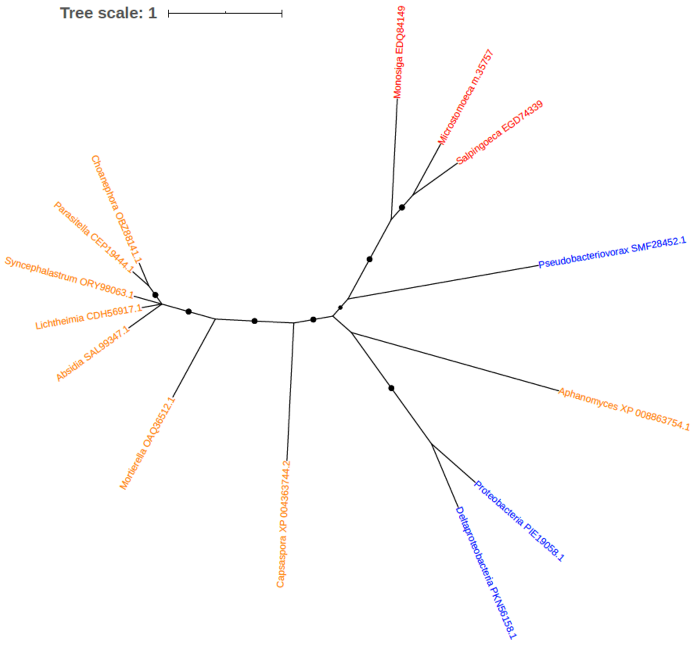


Figure S163. Phylogeny of Phospho-2-dehydro-3-deoxyheptonate aldolase (EC 2.5.1.54) (EGD74339)

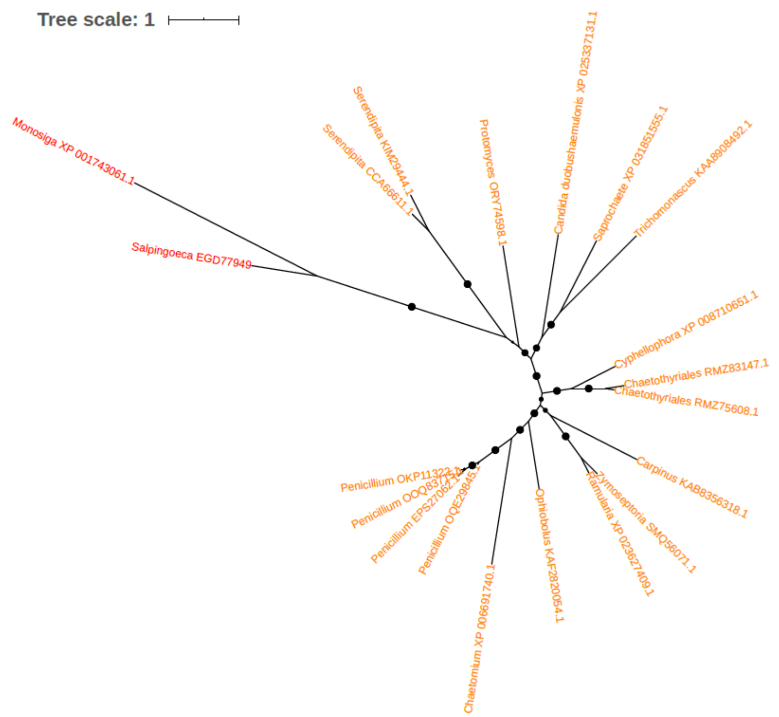


Figure S164. Phylogeny of Phosphoadenosine phosphosulfate reductase (EGD77949)

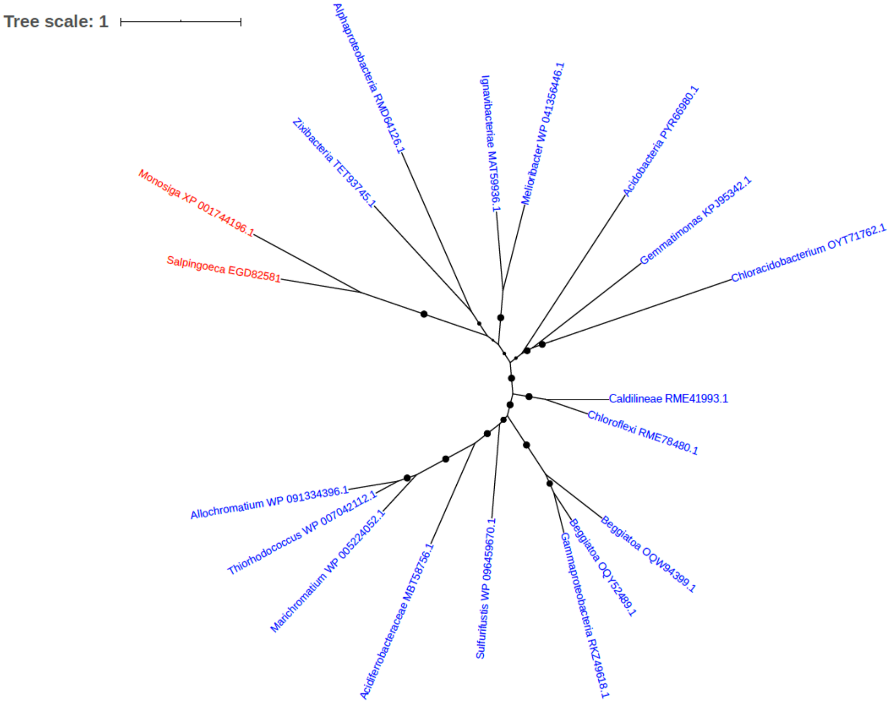


Figure S165. Phylogeny of Phosphoenolpyruvate carboxykinase (EGD82581)

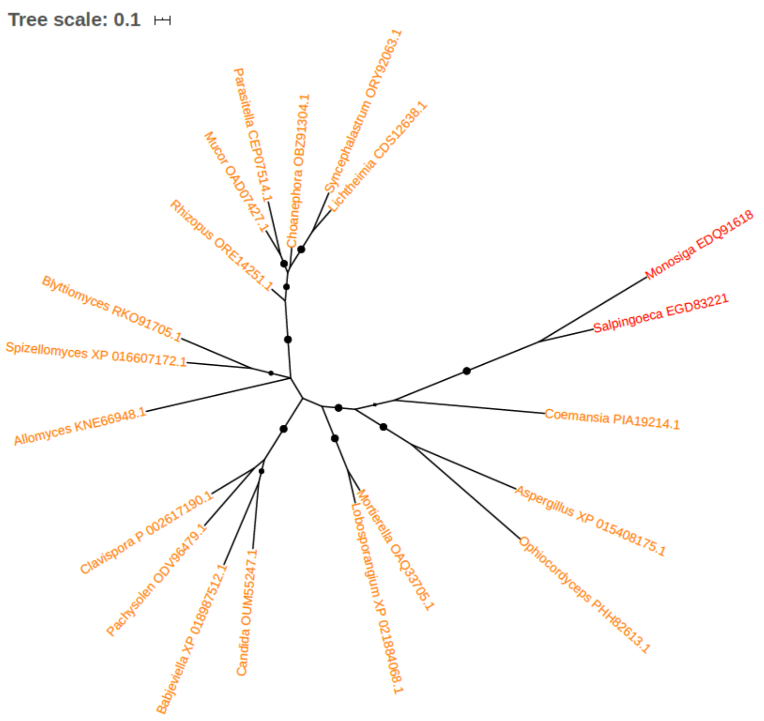


Figure S166. Phylogeny of Phosphomannomutase (EC 5.4.2.8) (EGD83221)

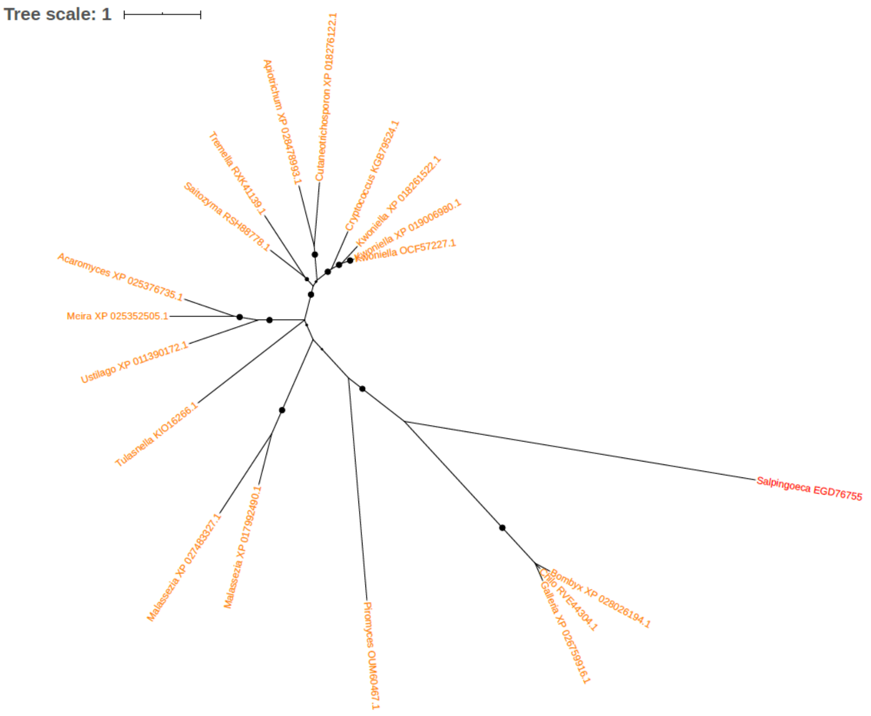


Figure S167. Phylogeny of Phosphopantothenate-cysteine ligase (EGD76755)

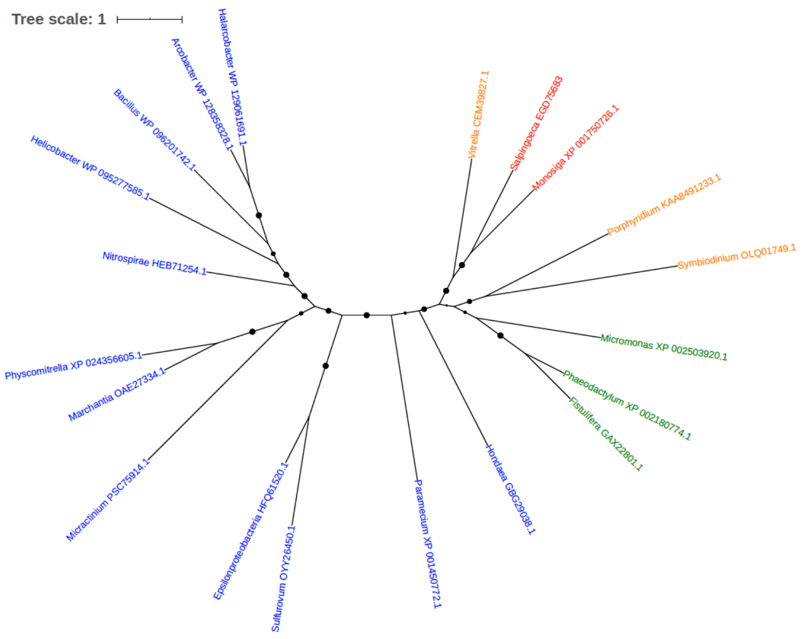


Figure S168. Phylogeny of Pirin (EGD75683)

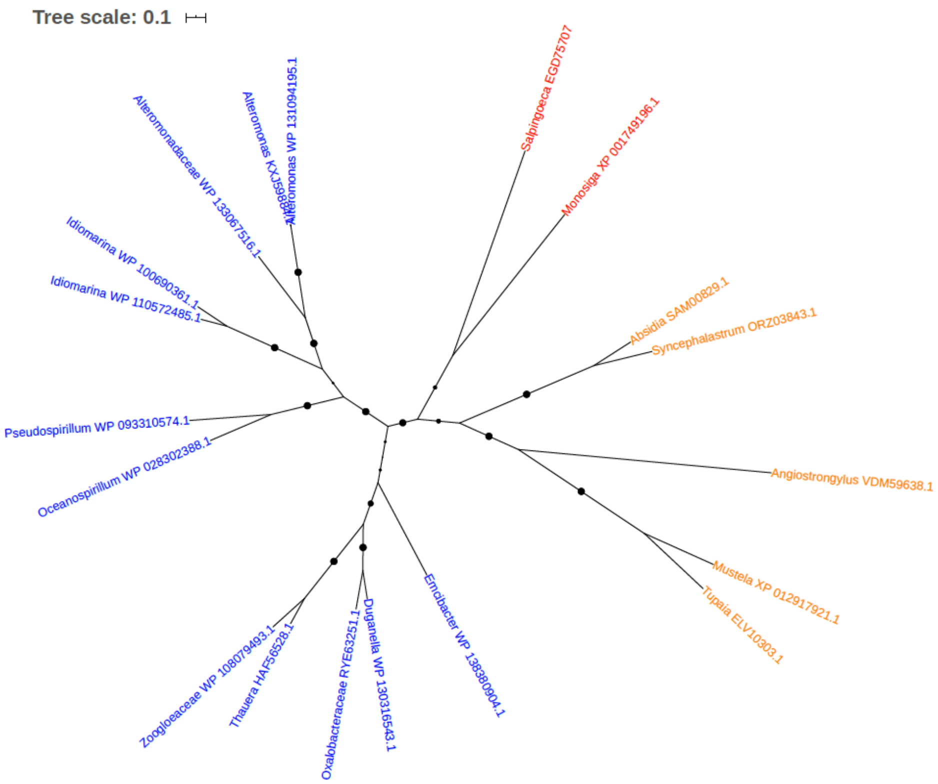


Figure S169. Phylogeny of Propionyl-CoA carboxylase (EGD75707)

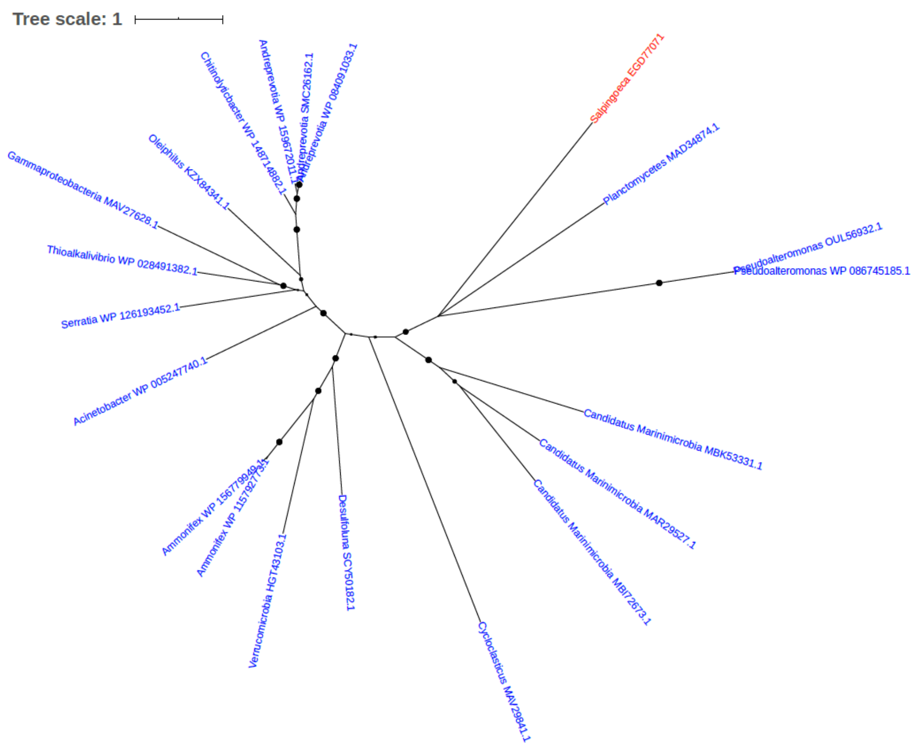


Figure S170. Phylogeny of Pseudouridine synthase (EC 5.4.99.-) (EGD77071)

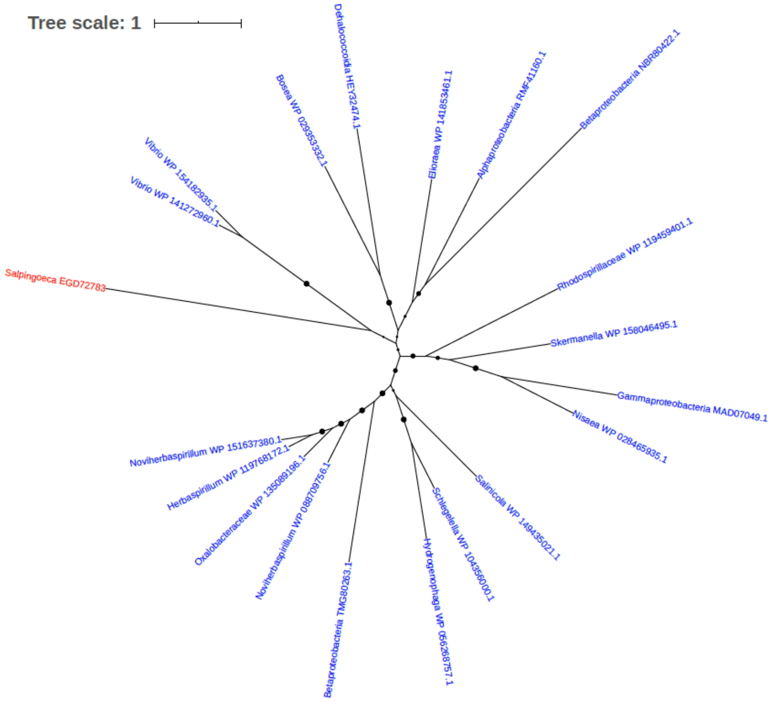


Figure S171. Phylogeny of Putative hydroxypyruvate isomerase (EC 5.3.1.22) (EGD82783)

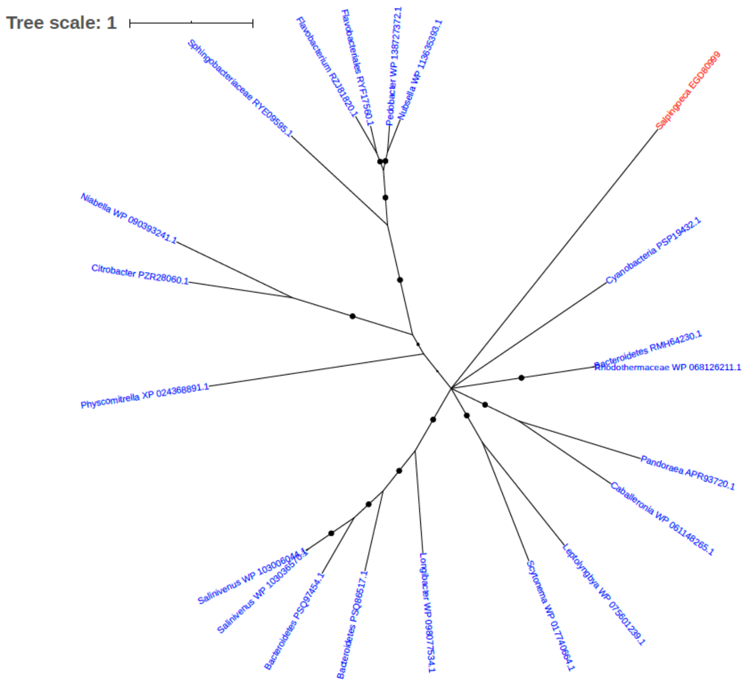


Figure S172. Phylogeny of Pyridoxine/pyridoxamine 5'-phosphate oxidase (EGD80999)

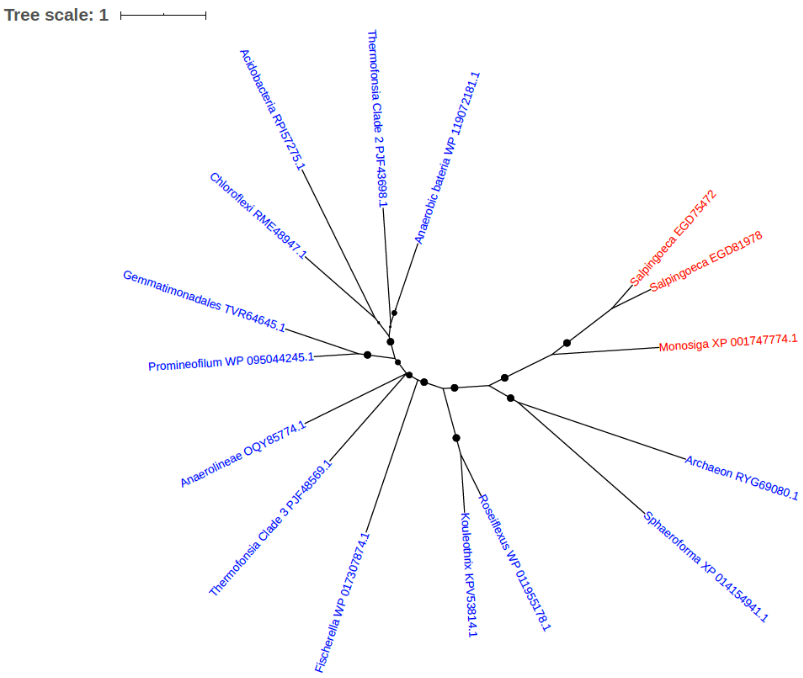


Figure S173. Phylogeny of Pyrroline-5-carboxylate reductase (EC 1.5.1.2) (EGD75472, EGD81978)

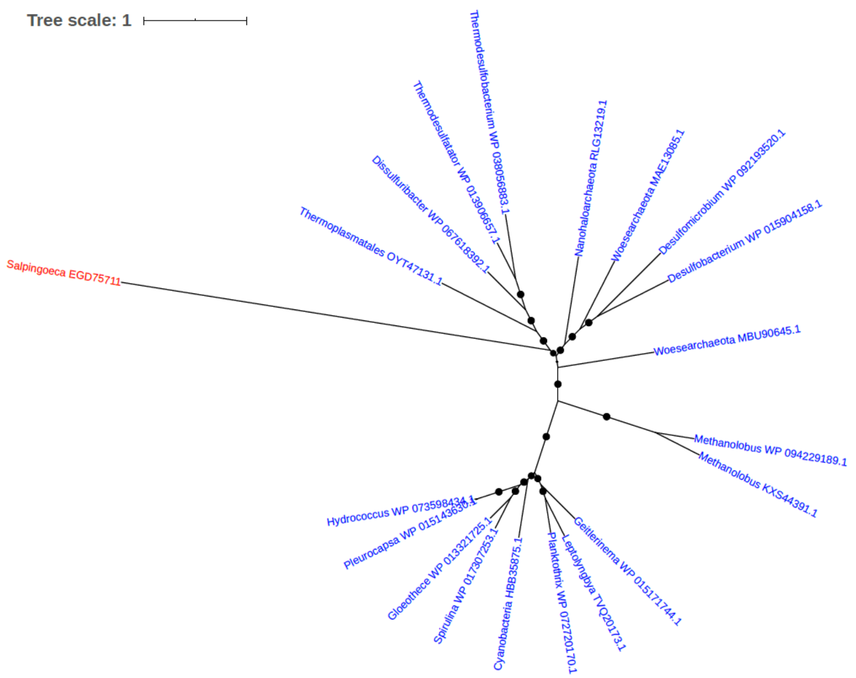


Figure S174. Phylogeny of Pyruvate water dikinase (EGD75711)

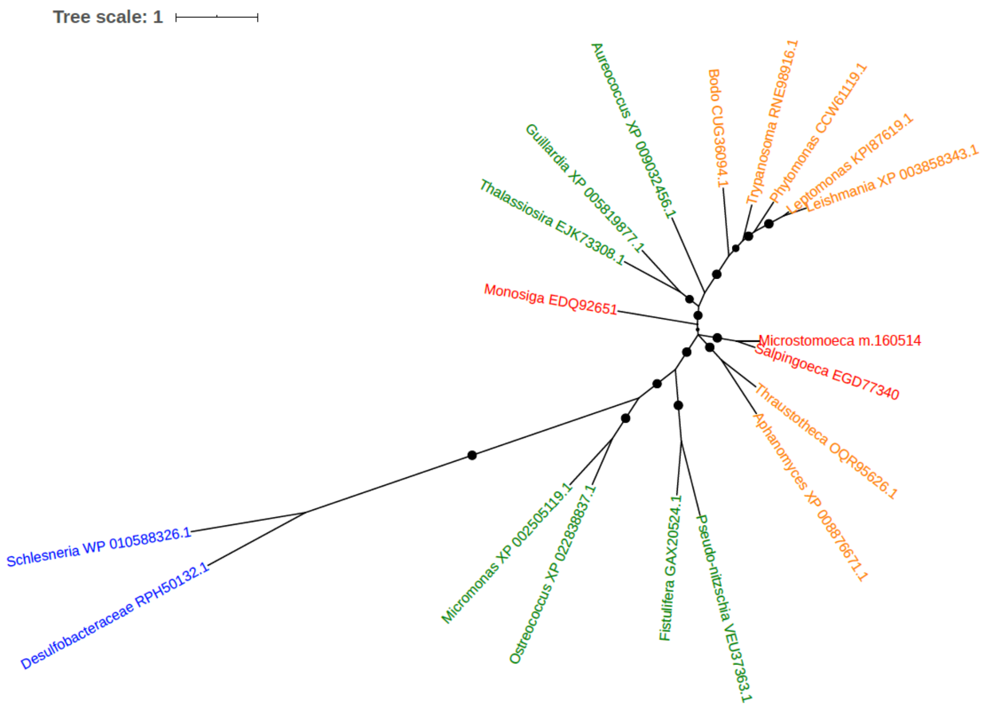


Figure S175. Phylogeny of Reductase with NAD or NADP as acceptor (EGD77304)

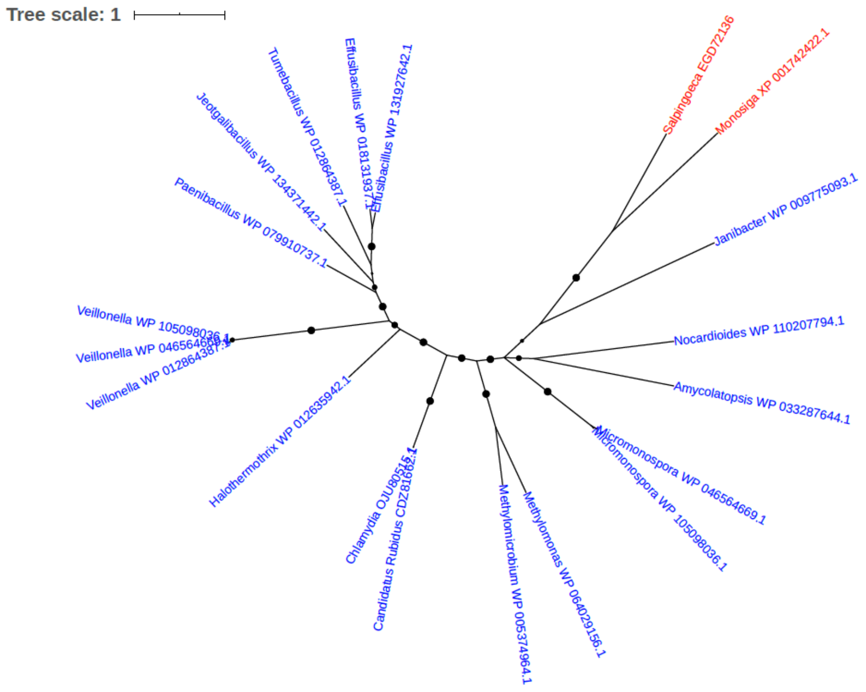


Figure S176. Phylogeny of Riboflavin biosynthesis protein RibA (EGD72136)

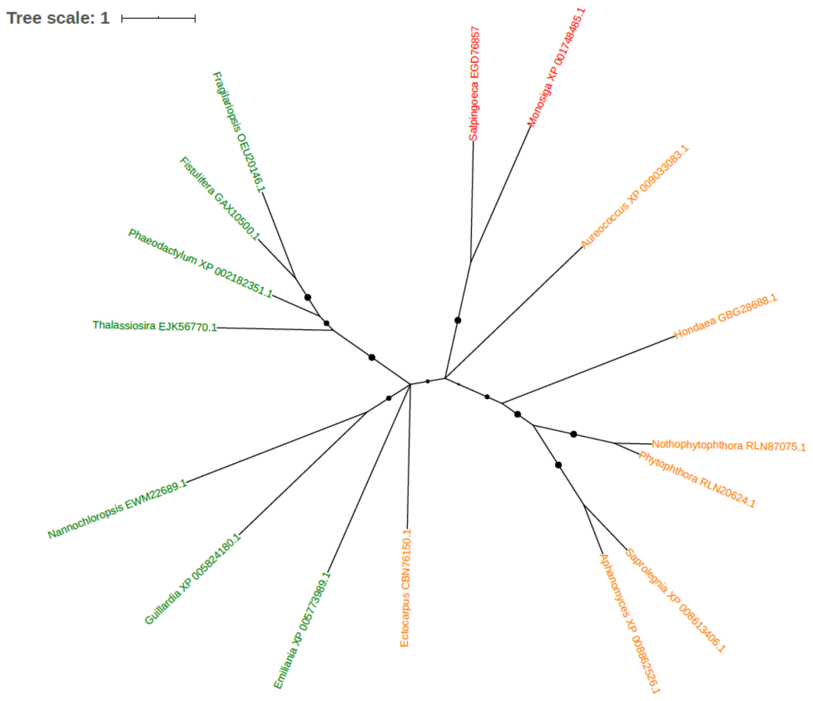


Figure S177. Phylogeny of Riboflavin synthase (EGD76857)

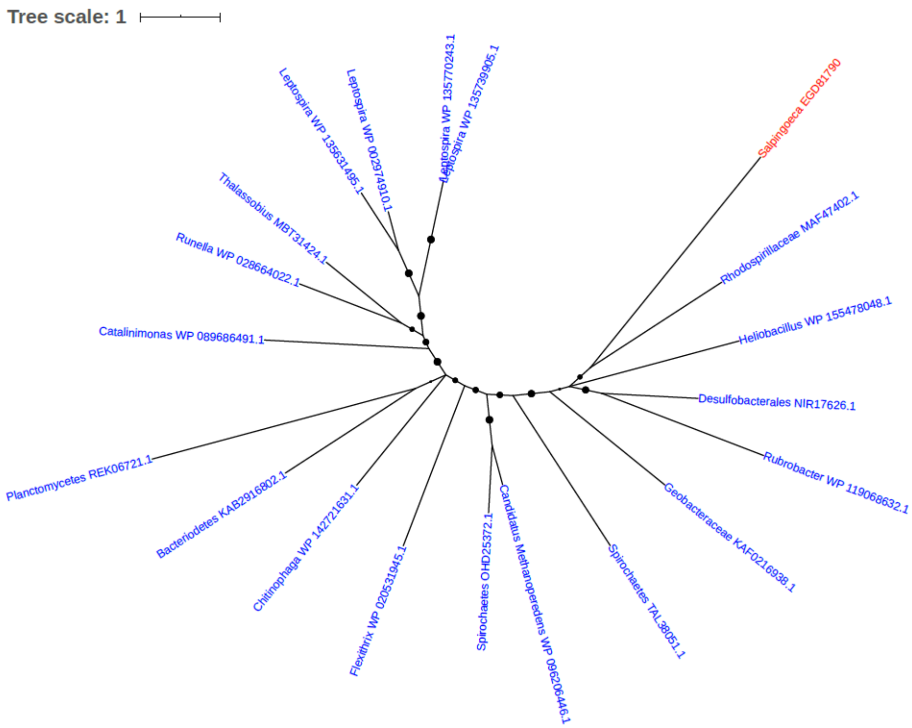


Figure S178. Phylogeny of SAM binding domain-containing protein (EGD81790)

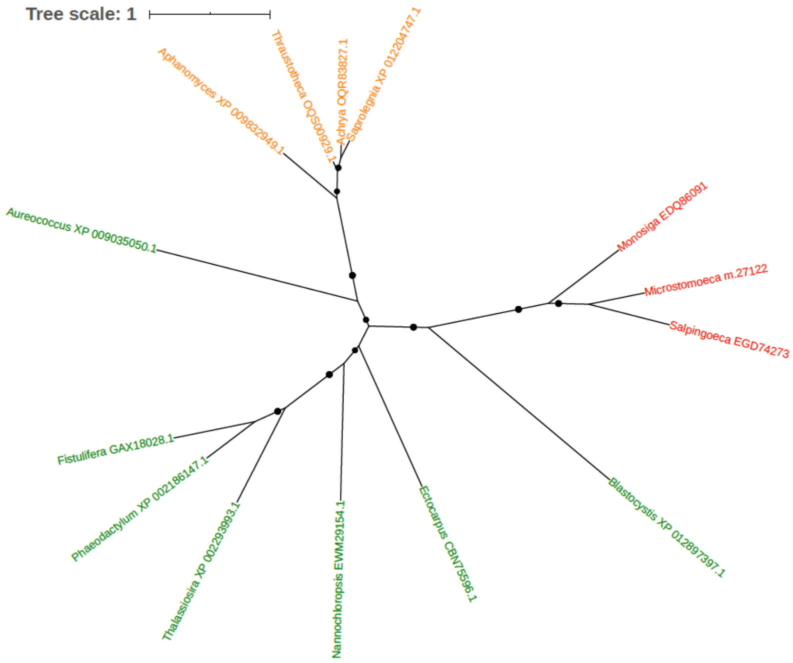


Figure S179. Phylogeny of Serine hydroxymethyltransferase (EC 2.1.2.1) (EGD74273)

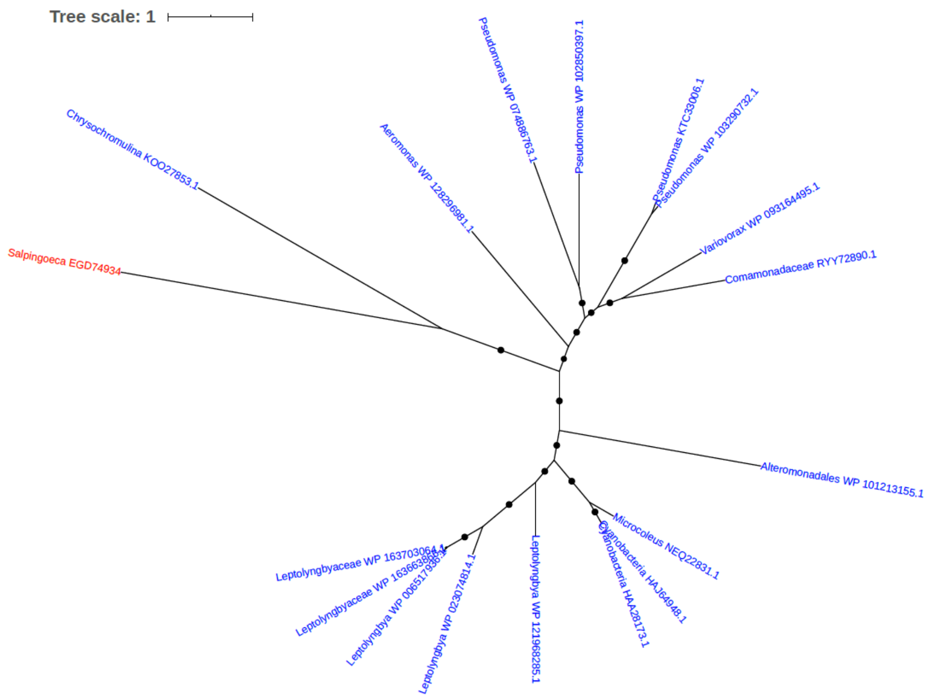


Figure S180. Phylogeny of Serine/threonine protein kinase (EGD74934)

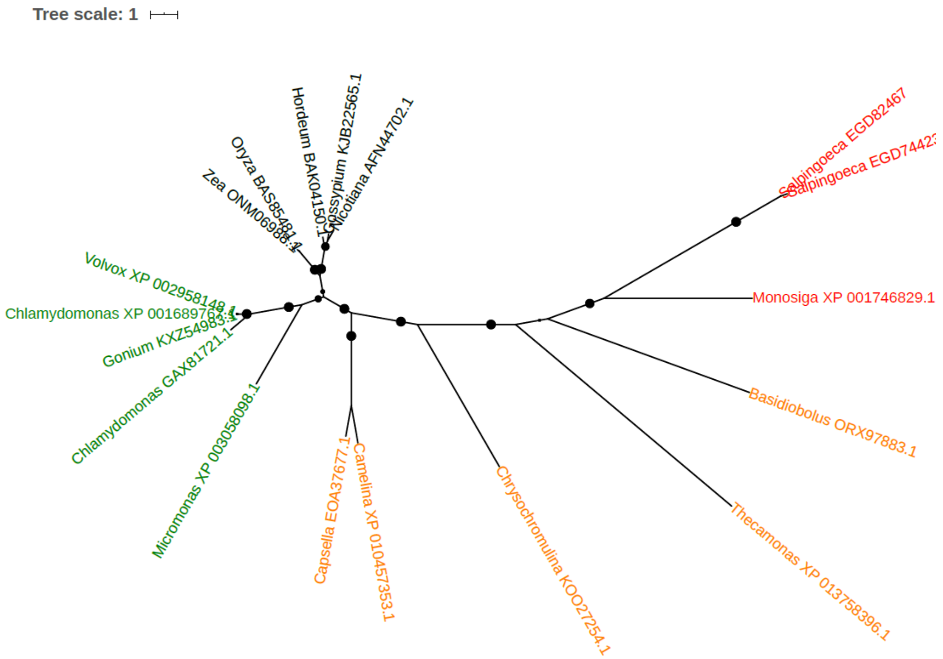


Figure S181. Phylogeny of Serine/threonine-protein phosphatase (EC 3.1.3.16) (EGD82467, EGD74423)

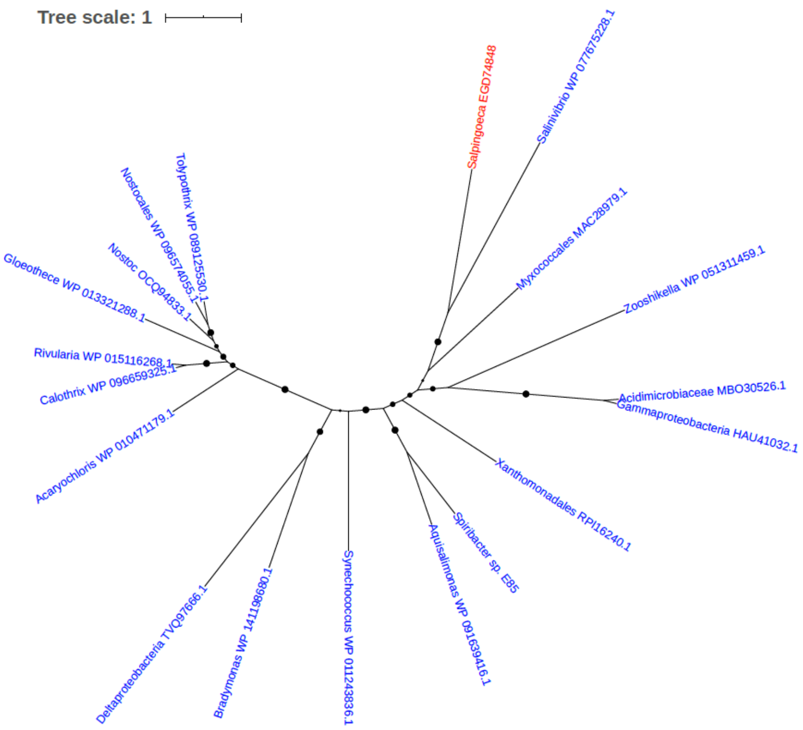


Figure S182. Phylogeny of Short chain dehydrogenase (EGD74848)

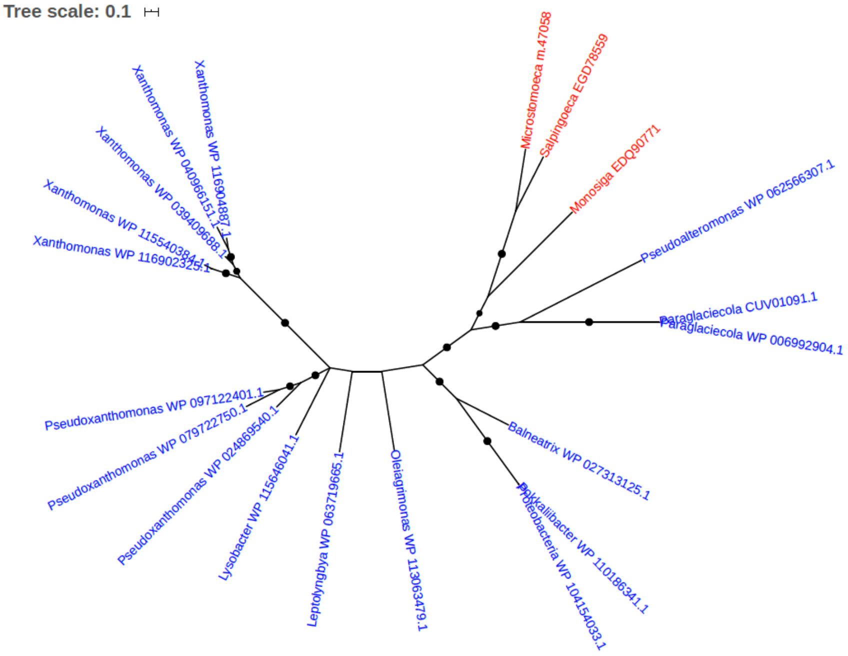


Figure S183. Phylogeny of Short chain dehydrogenase (EGD78559)

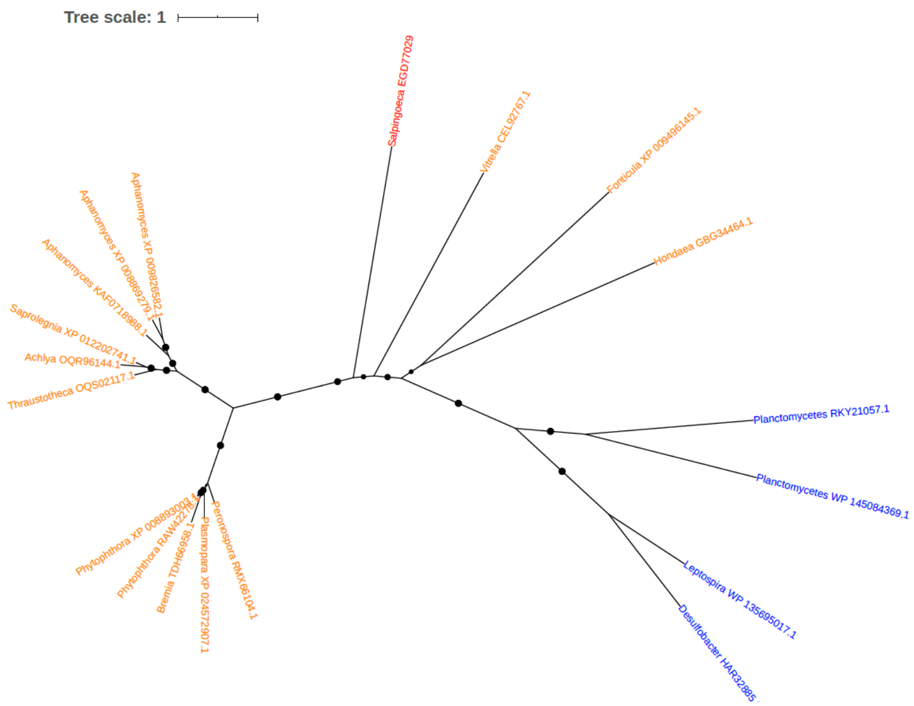


Figure S184. Phylogeny of Short chain dehydrogenase (EGD78559)

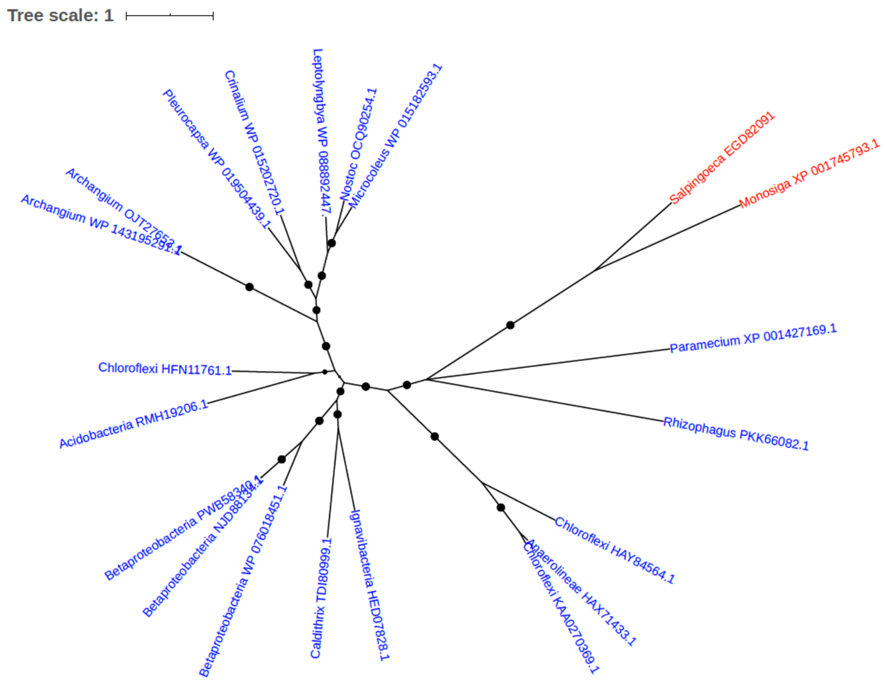


Figure S185. Phylogeny of Short-chain dehydrogenase/reductase 10c (EGD82091)

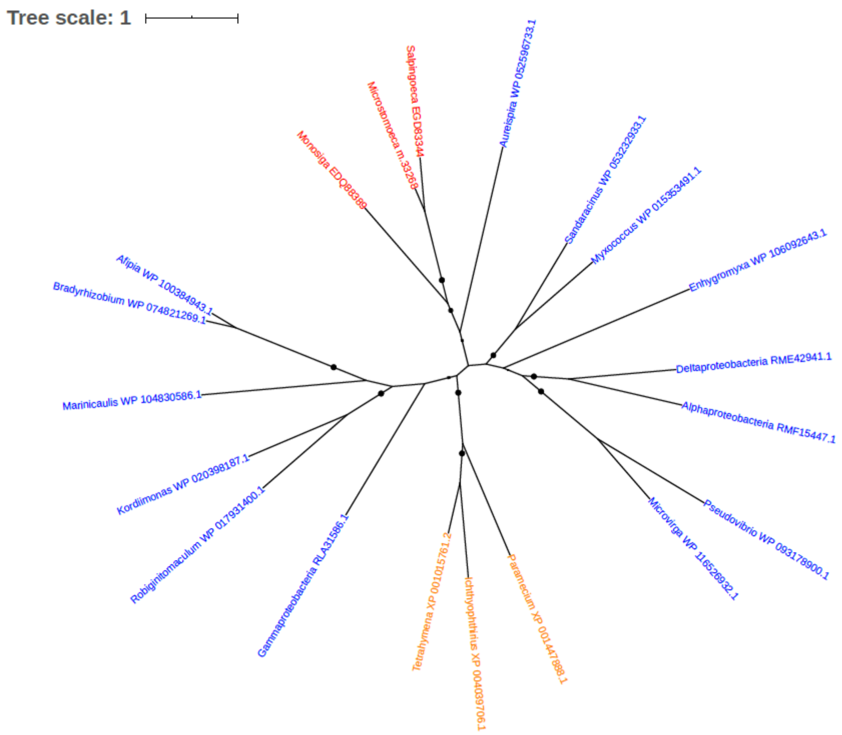


Figure S186. Phylogeny of Short-chain dehydrogenase/reductase SDR (EGD83344)

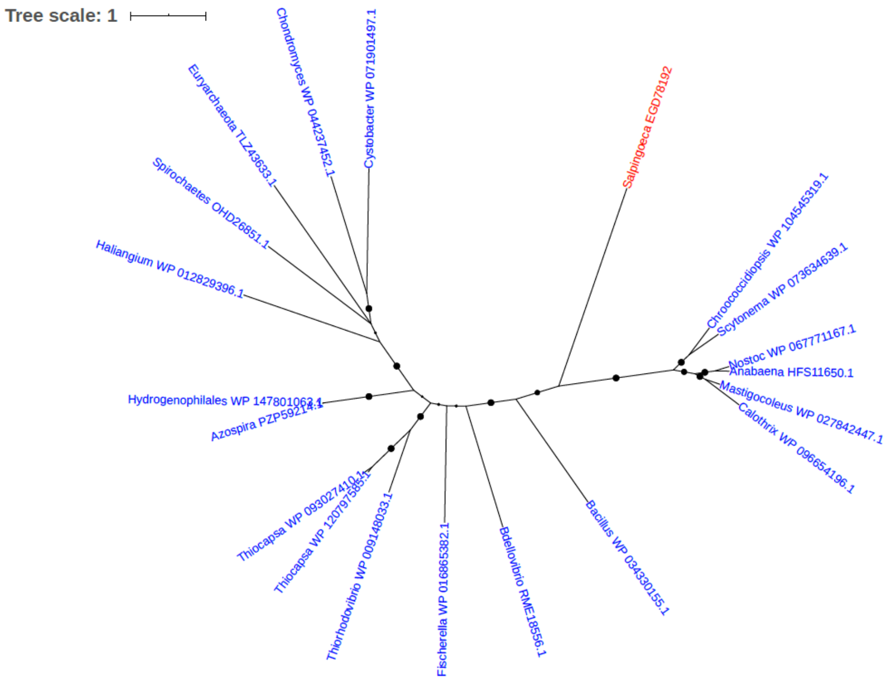


Figure S187. Phylogeny of Short-chain dehydrogenase/reductase SDR (EGD78192)

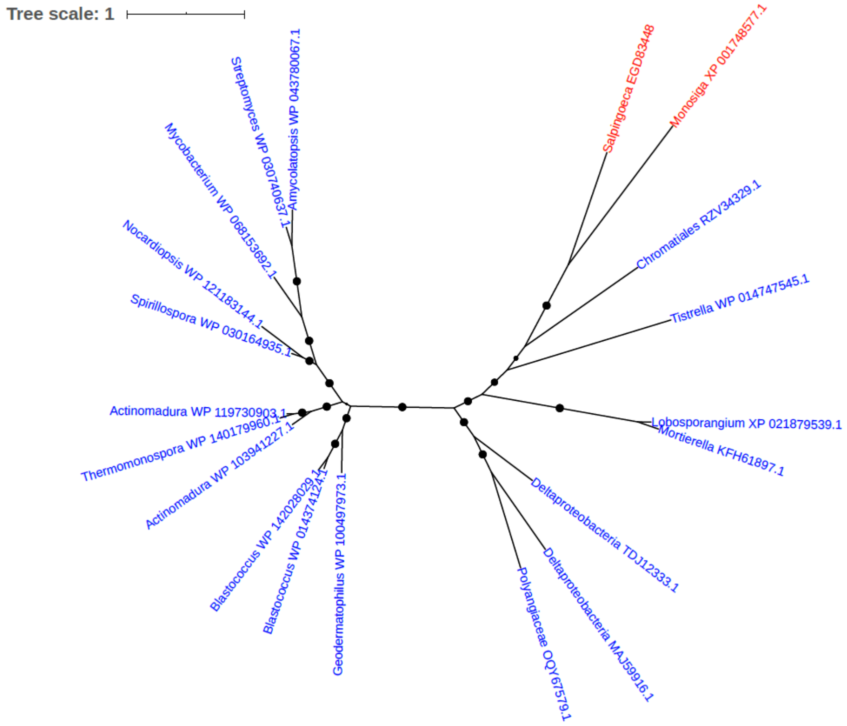


Figure S188. Phylogeny of Short-chain dehydrogenase/reductase SDR (EGD83448)

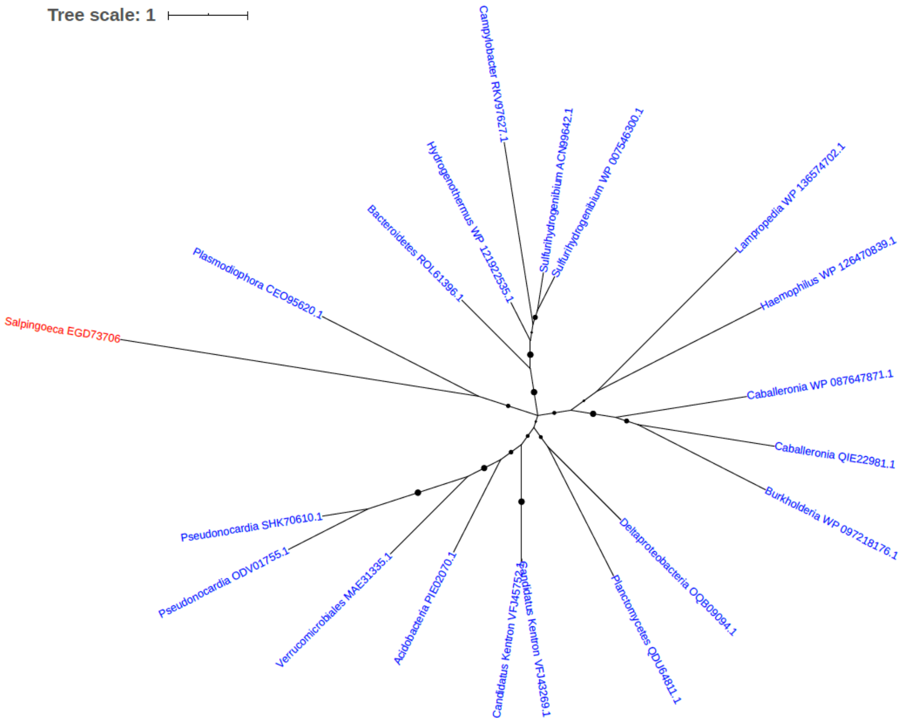


Figure S189. Phylogeny of Sir2 family transcriptional regulator (EGD73706)

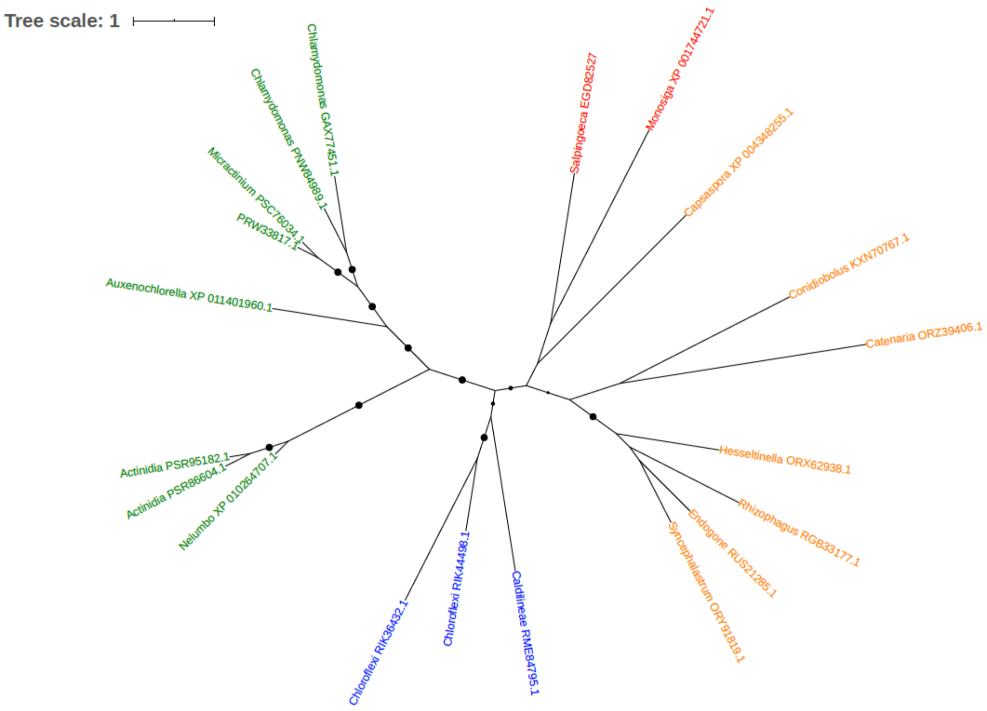


Figure S190. Phylogeny of SNO glutamine amidotransferase (Fragment) (EGD82527)

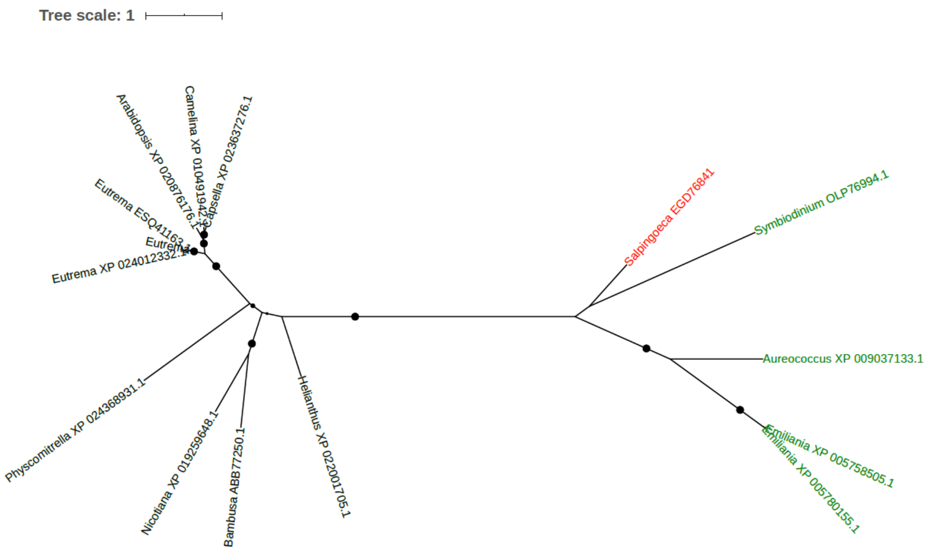


Figure S191. Phylogeny of Soluble acid invertase (EGD76841)

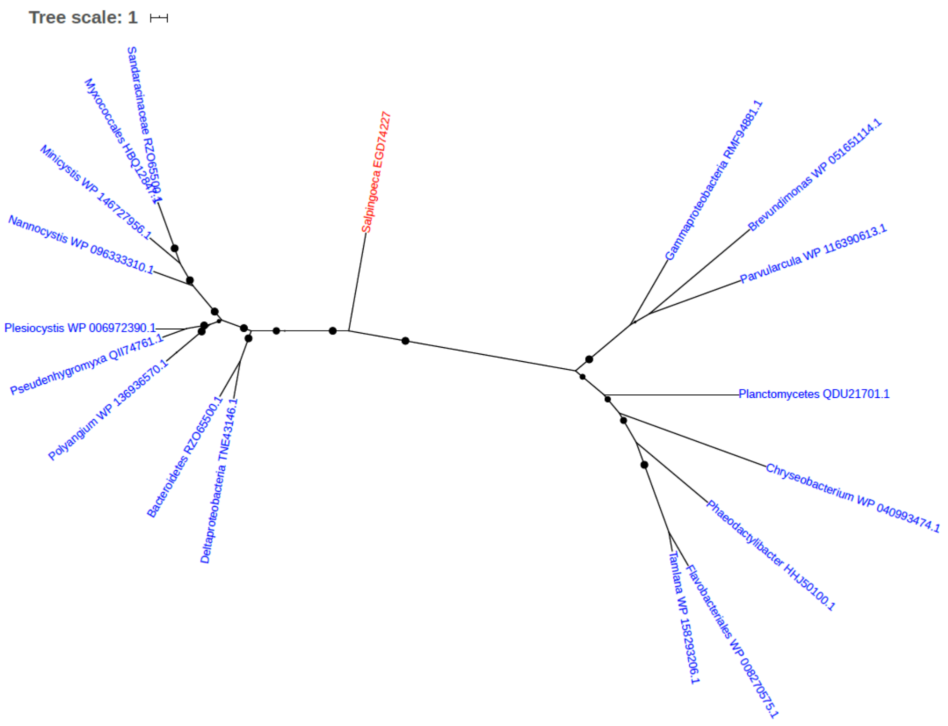


Figure S192. Phylogeny of Sterol desaturase (EGD74227)

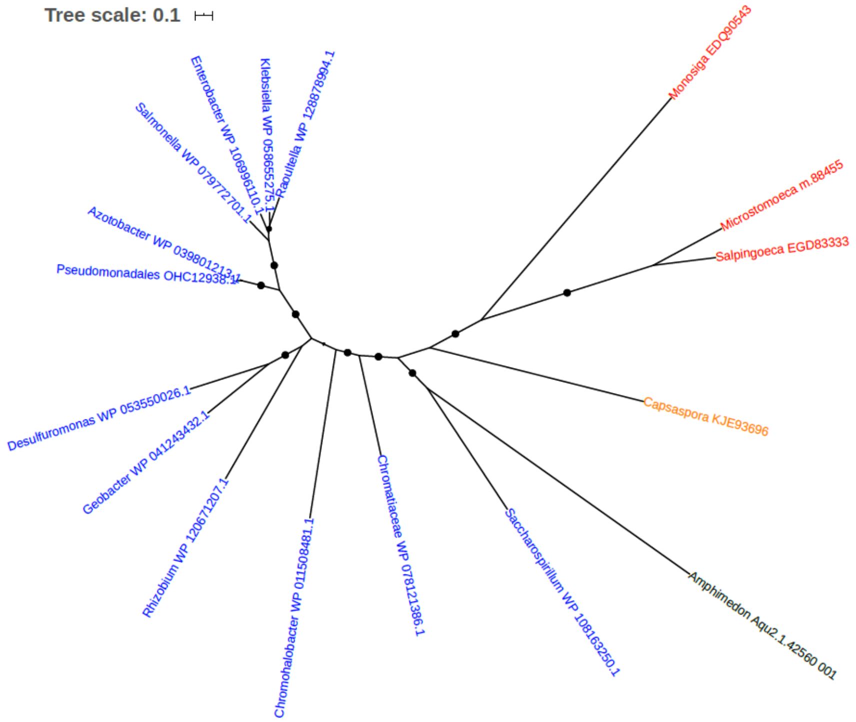


Figure S193. Phylogeny of Succinate-semialdehyde dehydrogenase (EC 1.2.1.24) (EGD83333)

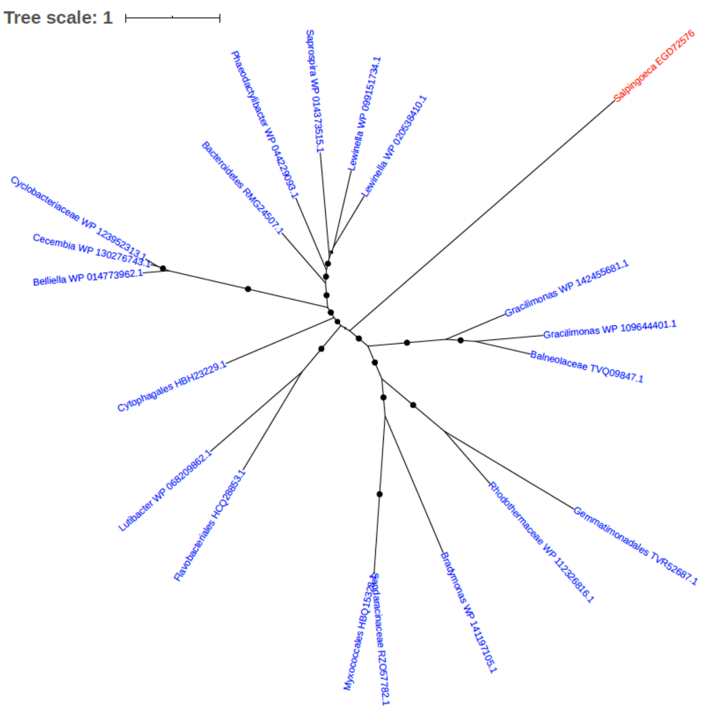


Figure S194. Phylogeny of Sulfate transporter (EGD72576)

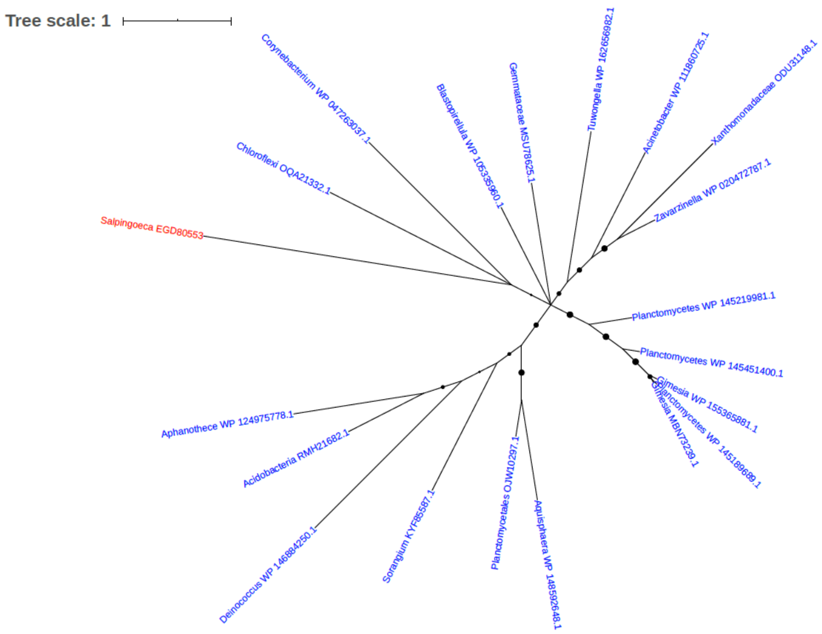


Figure S195. Phylogeny of Swarming motility protein YbiA (EGD80553)

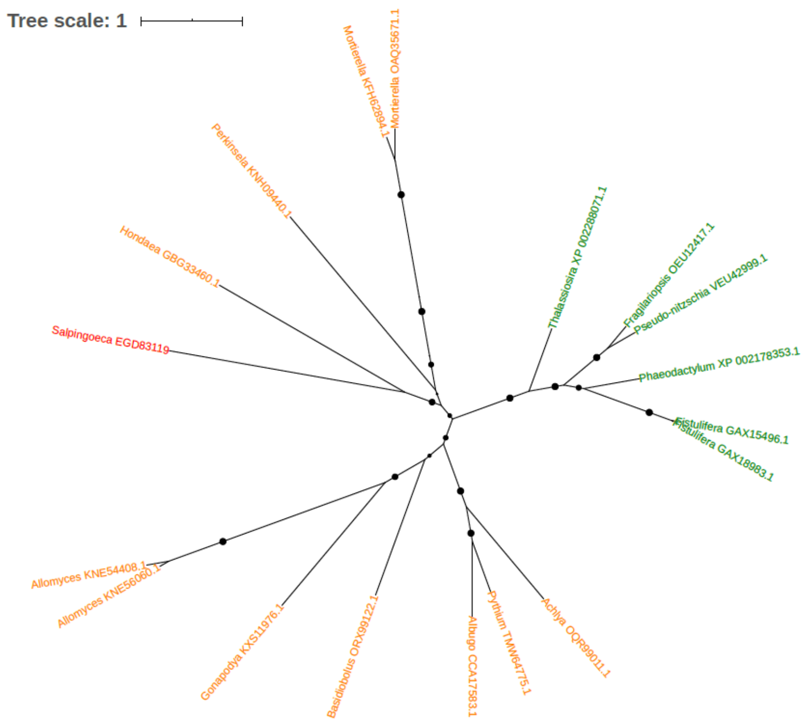


Figure S196. Phylogeny of T-cell receptor beta chain ANA 11 (EGD83119)

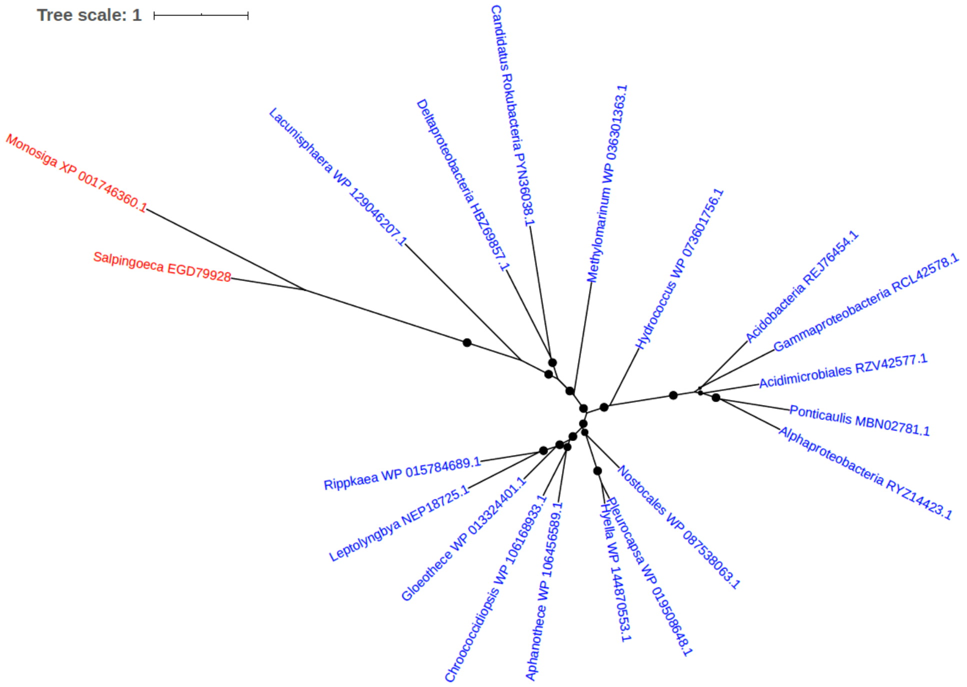


Figure S197. Phylogeny of Tetratricopeptide repeat domain-containing protein (EGD79928)

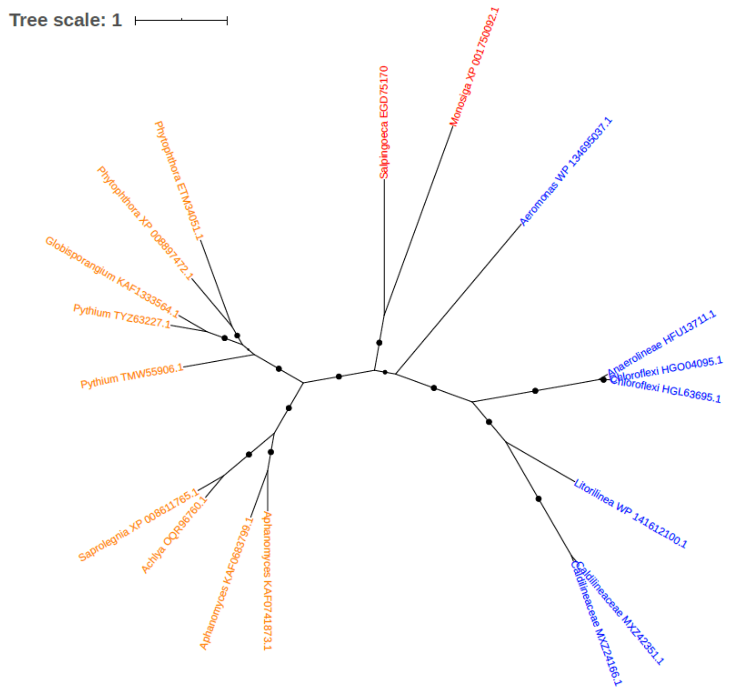


Figure S198. Phylogeny of Threonine aldolase (EGD75170)

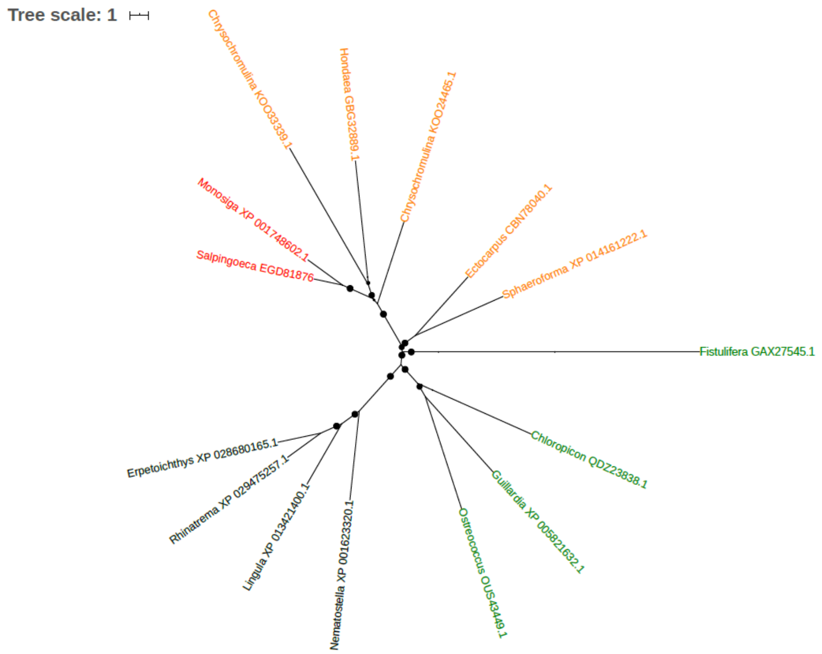


Figure S199. Phylogeny of Thymus specific serine peptidase (EGD81876)

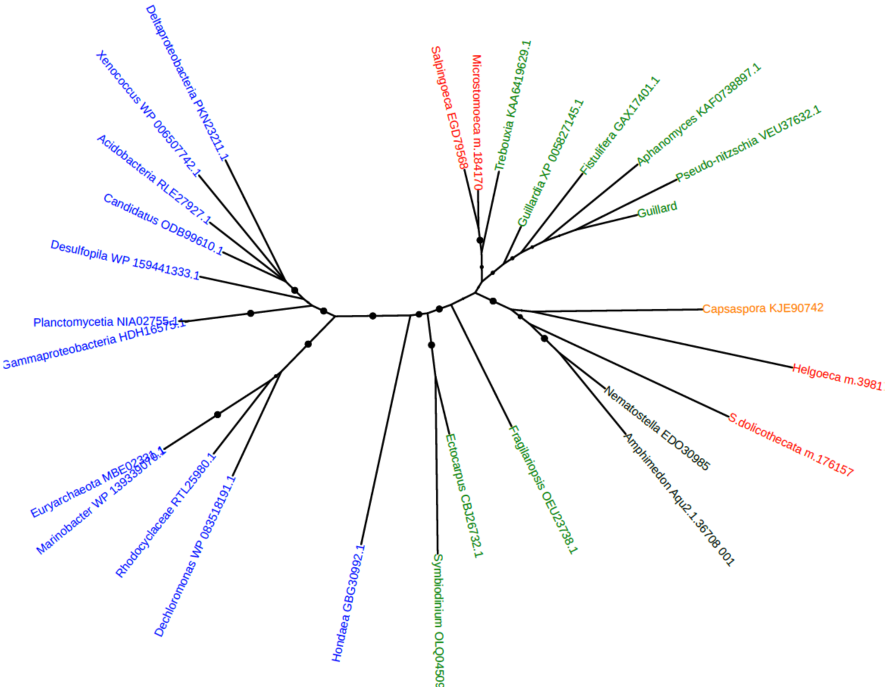


Figure S200. Phylogeny of TKL protein kinase (EGD79568)

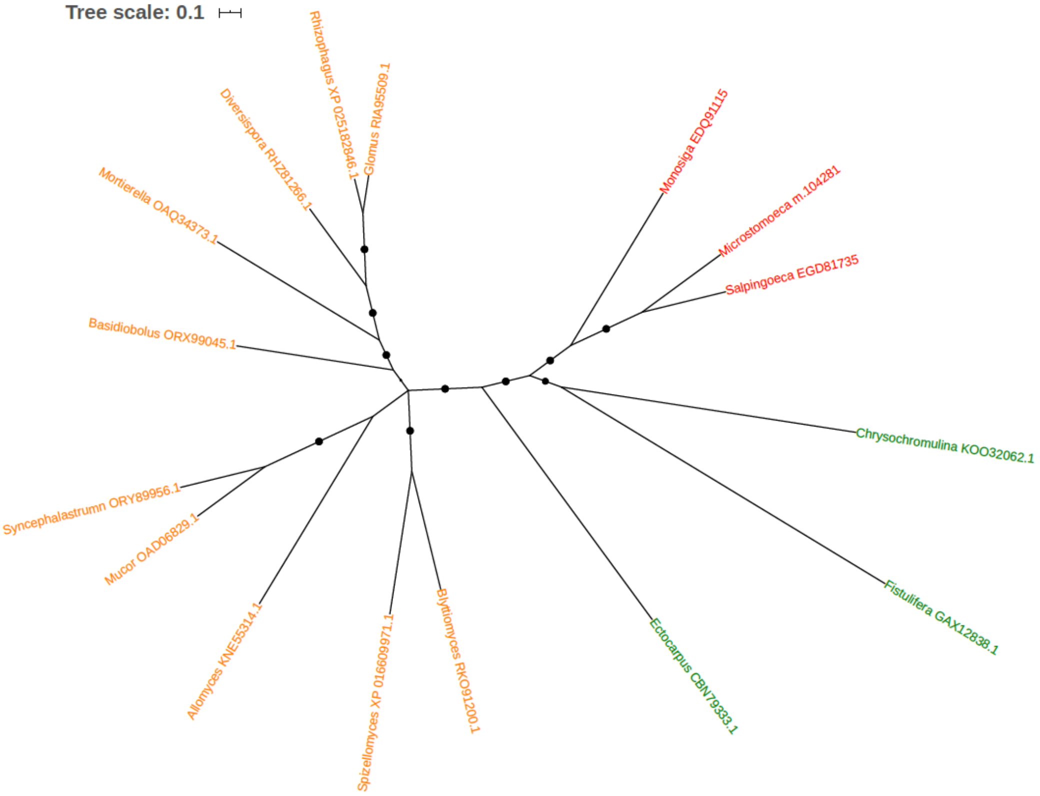


Figure S201. Phylogeny of Transketolase (EC 2.2.1.1) (EGD81735)

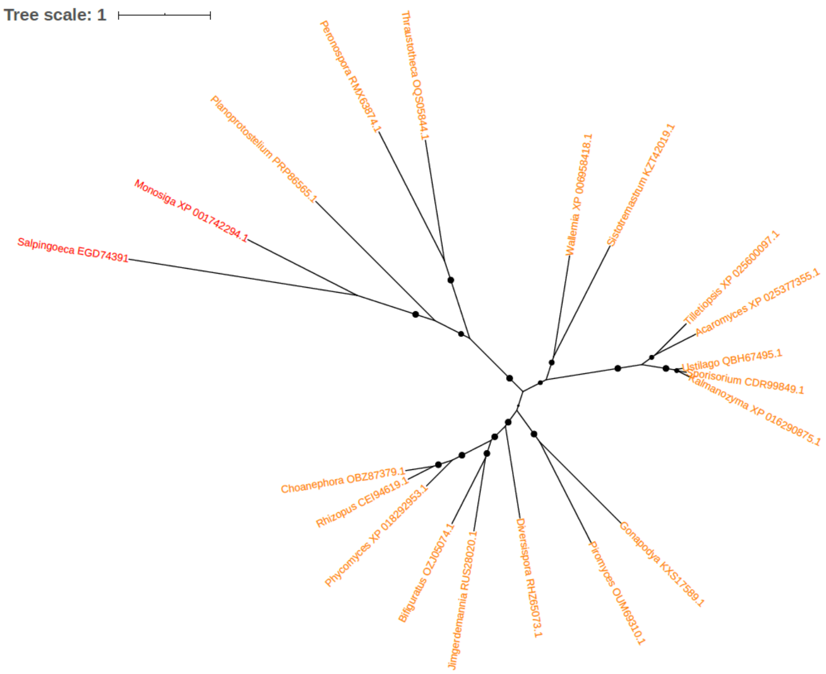


Figure S202. Phylogeny of Trifunctional protein with glutamine amidotransferase (EGD74391)

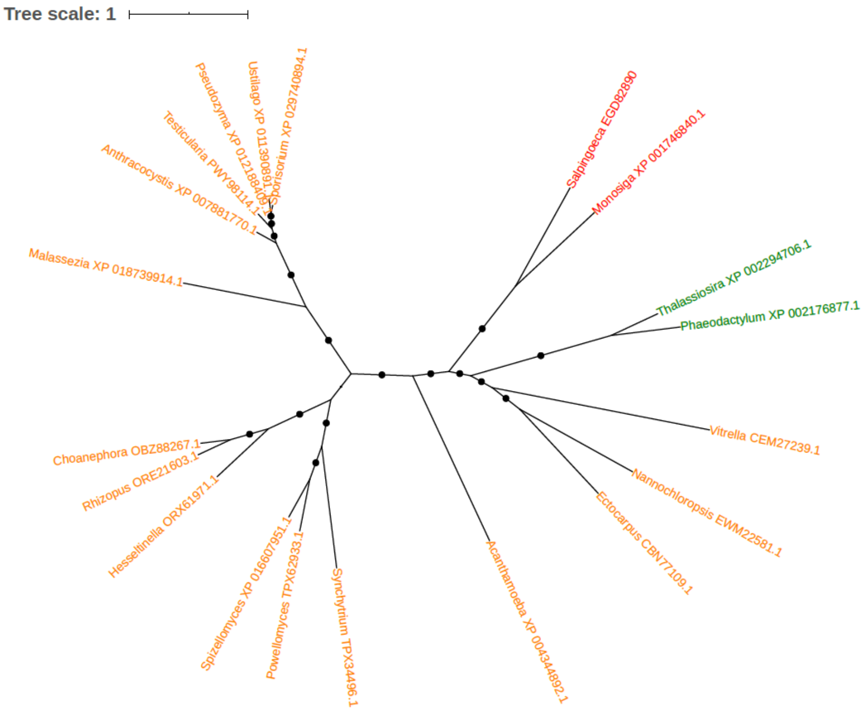


Figure S203. Phylogeny of TrpB (EGD82890)

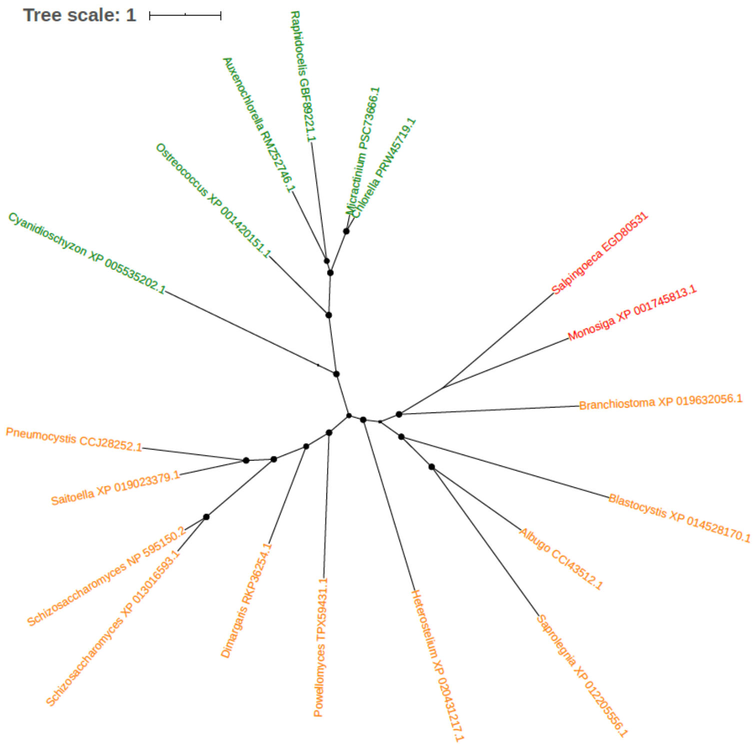


Figure S204. Phylogeny of TTK protein kinase (EGD80531)

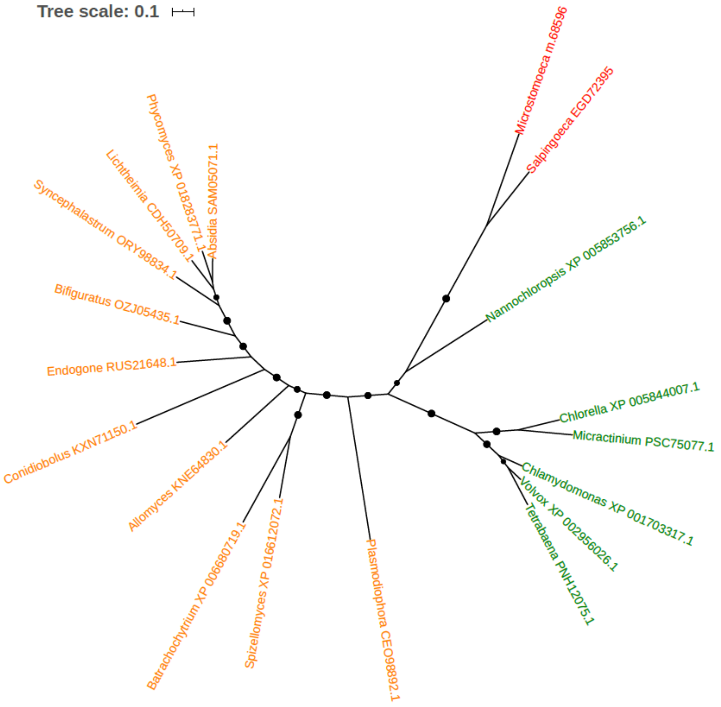


Figure S205. Phylogeny of U3 small nucleolar ribonucleoprotein IMP3 (EGD72395)

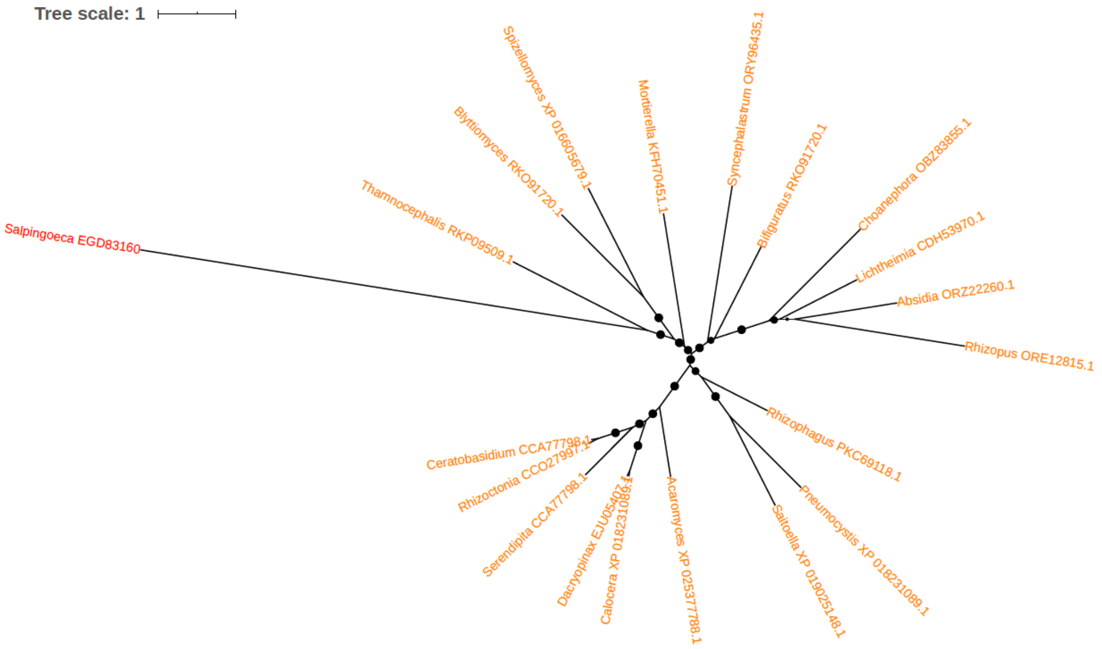


Figure S206. Phylogeny of Ubiquitin carboxyl-terminal hydrolase (EGD83160)

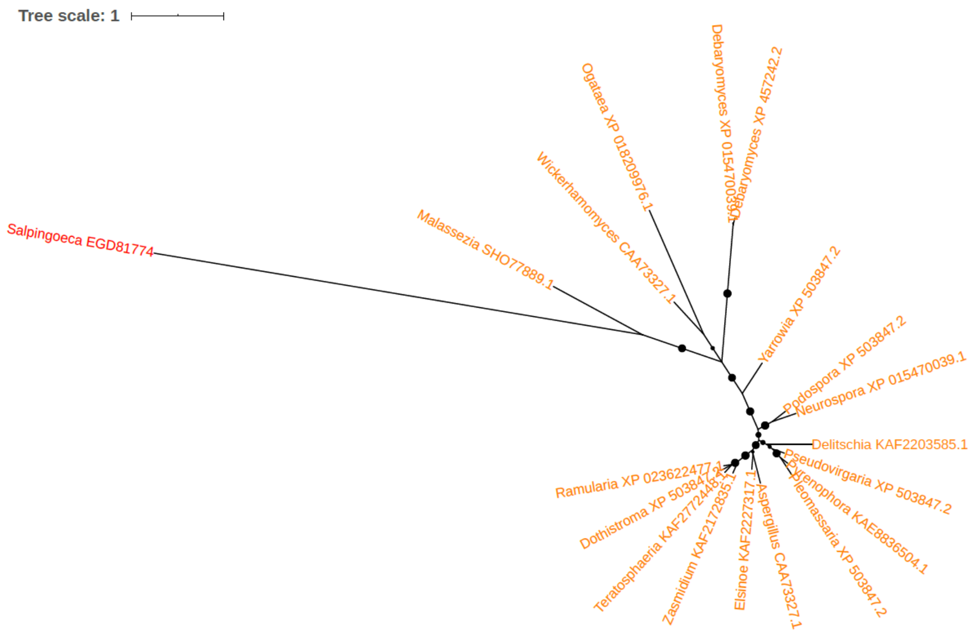


Figure S207. Phylogeny of Ubiquitin carrier protein (EGD81774)

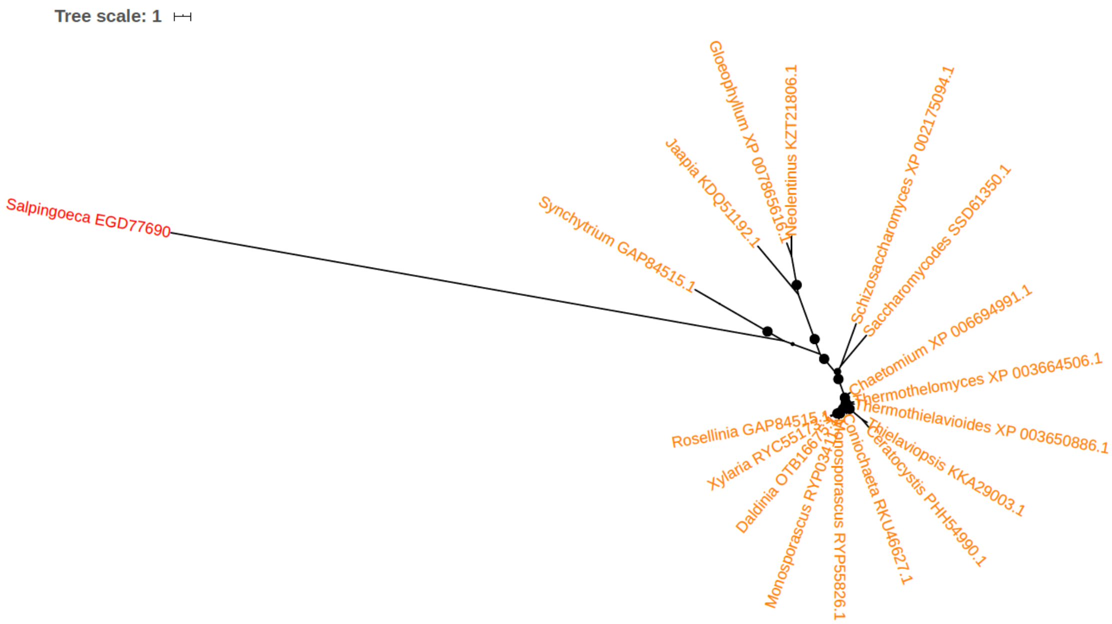


Figure S208. Phylogeny of Ubiquitin protein ligase (EGD77690)

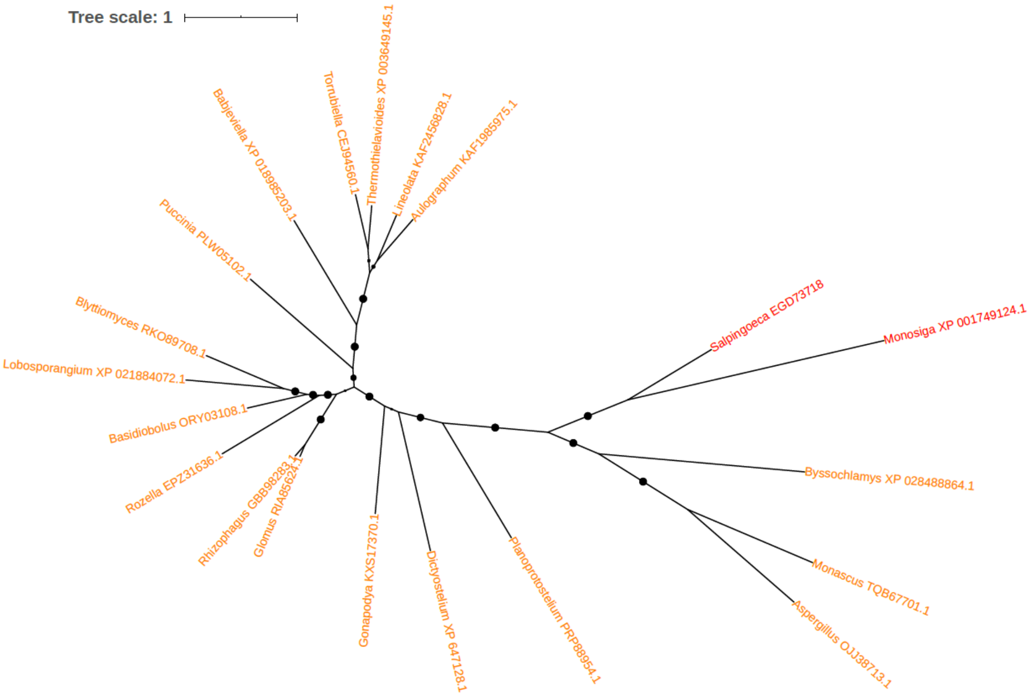


Figure S209. Phylogeny of Uracil phosphoribosyltransferase (EGD73718)

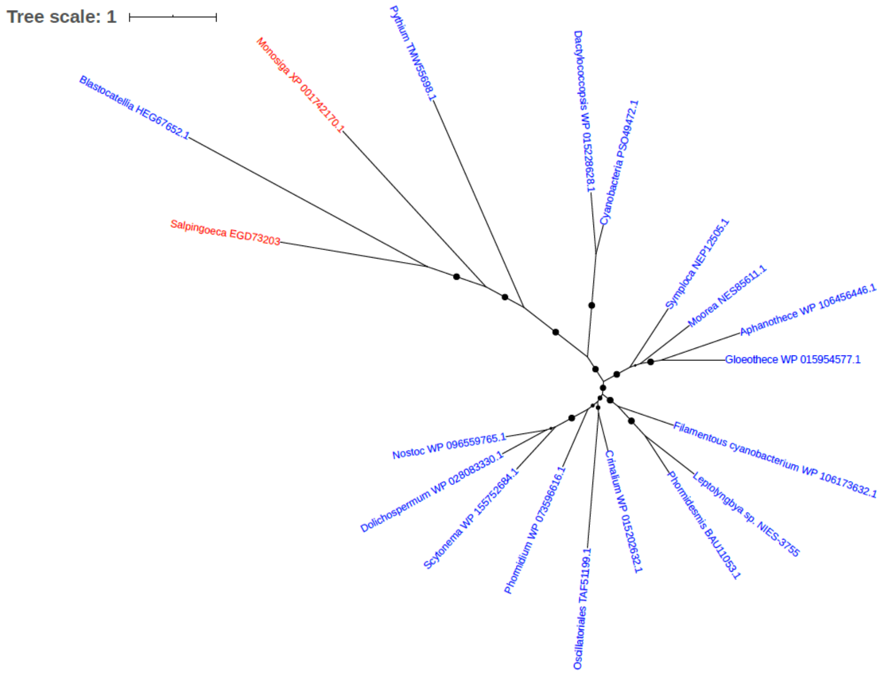


Figure S210. Phylogeny of Uroporphyrinogen-III C-methyltransferase (EGD73203)

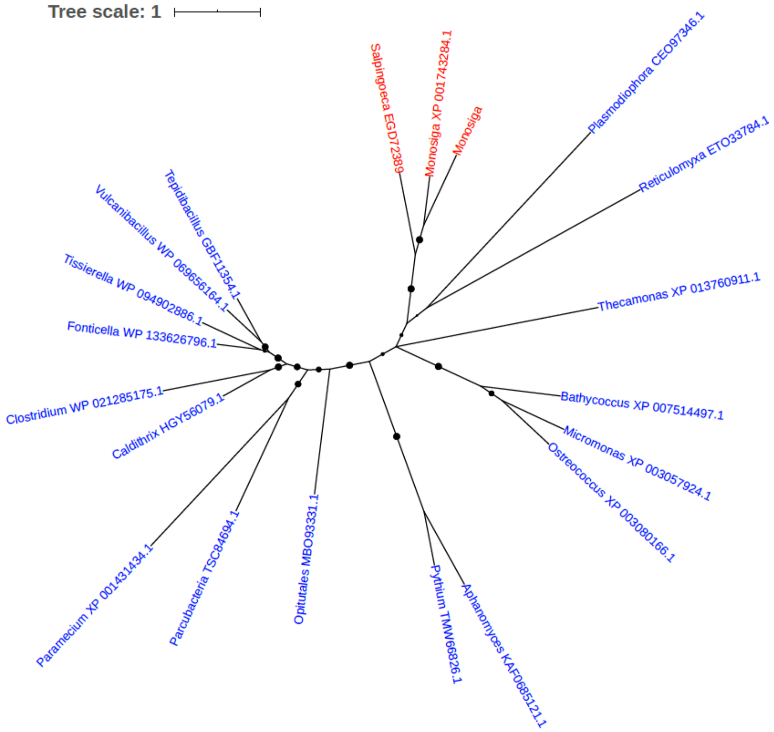


Figure S211. Phylogeny of Vacuolar iron family transporter (EGD72389)

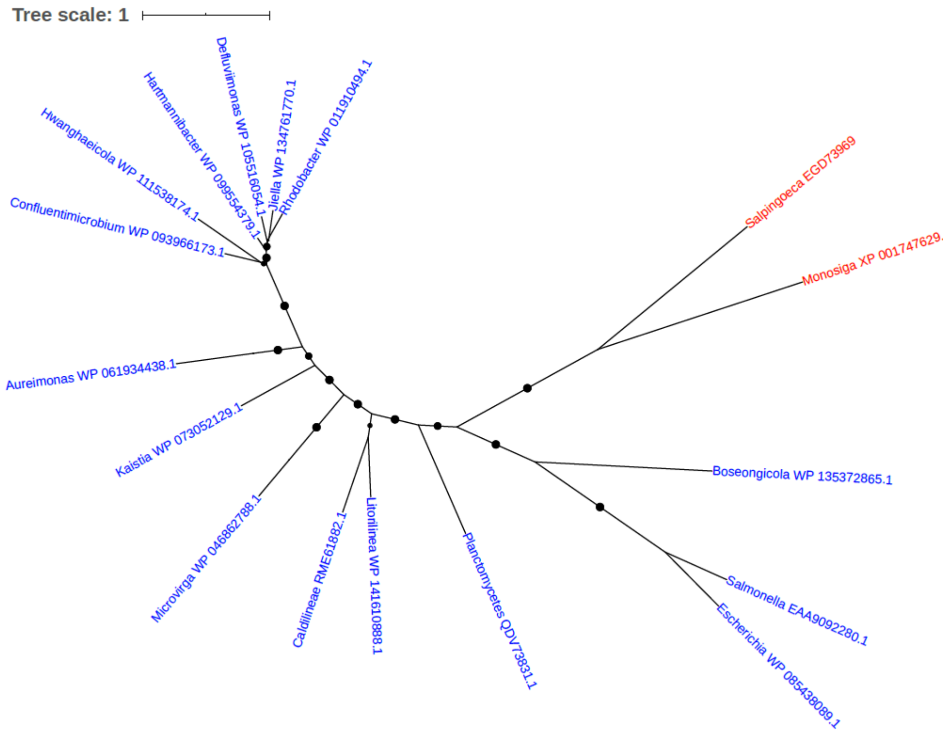


Figure S212. Phylogeny of Xylitol dehydrogenase (EGD73969)

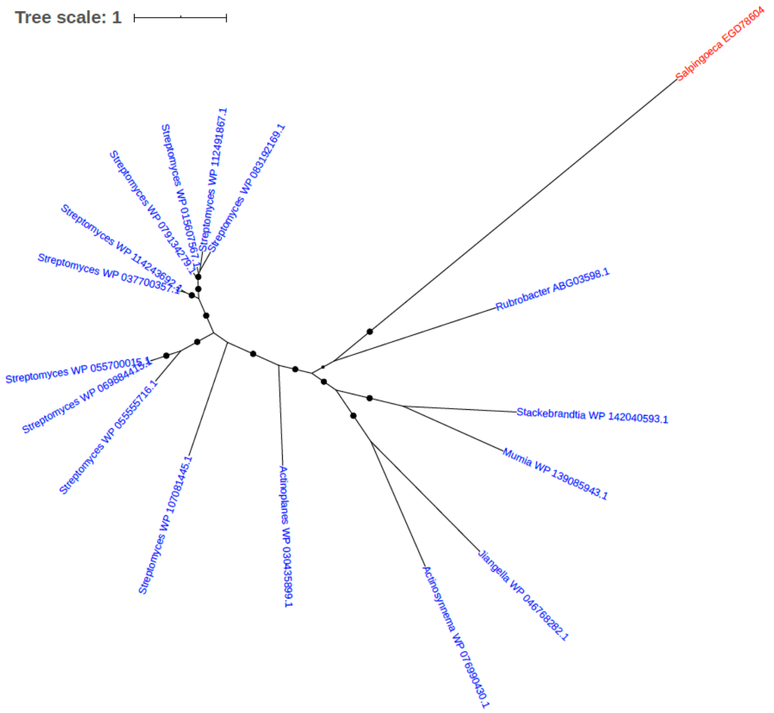


Figure S213. Phylogeny of YbbC (EGD78604)

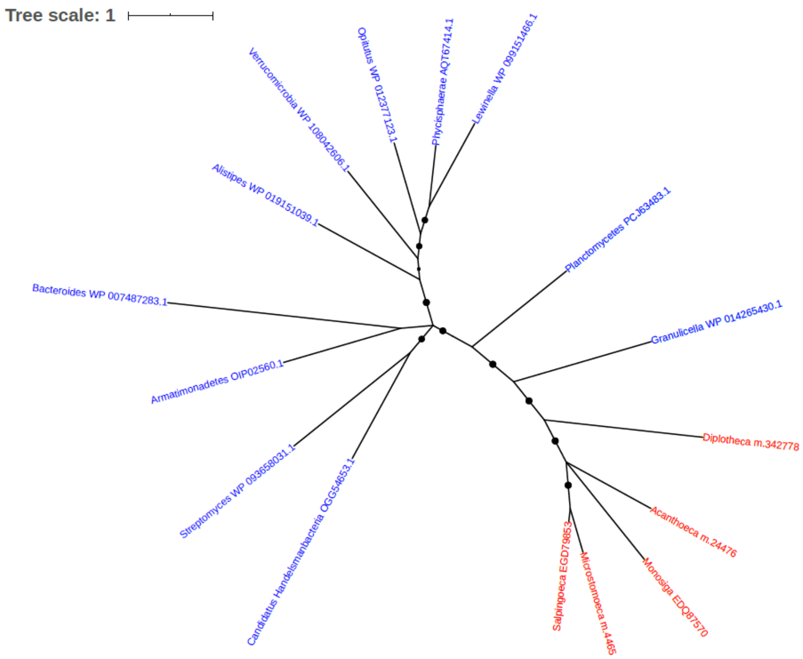


Figure S214. Phylogeny of uncharacterized protein (EGD79853)

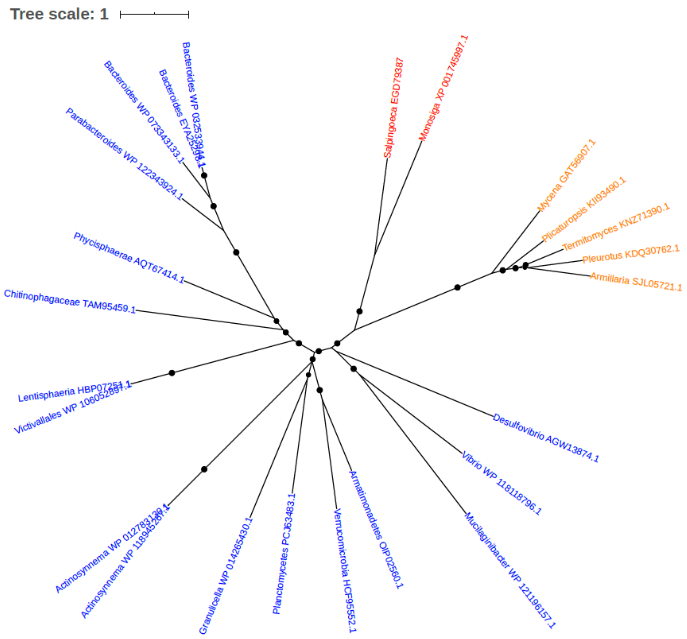


Figure S215. Phylogeny of uncharacterized protein (EGD79387)

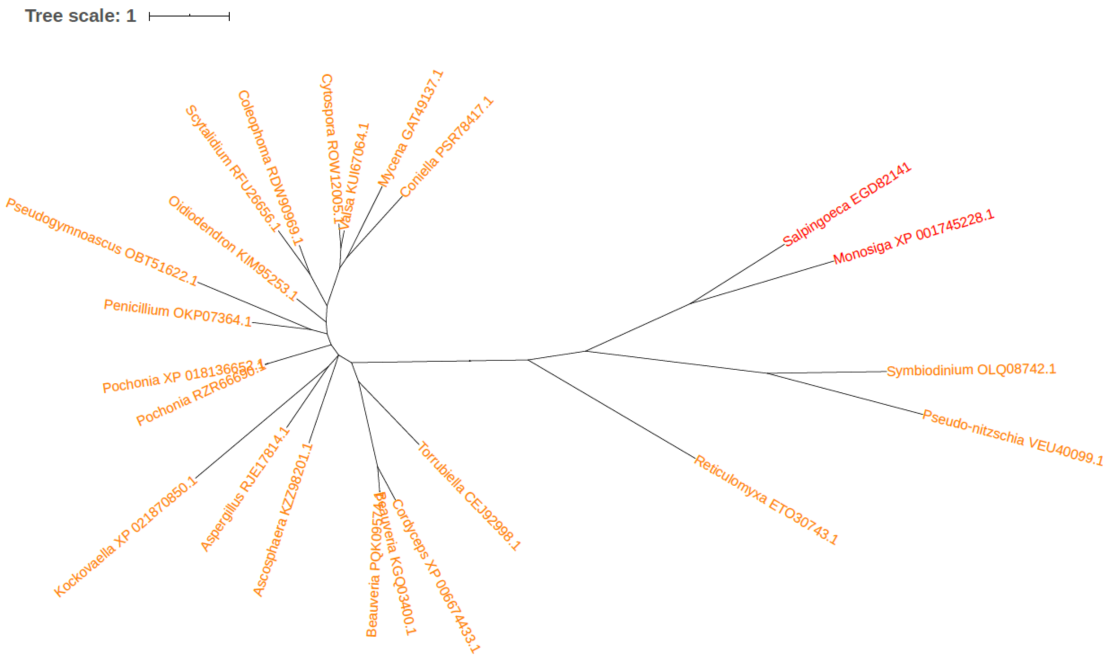


Figure S216. Phylogeny of uncharacterized protein (EGD82141)

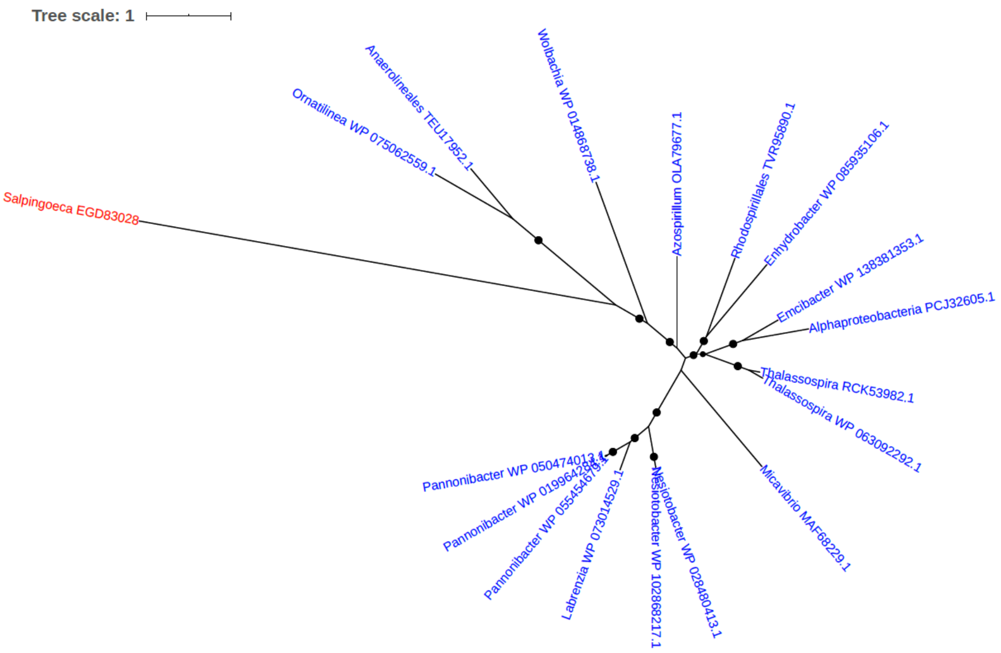


Figure S217. Phylogeny of uncharacterized protein (EGD83028)

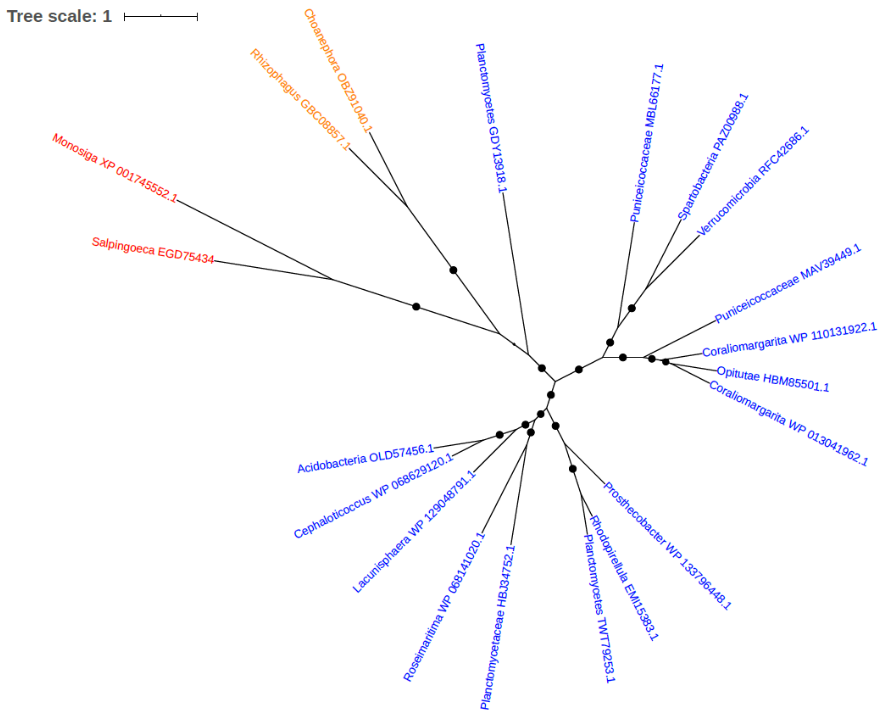


Figure S218. Phylogeny of uncharacterized protein (EGD75434)

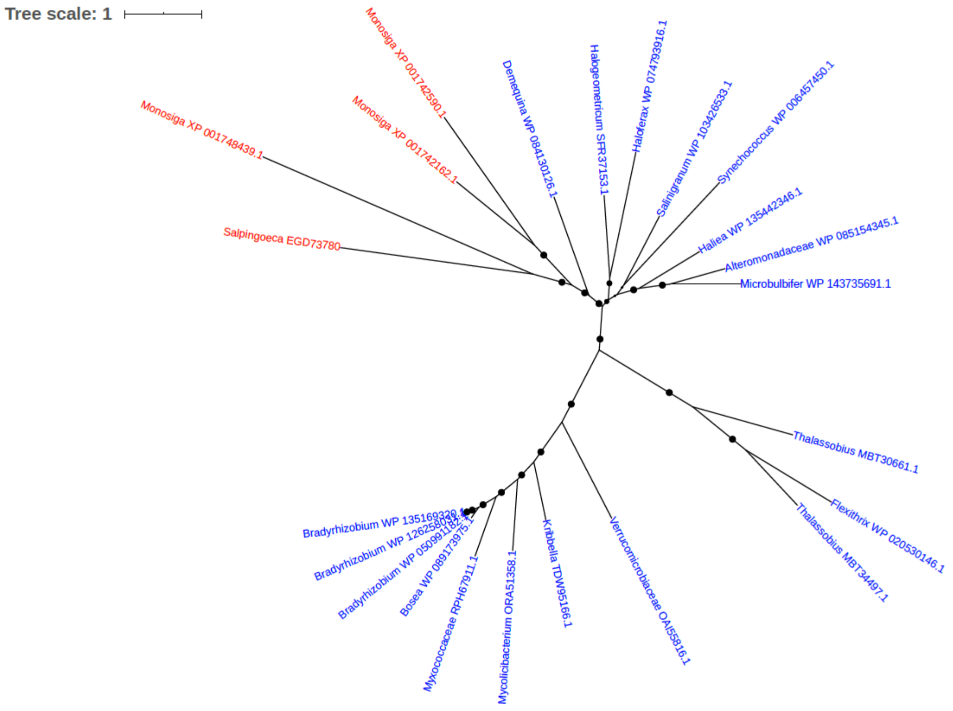


Figure S219. Phylogeny of uncharacterized protein (EGD73780)

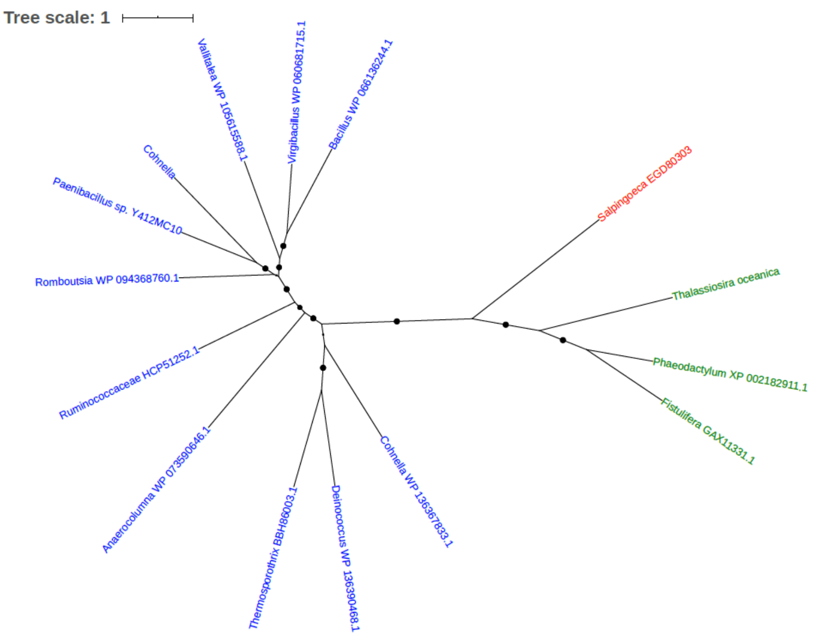


Figure S220. Phylogeny of uncharacterized protein (EGD80303)

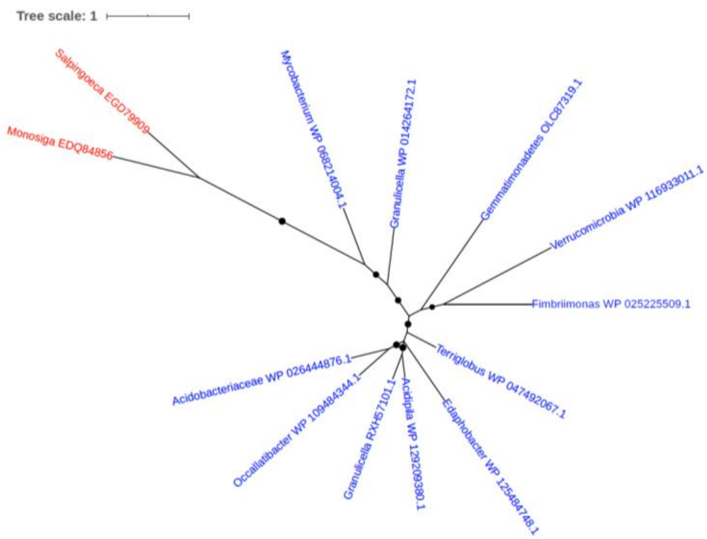


Figure S221. Phylogeny of uncharacterized protein (EGD79909)

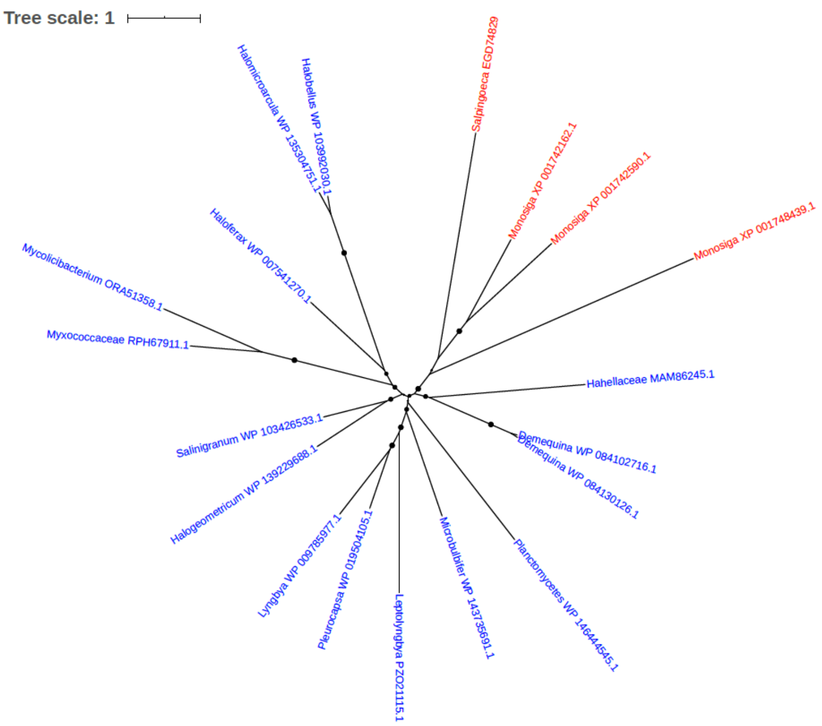


Figure S222. Phylogeny of uncharacterized protein (EGD74829)

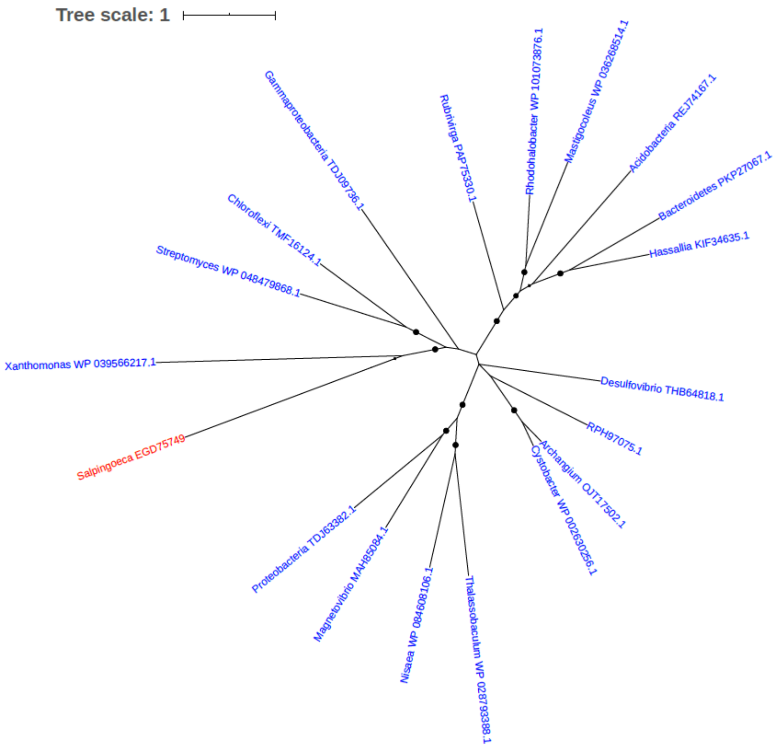


Figure S223. Phylogeny of uncharacterized protein (EGD75749)

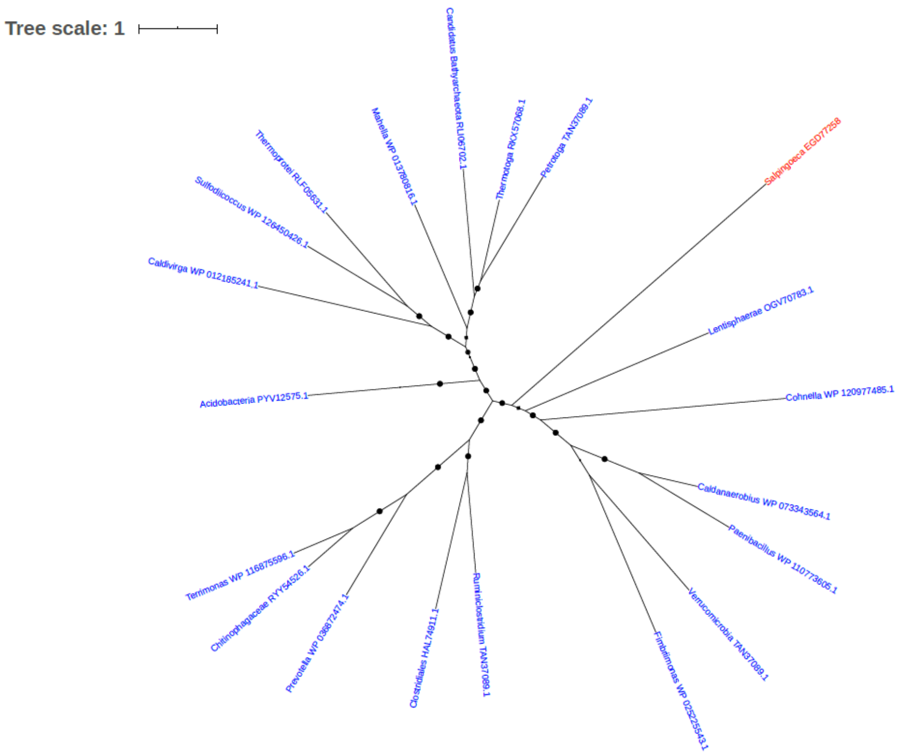


Figure S224. Phylogeny of uncharacterized protein (EGD77258)

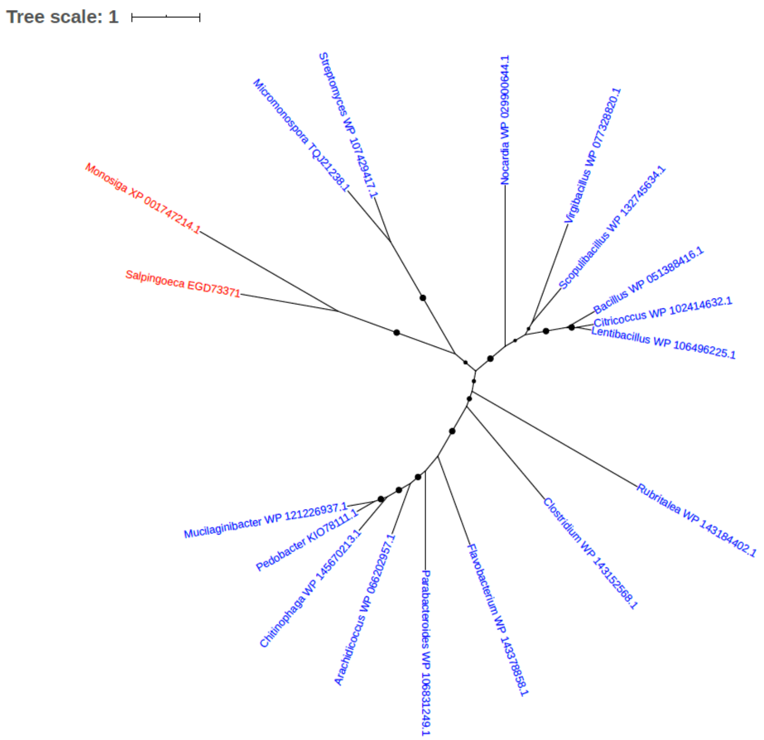


Figure S225. Phylogeny of uncharacterized protein (EGD73371)

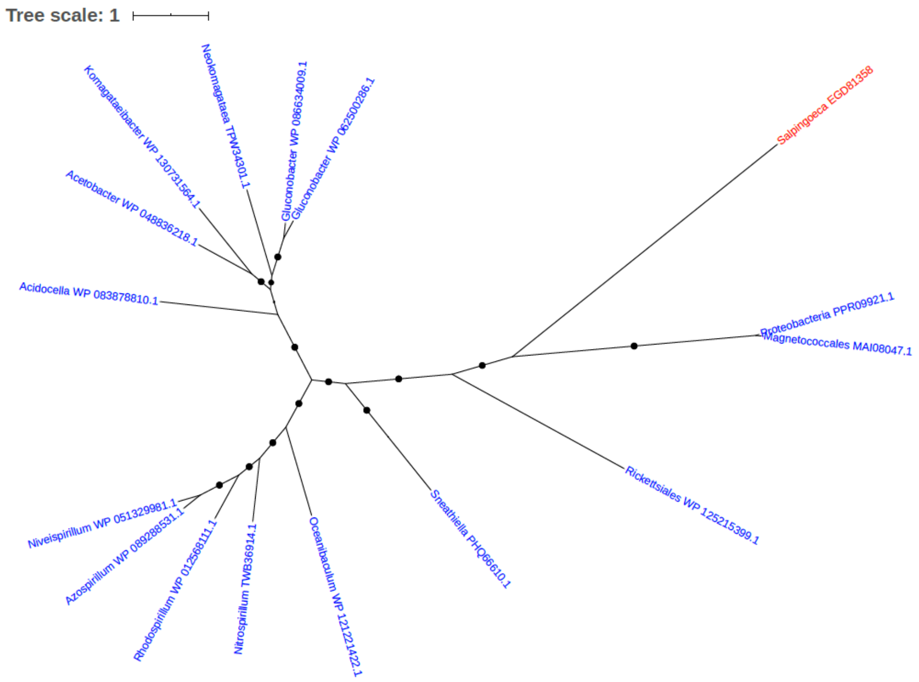


Figure S226. Phylogeny of uncharacterized protein (EGD81358)

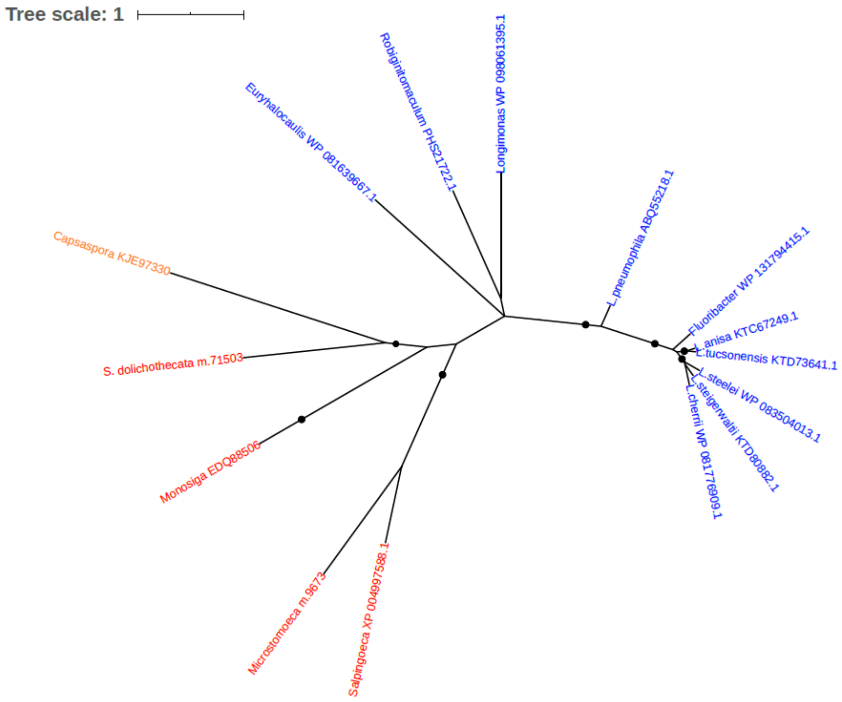


Figure S227. Phylogeny of uncharacterized protein

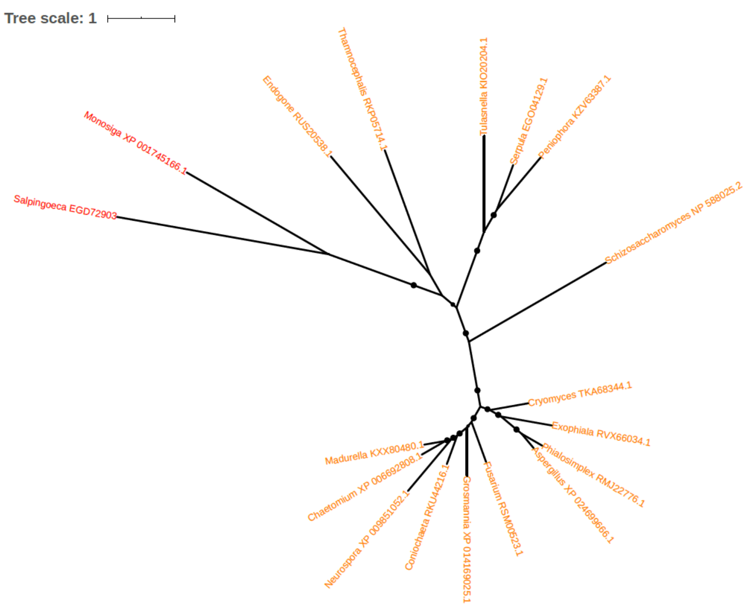


Figure S228. Phylogeny of uncharacterized protein (EGD72903)

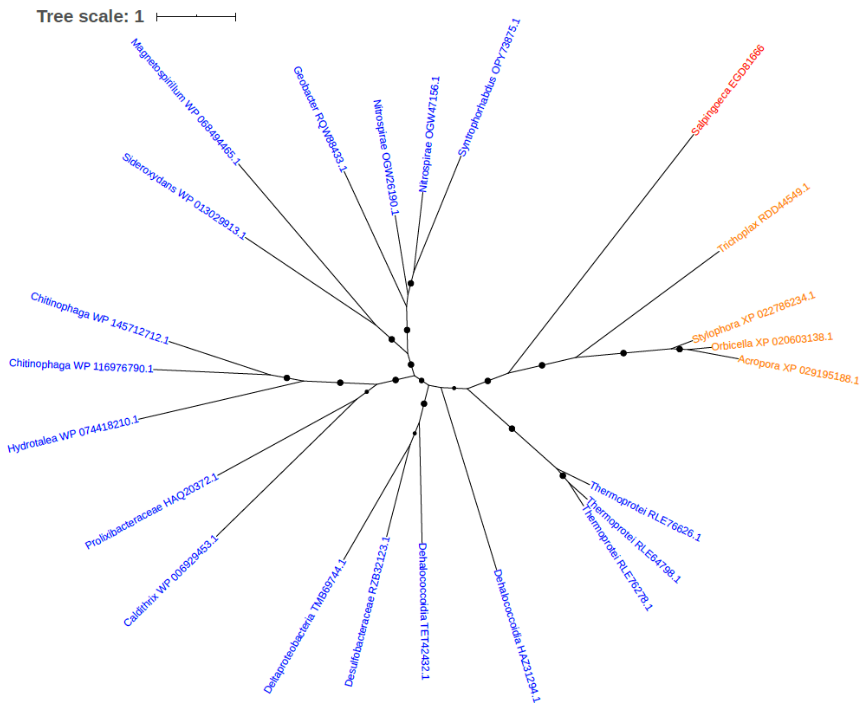


Figure S229. Phylogeny of uncharacterized protein (EGD81666)

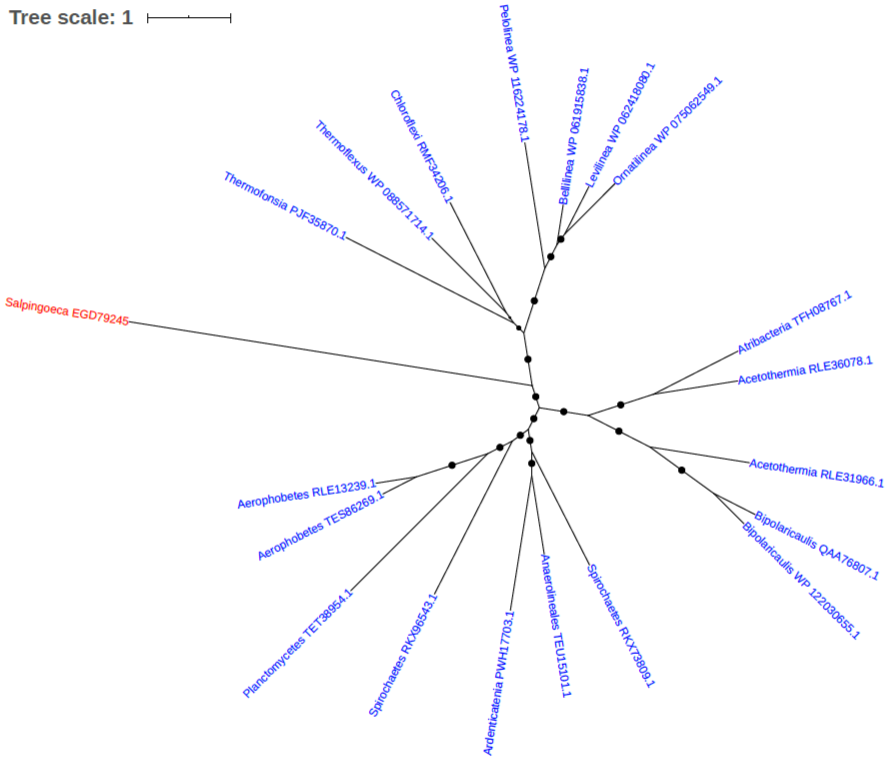


Figure S230. Phylogeny of uncharacterized protein (EGD79245)

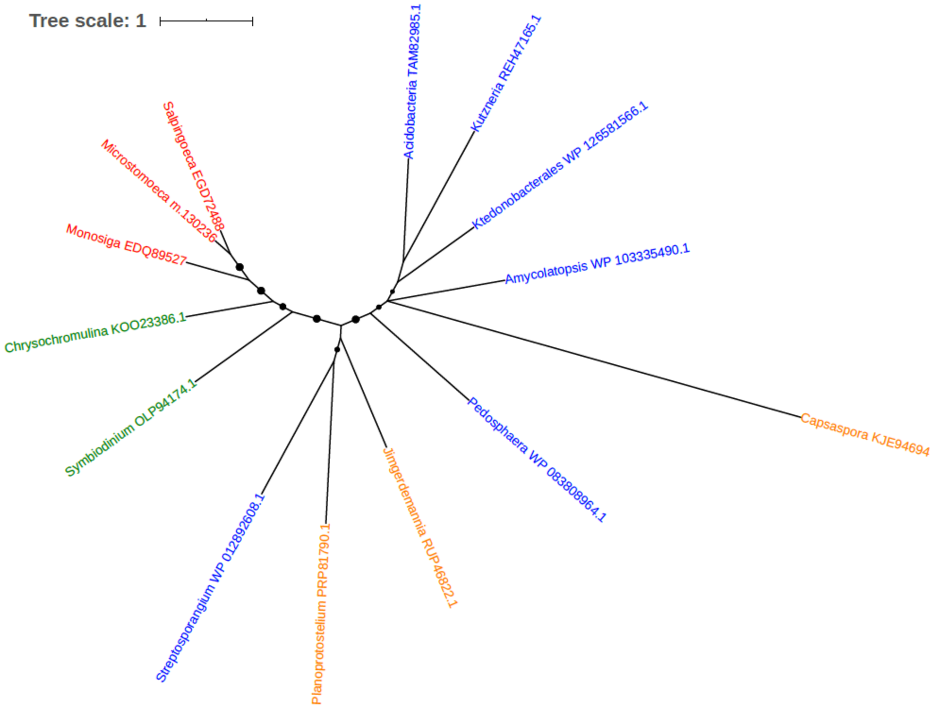


Figure S231. Phylogeny of uncharacterized protein (EGD72488)

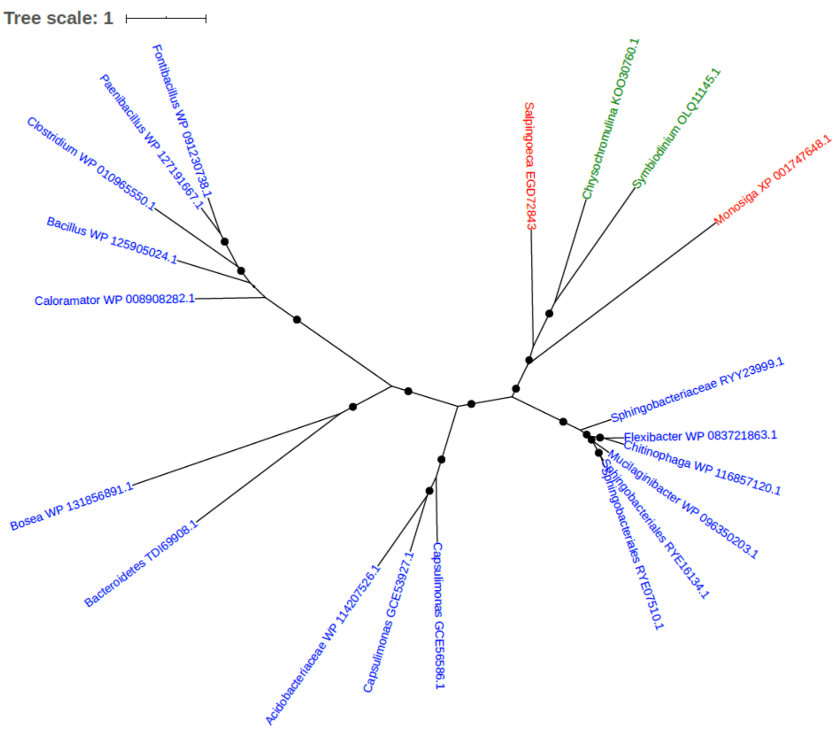


Figure S232. Phylogeny of uncharacterized protein (EGD72843)

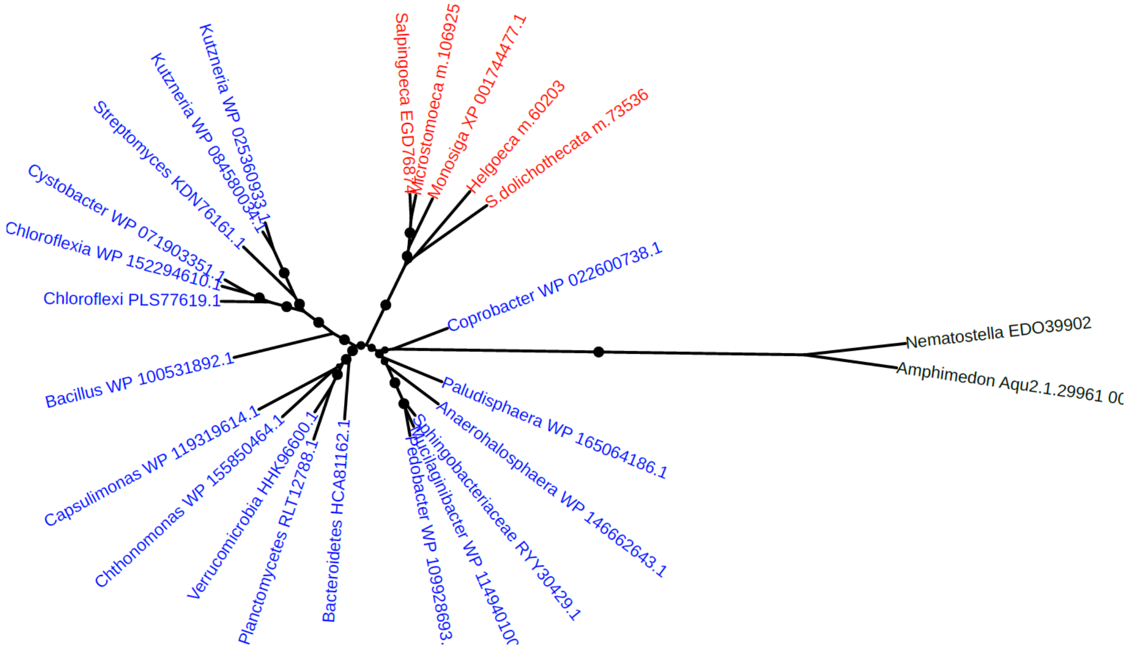


Figure S233. Phylogeny of uncharacterized protein (EGD76874)

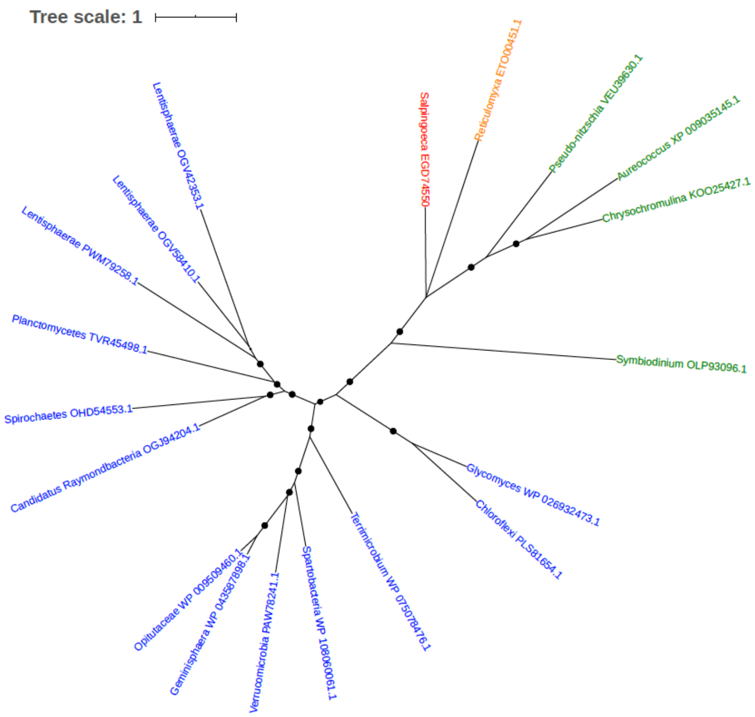


Figure S234. Phylogeny of uncharacterized protein (EGD74550)

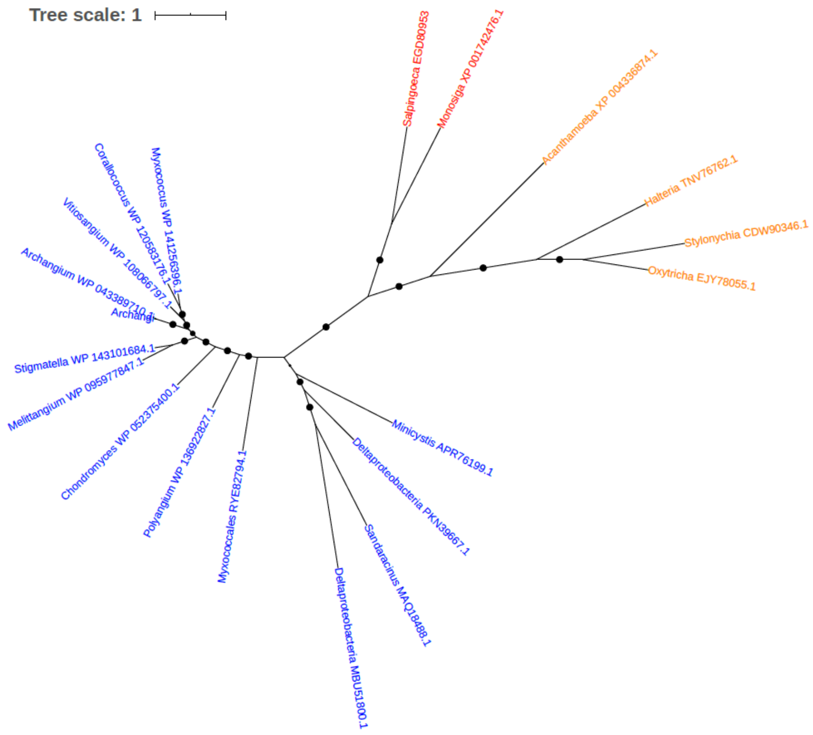


Figure S235. Phylogeny of uncharacterized protein (EGD80953)

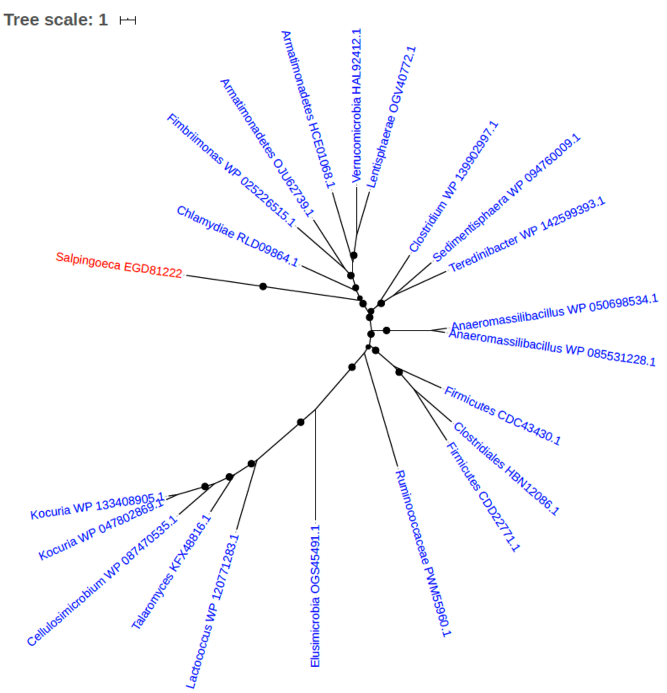


Figure S236. Phylogeny of uncharacterized protein (EGD81222)

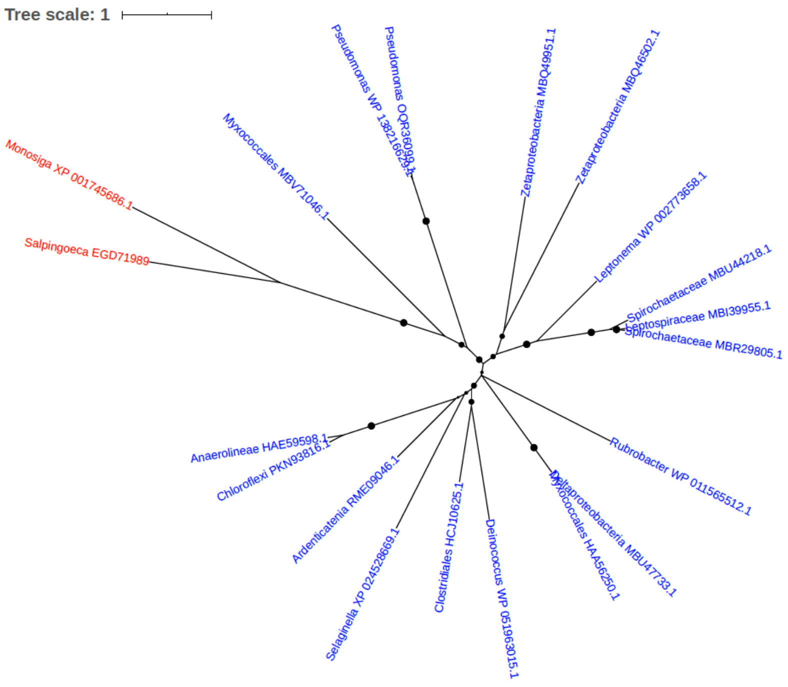


Figure S237. Phylogeny of uncharacterized protein (EGD71989)

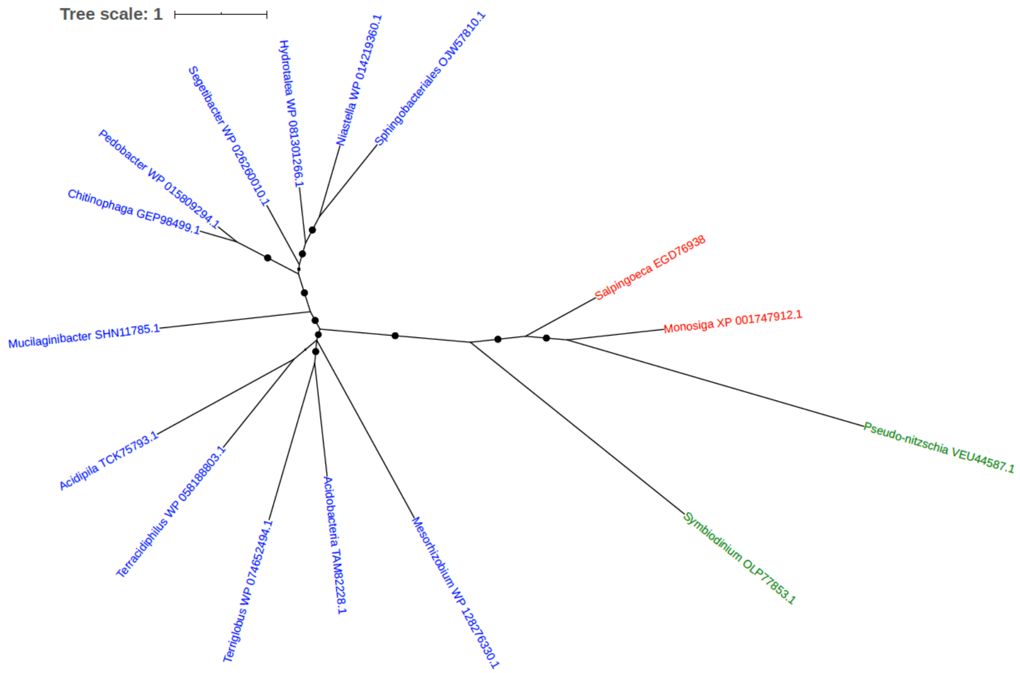


Figure S238. Phylogeny of uncharacterized protein (EGD76938)

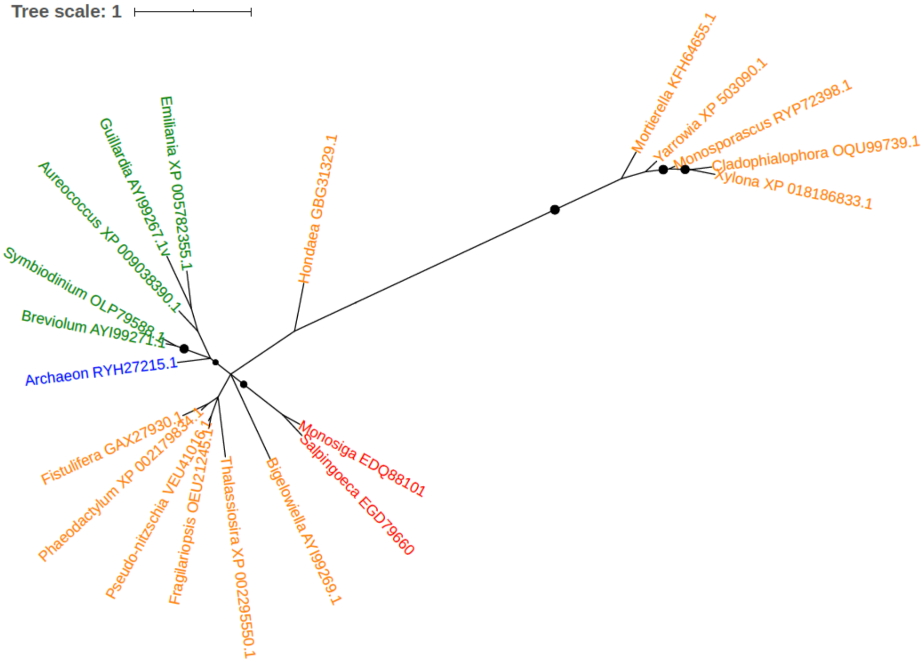


Figure S239. Phylogeny of uncharacterized protein (EGD79660)

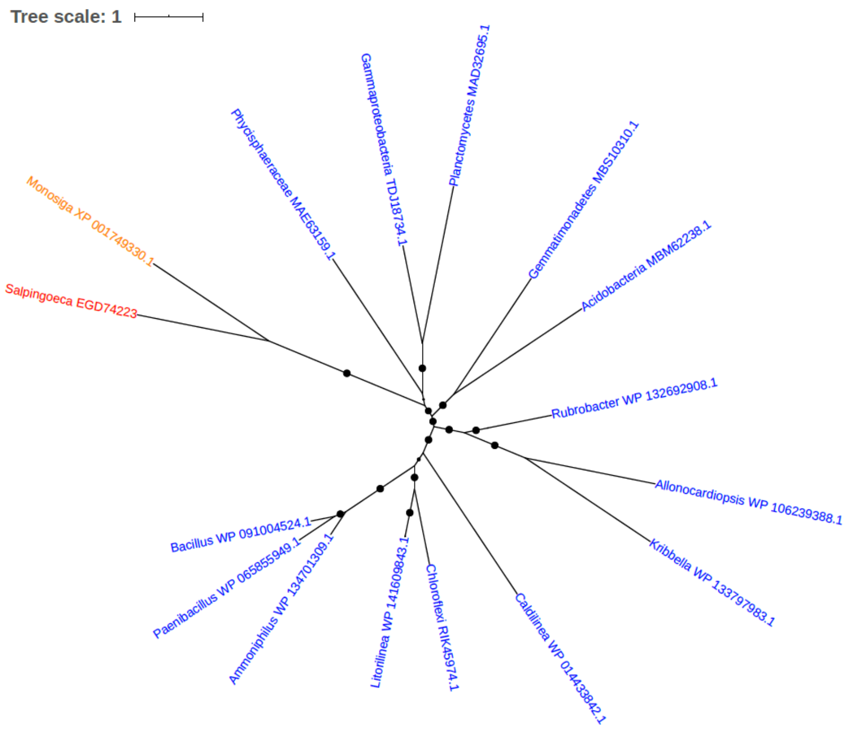


Figure S240. Phylogeny of uncharacterized protein (EGD74223)

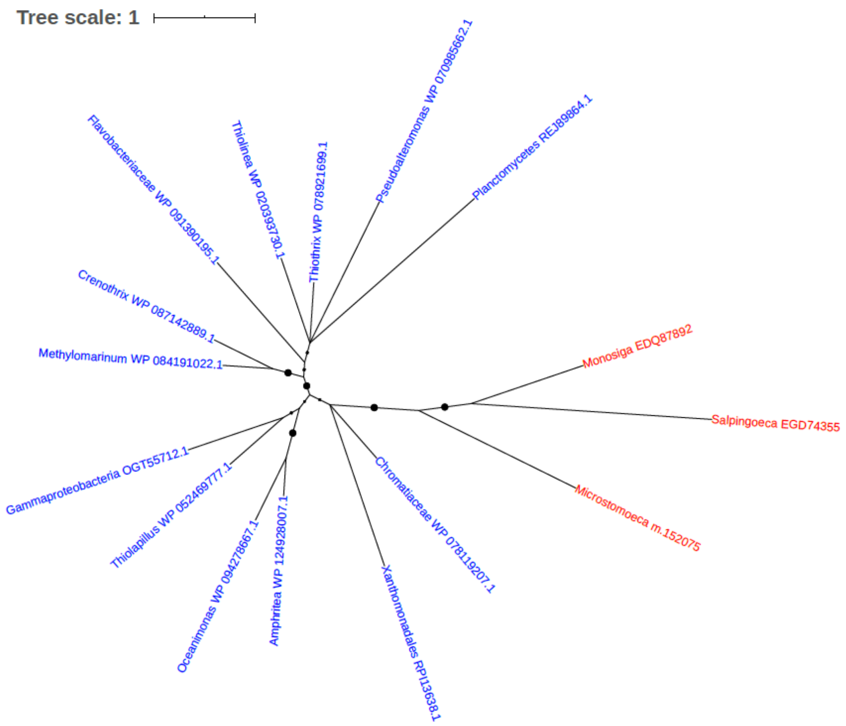


Figure S241. Phylogeny of uncharacterized protein (EGD74355)

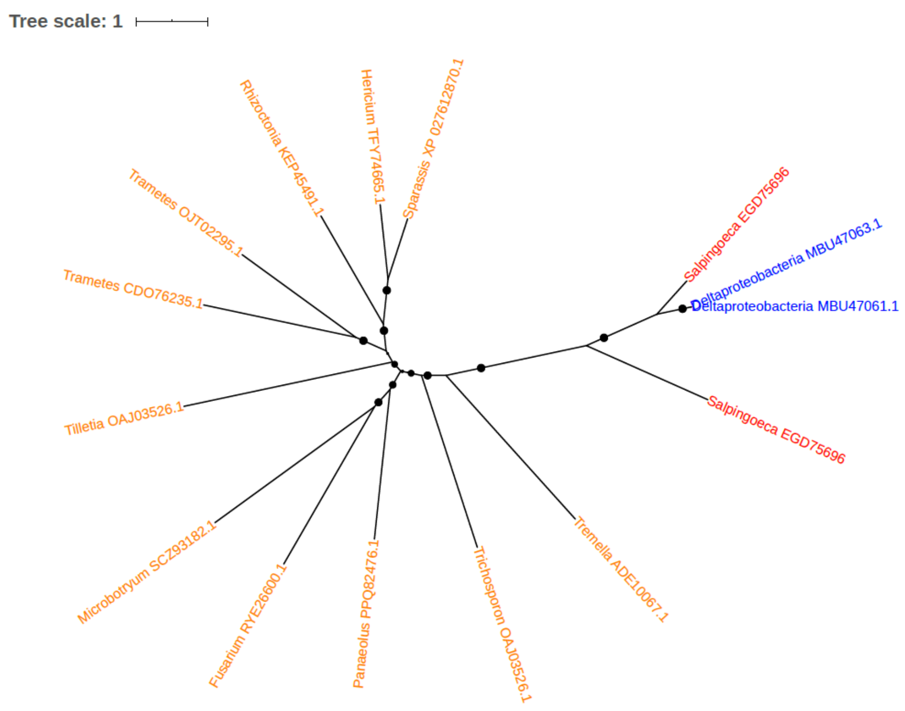


Figure S242. Phylogeny of uncharacterized protein (EGD75696)

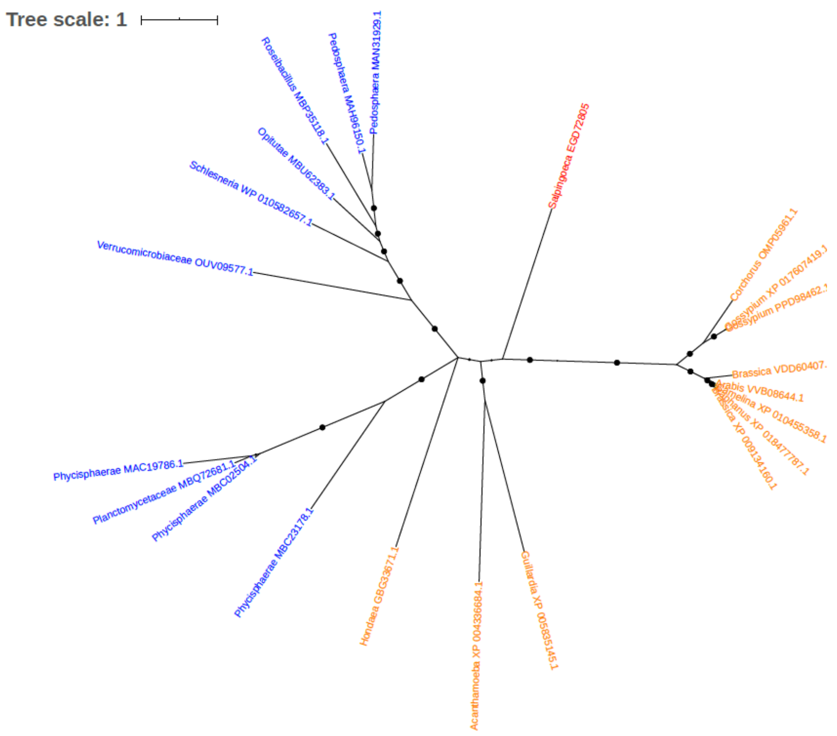


Figure S243. Phylogeny of uncharacterized protein (EGD72805)

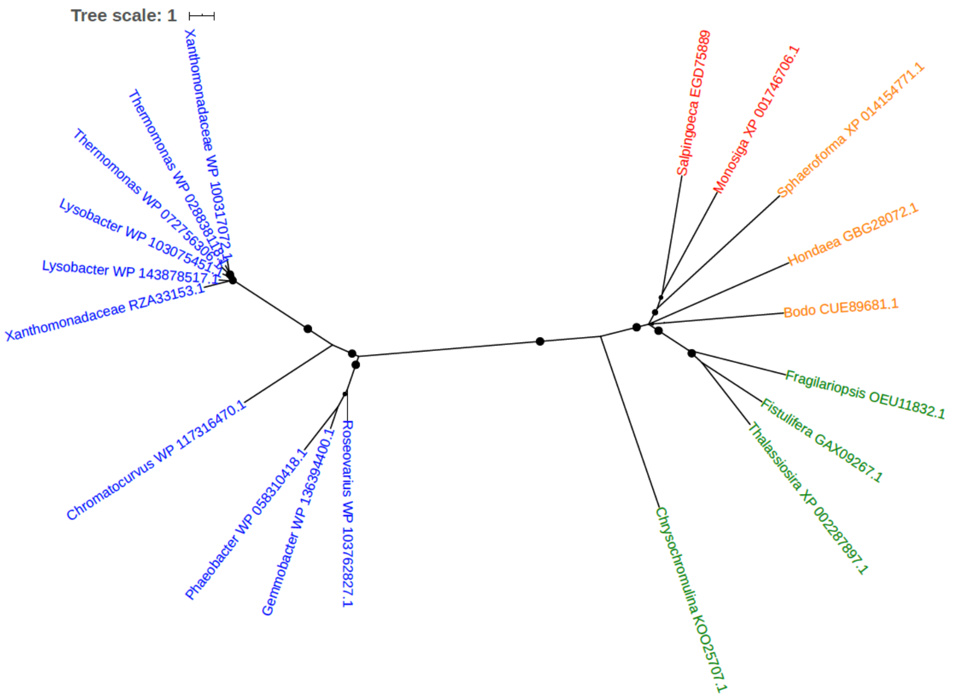


Figure S244. Phylogeny of uncharacterized protein (EGD75889)

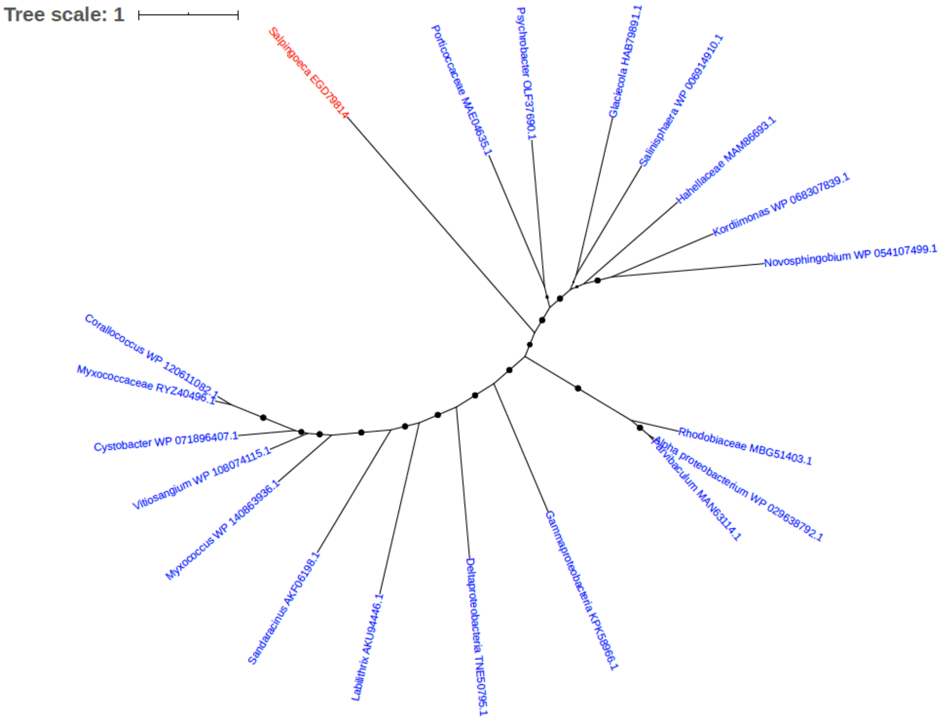


Figure S245. Phylogeny of uncharacterized protein (EGD79814)

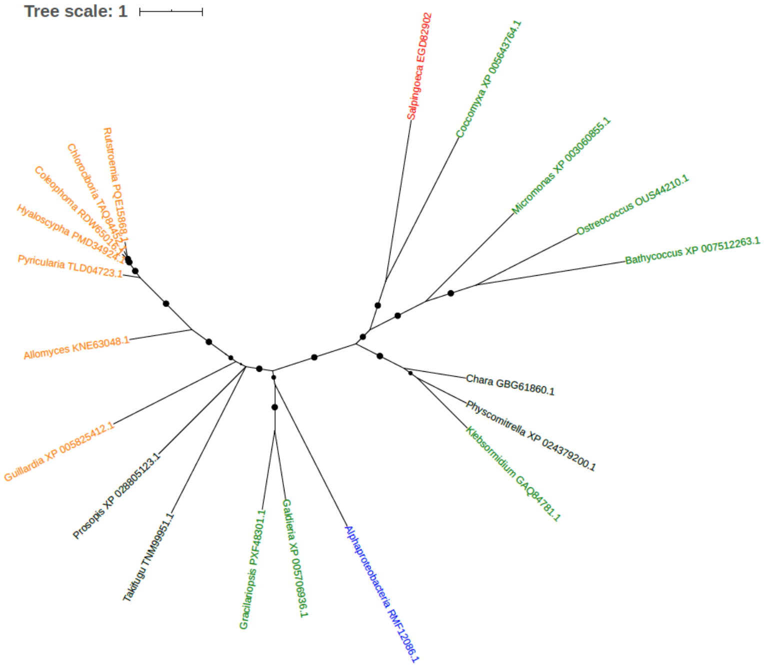


Figure S246. Phylogeny of uncharacterized protein (EGD82902)

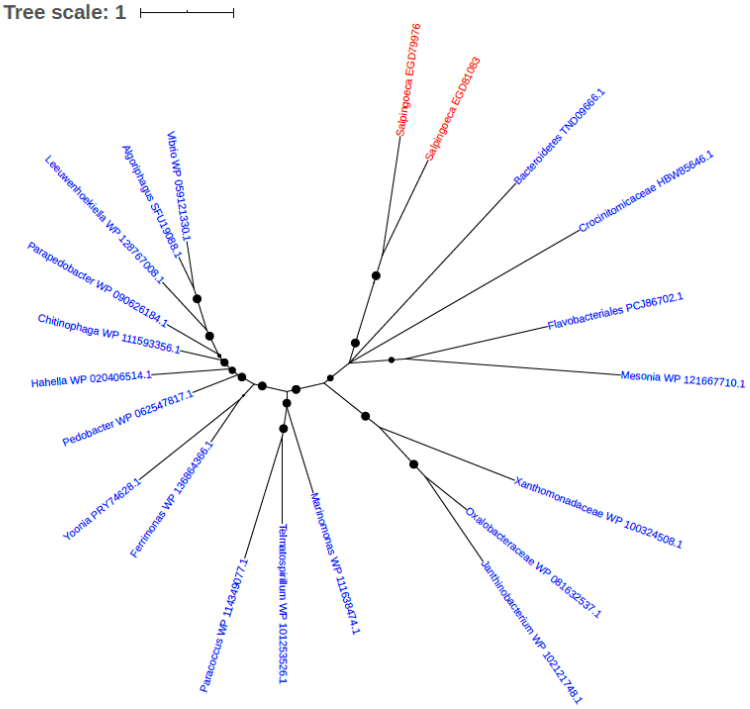


Figure S247. Phylogeny of uncharacterized protein (EGD89976, EGD81083)

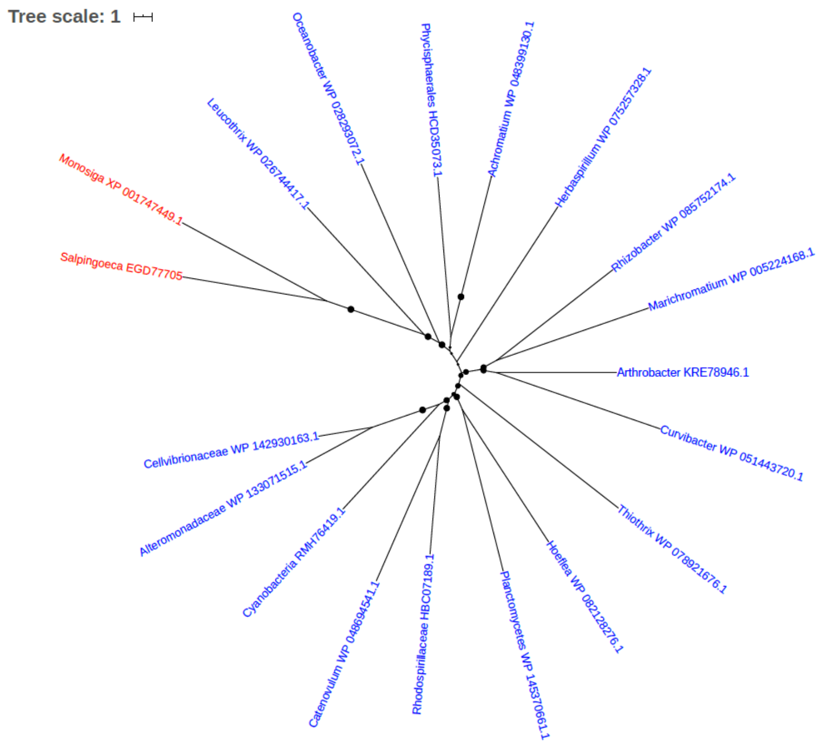


Figure S248. Phylogeny of uncharacterized protein (EGD77705)

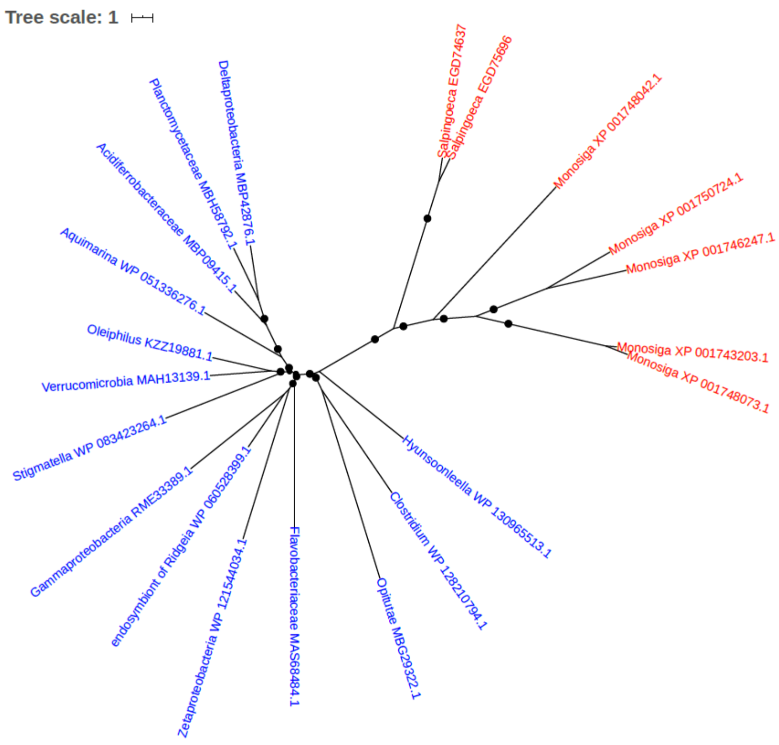


Figure S249. Phylogeny of uncharacterized protein (EGD74637, EGD75696)

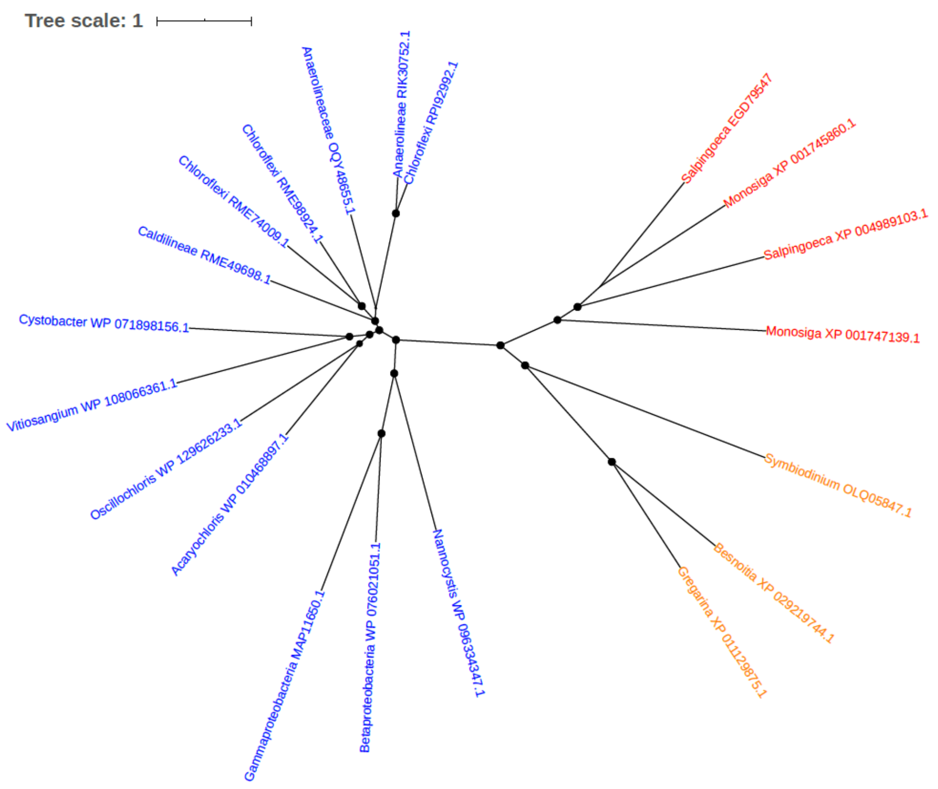


Figure S250. Phylogeny of uncharacterized protein (EGD79547)

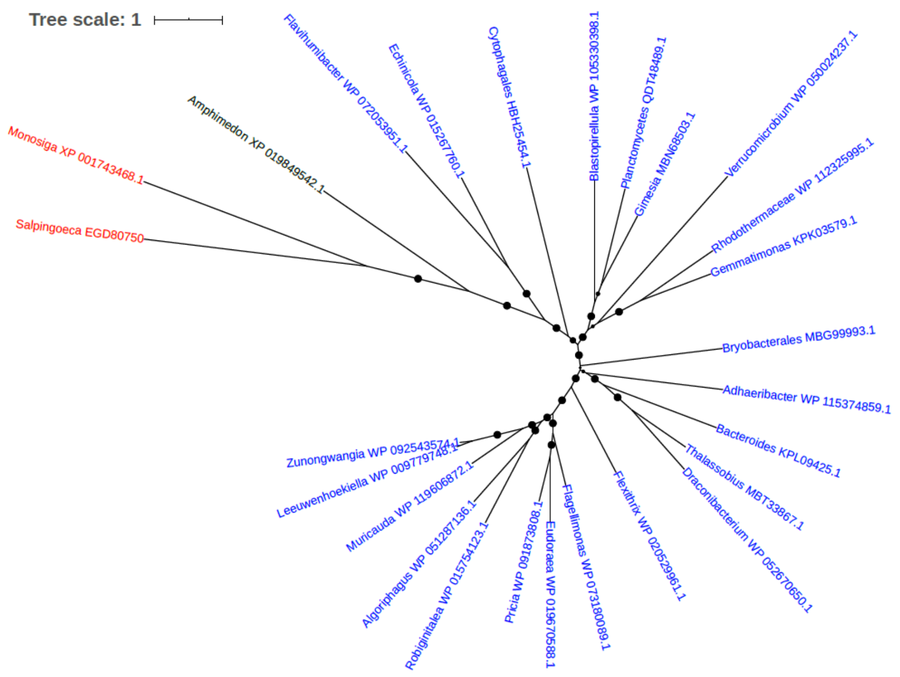


Figure S251. Phylogeny of uncharacterized protein (EGD80750)

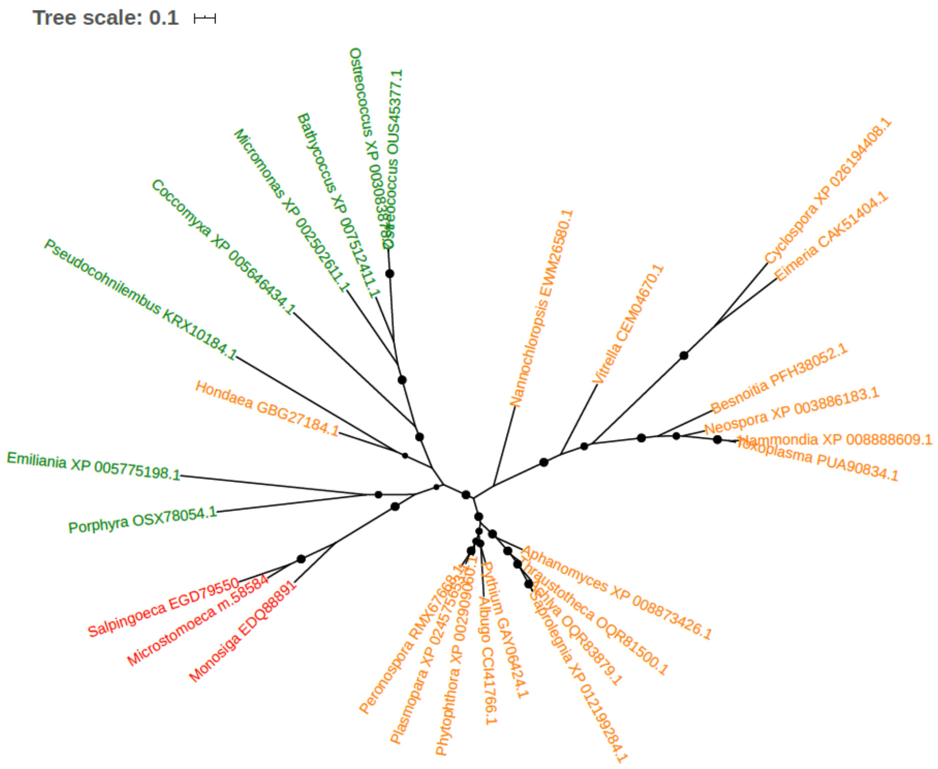


Figure S252. Phylogeny of uncharacterized protein (EGD79550)

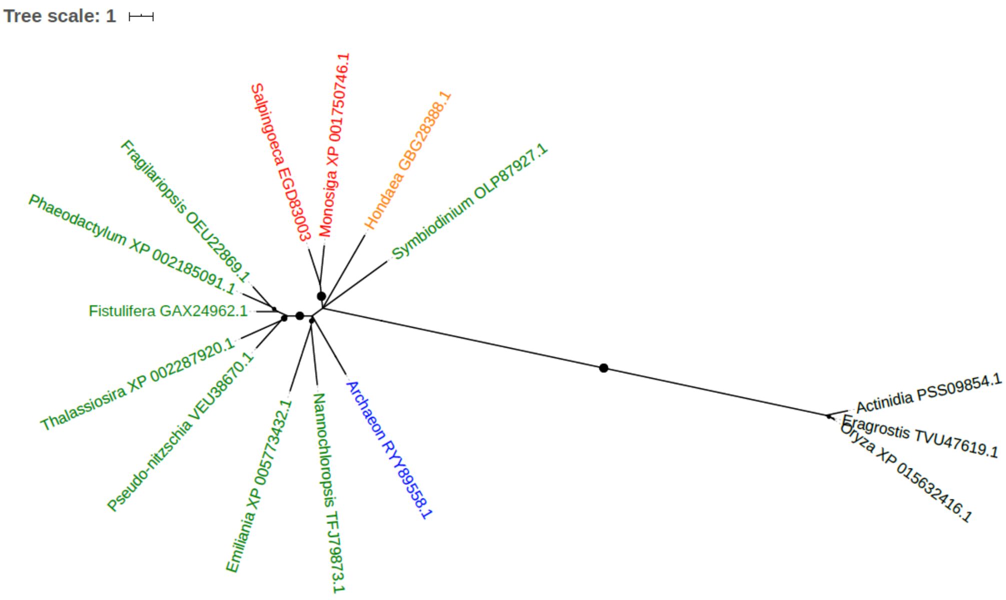


Figure S253. Phylogeny of uncharacterized protein (EGD83003)

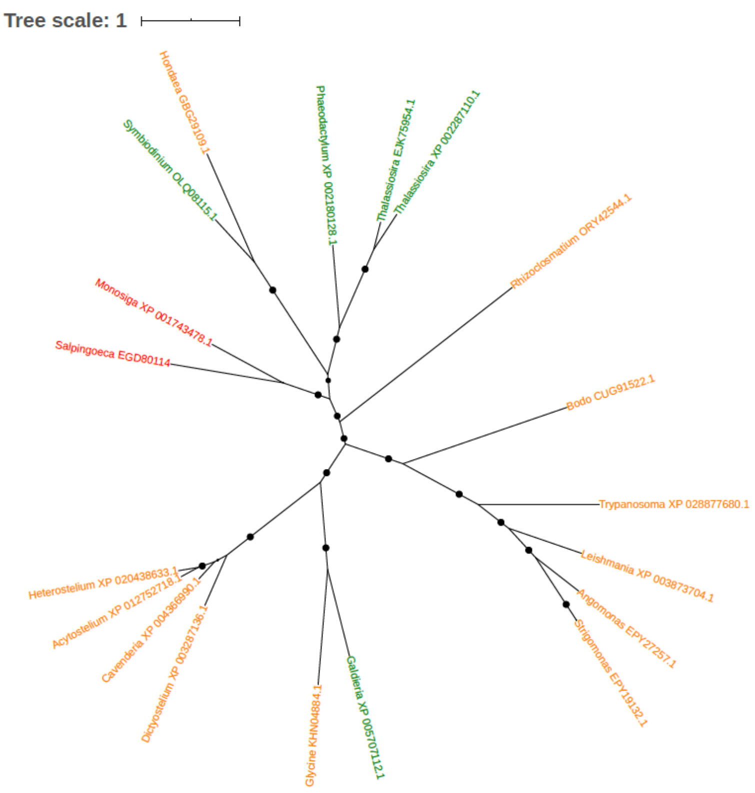


Figure S254. Phylogeny of uncharacterized protein (EGD80114)

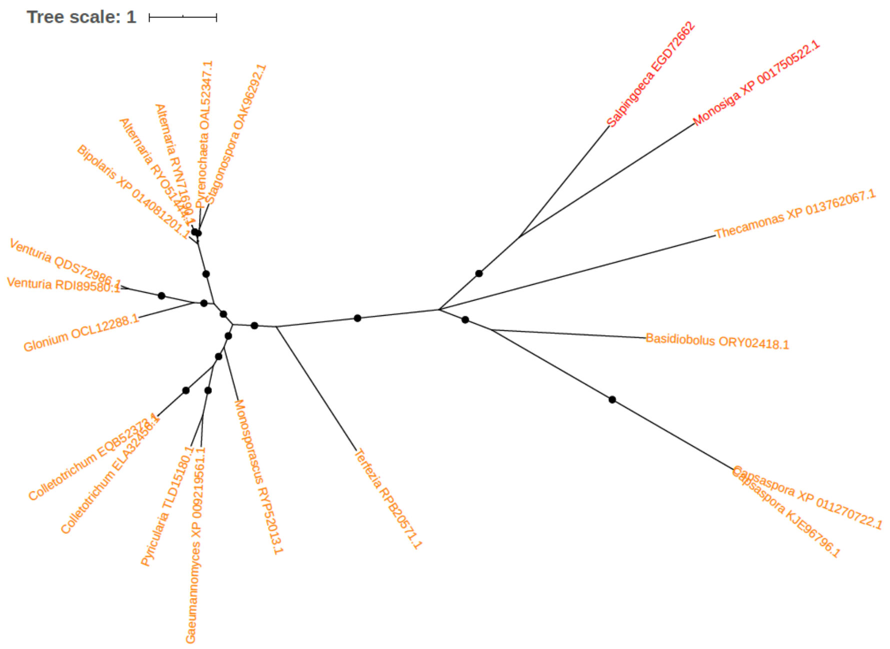


Figure S255. Phylogeny of uncharacterized protein (EGD72662)

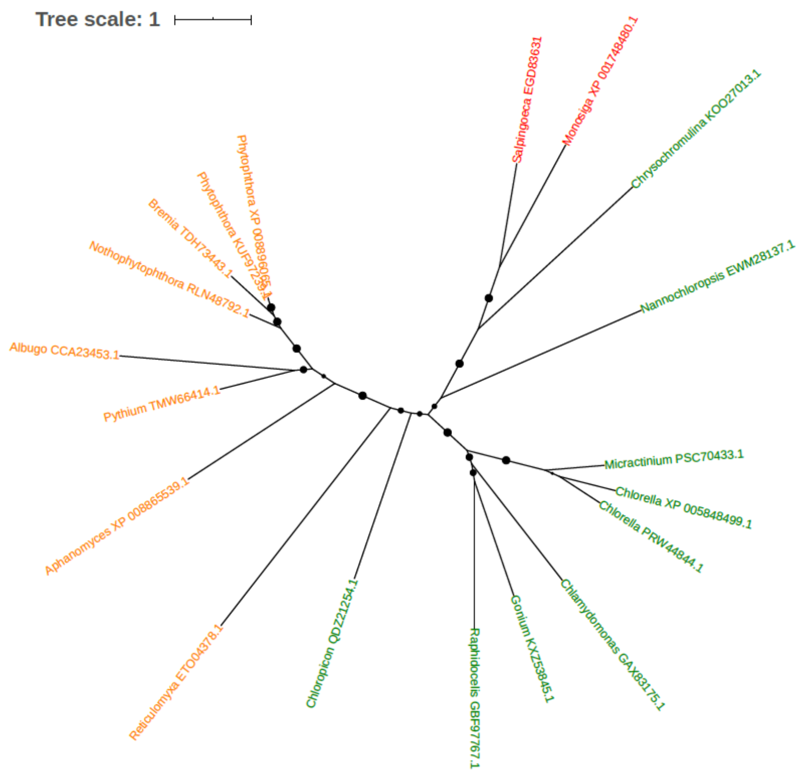


Figure S256. Phylogeny of uncharacterized protein (EGD83631)

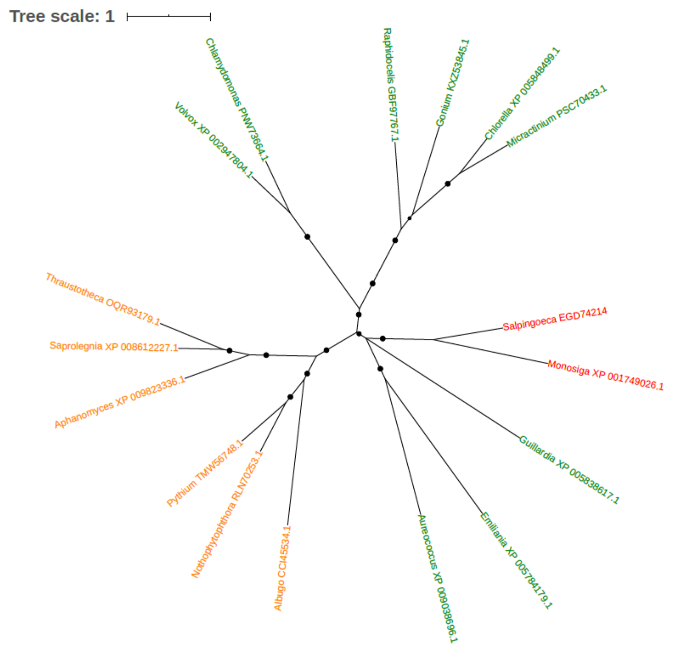


Figure S257. Phylogeny of uncharacterized protein (EGD74214)

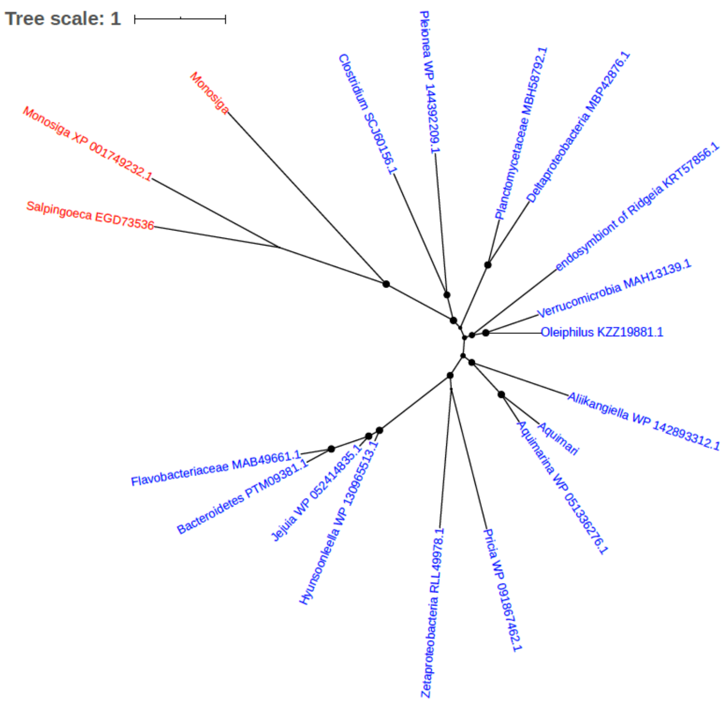


Figure S258. Phylogeny of uncharacterized protein (EGD73536)

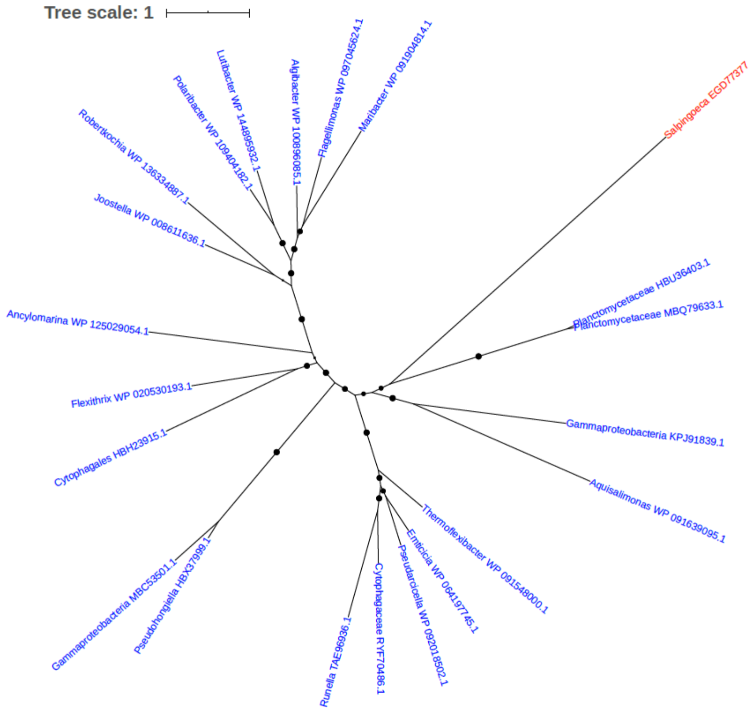


Figure S259. Phylogeny of uncharacterized protein (EGD77377)

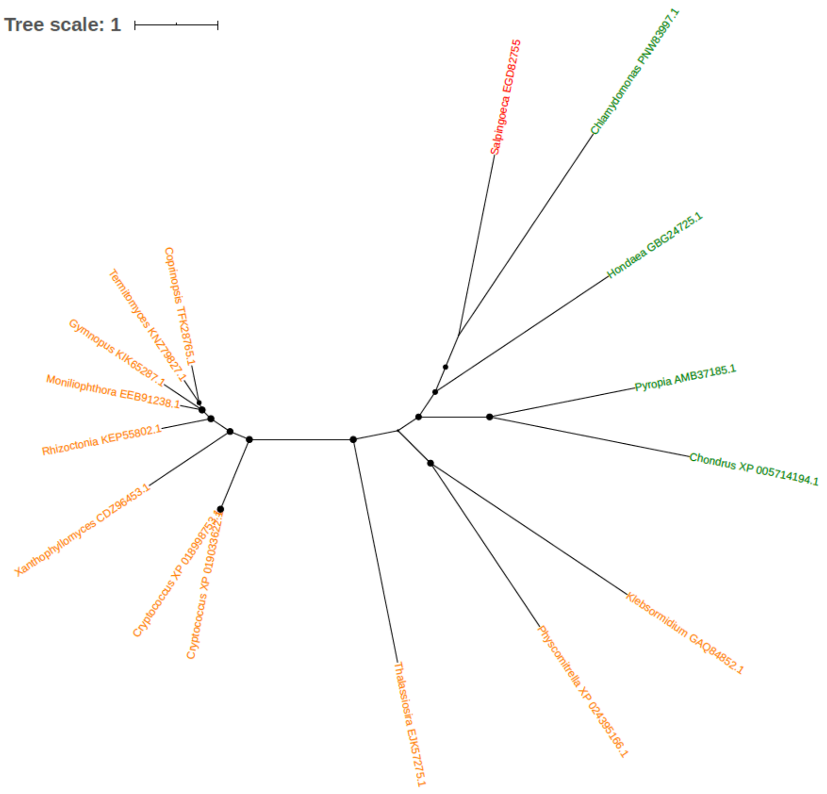


Figure S260. Phylogeny of uncharacterized protein (EGD82755)

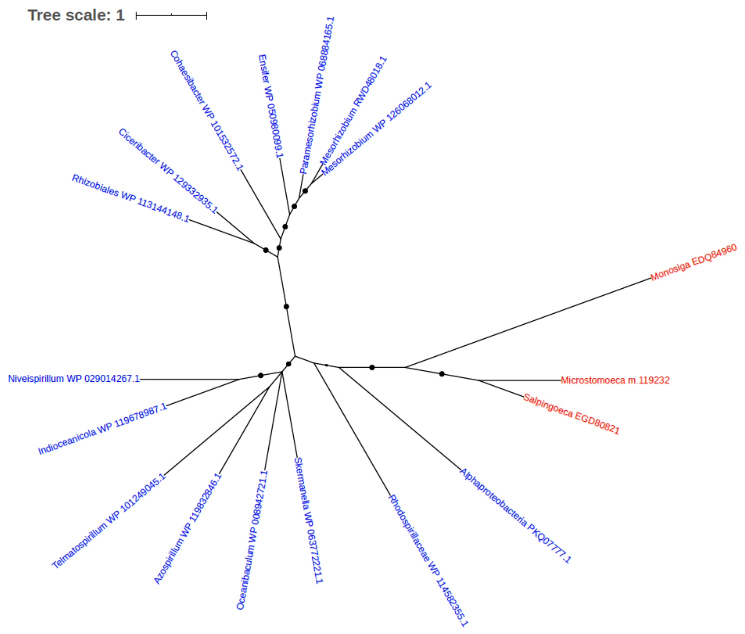


Figure S261. Phylogeny of uncharacterized protein (EGD80821)

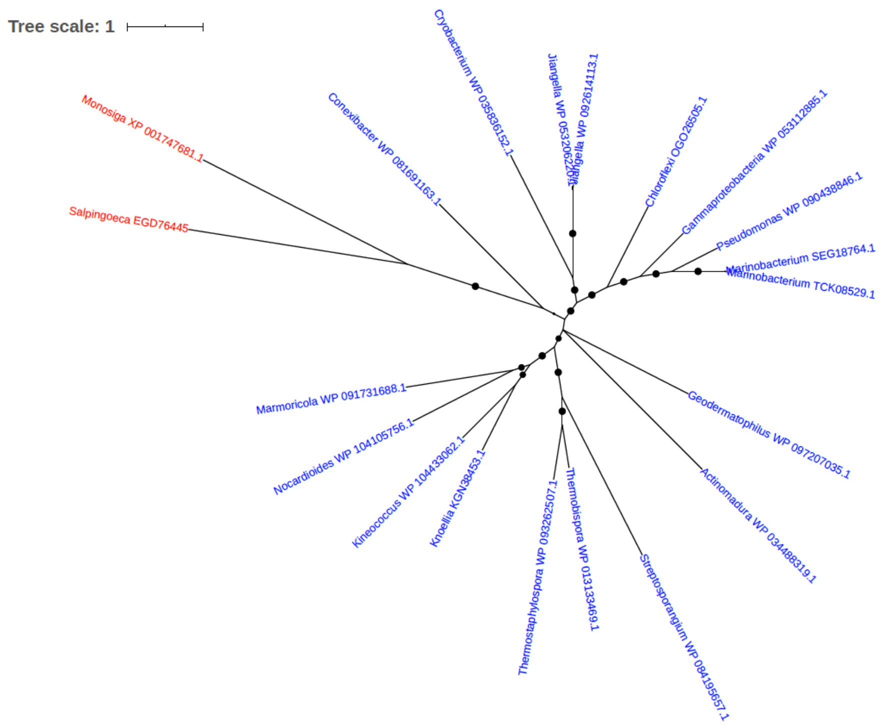

Figure S262. Phylogeny of uncharacterized protein (EGD76445)

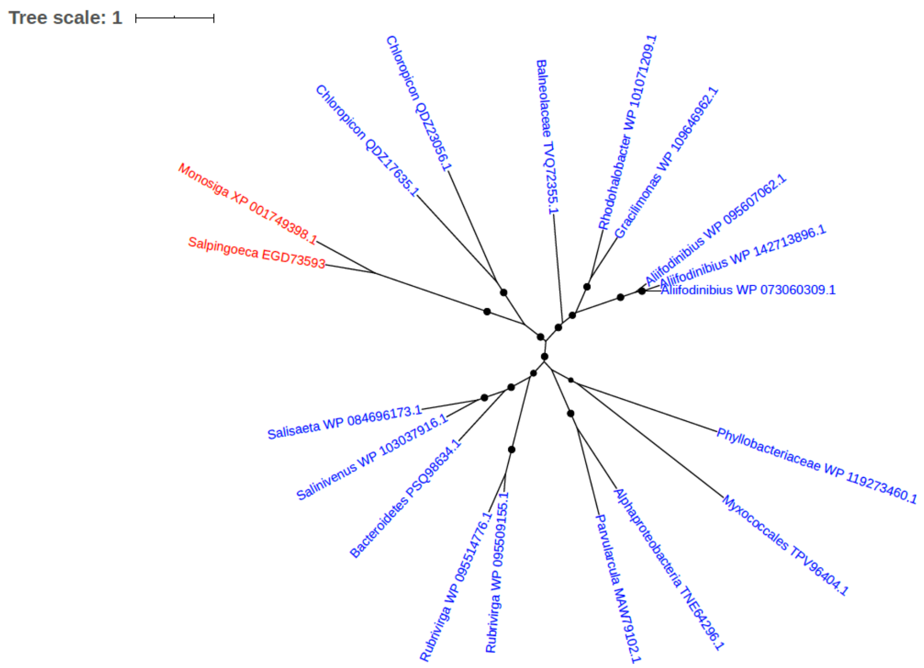

Figure S263. Phylogeny of uncharacterized protein (EGD73593)

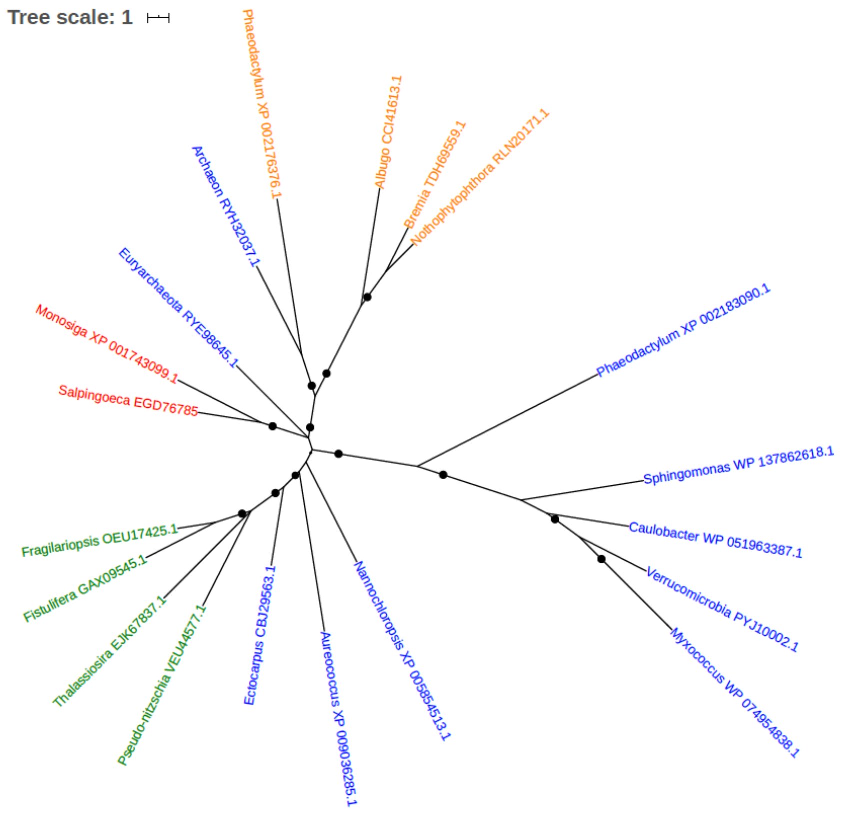


Figure S264. Phylogeny of uncharacterized protein (EGD76785)

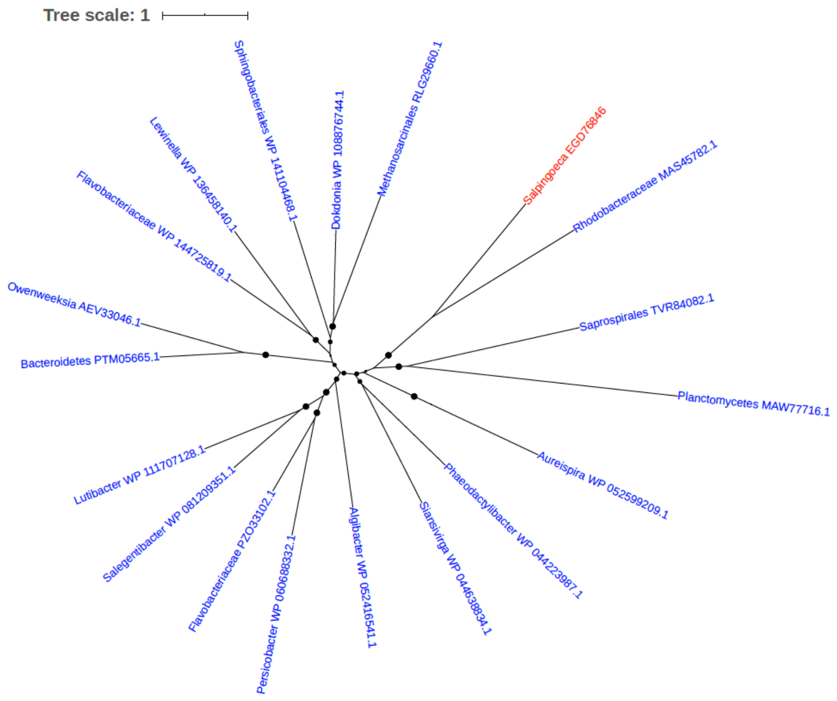


Figure S265. Phylogeny of uncharacterized protein (EGD76846)

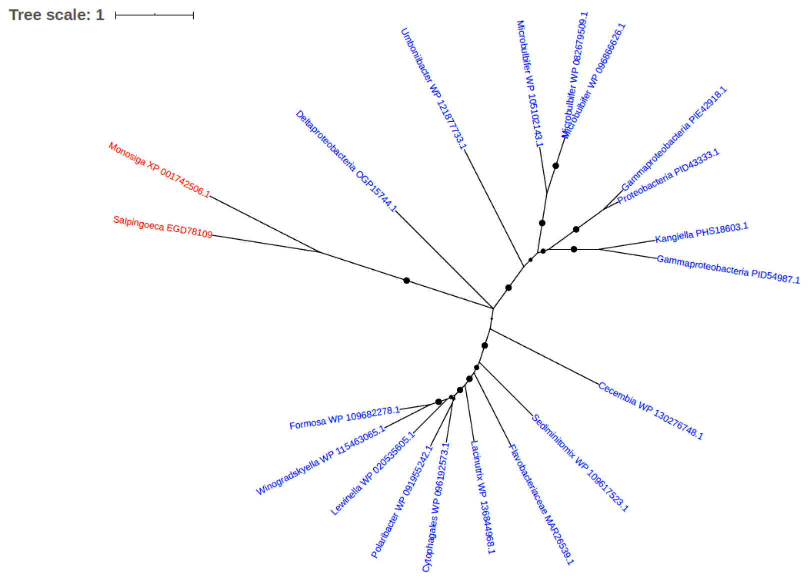


Figure S266. Phylogeny of uncharacterized protein (EGD78109)

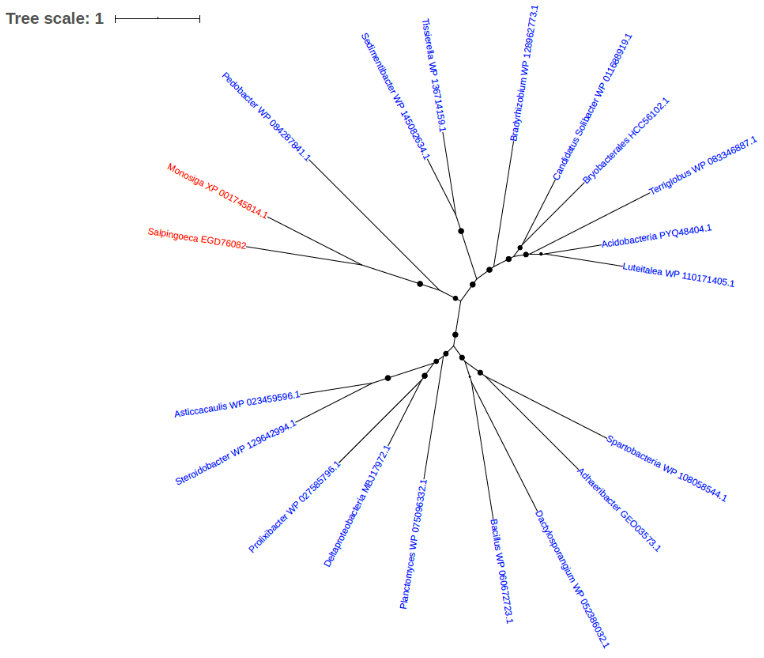


Figure S267. Phylogeny of uncharacterized protein (EGD76082)

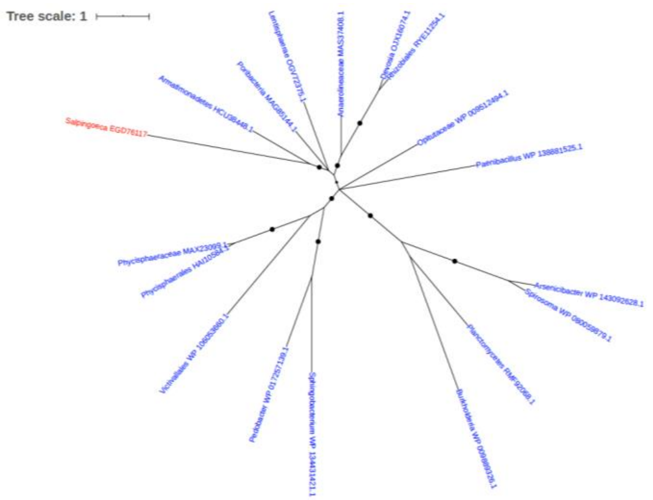


Figure S268. Phylogeny of uncharacterized protein (EGD76117)

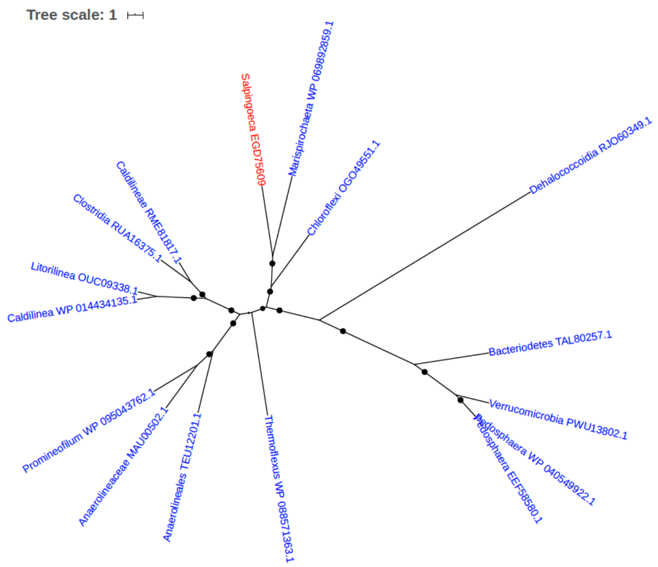


Figure S269. Phylogeny of uncharacterized protein (EGD75609)

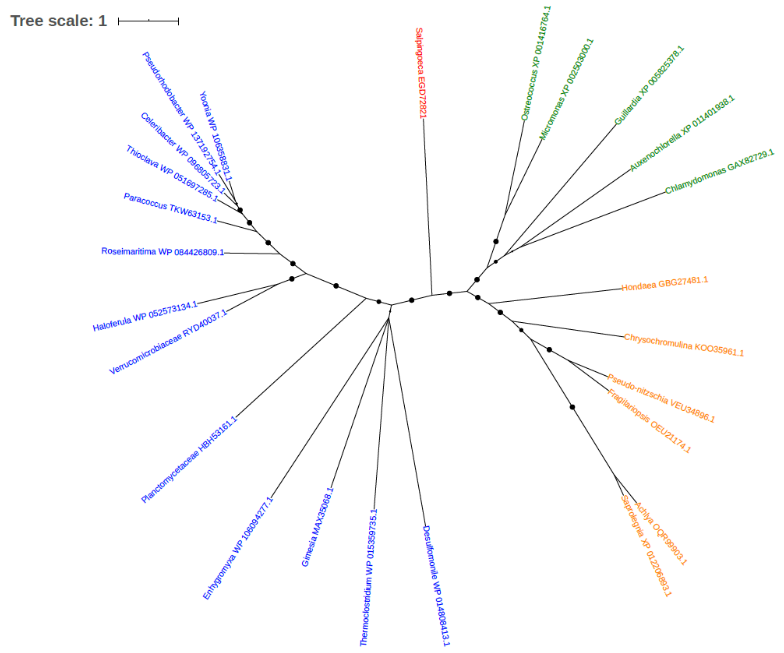


Figure S270. Phylogeny of uncharacterized protein (EGD72821)

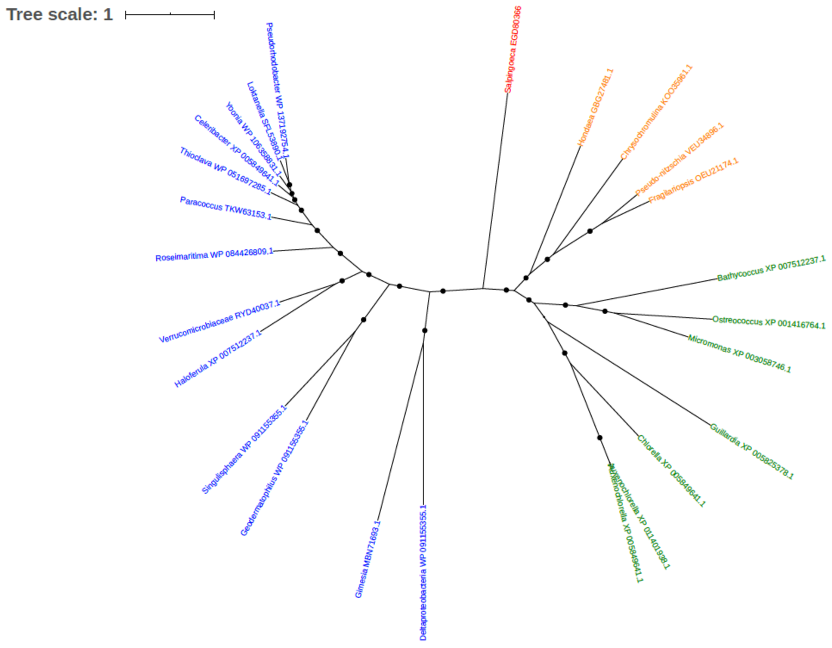


Figure S271. Phylogeny of uncharacterized protein (EGD80366)

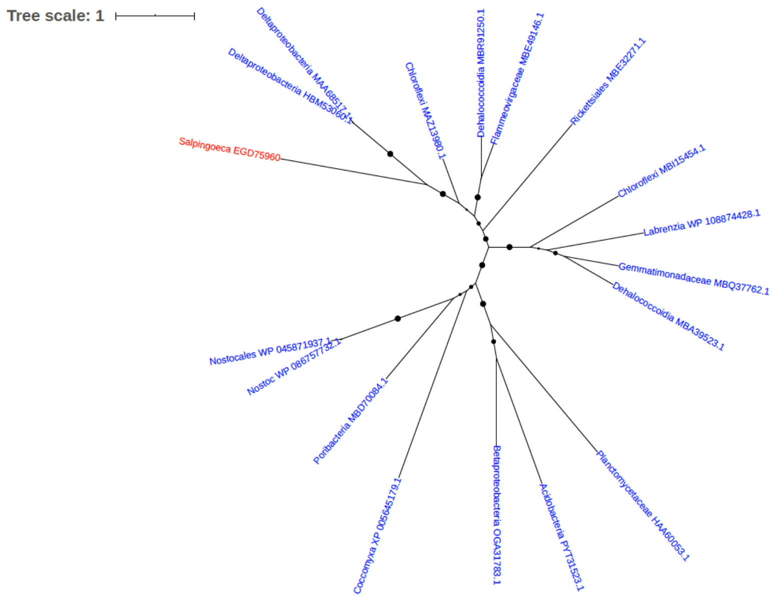


Figure S272. Phylogeny of uncharacterized protein (EGD75960)

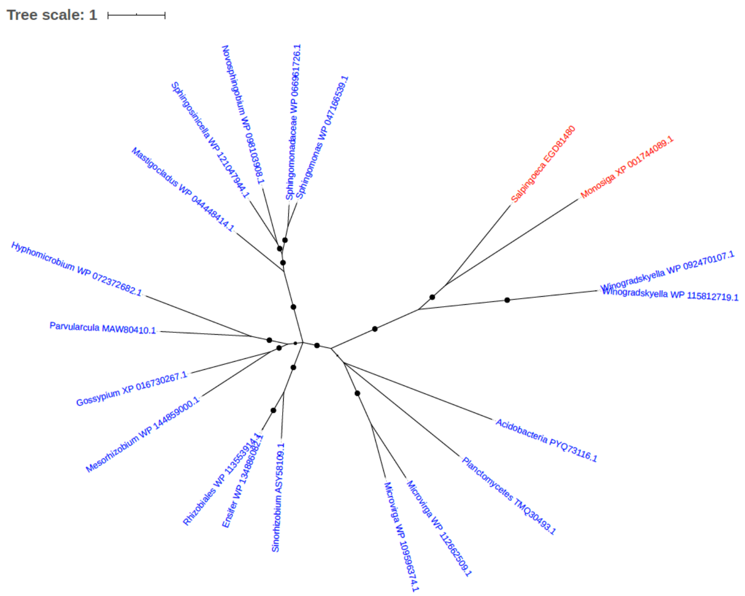


Figure S273. Phylogeny of uncharacterized protein (EGD81480)

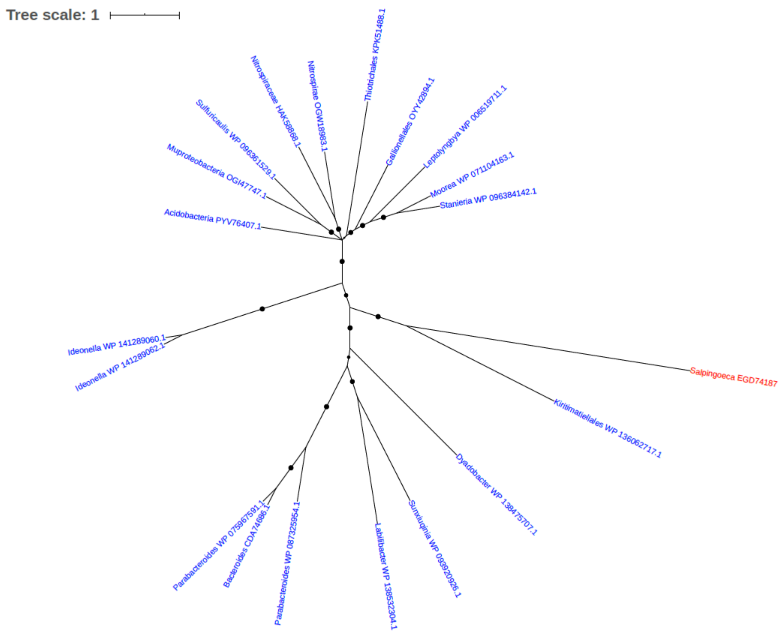


Figure S274. Phylogeny of uncharacterized protein (EGD74187)

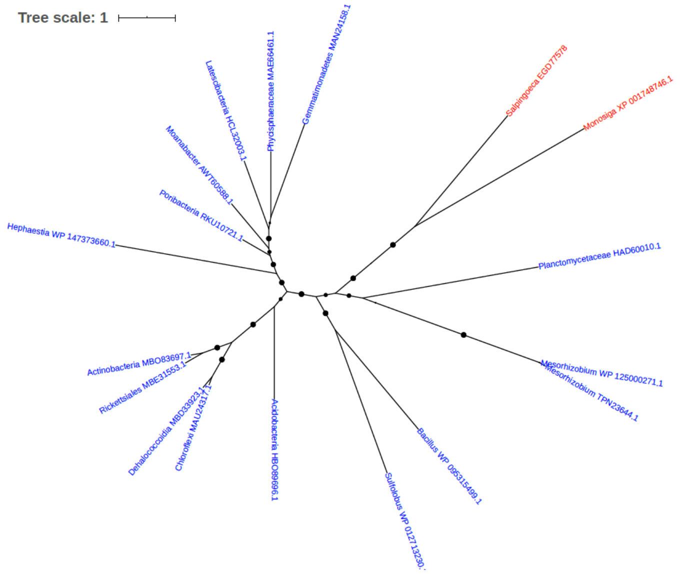


Figure S275. Phylogeny of uncharacterized protein (EGD77578)

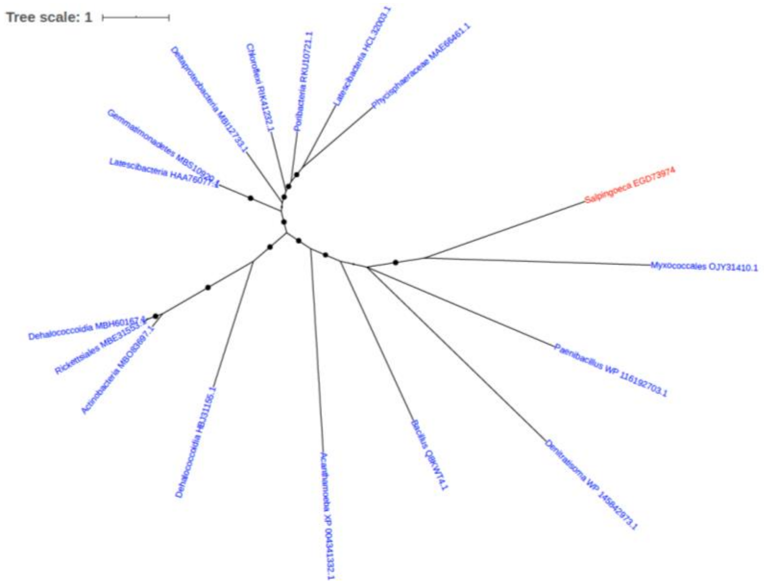


Figure S276. Phylogeny of uncharacterized protein (EGD73974)
