## Supplementary figures and images for "Detection of Horizontal Gene Transfer in the Genome of the Choanoflagellate *Salpingoeca rosetta*"

### Additional File 2

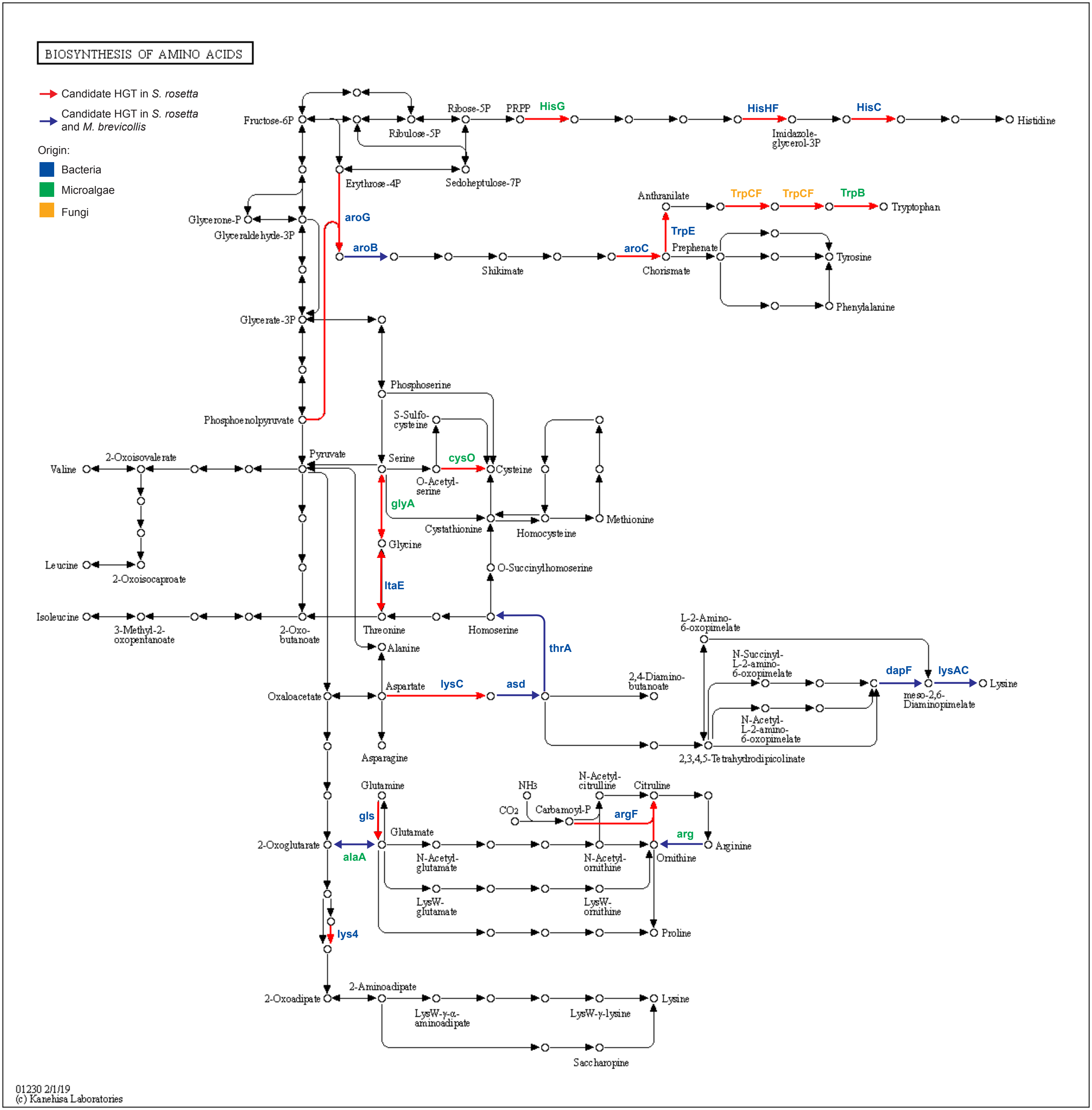
